# “The Green Band”: Using Production and Catch to Judge Distortive Pressure on an Ecosystem

**DOI:** 10.1101/2025.06.09.658403

**Authors:** E.A. Fulton, K. Sainsbury, C. Bulman, C. Novaglio, J. Porobic, D. Hayes, E.I. van Putten, L.X.C Dutra, L. Thomas, W. Norris, C. Pert, A. Willock, S.C. Montenegro, L. Garay-Narvaez, S. Hernández Concha, A. Hernández Saso, K.S. Mohamed, T.V. Sathianandan, S. Kuriakose, E. Varghese, T.M. Najmudeen, S. Vasudevan, M. Anish, S.S. Salim, I. Mandro, S. Zador, I. Ortiz, A. Whitehouse, K. Aydin, W. Tweit, D. Evans

## Abstract

Ecosystem based fisheries management is an increasingly accepted approach for many fisheries management agencies. Indicators and methods that can usefully support the aspiration of making ecosystem based fisheries management decisions on tactical time frames are slowly accumulating, but have primarily focused on biomass status and trends. We first verify that fishing can distort the cumulative distribution of connections within an ecosystem, and then show (via theoretical testing and application to four contrasting case study locations) how relating catch with production by defining an acceptable band of exploitation pressure – the “Green Band” – can help inform fisheries decision makers as to whether an ecosystem is under distortive patterns of fishing pressure that will ultimately undermine the structure of the ecosystem.

## Introduction

As more management agencies have committed to an ecosystem approach to fisheries (Garcia et al., 2003) or ecosystem-based fisheries management (EBFM; Pikitch et al., 2004) authorities, fishing industries and the scientific community have struggled with finding straightforward ways of defining how it can be achieved. Progress has been made (Patrick and Link, 2015), but one of the core challenges remaining in delivering EBFM is to find a means of pragmatically implementing the approach in a way that is feasible for the diversity of fisheries that operate in any one geographic region, nation, or ultimately the world. This management approach needs indicators and tools that can help deliver the science content in a form that is directly useful for management decision making.

Over the past two decades significant effort has been dedicated to finding suitable indicators that provide useful information on which to base ecosystem-based management decisions. While much of that effort has focused on ecological rather than social or economic considerations (Hornborg et al., 2019), the majority of those ecological indicators are directed towards biomasses and other aspects of status of an ecosystem (Link, 2005; Greenstreet and Rogers, 2006; Shin et al., 2010; Shin et al., 2018). In terms of indicators that give insight into ecosystem structure (and, by inference function), many of the most successful structural indicators have been indirect – such as those derived from biomass ratios (Fulton et al., 2005; Jennings, 2005; Link, 2005; Fogarty, 2014).

Over the past decade there has been a concerted effort to consider fundamentally different approaches to harvesting at the ecosystem scale, their compatibility with EBFM and the implications for ecosystem structure and status (Garcia *et al*., 2012; Froese *et al*., 2016; Burgess et al. 2016, Rehren and Gascuel, 2020; Law and Plank, 2023). A key aspect of this discourse has been debate around how to actually undertake ecosystem-based management – i.e. how to implement the alternative management approaches, how to provide practical guidance on the scale of the typical (annual) management cycle regarding whether fishing pressure is sustainable or not from an ecosystem perspective, and how to pragmatically changeover from single species management to EBFM.

The Lenfest Oceans Program and CSIRO Australia supported an international working group that explored potential indicators and guidelines for practical EBFM during 2017 and 2021. This paper reports on one aspect of that work. Building off network theory and work begun in Scotland (Heath et al., 2017) in the context of the ‘balanced harvest’ fisheries debate (Jacobsen *et al.,* 2014; Burgess *et al*., 2016; Froese *et al*., 2016; Burgess and Plank, 2020; Rehren and Gascuel, 2020; Law and Plank, 2023, *Sun et al*., 2023), the work summarised in this paper outlines how estimates of production and yield from a fishery can be used to provide guidance on whether a fishery is applying distortive pressure on ecosystem structure that will ultimately undermine its sustainability at an ecosystem level. The paper will first lay out the theoretical premise of the underlying concepts and the procedure for estimating the “Green Band” of fisheries’ operations, and assessing the pressure on an ecosystem’s structure. The methods will then be demonstrated using information and models from four case study regions.

### Network Premise

Networks – nodes and the edges (links) between them – are used to describe many aspects of the natural (e.g. food webs) and anthropogenic (e.g. electricity grids) world. Over the course of the last fifty years theoreticians have developed methods for analysing network structure and have identified a suite of well understood network properties. In particular, Ecological Network Analysis, a branch of network ecology (Borrett and Lau, 2014), highlights how the number and density of nodes in a system and the patterns of connections within it can be used to summarise the structural complexity of a network and the relative contribution of sub-webs to the overall system (Lau et al., 2017).

The similarity of methods and lessons learnt, regarding network properties, from across multiple disciplines highlights the universal nature of some network properties, including how their structure can influence resilience. While Ecological Network Analysis has had a rich academic history spanning many years (e.g. Dunne et al., 2002; Milo et al., 2002; Montoya and Sol, 2002; Johnson et al., 2003; Dunne et al., 2004; Jordan et al., 2006; Fath et al., 2007; D’Alelio et al., 2016; Marina et al., 2018; Fath et al., 2019; Niquil et al., 2020), it has not been applied to fisheries and conservation problems until recently (e.g. Navia et al., 2016; McDonald-Madden et al., 2015). This is unfortunate given the potential of the field.

For example, Erdös and Rényi (1960) showed that the structural properties of networks change above and below threshold points. Given fisheries change the relative species composition of an ecosystem, they will be changing the underlying weighted network of the ecosystem (i.e. the flows through the ecosystem), which means that they may also be changing fundamental ecosystem network properties.

Theoretical ecology has explored how the shape of the cumulative distribution of connections within a network changes as an ecosystem is put under increasing pressure (seminal papers can be found in Newman et al., 2006). This work has found that as the pressure from a stressor distorts the content of the network defining the ecosystem’s connections, it degraded fundamental network properties – such as aggregate connectivity – and this can be expressed in the power law coefficient of the system’s distribution of connections. Metrics of this type have been used to understand the dynamics of small world networks (Watts and Strogatz, 1998; Dunne et al., 2002; Navia et al., 2016). Whether or not food webs are small world networks, they appear to have many of the same network properties (Marina et al, 2018) and so the methods should be transferable.

Different forms of networks – such as the different classes of small world networks (Newman, 2005) – are distinguished by the shape of the cumulative distribution of their node connectivity, particularly the tail of that distribution. In exploring the potential for network theory to form a basis for EBFM metrics that focus on structural ecosystem properties, it is worth trying to understand whether varying degrees of fishing pressure change the nature of the network as expressed by the coefficients of their cumulative distribution of connections. Once it has been confirmed that the coefficients are informative with regard to maintenance of system structure, they can be used to verify whether more easily applied approaches – such as the “Green Band” approach described below – are a useful way to assess the pressure on an ecosystem and thereby guide management to achieve ecosystem structural sustainability within EBFM. The first section of the results presented here lays out our attempt to use network analysis metrics to verify the Green Band approach really does track whether the structure of a system is being distorted by fishing pressure.

Logically, an ecosystem is capable of sustaining the pressure (predation) profile of an unfished system – represented by the biomass-productivity profile and judged using the cumulative distribution of connections from network analysis. Pressure applied in anything other than this pattern will lead to potential distortions in the system structure and flows (as explored further below). The challenge for EBFM is to find a reasonable means of judging whether or not the pattern of pressure being applied to an ecosystem is distortive or not. In exploring the potential application of the balanced harvesting approach in Scottish fisheries Heath et al (2017) proposed a method for judging how balanced exploitation in a fishery was. While we are not engaging in the debate on the appropriateness of balanced harvesting as a practical multispecies fishing strategy, we saw potential in the method they put forward as a practical means of verifying whether realised fishing pressures were putting distortive pressure on the marine ecosystem - thereby providing EBFM guidance. For reasons that will become clear below, we term our approach the “Green Band”. We briefly describe the approach in the methods section below, before providing some example applications of the approach.

## Methods

### Network Analysis

The influence of fishing pressure on ecosystem network structure was explored by first running the Ecopath with Ecosim (EwE; Christensen et al., 2000) model of south eastern Australia (Bulman et al., 2006) under a wide range of fishing scenarios:

- unfished
- 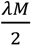 simultaneously for all exploited species where M is the average annual rate of natural mortality and λ ∼ [0.1, 0.2, 0.5, 0.65, 1.0, 1.5, 2.0, 5.0, 10.0]
- γFMMSY where γ ∼ [0.1, 0.2, 0.5, 0.65, 1.0, 1.5, 2.0, 5.0, 10.0, 20.0] and FMMSY is the multispecies maximum sustainable yield point (with full compensation, which means the rest of the system responds to changes in the fished species) estimated using the inbuilt EwE routine (Christensen et al., 2000).
- F = 𝑃^1+𝑎^ (i.e. consistent with the Green Band described below)
- historical fishing levels (also known as the base model run, this is the model run with the time series the model was originally fit to; this simulation was used to see how the results of the model are for realistic fished system patterns compared to those for more theoretical patterns)

All the simulations, except for the base model, were run for 100 years to ensure they reached equilibrium (or close, in some cases some of the slowest turnover species had not quite reached complete equilibrium by completion of the 100-year period). The base model simulation was halted at the end of the historical period.

The matrix defining the ecosystem network at the end of each run (*t*=100 years) was calculated with the weighted edges (diet connections between each predator-prey combination at time *t*; *Li,j,t*) calculated as:

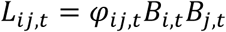

where *Bi,t* is the biomass of prey *i* at time *t*; *Bj,t* is the biomass of predator *j* at time *t*; and *φi,j,t* is the proportion of the diet of predator *j* made up of prey *i* at time *t*.

The cumulative distribution of 𝐿_𝑖j,100_was then calculated from each network using the *degree.distribution*() routine of the *igraph* R package (Csardi and Nepusz, 2006) and curve fitting scripts executed using the *nls*() (Baty et al., 2015) routine of R (version 4.0.2; R core Team 2021). There is an extensive body of literature from theoretical ecology on the shape of cumulative distributions of connections in networks and how they can be represented with power laws (Newman et al., 2006; Marina et al, 2018). Consequently, the candidate models considered were linear and power curves (both scaled and unscaled, and power with and without a tail), as described in Table 1. The best fitting curve was selected using AIC - calculated using the *AICcmodavg* (Mazerolle, 2020), *plyr* (Wickham, 2011) and *stringr* R packages (Wickham, 2021). The shape of these best fit distributions was compared across fishing scenarios. All scripts used to do analyses or generate plots in this paper are available from https://github.com/eafulton/GreenBand.git

**Table 1:**
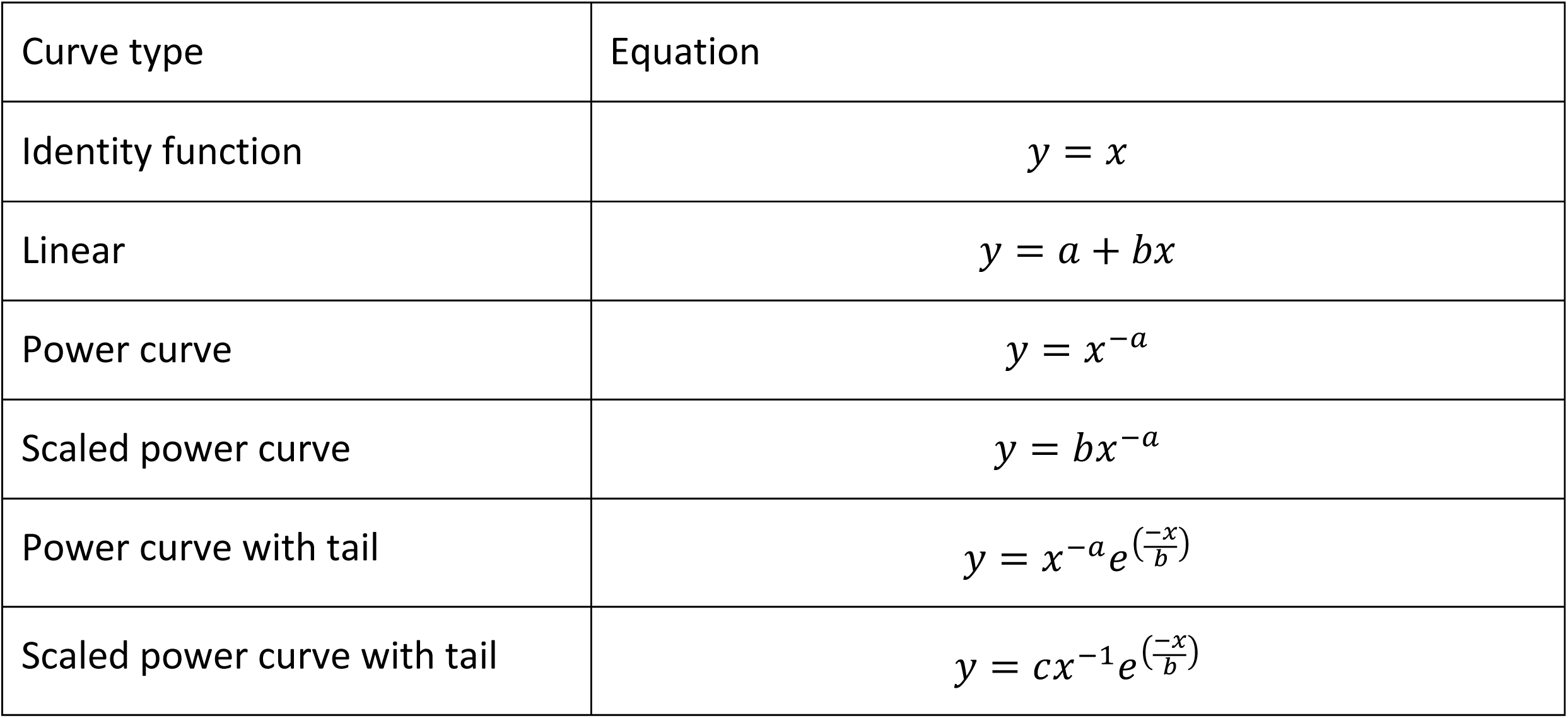
Curve forms considered during fitting of the form of the cumulative distribution of connections from the south eastern EwE model fishing scenarios.

| Curve type | Equation |
| --- | --- |
| Identity function | $y = x$ |
| Linear | $y = a + bx$ |
| Power curve | $y = x^{-a}$ |
| Scaled power curve | $y = bx^{-a}$ |
| Power curve with tail | $y = x^{-a}e^{\left(\frac{-x}{b}\right)}$ |
| Scaled power curve with tail | $y = cx^{-1}e^{\left(\frac{-x}{b}\right)}$ |

*“Green Band” Procedure*

Understanding how fishing may reshape a network (by placing differential, distortive, pressure across a network) is an important first step. However, to get management action there has to be a straightforward (and communicable) means of identifying whether distortive pressure is being applied and what can be done to intervene and rectify that situation. This is why we explored the rule of the Green Band approach. This procedure is summarised in Figure 1, but in brief:

**Figure 1:**
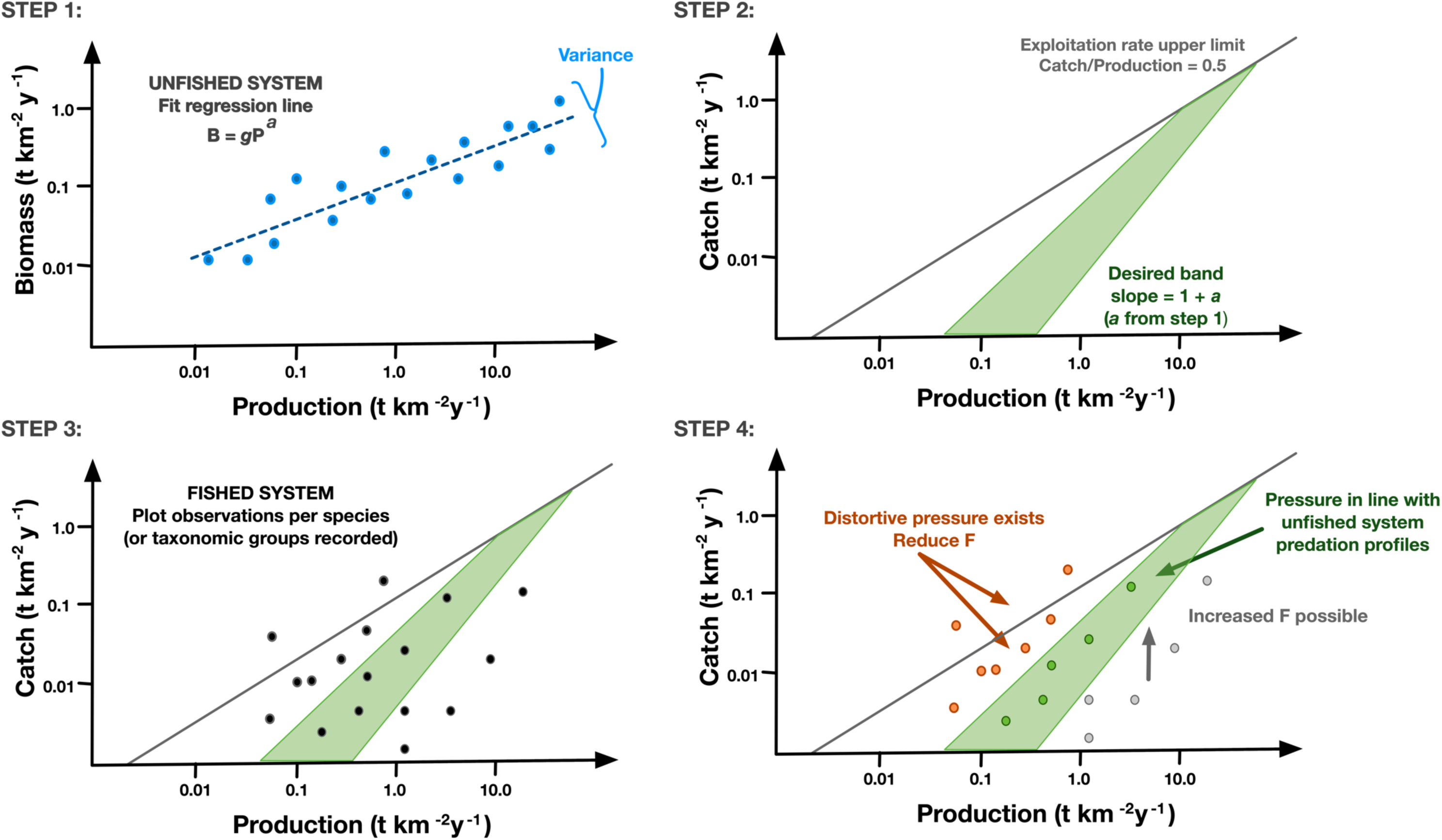
Schematic of Green Band procedure

**Step 1 – unfished profile:** In log-log space calculate the linear regression between biomass and production per species (or functional group, depending on the taxonomic resolution of the model) for the unfished ecosystem state. These relationships reflect the fact that (i) species with higher production rates support higher mortality rates, (ii) production rates scale with body size due to the size-dependency of many processes in marine ecosystems (Andersen et al 2016; Law et al. 2016). Note that the use of an unfished system is important as the intent is to determine the pressure profile the ecosystem has evolved to support in the absence of human exploitation.

**Step 2 – defining the Green Band:** Use the slope of the regression line in step 1 to define the “acceptable” pressure levels in the fished system using a plot of catch vs production (*P*). The left bound of the “acceptable” pressure levels in the fished system is given by:

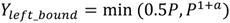

where a is the slope of the regression line in step 1. The right bound is given by:

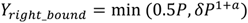

where *δ* is a scalar based on the variance in the points around the regression line in step 1 (i.e. spread of points either side) and typically has a value of approximately 0.01.

**Step 3 – plot species catch vs production:** In log-log space plot catch versus production per species (or functional group, depending on the taxonomic resolution of the model or available data).

**Step 4 – check location vs the Green Band:** The location of the points representing the individual species (or other taxonomic groups) on the catch vs production plot is used to judge whether the fishing pressure being applied is distortive or not. Species that are located in the Green Band are being fished with a level of pressure consistent with that of the natural system (i.e. it can be sustainably maintained indefinitely without influencing the ecosystem’s network metrics), while those species above the upper bound of the Green Band are being “structurally overfished” and fishing mortality should be reduced. Those species with points below the lower bound of the Green Band have scope for an increase in exploitation. The location of species vs the Green Band can be tracked through time to see whether fisheries management is moving stocks towards or away from the Green Band (i.e. whether management change is required to avoid the risk of undesirable ecosystem pressure).

Note that in the balanced harvest literature that inspired the original derivation of the method there has been much discussion over whether to use production (which would see the green band defined in terms of production alone) or species productivity per biomass (which would the green band defined in terms of production / biomass) in the various calculations. We have used production because early simulation testing, which concurs with the subsequent work of Law and Plank (2022), found that only production led to stable biomasses, while the other approach could see rarer species become heavily depleted due to density dependent effects and nonlinearities of responses.

In our application of the method, we have relied on ecosystem model (Ecopath with Ecosim) output, given the need for an unfished profile (generated in our models by running the modelled ecosystems without fishing pressure for an extended period of time). However, in theory, it could be calculated solely from observational data if suitably long time series of the appropriate measures were available. In many instances this will likely require using model derived values, but space for time substitution (such as used in Zamborain-Mason et al. 2023) offers an alternative approach for those preferring to use observations instead. This is discussed further in Law and Plank (2023) which also discusses the method of Heath et al (2017).

As the method was being applied here based solely on ecosystem models the sensitivity of the approach to the taxonomic resolution of the ecosystem model should be considered. This was achieved by simplifying with a highly resolved version of the EwE model of the eastern Bering Sea (Aydin et al., 2007), which had 111 taxonomic groups, 14 of which were split into juvenile and adult age life history stages (Table S1, Figure S1). The simplification involved using the aggregation methods typically employed by ecological modellers (combining age classes of a single species, combining species into functional groups etc). Following this approach, the most simplified model had 24 functional groups – still spanning from plankton to whales. The Green Band procedure was then repeated using the different model structures to see (i) how strongly the final slope and shape of the Green Band was influenced and (ii) whether it would have influenced any management advice (i.e. whether a species group was consistently judged as “acceptable”, “structurally overfished”, “capacity of expansion” in each case).

### Application - Case studies

The Green Band procedure was applied to EwE models fit to data from the four case study locations, which spanned a diverse range of ecosystem types and management contexts:

I. Northern central Chile (between 24°S and 32°10’S): This highly productive upwelling system is primarily fished by a quasi-commercial artisanal fishery. Formal fisheries management in the region has historically been on a species-by-species basis.
II. Kerala, India: This seasonal and productive tropical system has been fished for many decades by thousands of fishers. The gears used now and in recent years range from very simple traditional gears to highly mechanised offshore operations. The complexity of the ecosystem and the fishery sectors has led to some historical overfishing (though only some species and sectors are affected) and current fisheries management arrangements focus on effort control measures (including gear, seasonal and spatial management).
III. South eastern Australia: This is a low productivity temperate system that is, nevertheless, highly ecologically diverse. Fisheries in the region, commercial and recreational, use a wide range of fishing gears and interact with 100s of species. The most intensively targeted commercial species (some of which have been overexploited historically) are managed using individual transferrable quotas as part of an integrated management scheme that also includes, gear, seasonal and spatial controls and consideration of the broader ecosystem via an ecological risk assessment approach.
IV. Eastern Bering Sea, USA: This sub-Arctic shallow shelf, which extends 500km from coastal Alaska, is one of the most highly productive regions globally. It sustains high catches by volume, but is tightly managed using a regulatory system that relies on individual species catch limits, along with a cap on system level exploitation.

The base model run (the simulation fit to historical catches and driven by historical effort time series) was used to assess at how species in the ecosystem mapped against the Green Band for that system through time (see details of the case study areas and historical period in Table 2 and see Supplementary materials for the historical trajectories of biomass by species/functional group and system). This served as a test to assess how informative the approach was and what were any challenges or caveats to be aware of. To help understand how different parts of the ecosystem are affected, the species and functional groups in each model were classified as (a) fishery target species (the primary targets of the fisheries in the ecosystem), (b) secondary targets of the main fisheries in the ecosystem (this classification was needed in some ecosystems due to the large number of fished species), (c) bycatch species (species caught but not landed), (d) habitat forming species (benthic biogenic habitats such as corals, sponges, macroalgae etc), (e) species critical to system coherency (this classification is based on a method of finding in the hubs in the food web, defined using the method of Fulton et al., in prep) and (f) species of conservation concern (such as marine mammals and seabirds).

**Table 2:** Details of the case study locations and the historical period covered by the base run for the EwE model for each system.

| Attribute | Eastern Bering Sea | North central Chile | Kerala, India | Southeast Australia |
| --- | --- | --- | --- | --- |
| Ecosystem type | Subpolar | Upwelling | Tropical | Temperate |
| Spatial Extent | 500km from Alaskan coast<br>54°15' to 65°30'N | Chile EEZ<br>24°S to 32°10'S | Indian EEZ off Kerala state<br>8°S to 13°N | Australian EEZ<br>24°29'S to 49°S<br>115°08'E to 164°E |
| Historical period considered | 1971-2017 | 2006-2018 | 1985-2019 | 1985-2019 |

## Results

### Network Analysis

First we considered whether fishing distorts the structure of ecosystems and whether that is clear from network metrics. Figure 2 shows that the shape of the cumulative distribution curve for a system changes with the intensity and the pattern of fishing for south east Australia. The cumulative distributions are identical at low levels of fishing pressure and when in the green band, but as fishing intensifies to FMMSY or 0.325M (0.65×0.5xM) and beyond then the curve changes. This means the structure is maintained when fishing is light (<FMMSY Figure 2b and < “0.65 HalfM” in Figure 2c) and under the pattern of fishing consistent with the Green Band (F∼P in Figure 2a). However, as fishing pressure intensifies, more and more species are depleted, the shape of the cumulative distribution is modified, the system structure is distorted; the resulting curves no longer lie one on top of the other. This clearly shows that as fishing pressure increases the network characteristics (as captured by the cumulative distribution curve) changes. This is clear from the plots but was reinforced by the curve fitting (Table 3). Initially, the distortive pressure only changed the coefficients of the best fitting curves – all the curves from F0 to1.5FMMSY and “HalfM” are best fit by a scaled power curve with tail, though once F ≥ FMMSY or F ≥ 0.325M the coefficients differ to those for the curve best fit to lower F – but ultimately even the type of curve was affected (shifting first to linear relationships before shifting to either logarithmic or power curve relationships).

**Figure 2:**
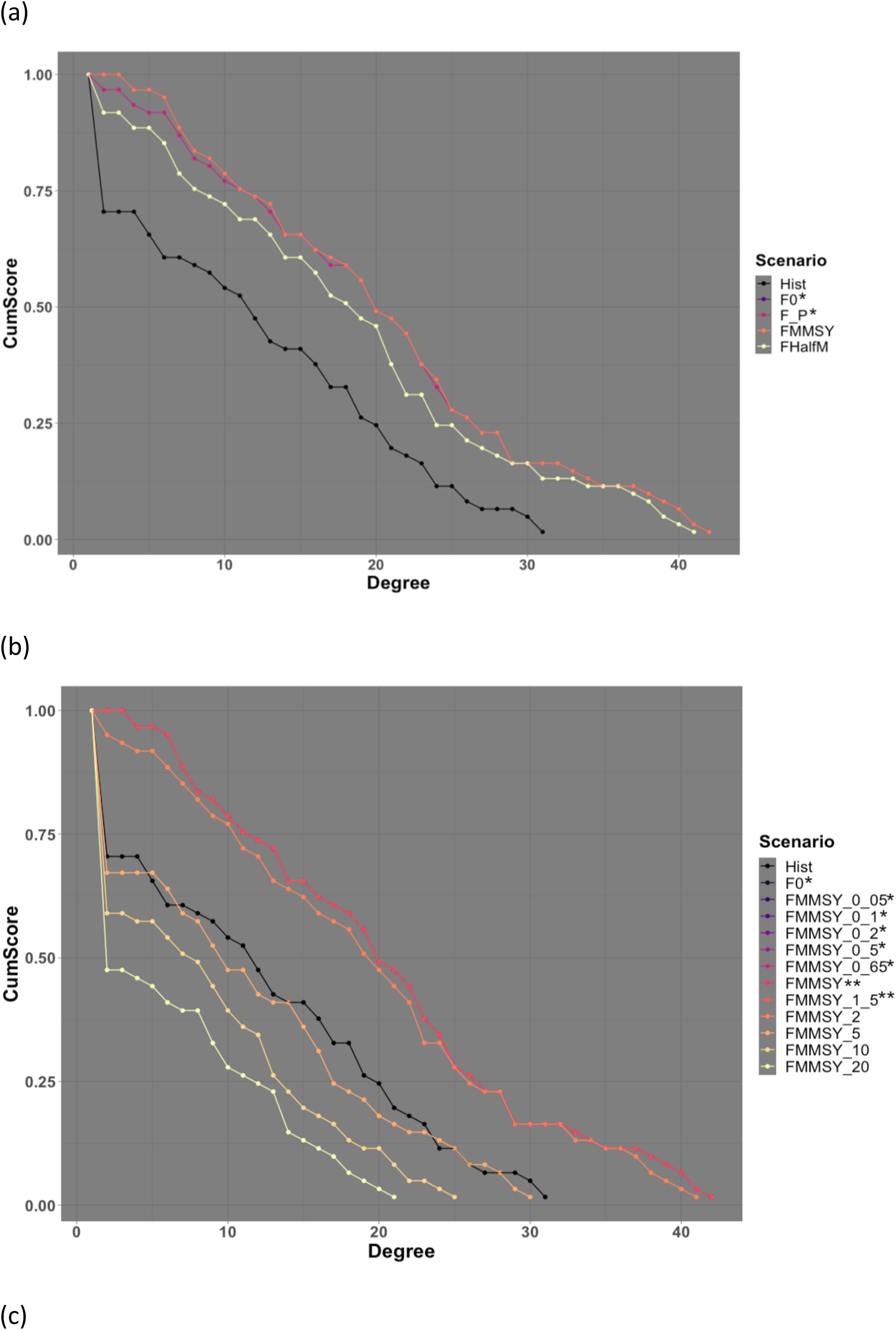
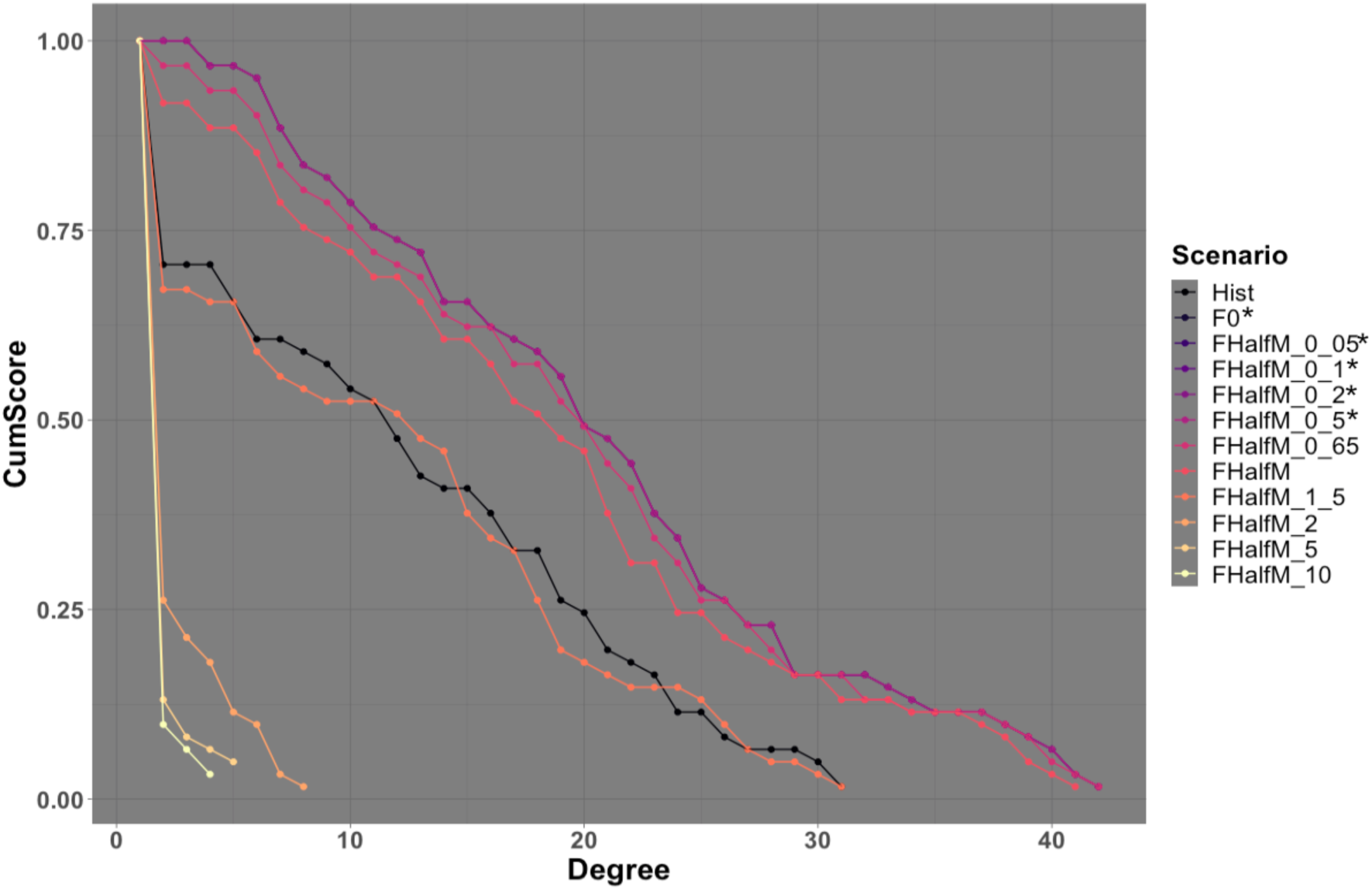
Plots of cumulative distribution of connections (as measured by degree, the number of connections to or from nodes in a network) from the southeast Australian EwE model under different levels and patterns of fishing pressure: (a) unfished, 𝑃^1+𝑎^, MMSY and half of M (natural mortality); (b) multiples of FMMSY; (c) multiples of half M. In each of these plots an * or ** indicates lines that lie one on top of the other. The Historical case is shown in all 3 plots for reference.

**Table 3:** Best fit curves for the cumulative distribution of connections from each south eastern EwE model fishing scenario.

| Fishing scenario | Best fit curve type | Measure of Fit |  |
| --- | --- | --- | --- |
|  |  | R<sup>2</sup> | AIC (AIC Range across models tested(min,max)) |
| F0 (unfished)<br>$F = P^{1+a}$ (consistent with Green Band)<br>$0.05 \cdot F_{MMSY}$<br>$0.1 \cdot F_{MMSY}$<br>$0.2 \cdot F_{MMSY}$<br>$0.5 \cdot F_{MMSY}$<br>$0.65 \cdot F_{MMSY}$<br>$0.05 \cdot \text{Half M}$<br>$0.1 \cdot \text{Half M}$<br>$0.2 \cdot \text{Half M}$<br>$0.5 \cdot \text{Half M}$ | $P(z) = z^{0.2442} e^{\frac{-z}{12.811}}$ | R<sup>2</sup>: 0.9797 | AIC: -129.87 (-129.87, -1.41) |
| $F_{MMSY}$<br>$1.5 \cdot F_{MMSY}$ | $P(z) = z^{0.243} e^{\frac{-z}{12.677}}$ | R<sup>2</sup>: 0.9853 | AIC: -123.24 (-123.24, -1.41) |
| $2 \cdot F_{MMSY}$ | $P(z) = 1.00574 - 0.02607z$ | R<sup>2</sup>: 0.9849 | AIC: -136.97 (-136.97, -7.40) |
| $5 \cdot F_{MMSY}$ | $P(z) = 0.78896 - 0.02781z$ | R<sup>2</sup>: 0.9537 | AIC: -80.64 (-80.64, -33.48) |
| $10 \cdot F_{MMSY}$ | $P(z) = 0.9786 - 0.2833 \log(z)$ | R<sup>2</sup>: 0.9296 | AIC: -59.58 (-59.58, -33.77) |
| $20 \cdot F_{MMSY}$ | $P(z) = 0.8601 - 0.2647 \log(z)$ | R<sup>2</sup>: 0.9088 | AIC: -46.59 (-46.59, -32.54) |
| $0.65 \cdot \text{Half M}$ | $P(z) = z^{0.2163} e^{\frac{-z}{13.0898}}$ | R<sup>2</sup>: 0.9783 | AIC: -127.82 (-127.82, -5.90) |
| Half M | $P(z) = z^{0.1842} e^{\frac{-z}{13.1548}}$ | R<sup>2</sup>: 0.9778 | AIC: -126.09 (-126.09, -11.99) |
| $1.5 \cdot \text{Half M}$ | $P(z) = 0.78879 - 0.02692z$ | R<sup>2</sup>: 0.9491 | AIC: -82.24 (-82.24, -31.87) |
| $2 \cdot \text{Half M}$ | $P(z) = 0.01887z^{-1.66274}$ | R<sup>2</sup>: 0.9450 | AIC: -16.16 (-16.16, 8.08) |
| $5 \cdot \text{Half M}$ | $P(z) = 0.0336z^{-3.1729}$ | R<sup>2</sup>: 0.9943 | AIC: 5.31 (5.31, 29.97) |
| $10 \cdot \text{Half M}$ | $P(z) = 0.01942z^{-3.48484}$ | R<sup>2</sup>: 0.9968 | AIC: -15.59 (-15.59, 5.87) |
| Historical fishing pressure (2014) | $P(z) = 0.81396 - 0.02751z$ | R<sup>2</sup>: 0.9792 | AIC: -94.39 (-94.39, -31.07) |

### Sensitivity to model structural complexity

Model simplifications had some effect on the slope of the unfished biomass vs production profiles – with the resulting slope parameters varying by less than 5% for the majority of simplified models. Only the simplest model – which had simplified 125 taxonomic/age groups to 24 aggregate functional groups – had a slope that differed by > 7% from the full case (its slope was 13% flatter than for the full model). This meant that the resulting Green Band produced was relatively robust to taxonomic resolution (Figure 3) with functional groups rated consistently versus their relative location (below, in or above the green band). Where detail was lost, at the individual species level, it was because a rare species rating was subsumed by the more dominant species in the functional group.

**Figure 3:**
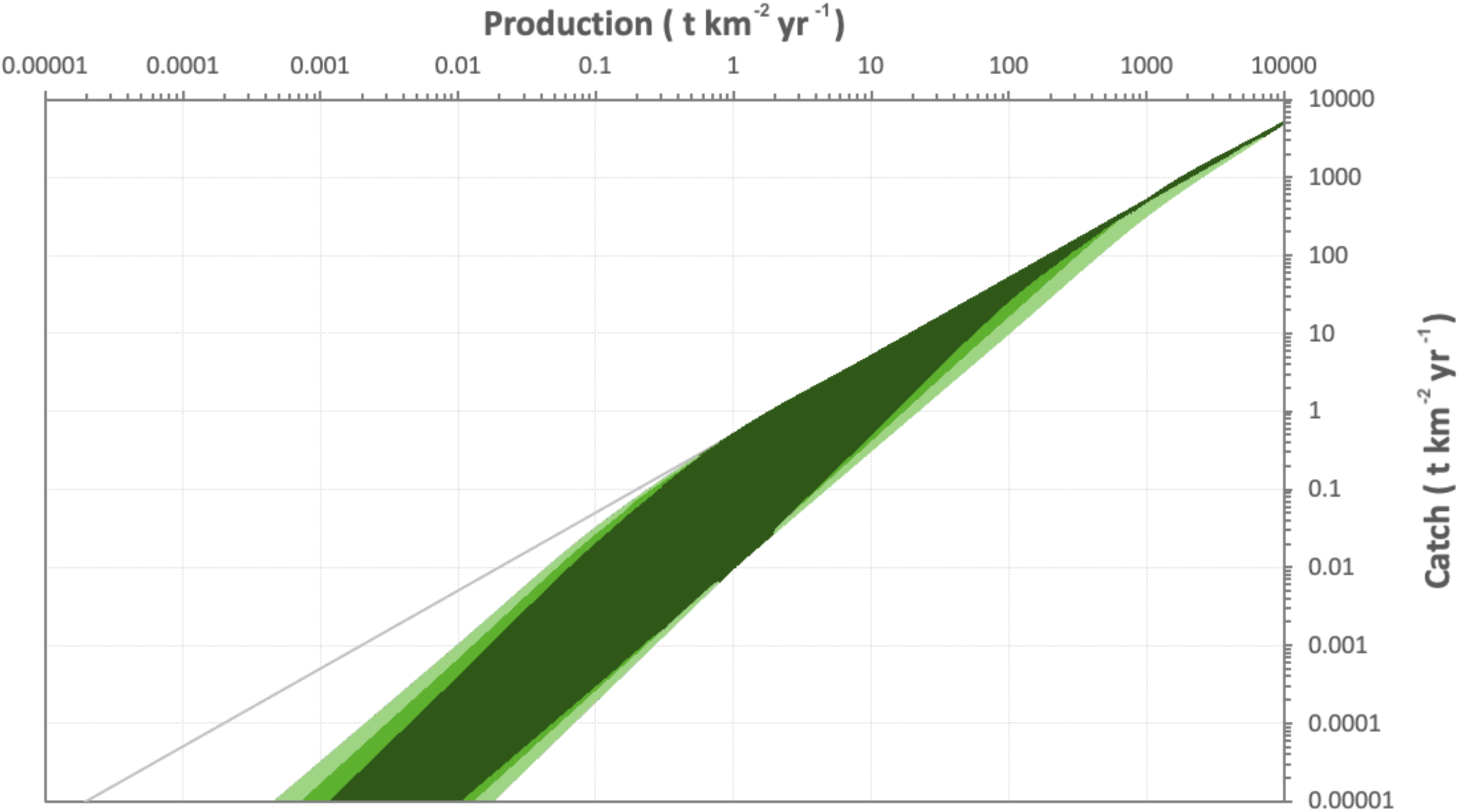
Plot showing the green band area produced by the differing levels of simplification. The dark area was present in all cases, the lighter area in less cases – the greater the simplification the flatter the slope (so the more it moved toward the half M reference line at the lower left and more toward the right axis in the upper right).

### Application - Case studies

Very few of the exploited groups are located within the “Green band” for the north central Chilean ecosystem (Figure 4). Several the highly productive target species are close to the upper margin (or are trending into it under current management), indicating that further action is needed to remove distortive pressure on this part of the structure of the ecosystem. Some of the species of conservation concern (e.g. sharks) are well above the green band and are apparently moving away from it. The fact that sea lions and seabirds are not directly harvested means they are not threatened in the same way.

**Figure 4:**
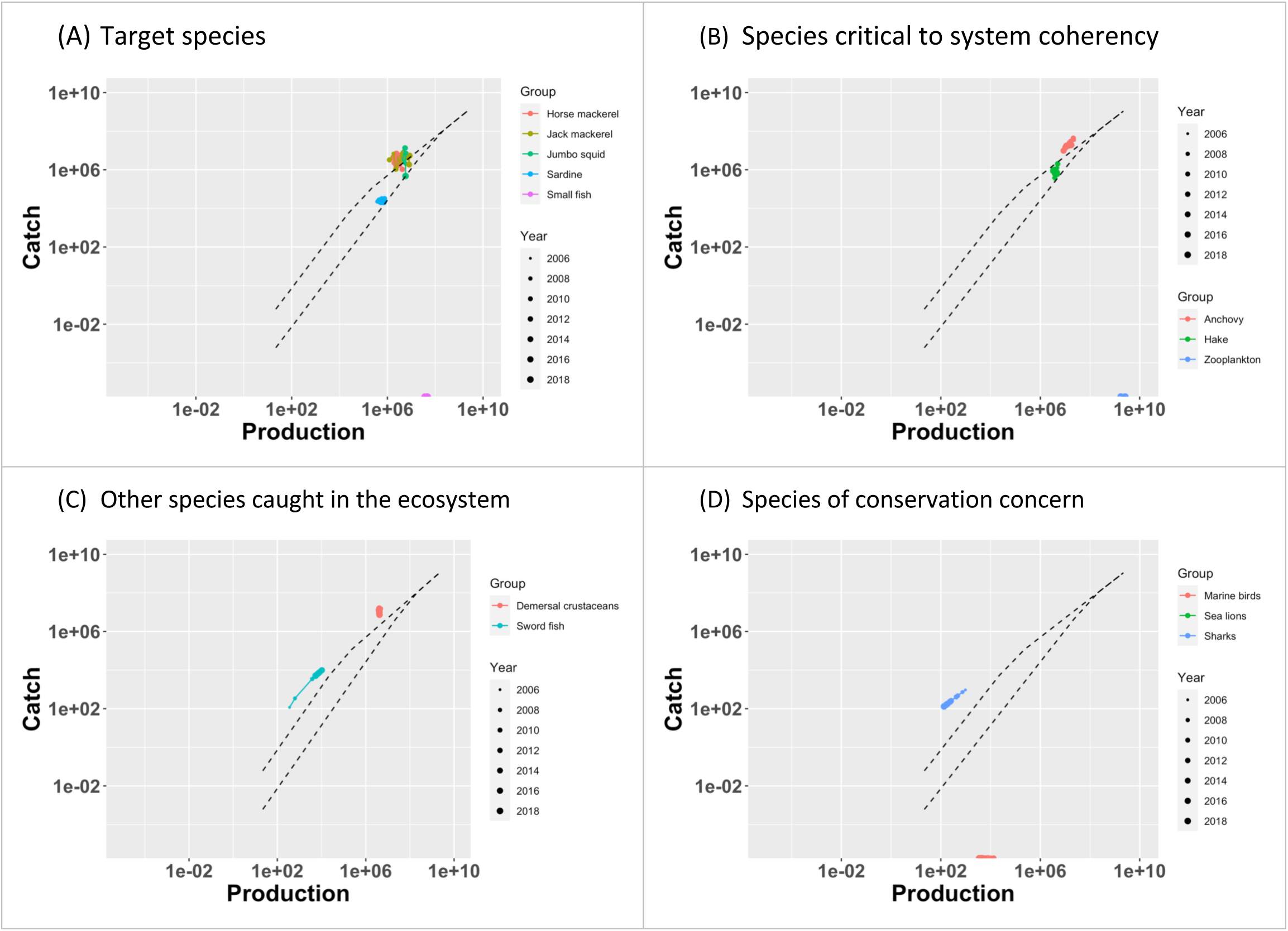
Green Band plots for the north central Chilean system between 2006 and 2018. The bounds of the Green Band are represented by the dashed lines.

Many of the species in the Kerala ecosystem are outside the “Green band” (Figure 5). Indeed, over the last three decades many species have not only sat above the Green Band but are trending away from it.

**Figure 5:**
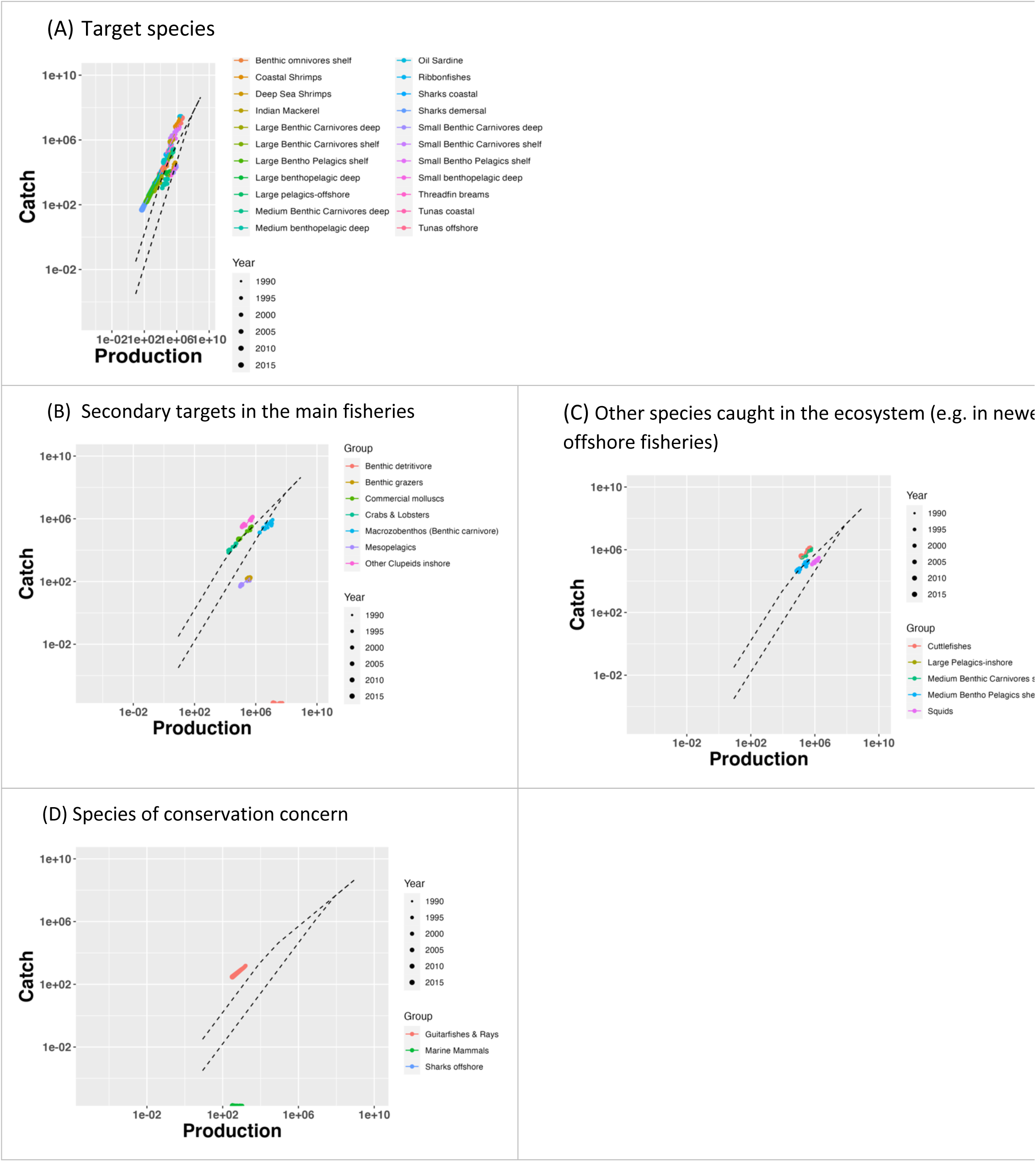
Green Band plots for Kerala, India between 1990 and 2017. The bounds of the Green Band are represented by the dashed lines.

The situation is more mixed in the south eastern Australian ecosystem. While many of the species groups considered less important by the fisheries of the region are inside the Green Band, or are only under light pressure (Figure 6), both the current and historically targeted species have been subject distortive pressure. Fortunately, more recent management actions seem to be rectifying that for several main target species, with more recent points sitting within the Green Band. A non-trivial proportion of the species of conservation concern are not faring as well, particularly the aggregate tuna and billfish group, pelagic sharks and seals.

**Figure 6:**
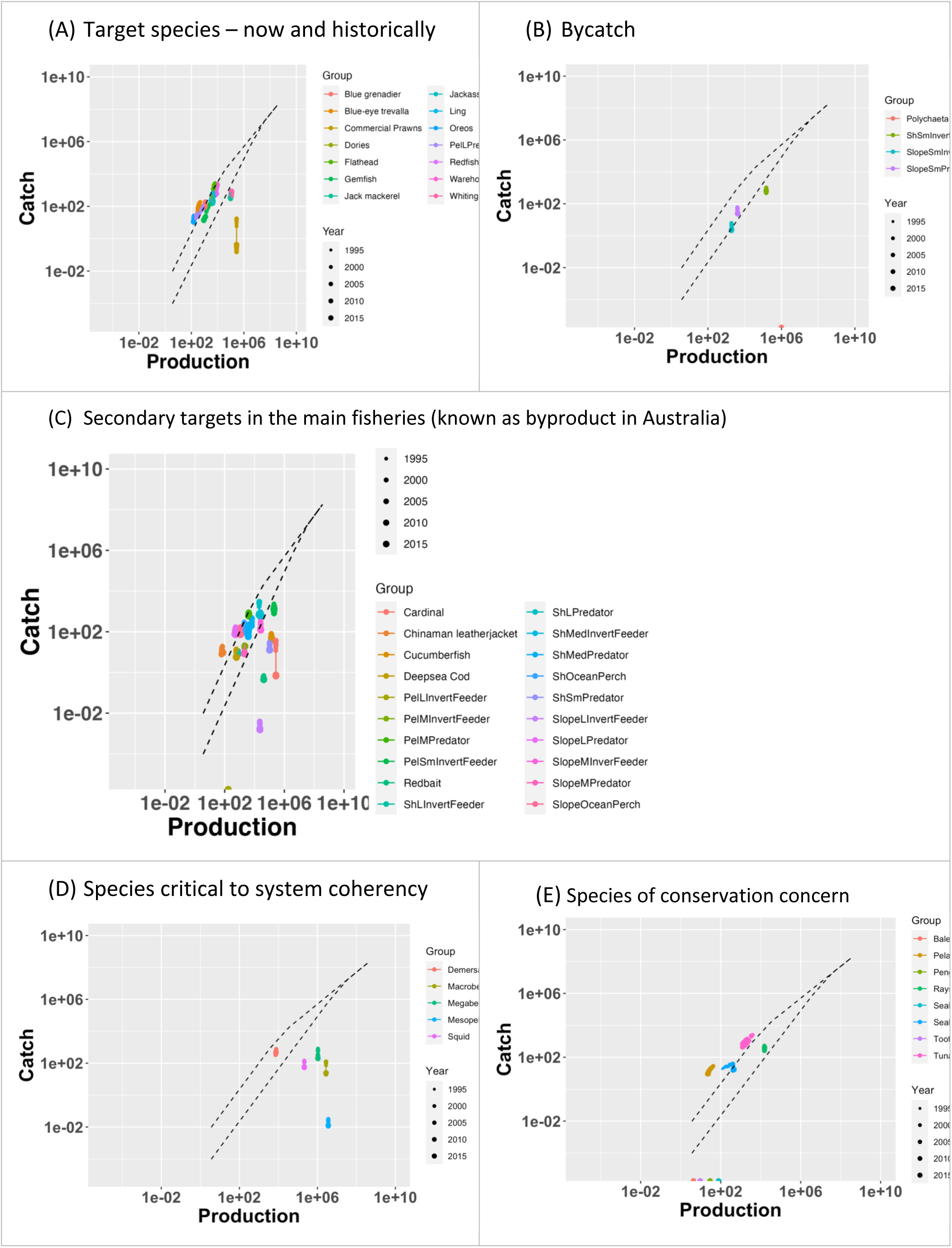
Green Band plots for the south east Australia ecosystem between 1994 and 2018. The bounds of the Green Band are represented by the dashed lines.

In common with Australia, the results for the eastern Being Sea were mixed. Many species were inside or below the “Green band” (Figure 7). However, even in the eastern Bering Sea not all groups are free from distortive pressure; several species of conservation concern are above the Green Band due to pressure from incidental interactions with fisheries. Similarly, corals and a small number of the most slow growing or vulnerable target species sit above the ‘Green band’. In the majority of cases (across all species) management has been taking species into or below the ‘Green band’ with the exception of Pacific ocean perch,some rockfish (Northern, Dusky, Rougheye, Sharpchin and Shortraker) and “Other Sebastes” (which are rockfish).

**Figure 7:**
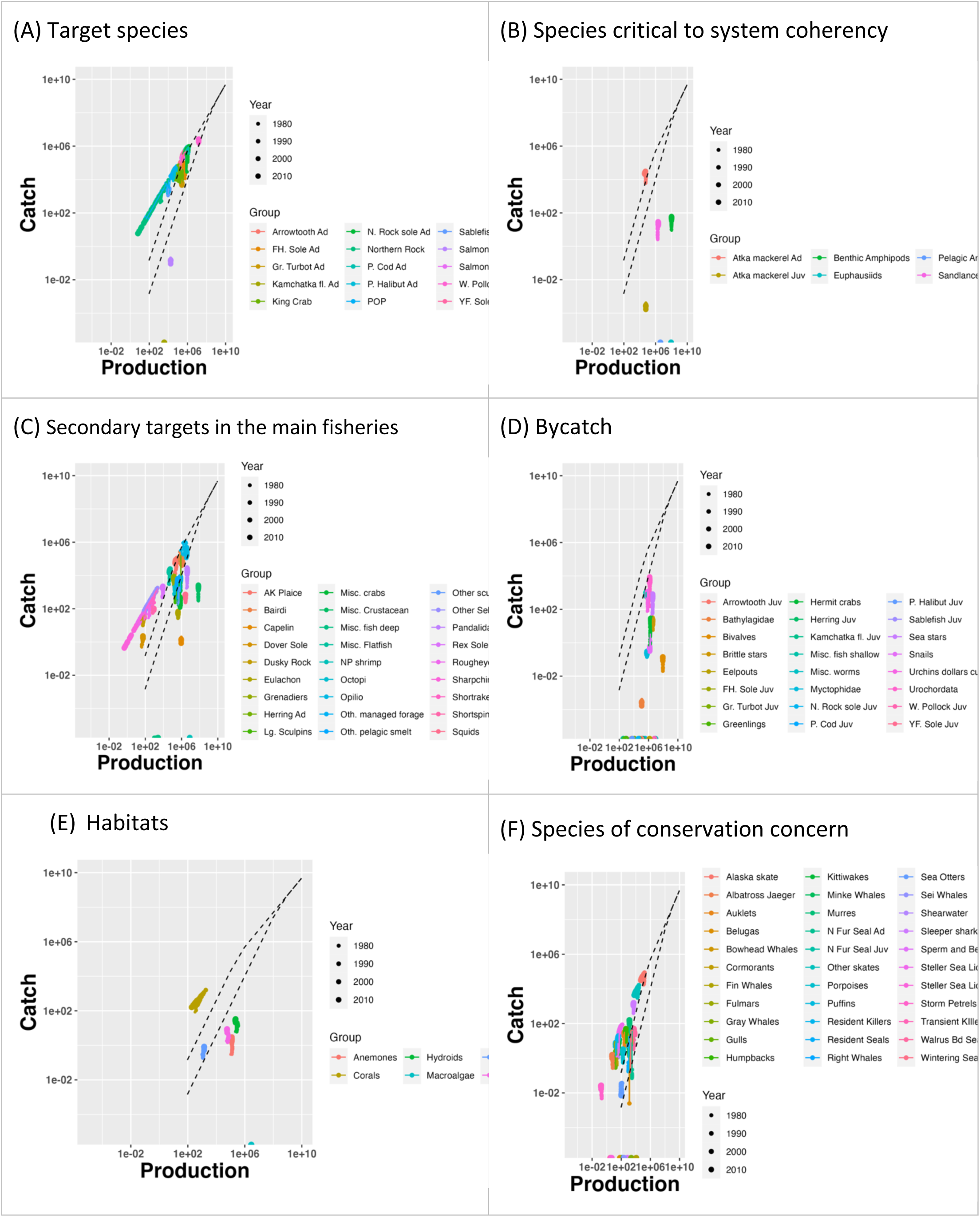
Green Band plots for a snapshot of the East Bering Sea from the 1970s. The bounds of the Green Band are represented by the dashed lines.

## Discussion

While there is broad agreement on the need for a systemic perspective to set a context for fisheries management actions (Patrick and Link, 2015), details regarding the specific implementation have been more challenging to identify (Lidström and Johnson, 2020). While there has been consider work over the past two decades on relevant indicators (e.g. Jennings, 2005; Link, 2005; Shin and Shannon, 2010; Shannon et al., 2010; Shin et al., 2018) the majority have focused on performance measures that are useful for tracking status and trends – such as relative abundance – rather than structural components of ecosystems.

In parallel, ecologists have been using network theory to explore ecosystem properties (e.g. Dunne et al., 2002; Milo et al., 2002; Montoya and Sol, 2002; Jordan et al., 2006; D’Alelio et al., 2016; Marina et al., 2018; Niquil et al., 2020). Network theory and associated indicators have been used to explore rules around food web formation (Dunne et al., 2004; Fath et al., 2007) as well as reporting on ecosystem status (Libralto et al., 2006) and theoretical consideration of management options (McDonald-Madden et al., 2015). Nevertheless, there have been few studies that directly consider how network analysis can elucidate the distortion of ecosystems due to fishing or other perturbations. Navia et al., (2016) are a rare exception. The analysis method presented here would do with empirical verification, but they do suggest that moderate to heavy fishing pressure can change patterns of ecosystem connectivity to the point that the cumulative distribution of connections is notably affected.

The modification of the shape of connections is in part related to the depletion and loss of species from the food web. However, it may also indicate more than network simplification; it could indicate that the maintenance or ongoing network formation is challenged. Food webs, including those modelled here, share properties with scale-free networks suggesting the rich literature on such networks could provide deeper insights for food webs. One insight is that scale-free networks form because new nodes attach links preferentially to already highly connected nodes (Amaral et al., 2000). Broad-scale and single-scale networks form from a similar process, but in those cases the highly connected nodes eventually stop receiving new links. For example, they may become less likely to accept a new link, or maintain an existing link, because there is a cost associated with those links and this eventually becomes overwhelming (Amaral et al., 2000). The preliminary work on the topic presented here suggests that disruption by fishing induces a cost in network formation or maintenance (e.g. in situations where new, rare or migratory species need to re-enter the food web periodically, such as is the case in the south east Australian system). In other instances, fishing simply truncates the nodes, artificially producing a pattern that implies a cost. Both mechanisms occur in the modelled food webs explored here. More work would be needed to clarify more generally what process holds under what circumstances and how sensitive the result is to defining where the tail of the cumulative distribution begins, which would require gaining a greater understanding of what induces the shift in the tail of the distribution.

While this topic has rich potential and could be scientifically fruitful it’s immediate capacity for informing management is more limited. While it is possible that tail type could be monitored, along with the point where the tail begins (to check for truncation), clearly communicating such an abstract and poorly understood concept would likely make it of limited decision-making utility. As a result, we have left this topic for future research and refinement. However, the fact that fishing appears to modify the distribution of ecosystem connections suggests that measures leveraging the theory of network connectivity or highlighting the application of distortive pressure on an ecosystem could usefully contribute to fisheries decision making in the context of EBFM. As the Green Band does exactly that and uses methods and information that are available in many systems it seems to present a promising tool for advising on EBFM.

Heath et al (2017) attempt to interpret how natural systems are structured and how fishing in proportion to production could in theory mimic that, and provides a way to identify when there is distortive pressure in an ecosystem. Law and Plank (in press) have since simulation tested the concept and shown it can maintain abundance and ecosystem structure through time. Similarly, the tests undertaken in this paper show that staying within the Green Band maintains the cumulative distribution of connectivity, while moving above it and overfishing parts of the system leads to a change in the. Moreover, the tests show the method is reasonably robust to issues of the resolution of the models used, so long as the models used follow best practice model construction with regard to defining functional groups and avoiding inappropriate aggregation of groups (Fulton et al., 2003; Link, 2010; Heymans et al., 2016).

The Green Band could be used in snap shot form to judge the pressure on an ecosystem (see Heath et al 2017). It could also be used through time to inform management agencies as to whether the interventions they have made are guiding the system to patterns of fishing that have less distortive pressure on ecosystem structure. The results across the case study ecosystems here show systems under different patterns and history of fishing pressure and management intervention success. In Kerala (and similarly in north central Chile) the number of groups above the Green Band reflects how intense fishing is leading to distortive pressure on the ecosystem structure, where very few species are in or below the Green Band. This is lead to quite substantial shifts in catch composition (Fulton et al., submitted) and on ecosystem resilience (Fulton et al., in prep). In contrast, a majority of groups within or below the Green Band for the eastern Bering Sea has, which is at least partly result of the precautionary ecosystem cap used in the system. Although it is noteworthy that many species of conservation concern are above the Green Band, showing how there can be a disconnect between dedicated fisheries and conservation management when not taking a systemic approach across the board. South eastern Australia has also had a mixed history in terms of the pattern of pressure it has applied to the ecosystem, although it appears that management reforms over the past decade have helped reduce the pressure on some species. These management actions were motivated by single-species management objectives (Smith et al., 2008) and the Green Band places these actions in a broader context, showing how the Green Band can complement existing management approaches, helping progress to an ecosystem approach without requiring a radical “all at once” overhaul of management processes.

The shape of the Green Band versus the reference line – which equates to the single species management bound on exploitation of half of production – clearly shows why simple single-species management rules of thumb fail to maintain many of the less productive species.

The lower the productivity the greater the divergence between the Green Band and the reference line, indicating that the constant across species assumptions regarding limits on exploitation are not conservative enough to sustain the less productive species. This is also seen in the work by Law and Plank (in press), where long-term simulations applying F ∼ P saw stable systems but F ∼ P/B saw some species decline, as did the application of a constant F=0.1 across all species. This illustrates how single species management as it is often applied in diverse systems with constrained resources will be insufficient to keep the entire ecosystem healthy (Hilborn, 2011) even if it does serve to recover some key target species (Hilborn et al., 2020). EBFM will involve more than just single-species management and there is a growing number of tools that can complement single species management by highlighting when single-species management is not delivering on ecosystem level objectives (e.g. Link et al., 2015; Libralto et al., 2019; Link and Watson, 2019; Link, 2021; Fulton et al., submitted). The Green Band is one example to add to this list of operational EBFM tools.

## Acknowledgements

This work was supported by the Lenfest Ocean Program and CSIRO Australia. Individual participants also supported by NOAA Alaska Fisheries Science Center, Cooperative Institute for Climate, Ocean, and Ecosystem Studies, University of Washington, North Pacific Fishery Management Council, Washington Department of Fish and Wildlife, Instituto de Fomento Pesquero, Subsecretaría de Pesca y Acuicultura, Central Marine Fisheries Research Institute, and the Directorate of Fisheries in India.

