## Supplementary materials for "“The Green Band”: Using Production and Catch to Judge Distortive Pressure on an Ecosystem"

Ecopath with Ecosim (EwE; Christensen and Walters, 2004) was used to model each of the case study ecosystems. Each of these models was constructed by the system experts and conditioned on historical data. Where sufficient information was available to fit the model closely to historical fisheries and survey data that was done, otherwise a fit considered plausible by the system experts given available data were used as a basis of the analyses. The base parameters, catch, discards and diet of the resulting models are shown in Tables S2-S15.

Ecosim models were initialised from the Ecopath base models. Unfished states for each ecosystem were obtained by running the model forward with no fishing for 100 years to ensure all groups reached equilibrium. Given species with very low production rates can take very long time periods to fully rebuild, in obtaining the unfished state, if any groups had not reached stable equilibrium after 100 years the model simulation was extended to 200 years to guarantee all species had reached an equilibrium state. This was important to make sure there was a clear and certain state to compare against. Moreover, in those cases where groups had not reached equilibrium by the end of 100 years they were still strongly trending and not approaching equilibrium yet.

When completing the other simulations (different levels of fishing pressure) the simulations were not extended to 200 years even in the (very) rare case where a species was not in an equilibrium state by the end of the 100 year simulation period.  This was because across all the simulations only a very small number of species had not reached absolute equilibrium by the end of the 100 years, but had already started to plateau and were obviously close to equilibrium (and running the simulations longer makes no material difference to the results).

The simulation of the historical pattern of fishing was represented by scaling historical effort levels so they were relative to the level of effort in the year(s) used to create the base Ecopath model. These relative effort time series were used to replicate the patterns of fishing through time and force the Ecosim model (using the method described in Ecopath User Guide; Christensen et al 2008)). The resulting biomass, catch and mortality time series formed the 'dataset' for the rest of the analyses (described in the main text). Effort rather than direct catch time series forcing was used so that the broader ecosystem footprint could be captured.

Effort time series were not available for the eastern Bering Sea ecosystem so a proxy was created by constructing a relative time series based on the pattern of catches seen in their primary target species. This was judged to create a plausible historical trajectory, with the sufficiency of this approach was judged based on whether these proxy time series reasonably reproduced observed catch time series for secondary species. An alternative approach could have been to use an F time-series, but this was not available for all taxonomic units in the model (including bycatch species).

***Model Variants – For Judging Sensitivity to Model Resolution***

**Table S1:** List of groups in each of the variant (sequentially simplified) models of the eastern Bering Sea

| 129 groups including age structure | 81 groups | 53 groups | 26 groups |
|---|---|---|---|
| Transient Killers | Killers Whales | Killers Whales | Baleen whales |
| Sperm and Beaked Whales | Toothed Whales | Toothed Whales | Toothed whales |
| Resident Killers | Porpoises | Porpoises | Sperm whales |
| Porpoises | Gray Whales | Gray Whales | Beaked whales |
| Belugas | Baleen whales | Baleen whales | Walrus & Bearded seals |
| Gray Whales | Sea Otters | Sea Otters | Seals |
| Humpbacks | Walrus Bd Seals | Walrus Bd Seals | Steller sea lions |
| Fin Whales | N. Fur Seal | N. Fur Seal | Pisc. Birds |
| Sei whales | Steller Sea Lion | Steller Sea Lion | Pollock |
| Right Whales | Resident seals | Seals | Adult pollock2+ |
| Minke Whales | Wintering seals | Seabirds | Juv. pollock0-1 |
| Bowhead Whales | Mobile Seabirds | Sleeper shark | Other Demersal Fish |
| Sea Otters | Inshore Seabirds | W. Pollock | Large Flatfish |
| Walrus Bd Seals | Sleeper shark | P. cod | Small Flatfish |
| N. Fur Seal | W. Pollock | Herring | Shallow Pelagics |
| N. Fur Seal juv | P. cod | Arrowtooth | Deep Pelagics |
| N. Fur Seal adults | Herring | Flatfish | Deepwater fish |
| SSL | Arrowtooth | Skates | Jellyfish |
| Steller Sea Lion juv | Kamchatka fl. | Sablefish | Cephalopods |
| Steller Sea Lion adults | Gr. Turbot | Eelpouts | Benth.P.Feeders |
| Resident seals | P. Halibut | Deepwater Fish | Infauna |
| Wintering seals | YF. Sole | POP | Epifauna |
| Shearwater | FH. Sole | Demersal fish | Large Zooplankton |
| Murres | N. Rock sole | Atka mackerel | Herbivorous Zooplankton |
| Kittiwakes | Flatfish | Sculpins | Phytoplankton |
| Auklets | Skates | Misc. fish shallow | Discards |
| Puffins | Sablefish | Cephalopods | Detritus |
| Fulmars | Eelpouts | Salmon | |
| Storm Petrels | Deepwater Fish | Mesopelagics | |
| Cormorants | POP | Forage fish | |
| Gulls | Rockfish | Crustaceans | |
| Albatross Jaeger | Shortspine Thorns | Sea stars | |
| Sleeper shark | Other Sebastes | Urchins dollars cucumbers | |
| W. Pollock | Atka mackerel | Snails | |
| W. Pollock juv | Greenlings | Benthic Amphipods | |
| W. Pollock adults | Sculpins | Coral-like | |
| P. cod | Misc. fish shallow | Urochordata | |
| P. cod juv | Octopi | Sea Pens & Sponges | |
| P. cod adults | Squids | Bivalves | |
| Herring | Salmon | Worms | |
| Herring juv | Bathylagidae | Scyphozoid | |
| Herring adults | Myctophid | Large Zooplankton | |
| Arrowtooth | Forage fish | Pelagic Amphipods | |
| Arrowtooth juv | Bairdi | Gelatinous filter feeders | |
| Arrowtooth adults | King Crab | Pteropods | |
| Kamchatka fl. | Opilio | Copepods | |
| Kamchatka fl. juv | Pandalidae | Microbes | |
| Kamchatka fl. adult | NP shrimp | Macroalgae | |
| Turbot | Sea stars | Phytoplankton | |
| Gr. Turbot juv | Brittle stars | Outside Production | |

| | | |
|---|---|---|
| Gr. Turbot adults | Urchins dollars cucumbers | Discards |
| P. Halibut | Snails | Detritus |
| P. Halibut juv | Hermit crabs | Outside Detritus |
| P. Halibut adults | Misc. crabs | |
| YF. Sole | Misc. Crustacean | |
| YF. Sole juv | Benthic Amphipods | |
| YF. Sole adults | Coral-like | |
| FH. sole | Urochordata | |
| FH. Sole juv | Sea Pens & Sponges | |
| FH. Sole adults | Bivalves | |
| Rock sole | Polychaetes | |
| N. Rock sole juv | Misc. worms | |
| N. Rock sole adults | Scyphozoid | |
| AK Plaice | Fish Larve | |
| Dover Sole | Chaetognaths | |
| Rex Sole | Euphasids | |
| Misc. Flatfish | Mysids | |
| Alaska skate | Pelagic Amphipods | |
| Other skates | Gelatinous filter feeders | |
| Sablefish | Pteropods | |
| Sablefish juv | Copepods | |
| Sablefish adults | Pelagic microbes | |
| Eelpouts | Benthic microbes | |
| Grenadiers | Macroalgae | |
| Misc. fish deep | Large Phytoplankton | |
| POP | Sm Phytoplankton | |
| Sharpchin Rock | Outside Production | |
| Northern Rock | Discards | |
| Dusty Rock | Offal | |
| Shortraker Rock | Detritus | |
| Rougheye Rock | Outside Detritus | |
| Shortspine Thorns | | |
| Other Sebastes | | |
| Atka mackerel | | |
| Atka mackerel juv | | |
| Atka mackerel adult | | |
| Greenlings | | |
| Lg. Sculpins | | |
| Other sculpins | | |
| Misc. fish shallow | | |
| Octopi | | |
| Squids | | |
| Salmon returning | | |
| Salmon outgoing | | |
| Bathylagidae | | |
| Myctophid | | |
| Capelin | | |
| Sandlance | | |
| Eulachon | | |
| Oth. manahed forage | | |
| Oth. pelagic smelt | | |
| Bairdi | | |
| King Crab | | |
| Opilio | | |
| Pandalidae | | |
| NP shrimp | | |
| Sea stars | | |

| |
|---|
| Brittle stars |
| Urchins dollars cucumbers |
| Snails |
| Hermit crabs |
| Misc. crabs |
| Misc. Crustacean |
| Benthic Amphipods |
| Anemones |
| Corals |
| Hydroids |
| Urochordata |
| Sea Pens |
| Sponges |
| Bivalves |
| Polychaetes |
| Misc. worms |
| Scyphozoid |
| Fish Larve |
| Chaetognaths |
| Euphasids |
| Mysids |
| Pelagic Amphipods |
| Gelatinous filter feeders |
| Pteropods |
| Copepods |
| Pelagic microbes |
| Benthic microbes |
| Macroalgae |
| Large Phytoplankton |
| Sm Phytoplankton |
| Outside Production |
| Discards |
| Offal |
| Pelagic Detritus |
| Benthic Detritus |
| Outside Detritus |

**Figure S1:** Mapping of groups in the East Bering Sea EwE model versions – original on the left most side and then moving right each sequentially simpler model.

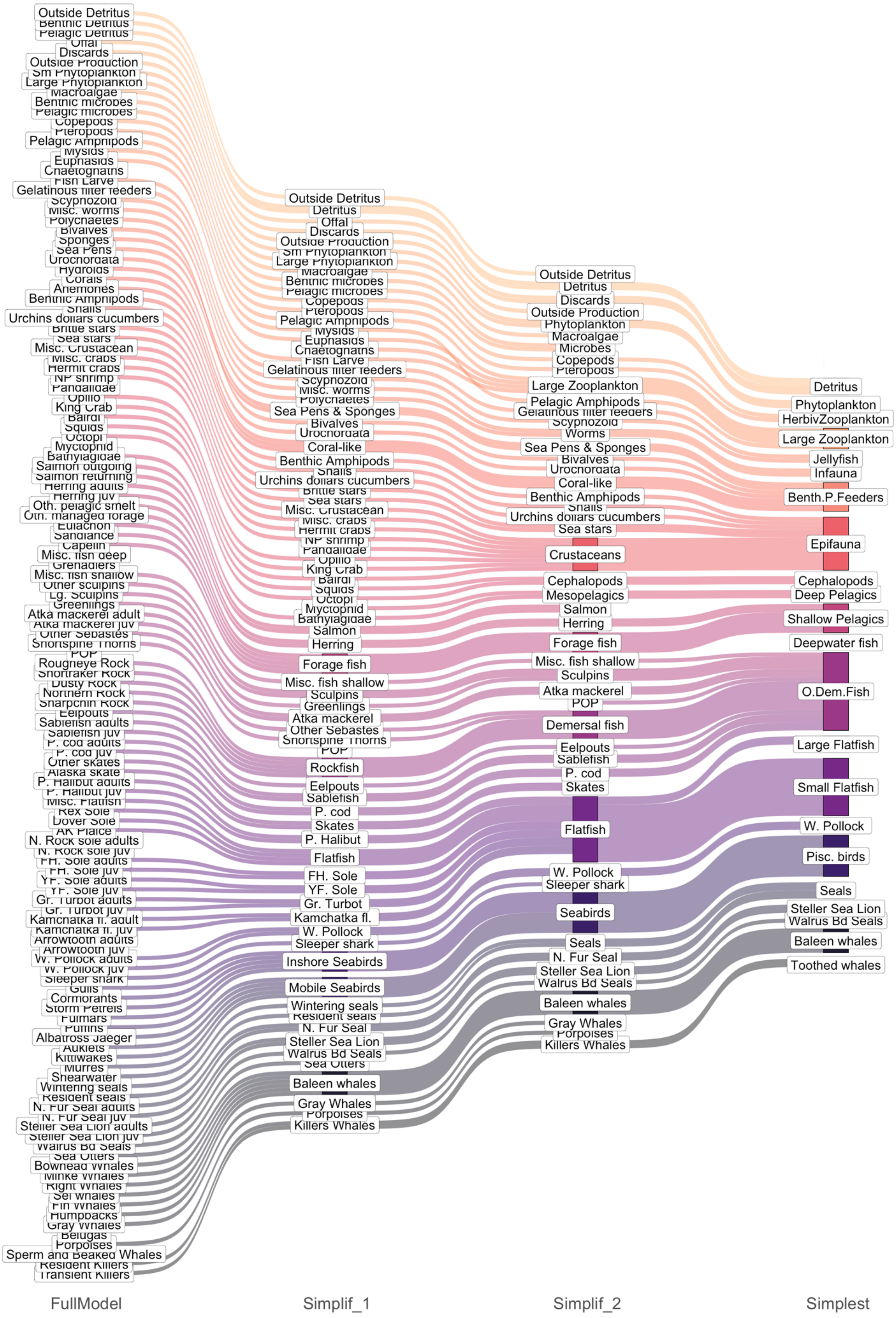

### *EwE Models*

**Table S2:** Ecopath base parameters for the East Bering Sea (converted from the Rpath version of the model)

| Group | Trophic Level | Biomass (tkm$^{-2}$) | *Mortality* or Production / Biomass (year$^{-1}$) | Consumption / Biomass (year$^{-1}$) | Ecotrophic Efficiency | Biomass Accumulation (tkm$^{-2}$) | Biomass Accumulation year$^{-1}$) |
|---|---|---|---|---|---|---|---|
| Transient KIllers | 4.825 | 0 | 0.025 | 11.157 | 0 | 0 | 0 |
| Sperm and Beaked Whales | 4.724 | 0.018 | 0.047 | 6.609 | 0 | 0 | 0 |
| Resident Killers | 4.683 | 0.001 | 0.025 | 11.157 | 0.17 | 0 | 0 |
| Porpoises | 4.63 | 0.004 | 0.05 | 30 | 0.055 | 0 | 0 |
| Belugas | 4.594 | 0.012 | 0.112 | 30 | 0.041 | 0 | 0 |
| Gray Whales | 3.511 | 0.033 | 0.063 | 8.873 | 0.039 | 0 | 0 |
| Humpbacks | 3.943 | 0.012 | 0.038 | 7.577 | 0.102 | 0 | 0 |
| Fin Whales | 3.731 | 0.455 | 0.027 | 6.517 | 0.093 | 0 | 0 |
| Sei Whales | 3.665 | 0.006 | 0.04 | 8.788 | 0.062 | 0 | 0 |
| Right Whales | 3.502 | 0.004 | 0.033 | 8 | 0.076 | 0 | 0 |
| Minke Whales | 4.12 | 0.024 | 0.051 | 7.782 | 0.052 | 0 | 0 |
| Bowhead Whales | 3.524 | 0.007 | 0.01 | 8.68 | 0.247 | 0 | 0 |
| Sea Otters | 3.739 | 0.001 | 0.117 | 73 | 0.022 | 0 | 0 |
| Walrus Bd Seals | 3.574 | 0.114 | 0.051 | 15.371 | 0.006 | 0 | 0 |
| N Fur Seal Juv | 4.583 | 0.001 | 0.252 | 82.2 | 0.021 | 0 | 0 |
| N Fur Seal Ad | 4.583 | 0.034 | 0.091 | 37.716 | 0.038 | 0 | 0 |
| Steller Sea Lions Juv | 4.653 | 0 | 0.194 | 55.252 | 0.063 | 0 | 0 |
| Steller Sea Lions Ad | 4.658 | 0.001 | 0.11 | 30.493 | 0.765 | 0 | 0 |
| Resident Seals | 4.529 | 0.013 | 0.083 | 17.439 | 0.045 | 0 | 0 |
| Wintering Seals | 4.579 | 0.03 | 0.069 | 19.197 | 0.87 | 0 | 0 |
| Shearwater | 4.536 | 0 | 0.1 | 73 | 0.085 | 0 | 0 |
| Murres | 4.433 | 0.008 | 0.169 | 72 | 0.05 | 0 | 0 |
| Kittiwakes | 4.4 | 0.001 | 0.077 | 110 | 0.111 | 0 | 0 |
| Auklets | 3.57 | 0.002 | 0.169 | 110 | 0.05 | 0 | 0 |
| Puffins | 4.397 | 0 | 0.04 | 73 | 0.213 | 0 | 0 |
| Fulmars | 4.588 | 0.001 | 0.055 | 73 | 0.154 | 0 | 0 |
| Storm Petrels | 4.281 | 0 | 0.12 | 144 | 0.07 | 0 | 0 |
| Cormorants | 4.485 | 0 | 0.159 | 73 | 0.054 | 0 | 0 |
| Gulls | 4.47 | 0 | 0.166 | 73 | 0.051 | 0 | 0 |
| Albatross Jaeger | 4.606 | 0 | 0.068 | 75 | 0.126 | 0 | 0 |
| Sleeper shark | 4.684 | 0.053 | 0.1 | 3 | 0.161 | 0 | 0 |
| W. Pollock Juv | 3.546 | 4.493 | 1.453 | 8.405 | 0.979 | 0 | 0 |
| W. Pollock Ad | 3.681 | 22.76 | 0.667 | 3.131 | 0.78 | 0 | 0 |
| P. Cod Juv | 3.62 | 0.186 | 1.782 | 8.828 | 0.798 | 0 | 0 |
| P. Cod Ad | 4.103 | 2.815 | 0.412 | 2.257 | 0.462 | 0 | 0 |
| Herring Juv | 3.522 | 0.171 | 1.052 | 7.79 | 0.804 | 0 | 0 |
| Herring Ad | 3.524 | 2.734 | 0.32 | 2.927 | 0.163 | 0 | 0 |
| Arrowtooth Juv | 3.977 | 0.006 | 0.81 | 6.139 | 0.983 | 0 | 0 |
| Arrowtooth Ad | 4.321 | 1.146 | 0.18 | 1.184 | 0.755 | 0 | 0 |
| Kamchatka fl. Juv | 4.128 | 0 | 0.81 | 6.139 | 0.034 | 0 | 0 |
| Kamchatka fl. Ad | 4.487 | 0.019 | 0.18 | 1.497 | 0.559 | 0 | 0 |
| Gr. Turbot Juv | 3.539 | 0.007 | 1.701 | 7.885 | 0.942 | 0 | 0 |
| Gr. Turbot Ad | 4.559 | 1.526 | 0.18 | 1.081 | 0.14 | 0 | 0 |
| P. Halibut Juv | 3.79 | 0.002 | 2.4 | 9.303 | 0.713 | 0 | 0 |
| P. Halibut Ad | 4.562 | 0.223 | 0.19 | 1.096 | 0.704 | 0 | 0 |
| YF. Sole Juv | 3.513 | 0.425 | 0.244 | 1.531 | 0.785 | 0 | 0 |

| Group | Trophic Level | Biomass (tkm$^{-2}$) | *Mortality* or Production / Biomass (year$^{-1}$) | Consumption / Biomass (year$^{-1}$) | Ecotrophic Efficiency | Biomass Accumulation (tkm$^{-2}$) | Biomass Accumulation year$^{-1}$) |
|---|---|---|---|---|---|---|---|
| YF. Sole Ad | 3.543 | 1.764 | 0.174 | 0.603 | 0.991 | 0 | 0 |
| FH. Sole Juv | 3.55 | 0.147 | 0.394 | 4.145 | 0.85 | 0 | 0 |
| FH. Sole Ad | 3.676 | 2.091 | 0.26 | 1.222 | 0.126 | 0 | 0 |
| N. Rock sole Juv | 3.505 | 0.157 | 0.292 | 3.482 | 0.761 | 0 | 0 |
| N. Rock sole Ad | 3.653 | 3.761 | 0.232 | 1.122 | 0.378 | 0 | 0 |
| AK Plaice | 3.51 | 1.068 | 0.2 | 2 | 0.52 | 0 | 0 |
| Dover Sole | 3.71 | 0 | 0.2 | 2 | 0.389 | 0 | 0 |
| Rex Sole | 3.725 | 0.041 | 0.2 | 2 | 0.475 | 0 | 0 |
| Misc. Flatfish | 3.706 | 0.222 | 0.2 | 2 | 0.819 | 0 | 0 |
| Alaska skate | 4.185 | 0.68 | 0.2 | 2 | 0.192 | 0 | 0 |
| Other skates | 4.425 | 0.093 | 0.2 | 2 | 0.232 | 0 | 0 |
| Sablefish Juv | 3.637 | 0.004 | 0.266 | 2.964 | 0.954 | 0 | 0 |
| Sablefish Ad | 4.519 | 0.054 | 0.19 | 1.002 | 0.303 | 0 | 0 |
| Eelpouts | 3.578 | 2.372 | 0.4 | 2 | 0.946 | 0 | 0 |
| Grenadiers | 4.293 | 0.968 | 0.15 | 2 | 0.268 | 0 | 0 |
| Misc. fish deep | 4.293 | 0.008 | 0.2 | 2 | 0.806 | 0 | 0 |
| POP | 3.548 | 0.167 | 0.1 | 2 | 0.845 | 0 | 0 |
| Sharpchin Rock | 3.769 | 0.002 | 0.1 | 2 | 0.798 | 0 | 0 |
| Northern Rock | 3.603 | 0.028 | 0.1 | 2 | 0.798 | 0 | 0 |
| Dusky Rock | 3.517 | 0.001 | 0.1 | 2 | 0.516 | 0 | 0 |
| Shortraker Rock | 3.864 | 0.009 | 0.1 | 2 | 0.447 | 0 | 0 |
| Rougheye Rock | 4.297 | 0.004 | 0.1 | 2 | 0.611 | 0 | 0 |
| Shortspine Thorns | 3.576 | 0.005 | 0.15 | 0.5 | 0.322 | 0 | 0 |
| Other Sebastes | 3.769 | 0.011 | 0.1 | 2 | 0.798 | 0 | 0 |
| Atka mackerel Juv | 3.523 | 0.071 | 0.9 | 12.019 | 0 | 0 | 0 |
| Atka mackerel Ad | 3.53 | 0.169 | 0.35 | 4.991 | 0.599 | 0 | 0 |
| Greenlings | 4.244 | 0.001 | 0.4 | 2 | 0.874 | 0 | 0 |
| Lg. Sculpins | 4.029 | 0.54 | 0.4 | 2 | 0.065 | 0 | 0 |
| Other sculpins | 3.879 | 1.137 | 0.4 | 2 | 0.88 | 0 | 0 |
| Misc. fish shallow | 3.675 | 1.168 | 0.4 | 2 | 0.838 | 0 | 0 |
| Octopi | 3.818 | 0.192 | 0.8 | 3.65 | 0.821 | 0 | 0 |
| Squids | 3.679 | 0.927 | 3.2 | 10.67 | 0.95 | 0 | 0 |
| Salmon returning | 3.827 | 0.164 | 1.65 | 11.6 | 0.975 | 0 | 0 |
| Salmon outgoing | 3.513 | 0.014 | 1.28 | 13.56 | 0.804 | 0 | 0 |
| Bathylagidae | 3.523 | 0.162 | 0.8 | 3.65 | 0.997 | 0 | 0 |
| Myctophidae | 3.523 | 0.826 | 0.8 | 3.65 | 0.986 | 0 | 0 |
| Capelin | 3.523 | 1.245 | 0.8 | 3.65 | 0.817 | 0 | 0 |
| Sandlance | 3.523 | 2.528 | 0.8 | 3.65 | 0.847 | 0 | 0 |
| Eulachon | 3.523 | 0.552 | 0.8 | 3.65 | 0.813 | 0 | 0 |
| Oth. managed forage | 3.523 | 1.051 | 0.8 | 3.65 | 0.839 | 0 | 0 |
| Oth. pelagic smelt | 3.523 | 0.499 | 0.8 | 3.65 | 0.802 | 0 | 0 |
| Bairdi | 3.369 | 0.522 | 1.601 | 3.092 | 0.749 | 0 | 0 |
| King Crab | 3.383 | 0.238 | 0.659 | 3.207 | 0.646 | 0 | 0 |
| Opilio | 3.281 | 2.179 | 1.295 | 3.117 | 0.524 | 0 | 0 |
| Pandalidae | 2.908 | 6.742 | 0.576 | 2.409 | 0.938 | 0 | 0 |
| NP shrimp | 2.908 | 12.702 | 0.576 | 2.409 | 0.806 | 0 | 0 |
| Sea stars | 3.53 | 2.471 | 1.21 | 6.05 | 0.005 | 0 | 0 |
| Brittle stars | 2.15 | 2.95 | 1.21 | 6.05 | 0.786 | 0 | 0 |
| Urchins dollars cucumbers | 2 | 1.167 | 0.61 | 3.05 | 0.595 | 0 | 0 |
| Snails | 2.905 | 0.807 | 1.81 | 9.05 | 0.784 | 0 | 0 |
| Hermit crabs | 3.131 | 1.774 | 0.82 | 4.1 | 0.79 | 0 | 0 |
| Misc. crabs | 3.131 | 0.71 | 0.82 | 4.1 | 0.814 | 0 | 0 |

| Group | Trophic Level | Biomass ($tkm^{-2}$) | *Mortality* or Production / Biomass ($year^{-1}$) | Consumption / Biomass ($year^{-1}$) | Ecotrophic Efficiency | Biomass Accumulation ($tkm^{-2}$) | Biomass Accumulation $year^{-1}$) |
|---|---|---|---|---|---|---|---|
| Misc. Crustacean | 2.5 | 8.886 | 7.4 | 37 | 0.8 | 0 | 0 |
| Benthic Amphipods | 2.5 | 12.796 | 7.4 | 37 | 0.801 | 0 | 0 |
| Anemones | 2.525 | 0.11 | 1 | 5 | 0.115 | 0 | 0 |
| Corals | 2.525 | 0.013 | 0.046 | 0.23 | 0.511 | 0 | 0 |
| Hydroids | 2.525 | 0.236 | 1 | 5 | 0.799 | 0 | 0 |
| Urochordata | 2.525 | 0.354 | 3.58 | 17.9 | 0.026 | 0 | 0 |
| Sea Pens | 2.525 | 0.013 | 0.092 | 0.461 | 0.003 | 0 | 0 |
| Sponges | 2.525 | 0.054 | 1 | 5 | 0.511 | 0 | 0 |
| Bivalves | 2.525 | 61.873 | 1.3 | 6.5 | 0.341 | 0 | 0 |
| Polychaetes | 2.5 | 21.687 | 2.97 | 14.85 | 0.204 | 0 | 0 |
| Misc. worms | 2.5 | 3.669 | 2.23 | 11.15 | 0.748 | 0 | 0 |
| Scyphozoid Jellies | 3.429 | 0.338 | 0.88 | 3 | 0.55 | 0 | 0 |
| Fish Larvae | 2.525 | 0.012 | 5.475 | 15.643 | 0.801 | 0 | 0 |
| Chaetognaths | 2.906 | 0.705 | 5.475 | 15.643 | 0.832 | 0 | 0 |
| Euphausiids | 2.525 | 17.831 | 5.475 | 15.643 | 0.905 | 0 | 0 |
| Mysids | 2.525 | 1.5 | 5.475 | 15.643 | 0.82 | 0 | 0 |
| Pelagic Amphipods | 2.525 | 1.695 | 2.5 | 7.143 | 0.848 | 0 | 0 |
| Gelatinous filter feeders | 2.525 | 0.774 | 5.475 | 15.643 | 0.842 | 0 | 0 |
| Pteropods | 2.525 | 0.232 | 5.475 | 15.643 | 0.795 | 0 | 0 |
| Copepods | 2.5 | 26.857 | 6 | 27.74 | 0.831 | 0 | 0 |
| Pelagic microbes | 2 | 45 | 36.5 | 104.286 | 0.258 | 0 | 0 |
| Benthic microbes | 2 | 22.061 | 36.5 | 104.286 | 0.8 | 0 | 0 |
| Macroalgae | 1 | 0.7501763 | 4 | | 0.8000059 | 0 | 0 |
| Lg Phytoplankton | 1 | 4.350356 | 101.794 | | 0.8 | 0 | 0 |
| Sm Phytoplankton | 1 | 39.5039 | 110.9194 | | 0.8000001 | 0 | 0 |
| Outside Production | 1 | 4 | 1 | | 0 | 0 | 0 |
| Discards | 1 | 0.6062543 | | | 0.08720814 | 0 | 0 |
| Offal | 1 | 2.054416 | | | 0.5810784 | 0 | 0 |
| Pelagic Detritus | 1 | 1 | | | 0.8773113 | 196.8834 | 196.8834 |
| Benthic Detritus | 1 | 1 | | | 0.92233 | 267.1387 | 267.1387 |
| Outside Detritus | 1 | 1.00E-05 | | | 0 | 0 | 0 |

**Table S3:** Diet matrix for the East Bering Sea Ecopath model (converted from the Rpath version of the model)

| Group | Transient Killers | Sperm and Beaked Whales | Resident Killers | Porpoises | Belugas | Gray Whales | Humpbacks | Fin Whales | Sei Whales | Right Whales | Minke Whales | Bowhead Whales | Sea Otters | Walrus Bd Seals | N Fur Seal Juv | N Fur Seal Ad | Steller Sea Lions Juv | Steller Sea Lions Ad | Resident Seals | Wintering Seals | Shearwater | Murres | Kittiwakes | Auklets | Puffins | Fulmars | Storm Petrels | Cormorants | Gulls | Albatross Jaeger |
|---|---|---|---|---|---|---|---|---|---|---|---|---|---|---|---|---|---|---|---|---|---|---|---|---|---|---|---|---|---|---|
| Transient KIllers | 0 | 0 | 0 | 0 | 0 | 0 | 0 | 0 | 0 | 0 | 0 | 0 | 0 | 0 | 0 | 0 | 0 | 0 | 0 | 0 | 0 | 0 | 0 | 0 | 0 | 0 | 0 | 0 | 0 | 0 |
| Sperm and Beaked Whales | 0 | 0 | 0 | 0 | 0 | 0 | 0 | 0 | 0 | 0 | 0 | 0 | 0 | 0 | 0 | 0 | 0 | 0 | 0 | 0 | 0 | 0 | 0 | 0 | 0 | 0 | 0 | 0 | 0 | 0 |
| Resident Killers | 0 | 0 | 0 | 0 | 0 | 0 | 0 | 0 | 0 | 0 | 0 | 0 | 0 | 0 | 0 | 0 | 0 | 0 | 0 | 0 | 0 | 0 | 0 | 0 | 0 | 0 | 0 | 0 | 0 | 0 |
| Porpoises | 0.006 | 0 | 0 | 0 | 0 | 0 | 0 | 0 | 0 | 0 | 0 | 0 | 0 | 0 | 0 | 0 | 0 | 0 | 0 | 0 | 0 | 0 | 0 | 0 | 0 | 0 | 0 | 0 | 0 | 0 |
| Belugas | 0.02 | 0 | 0 | 0 | 0 | 0 | 0 | 0 | 0 | 0 | 0 | 0 | 0 | 0 | 0 | 0 | 0 | 0 | 0 | 0 | 0 | 0 | 0 | 0 | 0 | 0 | 0 | 0 | 0 | 0 |
| Gray Whales | 0.054 | 0 | 0 | 0 | 0 | 0 | 0 | 0 | 0 | 0 | 0 | 0 | 0 | 0 | 0 | 0 | 0 | 0 | 0 | 0 | 0 | 0 | 0 | 0 | 0 | 0 | 0 | 0 | 0 | 0 |
| Humpbacks | 0.02 | 0 | 0 | 0 | 0 | 0 | 0 | 0 | 0 | 0 | 0 | 0 | 0 | 0 | 0 | 0 | 0 | 0 | 0 | 0 | 0 | 0 | 0 | 0 | 0 | 0 | 0 | 0 | 0 | 0 |
| Fin Whales | 0.749 | 0 | 0 | 0 | 0 | 0 | 0 | 0 | 0 | 0 | 0 | 0 | 0 | 0 | 0 | 0 | 0 | 0 | 0 | 0 | 0 | 0 | 0 | 0 | 0 | 0 | 0 | 0 | 0 | 0 |
| Sei Whales | 0.01 | 0 | 0 | 0 | 0 | 0 | 0 | 0 | 0 | 0 | 0 | 0 | 0 | 0 | 0 | 0 | 0 | 0 | 0 | 0 | 0 | 0 | 0 | 0 | 0 | 0 | 0 | 0 | 0 | 0 |
| Right Whales | 0.006 | 0 | 0 | 0 | 0 | 0 | 0 | 0 | 0 | 0 | 0 | 0 | 0 | 0 | 0 | 0 | 0 | 0 | 0 | 0 | 0 | 0 | 0 | 0 | 0 | 0 | 0 | 0 | 0 | 0 |
| Minke Whales | 0.041 | 0 | 0 | 0 | 0 | 0 | 0 | 0 | 0 | 0 | 0 | 0 | 0 | 0 | 0 | 0 | 0 | 0 | 0 | 0 | 0 | 0 | 0 | 0 | 0 | 0 | 0 | 0 | 0 | 0 |
| Bowhead Whales | 0.012 | 0 | 0 | 0 | 0 | 0 | 0 | 0 | 0 | 0 | 0 | 0 | 0 | 0 | 0 | 0 | 0 | 0 | 0 | 0 | 0 | 0 | 0 | 0 | 0 | 0 | 0 | 0 | 0 | 0 |
| Sea Otters | 0.001 | 0 | 0 | 0 | 0 | 0 | 0 | 0 | 0 | 0 | 0 | 0 | 0 | 0 | 0 | 0 | 0 | 0 | 0 | 0 | 0 | 0 | 0 | 0 | 0 | 0 | 0 | 0 | 0 | 0 |
| Walrus Bd Seals | 0 | 0 | 0 | 0 | 0 | 0 | 0 | 0 | 0 | 0 | 0 | 0 | 0 | 0 | 0 | 0 | 0 | 0 | 0 | 0 | 0 | 0 | 0 | 0 | 0 | 0 | 0 | 0 | 0 | 0 |
| N Fur Seal Juv | 0.003 | 0 | 0 | 0 | 0 | 0 | 0 | 0 | 0 | 0 | 0 | 0 | 0 | 0 | 0 | 0 | 0 | 0 | 0 | 0 | 0 | 0 | 0 | 0 | 0 | 0 | 0 | 0 | 0 | 0 |
| N Fur Seal Ad | 0.054 | 0 | 0 | 0 | 0 | 0 | 0 | 0 | 0 | 0 | 0 | 0 | 0 | 0 | 0 | 0 | 0 | 0 | 0 | 0 | 0 | 0 | 0 | 0 | 0 | 0 | 0 | 0 | 0 | 0 |
| Steller Sea Lions Juv | 0 | 0 | 0 | 0 | 0 | 0 | 0 | 0 | 0 | 0 | 0 | 0 | 0 | 0 | 0 | 0 | 0 | 0 | 0 | 0 | 0 | 0 | 0 | 0 | 0 | 0 | 0 | 0 | 0 | 0 |
| Steller Sea Lions Ad | 0 | 0 | 0 | 0 | 0 | 0 | 0 | 0 | 0 | 0 | 0 | 0 | 0 | 0 | 0 | 0 | 0 | 0 | 0 | 0 | 0 | 0 | 0 | 0 | 0 | 0 | 0 | 0 | 0 | 0 |
| Resident Seals | 0.023 | 0 | 0 | 0 | 0 | 0 | 0 | 0 | 0 | 0 | 0 | 0 | 0 | 0 | 0 | 0 | 0 | 0 | 0 | 0 | 0 | 0 | 0 | 0 | 0 | 0 | 0 | 0 | 0 | 0 |
| Wintering Seals | 0 | 0 | 0 | 0 | 0 | 0 | 0 | 0 | 0 | 0 | 0 | 0 | 0 | 0.001 | 0 | 0 | 0 | 0 | 0 | 0 | 0 | 0 | 0 | 0 | 0 | 0 | 0 | 0 | 0 | 0 |
| Shearwater | 0 | 0 | 0 | 0 | 0 | 0 | 0 | 0 | 0 | 0 | 0 | 0 | 0 | 0 | 0 | 0 | 0 | 0 | 0 | 0 | 0 | 0 | 0 | 0 | 0 | 0 | 0 | 0 | 0 | 0 |
| Murres | 0 | 0 | 0 | 0 | 0 | 0 | 0 | 0 | 0 | 0 | 0 | 0 | 0 | 0 | 0 | 0 | 0 | 0 | 0 | 0 | 0 | 0 | 0 | 0 | 0 | 0.001 | 0 | 0 | 0 | 0 |
| Kittiwakes | 0 | 0 | 0 | 0 | 0 | 0 | 0 | 0 | 0 | 0 | 0 | 0 | 0 | 0 | 0 | 0 | 0 | 0 | 0 | 0 | 0 | 0 | 0 | 0 | 0 | 0 | 0 | 0 | 0 | 0 |
| Auklets | 0 | 0 | 0 | 0 | 0 | 0 | 0 | 0 | 0 | 0 | 0 | 0 | 0 | 0 | 0 | 0 | 0 | 0 | 0 | 0 | 0 | 0 | 0 | 0 | 0 | 0 | 0 | 0 | 0 | 0 |
| Puffins | 0 | 0 | 0 | 0 | 0 | 0 | 0 | 0 | 0 | 0 | 0 | 0 | 0 | 0 | 0 | 0 | 0 | 0 | 0 | 0 | 0 | 0 | 0 | 0 | 0 | 0 | 0 | 0 | 0 | 0 |
| Fulmars | 0 | 0 | 0 | 0 | 0 | 0 | 0 | 0 | 0 | 0 | 0 | 0 | 0 | 0 | 0 | 0 | 0 | 0 | 0 | 0 | 0 | 0 | 0 | 0 | 0 | 0 | 0 | 0 | 0 | 0 |
| Storm Petrels | 0 | 0 | 0 | 0 | 0 | 0 | 0 | 0 | 0 | 0 | 0 | 0 | 0 | 0 | 0 | 0 | 0 | 0 | 0 | 0 | 0 | 0 | 0 | 0 | 0 | 0 | 0 | 0 | 0 | 0 |
| Cormorants | 0 | 0 | 0 | 0 | 0 | 0 | 0 | 0 | 0 | 0 | 0 | 0 | 0 | 0 | 0 | 0 | 0 | 0 | 0 | 0 | 0 | 0 | 0 | 0 | 0 | 0 | 0 | 0 | 0 | 0 |
| Gulls | 0 | 0 | 0 | 0 | 0 | 0 | 0 | 0 | 0 | 0 | 0 | 0 | 0 | 0 | 0 | 0 | 0 | 0 | 0 | 0 | 0 | 0 | 0 | 0 | 0 | 0 | 0 | 0 | 0 | 0 |
| Albatross Jaeger | 0 | 0 | 0 | 0 | 0 | 0 | 0 | 0 | 0 | 0 | 0 | 0 | 0 | 0 | 0 | 0 | 0 | 0 | 0 | 0 | 0 | 0 | 0 | 0 | 0 | 0 | 0 | 0 | 0 | 0 |
| Sleeper shark | 0 | 0.001 | 0 | 0 | 0 | 0 | 0 | 0 | 0 | 0 | 0 | 0 | 0 | 0 | 0 | 0 | 0 | 0 | 0 | 0 | 0 | 0 | 0 | 0 | 0 | 0 | 0 | 0 | 0 | 0 |
| W. Pollock Juv | 0 | 0 | 0.054 | 0.114 | 0.227 | 0 | 0.14 | 0.071 | 0.099 | 0 | 0.216 | 0 | 0.018 | 0 | 0.305 | 0.305 | 0.035 | 0 | 0.017 | 0.052 | 0 | 0.27 | 0.223 | 0.003 | 0.111 | 0.293 | 0.016 | 0.024 | 0.002 | 0.173 |

| Group | Transient Killers | Sperm and Beaked Whales | Resident Killers | Porpoises | Belugas | Gray Whales | Humpbacks | Fin Whales | Sei Whales | Right Whales | Minke Whales | Bowhead Whales | Sea Otters | Walrus Bd Seals | N Fur Seal Juv | N Fur Seal Ad | Steller Sea Lions Juv | Steller Sea Lions Ad | Resident Seals | Wintering Seals | Shearwater | Murres | Kittiwakes | Auklets | Puffins | Fulmars | Storm Petrels | Cormorants | Gulls | Albatross Jaeger |
|---|---|---|---|---|---|---|---|---|---|---|---|---|---|---|---|---|---|---|---|---|---|---|---|---|---|---|---|---|---|---|
| W. Pollock Ad | 0 | 0 | 0.447 | 0 | 0 | 0 | 0 | 0 | 0 | 0 | 0 | 0 | 0.149 | 0 | 0 | 0 | 0.295 | 0.33 | 0.145 | 0.436 | 0 | 0 | 0 | 0 | 0 | 0 | 0 | 0 | 0 | 0 |
| P. Cod Juv | 0 | 0 | 0.002 | 0.005 | 0.01 | 0 | 0 | 0 | 0 | 0 | 0 | 0 | 0.001 | 0 | 0.007 | 0.007 | 0.07 | 0.07 | 0.045 | 0.005 | 0 | 0.2 | 0.01 | 0 | 0.005 | 0.013 | 0.001 | 0.001 | 0 | 0.008 |
| P. Cod Ad | 0 | 0.058 | 0.06 | 0 | 0 | 0 | 0 | 0 | 0 | 0 | 0 | 0 | 0 | 0 | 0 | 0 | 0 | 0 | 0 | 0 | 0 | 0 | 0 | 0 | 0 | 0 | 0 | 0 | 0 | 0 |
| Herring Juv | 0 | 0 | 0 | 0.001 | 0.001 | 0 | 0.001 | 0 | 0.001 | 0 | 0.001 | 0 | 0 | 0 | 0.061 | 0.061 | 0.03 | 0.03 | 0.028 | 0.044 | 0 | 0 | 0.003 | 0 | 0 | 0 | 0 | 0.127 | 0 | 0.001 |
| Herring Ad | 0 | 0 | 0.015 | 0 | 0 | 0 | 0 | 0 | 0 | 0 | 0 | 0 | 0.005 | 0 | 0 | 0 | 0 | 0 | 0 | 0 | 0 | 0 | 0 | 0 | 0 | 0 | 0 | 0 | 0 | 0 |
| Arrowtooth Juv | 0 | 0 | 0.002 | 0 | 0 | 0 | 0 | 0 | 0 | 0 | 0 | 0 | 0 | 0 | 0 | 0 | 0 | 0 | 0 | 0 | 0 | 0 | 0 | 0 | 0 | 0 | 0 | 0 | 0 | 0 |
| Arrowtooth Ad | 0 | 0 | 0.023 | 0 | 0 | 0 | 0 | 0 | 0 | 0 | 0 | 0 | 0 | 0 | 0 | 0 | 0 | 0 | 0 | 0 | 0 | 0 | 0 | 0 | 0 | 0 | 0 | 0 | 0 | 0 |
| Kamchatka fl. Juv | 0 | 0 | 0 | 0 | 0 | 0 | 0 | 0 | 0 | 0 | 0 | 0 | 0 | 0 | 0 | 0 | 0 | 0 | 0 | 0 | 0 | 0 | 0 | 0 | 0 | 0 | 0 | 0 | 0 | 0 |
| Kamchatka fl. Ad | 0 | 0 | 0.001 | 0 | 0 | 0 | 0 | 0 | 0 | 0 | 0 | 0 | 0 | 0 | 0 | 0 | 0 | 0 | 0 | 0 | 0 | 0 | 0 | 0 | 0 | 0 | 0 | 0 | 0 | 0 |
| Gr. Turbot Juv | 0 | 0 | 0.001 | 0 | 0 | 0 | 0 | 0 | 0 | 0 | 0 | 0 | 0 | 0 | 0 | 0 | 0 | 0 | 0 | 0 | 0 | 0 | 0 | 0 | 0 | 0 | 0 | 0 | 0 | 0 |
| Gr. Turbot Ad | 0 | 0 | 0.008 | 0 | 0 | 0 | 0 | 0 | 0 | 0 | 0 | 0 | 0 | 0 | 0 | 0 | 0 | 0 | 0 | 0 | 0 | 0 | 0 | 0 | 0 | 0 | 0 | 0 | 0 | 0 |
| P. Halibut Juv | 0 | 0 | 0.001 | 0 | 0 | 0 | 0 | 0 | 0 | 0 | 0 | 0 | 0 | 0 | 0 | 0 | 0 | 0 | 0 | 0 | 0 | 0 | 0 | 0 | 0 | 0 | 0 | 0 | 0 | 0 |
| P. Halibut Ad | 0 | 0 | 0.005 | 0 | 0 | 0 | 0 | 0 | 0 | 0 | 0 | 0 | 0 | 0 | 0 | 0 | 0 | 0 | 0 | 0 | 0 | 0 | 0 | 0 | 0 | 0 | 0 | 0 | 0 | 0 |
| YF. Sole Juv | 0 | 0 | 0.012 | 0 | 0 | 0 | 0 | 0 | 0 | 0 | 0 | 0 | 0 | 0 | 0.018 | 0.018 | 0.027 | 0.027 | 0.02 | 0.009 | 0 | 0 | 0 | 0 | 0 | 0 | 0 | 0 | 0 | 0 |
| YF. Sole Ad | 0 | 0 | 0.117 | 0 | 0 | 0 | 0 | 0 | 0 | 0 | 0 | 0 | 0 | 0 | 0 | 0 | 0 | 0 | 0 | 0 | 0 | 0 | 0 | 0 | 0 | 0 | 0 | 0 | 0 | 0 |
| FH. Sole Juv | 0 | 0 | 0.003 | 0 | 0 | 0 | 0 | 0 | 0 | 0 | 0 | 0 | 0 | 0 | 0.004 | 0.004 | 0.007 | 0.007 | 0.005 | 0.002 | 0 | 0 | 0 | 0 | 0 | 0 | 0 | 0 | 0 | 0 |
| FH. Sole Ad | 0 | 0 | 0.029 | 0 | 0 | 0 | 0 | 0 | 0 | 0 | 0 | 0 | 0 | 0 | 0 | 0 | 0 | 0 | 0 | 0 | 0 | 0 | 0 | 0 | 0 | 0 | 0 | 0 | 0 | 0 |
| N. Rock sole Juv | 0 | 0 | 0.008 | 0 | 0 | 0 | 0 | 0 | 0 | 0 | 0 | 0 | 0 | 0 | 0.012 | 0.012 | 0.018 | 0.018 | 0.013 | 0.006 | 0 | 0 | 0 | 0 | 0 | 0 | 0 | 0 | 0 | 0 |
| N. Rock sole Ad | 0 | 0 | 0.078 | 0 | 0 | 0 | 0 | 0 | 0 | 0 | 0 | 0 | 0 | 0 | 0 | 0 | 0 | 0 | 0 | 0 | 0 | 0 | 0 | 0 | 0 | 0 | 0 | 0 | 0 | 0 |
| AK Plaice | 0 | 0 | 0.026 | 0 | 0 | 0 | 0 | 0 | 0 | 0 | 0 | 0 | 0 | 0 | 0.04 | 0.04 | 0.059 | 0.059 | 0.044 | 0.019 | 0 | 0 | 0 | 0 | 0 | 0 | 0 | 0 | 0 | 0 |
| Dover Sole | 0 | 0 | 0 | 0 | 0 | 0 | 0 | 0 | 0 | 0 | 0 | 0 | 0 | 0 | 0 | 0 | 0 | 0 | 0 | 0 | 0 | 0 | 0 | 0 | 0 | 0 | 0 | 0 | 0 | 0 |
| Rex Sole | 0 | 0 | 0.001 | 0 | 0 | 0 | 0 | 0 | 0 | 0 | 0 | 0 | 0 | 0 | 0.002 | 0.002 | 0.002 | 0.002 | 0.002 | 0.001 | 0 | 0 | 0 | 0 | 0 | 0 | 0 | 0 | 0 | 0 |
| Misc. Flatfish | 0 | 0 | 0.004 | 0 | 0 | 0 | 0 | 0 | 0 | 0 | 0 | 0 | 0 | 0 | 0.006 | 0.006 | 0.008 | 0.008 | 0.006 | 0.003 | 0 | 0 | 0 | 0 | 0 | 0 | 0 | 0 | 0 | 0 |
| Alaska skate | 0 | 0.016 | 0 | 0 | 0 | 0 | 0 | 0 | 0 | 0 | 0 | 0 | 0 | 0 | 0 | 0 | 0 | 0 | 0 | 0 | 0 | 0 | 0 | 0 | 0 | 0 | 0 | 0 | 0 | 0 |
| Other skates | 0 | 0.002 | 0 | 0 | 0 | 0 | 0 | 0 | 0 | 0 | 0 | 0 | 0 | 0 | 0 | 0 | 0 | 0 | 0 | 0 | 0 | 0 | 0 | 0 | 0 | 0 | 0 | 0 | 0 | 0 |
| Sablefish Juv | 0 | 0 | 0 | 0 | 0 | 0 | 0 | 0 | 0 | 0 | 0 | 0 | 0 | 0 | 0.005 | 0 | 0 | 0 | 0 | 0 | 0 | 0 | 0 | 0 | 0 | 0 | 0 | 0 | 0 | 0 |
| Sablefish Ad | 0 | 0.001 | 0 | 0 | 0 | 0 | 0 | 0 | 0 | 0 | 0 | 0 | 0 | 0 | 0 | 0 | 0 | 0 | 0 | 0 | 0 | 0 | 0 | 0 | 0 | 0 | 0 | 0 | 0 | 0 |
| Eelpouts | 0 | 0.044 | 0 | 0.096 | 0.192 | 0 | 0 | 0 | 0 | 0 | 0 | 0 | 0 | 0 | 0 | 0 | 0.01 | 0.01 | 0.057 | 0.06 | 0 | 0 | 0 | 0 | 0 | 0 | 0 | 0 | 0 | 0 |
| Grenadiers | 0 | 0.023 | 0 | 0 | 0 | 0 | 0 | 0 | 0 | 0 | 0 | 0 | 0 | 0 | 0 | 0 | 0 | 0 | 0 | 0 | 0 | 0 | 0 | 0 | 0 | 0 | 0 | 0 | 0 | 0 |
| Misc. fish deep | 0 | 0 | 0 | 0 | 0 | 0 | 0 | 0 | 0 | 0 | 0 | 0 | 0 | 0 | 0 | 0 | 0 | 0 | 0 | 0 | 0 | 0 | 0 | 0 | 0 | 0 | 0 | 0 | 0 | 0 |
| POP | 0 | 0.004 | 0 | 0 | 0 | 0 | 0 | 0 | 0 | 0 | 0 | 0 | 0 | 0 | 0.001 | 0.001 | 0.003 | 0.003 | 0.003 | 0 | 0 | 0.001 | 0.001 | 0.001 | 0.001 | 0.001 | 0.001 | 0.001 | 0.001 | 0 |
| Sharpchin Rock | 0 | 0 | 0 | 0 | 0 | 0 | 0 | 0 | 0 | 0 | 0 | 0 | 0 | 0 | 0 | 0 | 0 | 0 | 0 | 0 | 0 | 0 | 0 | 0 | 0 | 0 | 0 | 0 | 0 | 0 |
| Northern Rock | 0 | 0.001 | 0 | 0 | 0 | 0 | 0 | 0 | 0 | 0 | 0 | 0 | 0 | 0 | 0 | 0 | 0 | 0 | 0 | 0 | 0 | 0 | 0 | 0 | 0 | 0 | 0 | 0 | 0 | 0 |
| Dusky Rock | 0 | 0 | 0 | 0 | 0 | 0 | 0 | 0 | 0 | 0 | 0 | 0 | 0 | 0 | 0 | 0 | 0 | 0 | 0 | 0 | 0 | 0 | 0 | 0 | 0 | 0 | 0 | 0 | 0 | 0 |
| Shortraker Rock | 0 | 0 | 0 | 0 | 0 | 0 | 0 | 0 | 0 | 0 | 0 | 0 | 0 | 0 | 0 | 0 | 0 | 0 | 0 | 0 | 0 | 0 | 0 | 0 | 0 | 0 | 0 | 0 | 0 | 0 |

| Group | Transient Killers | Sperm and Beaked Whales | Resident Killers | Porpoises | Belugas | Gray Whales | Humpbacks | Fin Whales | Sei Whales | Right Whales | Minke Whales | Bowhead Whales | Sea Otters | Walrus Bd Seals | N Fur Seal Juv | N Fur Seal Ad | Steller Sea Lions Juv | Steller Sea Lions Ad | Resident Seals | Wintering Seals | Shearwater | Murres | Kittiwakes | Auklets | Puffins | Fulmars | Storm Petrels | Cormorants | Gulls | Albatross Jaeger |
|---|---|---|---|---|---|---|---|---|---|---|---|---|---|---|---|---|---|---|---|---|---|---|---|---|---|---|---|---|---|---|
| Rougheye Rock | 0 | 0 | 0 | 0 | 0 | 0 | 0 | 0 | 0 | 0 | 0 | 0 | 0 | 0 | 0 | 0 | 0 | 0 | 0 | 0 | 0 | 0 | 0 | 0 | 0 | 0 | 0 | 0 | 0 | 0 |
| Shortspine Thorns | 0 | 0 | 0 | 0 | 0 | 0 | 0 | 0 | 0 | 0 | 0 | 0 | 0 | 0 | 0 | 0 | 0 | 0 | 0 | 0 | 0 | 0 | 0 | 0 | 0 | 0 | 0 | 0 | 0 | 0 |
| Other Sebastes | 0 | 0 | 0 | 0 | 0 | 0 | 0 | 0 | 0 | 0 | 0 | 0 | 0 | 0 | 0 | 0 | 0 | 0 | 0 | 0 | 0 | 0 | 0 | 0 | 0 | 0 | 0 | 0 | 0 | 0 |
| Atka mackerel Juv | 0 | 0 | 0 | 0 | 0 | 0 | 0 | 0 | 0 | 0 | 0 | 0 | 0 | 0 | 0 | 0 | 0 | 0 | 0 | 0 | 0 | 0 | 0 | 0 | 0 | 0 | 0 | 0 | 0 | 0 |
| Atka mackerel Ad | 0 | 0 | 0 | 0 | 0 | 0 | 0 | 0 | 0 | 0 | 0 | 0 | 0 | 0 | 0.005 | 0.005 | 0.001 | 0.001 | 0.011 | 0 | 0 | 0 | 0 | 0 | 0 | 0 | 0 | 0 | 0 | 0 |
| Greenlings | 0 | 0 | 0 | 0 | 0 | 0 | 0 | 0 | 0 | 0 | 0 | 0 | 0 | 0 | 0 | 0 | 0 | 0 | 0 | 0 | 0 | 0 | 0 | 0 | 0 | 0 | 0 | 0 | 0 | 0 |
| Lg. Sculpins | 0 | 0 | 0 | 0 | 0 | 0 | 0 | 0 | 0 | 0 | 0 | 0 | 0 | 0 | 0 | 0 | 0 | 0 | 0 | 0 | 0 | 0 | 0 | 0 | 0 | 0 | 0 | 0 | 0 | 0 |
| Other sculpins | 0 | 0 | 0 | 0.04 | 0.08 | 0 | 0 | 0 | 0 | 0 | 0 | 0 | 0 | 0 | 0.001 | 0.001 | 0.06 | 0.06 | 0.057 | 0.079 | 0 | 0 | 0 | 0 | 0 | 0 | 0 | 0 | 0 | 0 |
| Misc. fish shallow | 0 | 0 | 0 | 0.026 | 0.053 | 0 | 0.033 | 0.016 | 0 | 0 | 0 | 0 | 0 | 0 | 0 | 0 | 0.03 | 0.03 | 0.153 | 0.099 | 0 | 0 | 0 | 0 | 0 | 0 | 0 | 0 | 0 | 0 |
| Octopi | 0 | 0 | 0 | 0 | 0.009 | 0 | 0 | 0 | 0 | 0 | 0 | 0 | 0 | 0.01 | 0.001 | 0.001 | 0.181 | 0.181 | 0.113 | 0.01 | 0 | 0 | 0 | 0 | 0 | 0 | 0 | 0 | 0 | 0 |
| Squids | 0 | 0.85 | 0 | 0.5 | 0.067 | 0 | 0.005 | 0 | 0.05 | 0 | 0.02 | 0 | 0.05 | 0 | 0.305 | 0.305 | 0.03 | 0.03 | 0.028 | 0 | 0.26 | 0.034 | 0.007 | 0.001 | 0.038 | 0.585 | 0.606 | 0 | 0 | 0.5 |
| Salmon returning | 0 | 0 | 0.004 | 0.008 | 0.017 | 0 | 0.01 | 0.005 | 0 | 0 | 0.016 | 0 | 0 | 0 | 0.01 | 0.01 | 0.005 | 0.005 | 0.003 | 0 | 0 | 0 | 0 | 0 | 0 | 0 | 0 | 0 | 0 | 0 |
| Salmon outgoing | 0 | 0 | 0 | 0 | 0 | 0 | 0 | 0 | 0 | 0 | 0 | 0 | 0 | 0 | 0.01 | 0.01 | 0.005 | 0.005 | 0.003 | 0 | 0 | 0 | 0 | 0 | 0 | 0 | 0 | 0 | 0 | 0 |
| Bathylagidae | 0 | 0 | 0.004 | 0.008 | 0 | 0 | 0 | 0 | 0 | 0 | 0 | 0 | 0 | 0 | 0 | 0 | 0 | 0 | 0 | 0 | 0.014 | 0 | 0 | 0 | 0 | 0 | 0 | 0.025 | 0 | 0.012 |
| Myctophidae | 0 | 0 | 0.014 | 0.029 | 0 | 0 | 0 | 0 | 0 | 0 | 0 | 0 | 0 | 0 | 0.041 | 0.045 | 0 | 0 | 0 | 0 | 0.048 | 0.002 | 0.12 | 0 | 0 | 0 | 0 | 0.089 | 0 | 0.044 |
| Capelin | 0 | 0 | 0.011 | 0.024 | 0.048 | 0 | 0.03 | 0.015 | 0 | 0 | 0.046 | 0 | 0.004 | 0 | 0.156 | 0.156 | 0.06 | 0.06 | 0.062 | 0.084 | 0.406 | 0.052 | 0.071 | 0.008 | 0.099 | 0.01 | 0.006 | 0.074 | 0.134 | 0.037 |
| Sandlance | 0 | 0 | 0.04 | 0.085 | 0.17 | 0 | 0.105 | 0.053 | 0 | 0 | 0.161 | 0 | 0.013 | 0 | 0.01 | 0.01 | 0.04 | 0.04 | 0 | 0 | 0.141 | 0.184 | 0.25 | 0.027 | 0.351 | 0.035 | 0.021 | 0.423 | 0.473 | 0.129 |
| Eulachon | 0 | 0 | 0.008 | 0.016 | 0.032 | 0 | 0.02 | 0.01 | 0 | 0 | 0.03 | 0 | 0.003 | 0 | 0 | 0 | 0 | 0 | 0 | 0 | 0.027 | 0.035 | 0.047 | 0.005 | 0.066 | 0.007 | 0.004 | 0.049 | 0.089 | 0.024 |
| Oth. managed forage | 0 | 0 | 0.016 | 0.033 | 0.067 | 0 | 0.041 | 0.021 | 0 | 0 | 0.063 | 0 | 0.005 | 0 | 0 | 0 | 0 | 0 | 0 | 0 | 0.055 | 0.072 | 0.098 | 0.01 | 0.138 | 0.014 | 0.008 | 0.102 | 0.186 | 0.051 |
| Oth. pelagic smelt | 0 | 0 | 0.007 | 0.014 | 0.028 | 0 | 0.017 | 0.009 | 0 | 0 | 0.027 | 0 | 0.002 | 0 | 0 | 0 | 0 | 0 | 0 | 0 | 0.023 | 0.03 | 0.041 | 0.004 | 0.058 | 0.006 | 0.003 | 0.043 | 0.078 | 0.021 |
| Bairdi | 0 | 0 | 0 | 0 | 0 | 0.001 | 0 | 0 | 0 | 0 | 0 | 0 | 0 | 0.001 | 0 | 0 | 0.001 | 0.001 | 0.001 | 0 | 0 | 0 | 0 | 0 | 0 | 0 | 0 | 0 | 0 | 0 |
| King Crab | 0 | 0 | 0 | 0 | 0 | 0 | 0 | 0 | 0 | 0 | 0 | 0 | 0 | 0 | 0 | 0 | 0 | 0 | 0.001 | 0 | 0 | 0 | 0 | 0 | 0 | 0 | 0 | 0 | 0 | 0 |
| Opilio | 0 | 0 | 0 | 0 | 0 | 0.002 | 0 | 0 | 0 | 0 | 0 | 0.001 | 0 | 0.004 | 0 | 0 | 0.004 | 0.004 | 0.004 | 0 | 0 | 0 | 0 | 0 | 0 | 0 | 0 | 0.001 | 0 | 0 |
| Pandalidae | 0 | 0 | 0 | 0 | 0 | 0.005 | 0 | 0 | 0 | 0 | 0 | 0.002 | 0 | 0.008 | 0 | 0 | 0.004 | 0.004 | 0.064 | 0.02 | 0 | 0 | 0 | 0 | 0 | 0 | 0 | 0.003 | 0 | 0 |
| NP shrimp | 0 | 0 | 0 | 0 | 0 | 0.008 | 0 | 0 | 0 | 0 | 0 | 0.004 | 0 | 0.013 | 0 | 0 | 0.006 | 0.006 | 0.106 | 0.034 | 0 | 0 | 0 | 0 | 0 | 0 | 0 | 0.005 | 0 | 0 |
| Sea stars | 0 | 0 | 0 | 0 | 0 | 0.002 | 0 | 0 | 0 | 0 | 0 | 0.001 | 0 | 0 | 0 | 0 | 0 | 0 | 0 | 0 | 0 | 0 | 0 | 0 | 0 | 0 | 0 | 0 | 0 | 0 |
| Brittle stars | 0 | 0 | 0 | 0 | 0 | 0.002 | 0 | 0 | 0 | 0 | 0 | 0.001 | 0 | 0 | 0 | 0 | 0 | 0 | 0 | 0 | 0 | 0 | 0 | 0 | 0 | 0 | 0 | 0 | 0 | 0 |
| Urchins dollars cucumbers | 0 | 0 | 0 | 0 | 0 | 0.001 | 0 | 0 | 0 | 0 | 0 | 0 | 0.462 | 0 | 0 | 0 | 0 | 0 | 0 | 0 | 0 | 0 | 0 | 0 | 0 | 0 | 0 | 0 | 0 | 0 |
| Snails | 0 | 0 | 0 | 0 | 0 | 0.001 | 0 | 0 | 0 | 0 | 0 | 0 | 0 | 0.061 | 0 | 0 | 0.001 | 0.001 | 0 | 0 | 0 | 0 | 0 | 0 | 0 | 0 | 0 | 0 | 0 | 0 |
| Hermit crabs | 0 | 0 | 0 | 0 | 0 | 0.001 | 0 | 0 | 0 | 0 | 0 | 0.001 | 0 | 0.004 | 0 | 0 | 0.003 | 0.003 | 0.004 | 0.007 | 0 | 0 | 0 | 0 | 0 | 0 | 0 | 0 | 0 | 0 |
| Misc. crabs | 0 | 0 | 0 | 0 | 0 | 0.001 | 0 | 0 | 0 | 0 | 0 | 0 | 0.288 | 0.001 | 0 | 0 | 0.001 | 0.001 | 0.002 | 0.003 | 0 | 0 | 0 | 0 | 0 | 0 | 0 | 0 | 0 | 0 |
| Misc. Crustacean | 0 | 0 | 0 | 0 | 0 | 0.007 | 0 | 0 | 0 | 0 | 0 | 0.003 | 0 | 0 | 0 | 0 | 0 | 0 | 0 | 0 | 0.01 | 0.006 | 0.004 | 0.001 | 0.063 | 0 | 0.001 | 0.004 | 0.001 | 0 |
| Benthic Amphipods | 0 | 0 | 0 | 0 | 0 | 0.9 | 0 | 0 | 0 | 0 | 0 | 0.004 | 0 | 0 | 0 | 0 | 0 | 0 | 0 | 0 | 0 | 0 | 0.005 | 0.001 | 0 | 0 | 0.002 | 0.006 | 0.002 | 0 |
| Anemones | 0 | 0 | 0 | 0 | 0 | 0 | 0 | 0 | 0 | 0 | 0 | 0 | 0 | 0 | 0 | 0 | 0 | 0 | 0 | 0 | 0 | 0 | 0 | 0 | 0 | 0 | 0 | 0 | 0 | 0 |
| Corals | 0 | 0 | 0 | 0 | 0 | 0 | 0 | 0 | 0 | 0 | 0 | 0 | 0 | 0 | 0 | 0 | 0 | 0 | 0 | 0 | 0 | 0 | 0 | 0 | 0 | 0 | 0 | 0 | 0 | 0 |

| Group | Transient Killers | Sperm and Beaked Whales | Resident Killers | Porpoises | Belugas | Gray Whales | Humpbacks | Fin Whales | Sei Whales | Right Whales | Minke Whales | Bowhead Whales | Sea Otters | Walrus Bd Seals | N Fur Seal Juv | N Fur Seal Ad | Steller Sea Lions Juv | Steller Sea Lions Ad | Resident Seals | Wintering Seals | Shearwater | Murres | Kittiwakes | Auklets | Puffins | Fulmars | Storm Petrels | Cormorants | Gulls | Albatross Jaeger |
|---|---|---|---|---|---|---|---|---|---|---|---|---|---|---|---|---|---|---|---|---|---|---|---|---|---|---|---|---|---|---|
| Hydroids | 0 | 0 | 0 | 0 | 0 | 0 | 0 | 0 | 0 | 0 | 0 | 0 | 0 | 0 | 0 | 0 | 0 | 0 | 0 | 0 | 0 | 0 | 0 | 0 | 0 | 0 | 0 | 0 | 0 | 0 |
| Urochordata | 0 | 0 | 0 | 0 | 0 | 0 | 0 | 0 | 0 | 0 | 0 | 0 | 0 | 0 | 0 | 0 | 0 | 0 | 0 | 0 | 0 | 0 | 0 | 0 | 0 | 0 | 0 | 0 | 0 | 0 |
| Sea Pens | 0 | 0 | 0 | 0 | 0 | 0 | 0 | 0 | 0 | 0 | 0 | 0 | 0 | 0 | 0 | 0 | 0 | 0 | 0 | 0 | 0 | 0 | 0 | 0 | 0 | 0 | 0 | 0 | 0 | 0 |
| Sponges | 0 | 0 | 0 | 0 | 0 | 0 | 0 | 0 | 0 | 0 | 0 | 0 | 0 | 0 | 0 | 0 | 0 | 0 | 0 | 0 | 0 | 0 | 0 | 0 | 0 | 0 | 0 | 0 | 0 | 0 |
| Bivalves | 0 | 0 | 0 | 0 | 0 | 0.049 | 0 | 0 | 0 | 0 | 0 | 0.022 | 0 | 0.714 | 0 | 0 | 0.001 | 0.001 | 0 | 0 | 0 | 0 | 0.027 | 0.005 | 0 | 0.001 | 0.008 | 0 | 0.009 | 0 |
| Polychaetes | 0 | 0 | 0 | 0 | 0 | 0.017 | 0 | 0 | 0 | 0 | 0 | 0.008 | 0 | 0 | 0 | 0 | 0 | 0 | 0 | 0 | 0 | 0 | 0.009 | 0.002 | 0 | 0 | 0.003 | 0 | 0.003 | 0 |
| Misc. worms | 0 | 0 | 0 | 0 | 0 | 0.003 | 0 | 0 | 0 | 0 | 0 | 0.001 | 0 | 0.183 | 0 | 0 | 0 | 0 | 0 | 0.026 | 0 | 0 | 0.001 | 0 | 0 | 0 | 0 | 0 | 0 | 0 |
| Scyphozoid Jellies | 0 | 0 | 0 | 0 | 0 | 0 | 0 | 0 | 0 | 0 | 0 | 0 | 0 | 0 | 0 | 0 | 0 | 0 | 0 | 0 | 0 | 0 | 0 | 0 | 0 | 0 | 0 | 0 | 0 | 0 |
| Fish Larvae | 0 | 0 | 0 | 0 | 0 | 0 | 0 | 0 | 0 | 0 | 0 | 0 | 0 | 0 | 0 | 0 | 0 | 0 | 0 | 0 | 0 | 0 | 0 | 0 | 0 | 0 | 0 | 0 | 0 | 0 |
| Chaetognaths | 0 | 0 | 0 | 0 | 0 | 0 | 0.016 | 0.016 | 0.001 | 0.001 | 0.01 | 0.014 | 0 | 0 | 0 | 0 | 0 | 0 | 0 | 0 | 0 | 0 | 0 | 0 | 0 | 0 | 0 | 0 | 0 | 0 |
| Euphausiids | 0 | 0 | 0 | 0 | 0 | 0 | 0.495 | 0.5 | 0.042 | 0.042 | 0.333 | 0.458 | 0 | 0 | 0 | 0 | 0 | 0 | 0 | 0 | 0.007 | 0.043 | 0.031 | 0.355 | 0.026 | 0.013 | 0.122 | 0.009 | 0.003 | 0 |
| Mysids | 0 | 0 | 0 | 0 | 0 | 0 | 0.03 | 0.03 | 0.003 | 0.003 | 0.02 | 0.028 | 0 | 0 | 0 | 0 | 0 | 0 | 0 | 0 | 0 | 0.003 | 0.002 | 0.022 | 0.002 | 0.001 | 0.007 | 0 | 0 | 0 |
| Pelagic Amphipods | 0 | 0 | 0 | 0 | 0 | 0 | 0.047 | 0.047 | 0.004 | 0.004 | 0.032 | 0.043 | 0 | 0 | 0 | 0 | 0 | 0 | 0 | 0 | 0 | 0.004 | 0.003 | 0.034 | 0.003 | 0.001 | 0.012 | 0 | 0 | 0 |
| Gelatinous filter feeders | 0 | 0 | 0 | 0 | 0 | 0 | 0 | 0 | 0 | 0 | 0 | 0 | 0 | 0 | 0 | 0 | 0 | 0 | 0 | 0 | 0 | 0 | 0 | 0 | 0 | 0 | 0 | 0 | 0 | 0 |
| Pteropods | 0 | 0 | 0 | 0 | 0 | 0 | 0.007 | 0.007 | 0.001 | 0.001 | 0.005 | 0.006 | 0 | 0 | 0 | 0 | 0 | 0 | 0 | 0 | 0 | 0 | 0 | 0 | 0 | 0 | 0 | 0 | 0 | 0 |
| Copepods | 0 | 0 | 0 | 0 | 0 | 0 | 0.005 | 0.2 | 0.8 | 0.95 | 0.02 | 0.4 | 0 | 0 | 0 | 0 | 0 | 0 | 0 | 0 | 0.01 | 0.063 | 0.046 | 0.522 | 0.039 | 0.019 | 0.179 | 0.013 | 0.005 | 0 |
| Pelagic microbes | 0 | 0 | 0 | 0 | 0 | 0 | 0 | 0 | 0 | 0 | 0 | 0 | 0 | 0 | 0 | 0 | 0 | 0 | 0 | 0 | 0 | 0 | 0 | 0 | 0 | 0 | 0 | 0 | 0 | 0 |
| Benthic microbes | 0 | 0 | 0 | 0 | 0 | 0 | 0 | 0 | 0 | 0 | 0 | 0 | 0 | 0 | 0 | 0 | 0 | 0 | 0 | 0 | 0 | 0 | 0 | 0 | 0 | 0 | 0 | 0 | 0 | 0 |
| Macroalgae | 0 | 0 | 0 | 0 | 0 | 0 | 0 | 0 | 0 | 0 | 0 | 0 | 0 | 0 | 0 | 0 | 0 | 0 | 0 | 0 | 0 | 0 | 0 | 0 | 0 | 0 | 0 | 0 | 0 | 0 |
| Lg Phytoplankton | 0 | 0 | 0 | 0 | 0 | 0 | 0 | 0 | 0 | 0 | 0 | 0 | 0 | 0 | 0 | 0 | 0 | 0 | 0 | 0 | 0 | 0 | 0 | 0 | 0 | 0 | 0 | 0 | 0 | 0 |
| Sm Phytoplankton | 0 | 0 | 0 | 0 | 0 | 0 | 0 | 0 | 0 | 0 | 0 | 0 | 0 | 0 | 0 | 0 | 0 | 0 | 0 | 0 | 0 | 0 | 0 | 0 | 0 | 0 | 0 | 0 | 0 | 0 |
| Outside Production | 0 | 0 | 0 | 0 | 0 | 0 | 0 | 0 | 0 | 0 | 0 | 0 | 0 | 0 | 0 | 0 | 0 | 0 | 0 | 0 | 0 | 0 | 0 | 0 | 0 | 0 | 0 | 0 | 0 | 0 |
| Discards | 0 | 0 | 0 | 0 | 0 | 0 | 0 | 0 | 0 | 0 | 0 | 0 | 0 | 0 | 0 | 0 | 0 | 0 | 0 | 0 | 0 | 0 | 0 | 0 | 0 | 0 | 0 | 0 | 0.011 | 0 |
| Offal | 0 | 0 | 0 | 0 | 0 | 0 | 0 | 0 | 0 | 0 | 0 | 0 | 0 | 0 | 0 | 0 | 0 | 0 | 0 | 0 | 0 | 0 | 0 | 0 | 0 | 0 | 0 | 0 | 0 | 0 |
| Pelagic Detritus | 0 | 0 | 0 | 0 | 0 | 0 | 0 | 0 | 0 | 0 | 0 | 0 | 0 | 0 | 0 | 0 | 0 | 0 | 0 | 0 | 0 | 0 | 0 | 0 | 0 | 0 | 0 | 0 | 0 | 0 |
| Benthic Detritus | 0 | 0 | 0 | 0 | 0 | 0 | 0 | 0 | 0 | 0 | 0 | 0 | 0 | 0 | 0 | 0 | 0 | 0 | 0 | 0 | 0 | 0 | 0 | 0 | 0 | 0 | 0 | 0 | 0 | 0 |
| Outside Detritus | 0 | 0 | 0 | 0 | 0 | 0 | 0 | 0 | 0 | 0 | 0 | 0 | 0 | 0 | 0 | 0 | 0 | 0 | 0 | 0 | 0 | 0 | 0 | 0 | 0 | 0 | 0 | 0 | 0 | 0 |

**Table S3:** Diet matrix for the East Bering Sea - continued

| Group | Sleeper shark | W. Pollock Juv | W. Pollock Ad | P. Cod Juv | P. Cod Ad | Herring Juv | Herring Ad | Arrowtooth Juv | Arrowtooth Ad | Kamchatka fl. Juv | Kamchatka fl. Ad | Gr. Turbot Juv | Gr. Turbot Ad | P. Halibut Juv | P. Halibut Ad | YF. Sole Juv | YF. Sole Ad | FH. Sole Juv | FH. Sole Ad | N. Rock sole Juv | N. Rock sole Ad | AK Plaice | Dover Sole | Rex Sole | Misc. Flatfish | Alaska skate | Other skates | Sablefish Juv | Sablefish Ad | Eelpouts |
|---|---|---|---|---|---|---|---|---|---|---|---|---|---|---|---|---|---|---|---|---|---|---|---|---|---|---|---|---|---|---|
| Transient KIllers | 0 | 0 | 0 | 0 | 0 | 0 | 0 | 0 | 0 | 0 | 0 | 0 | 0 | 0 | 0 | 0 | 0 | 0 | 0 | 0 | 0 | 0 | 0 | 0 | 0 | 0 | 0 | 0 | 0 | 0 |
| Sperm and Beaked Whales | 0 | 0 | 0 | 0 | 0 | 0 | 0 | 0 | 0 | 0 | 0 | 0 | 0 | 0 | 0 | 0 | 0 | 0 | 0 | 0 | 0 | 0 | 0 | 0 | 0 | 0 | 0 | 0 | 0 | 0 |
| Resident Killers | 0 | 0 | 0 | 0 | 0 | 0 | 0 | 0 | 0 | 0 | 0 | 0 | 0 | 0 | 0 | 0 | 0 | 0 | 0 | 0 | 0 | 0 | 0 | 0 | 0 | 0 | 0 | 0 | 0 | 0 |
| Porpoises | 0 | 0 | 0 | 0 | 0 | 0 | 0 | 0 | 0 | 0 | 0 | 0 | 0 | 0 | 0 | 0 | 0 | 0 | 0 | 0 | 0 | 0 | 0 | 0 | 0 | 0 | 0 | 0 | 0 | 0 |
| Belugas | 0 | 0 | 0 | 0 | 0 | 0 | 0 | 0 | 0 | 0 | 0 | 0 | 0 | 0 | 0 | 0 | 0 | 0 | 0 | 0 | 0 | 0 | 0 | 0 | 0 | 0 | 0 | 0 | 0 | 0 |
| Gray Whales | 0 | 0 | 0 | 0 | 0 | 0 | 0 | 0 | 0 | 0 | 0 | 0 | 0 | 0 | 0 | 0 | 0 | 0 | 0 | 0 | 0 | 0 | 0 | 0 | 0 | 0 | 0 | 0 | 0 | 0 |
| Humpbacks | 0 | 0 | 0 | 0 | 0 | 0 | 0 | 0 | 0 | 0 | 0 | 0 | 0 | 0 | 0 | 0 | 0 | 0 | 0 | 0 | 0 | 0 | 0 | 0 | 0 | 0 | 0 | 0 | 0 | 0 |
| Fin Whales | 0 | 0 | 0 | 0 | 0 | 0 | 0 | 0 | 0 | 0 | 0 | 0 | 0 | 0 | 0 | 0 | 0 | 0 | 0 | 0 | 0 | 0 | 0 | 0 | 0 | 0 | 0 | 0 | 0 | 0 |
| Sei Whales | 0 | 0 | 0 | 0 | 0 | 0 | 0 | 0 | 0 | 0 | 0 | 0 | 0 | 0 | 0 | 0 | 0 | 0 | 0 | 0 | 0 | 0 | 0 | 0 | 0 | 0 | 0 | 0 | 0 | 0 |
| Right Whales | 0 | 0 | 0 | 0 | 0 | 0 | 0 | 0 | 0 | 0 | 0 | 0 | 0 | 0 | 0 | 0 | 0 | 0 | 0 | 0 | 0 | 0 | 0 | 0 | 0 | 0 | 0 | 0 | 0 | 0 |
| Minke Whales | 0 | 0 | 0 | 0 | 0 | 0 | 0 | 0 | 0 | 0 | 0 | 0 | 0 | 0 | 0 | 0 | 0 | 0 | 0 | 0 | 0 | 0 | 0 | 0 | 0 | 0 | 0 | 0 | 0 | 0 |
| Bowhead Whales | 0 | 0 | 0 | 0 | 0 | 0 | 0 | 0 | 0 | 0 | 0 | 0 | 0 | 0 | 0 | 0 | 0 | 0 | 0 | 0 | 0 | 0 | 0 | 0 | 0 | 0 | 0 | 0 | 0 | 0 |
| Sea Otters | 0 | 0 | 0 | 0 | 0 | 0 | 0 | 0 | 0 | 0 | 0 | 0 | 0 | 0 | 0 | 0 | 0 | 0 | 0 | 0 | 0 | 0 | 0 | 0 | 0 | 0 | 0 | 0 | 0 | 0 |
| Walrus Bd Seals | 0 | 0 | 0 | 0 | 0 | 0 | 0 | 0 | 0 | 0 | 0 | 0 | 0 | 0 | 0 | 0 | 0 | 0 | 0 | 0 | 0 | 0 | 0 | 0 | 0 | 0 | 0 | 0 | 0 | 0 |
| N Fur Seal Juv | 0 | 0 | 0 | 0 | 0 | 0 | 0 | 0 | 0 | 0 | 0 | 0 | 0 | 0 | 0 | 0 | 0 | 0 | 0 | 0 | 0 | 0 | 0 | 0 | 0 | 0 | 0 | 0 | 0 | 0 |
| N Fur Seal Ad | 0 | 0 | 0 | 0 | 0 | 0 | 0 | 0 | 0 | 0 | 0 | 0 | 0 | 0 | 0 | 0 | 0 | 0 | 0 | 0 | 0 | 0 | 0 | 0 | 0 | 0 | 0 | 0 | 0 | 0 |
| Steller Sea Lions Juv | 0 | 0 | 0 | 0 | 0 | 0 | 0 | 0 | 0 | 0 | 0 | 0 | 0 | 0 | 0 | 0 | 0 | 0 | 0 | 0 | 0 | 0 | 0 | 0 | 0 | 0 | 0 | 0 | 0 | 0 |
| Steller Sea Lions Ad | 0 | 0 | 0 | 0 | 0 | 0 | 0 | 0 | 0 | 0 | 0 | 0 | 0 | 0 | 0 | 0 | 0 | 0 | 0 | 0 | 0 | 0 | 0 | 0 | 0 | 0 | 0 | 0 | 0 | 0 |
| Resident Seals | 0 | 0 | 0 | 0 | 0 | 0 | 0 | 0 | 0 | 0 | 0 | 0 | 0 | 0 | 0 | 0 | 0 | 0 | 0 | 0 | 0 | 0 | 0 | 0 | 0 | 0 | 0 | 0 | 0 | 0 |
| Wintering Seals | 0 | 0 | 0 | 0 | 0 | 0 | 0 | 0 | 0 | 0 | 0 | 0 | 0 | 0 | 0 | 0 | 0 | 0 | 0 | 0 | 0 | 0 | 0 | 0 | 0 | 0 | 0 | 0 | 0 | 0 |
| Shearwater | 0 | 0 | 0 | 0 | 0 | 0 | 0 | 0 | 0 | 0 | 0 | 0 | 0 | 0 | 0 | 0 | 0 | 0 | 0 | 0 | 0 | 0 | 0 | 0 | 0 | 0 | 0 | 0 | 0 | 0 |
| Murres | 0 | 0 | 0 | 0 | 0 | 0 | 0 | 0 | 0 | 0 | 0 | 0 | 0 | 0 | 0 | 0 | 0 | 0 | 0 | 0 | 0 | 0 | 0 | 0 | 0 | 0 | 0 | 0 | 0 | 0 |
| Kittiwakes | 0 | 0 | 0 | 0 | 0 | 0 | 0 | 0 | 0 | 0 | 0 | 0 | 0 | 0 | 0 | 0 | 0 | 0 | 0 | 0 | 0 | 0 | 0 | 0 | 0 | 0 | 0 | 0 | 0 | 0 |
| Auklets | 0 | 0 | 0 | 0 | 0 | 0 | 0 | 0 | 0 | 0 | 0 | 0 | 0 | 0 | 0 | 0 | 0 | 0 | 0 | 0 | 0 | 0 | 0 | 0 | 0 | 0 | 0 | 0 | 0 | 0 |
| Puffins | 0 | 0 | 0 | 0 | 0 | 0 | 0 | 0 | 0 | 0 | 0 | 0 | 0 | 0 | 0 | 0 | 0 | 0 | 0 | 0 | 0 | 0 | 0 | 0 | 0 | 0 | 0 | 0 | 0 | 0 |
| Fulmars | 0 | 0 | 0 | 0 | 0 | 0 | 0 | 0 | 0 | 0 | 0 | 0 | 0 | 0 | 0 | 0 | 0 | 0 | 0 | 0 | 0 | 0 | 0 | 0 | 0 | 0 | 0 | 0 | 0 | 0 |
| Storm Petrels | 0 | 0 | 0 | 0 | 0 | 0 | 0 | 0 | 0 | 0 | 0 | 0 | 0 | 0 | 0 | 0 | 0 | 0 | 0 | 0 | 0 | 0 | 0 | 0 | 0 | 0 | 0 | 0 | 0 | 0 |
| Cormorants | 0 | 0 | 0 | 0 | 0 | 0 | 0 | 0 | 0 | 0 | 0 | 0 | 0 | 0 | 0 | 0 | 0 | 0 | 0 | 0 | 0 | 0 | 0 | 0 | 0 | 0 | 0 | 0 | 0 | 0 |
| Gulls | 0 | 0 | 0 | 0 | 0 | 0 | 0 | 0 | 0 | 0 | 0 | 0 | 0 | 0 | 0 | 0 | 0 | 0 | 0 | 0 | 0 | 0 | 0 | 0 | 0 | 0 | 0 | 0 | 0 | 0 |
| Albatross Jaeger | 0 | 0 | 0 | 0 | 0 | 0 | 0 | 0 | 0 | 0 | 0 | 0 | 0 | 0 | 0 | 0 | 0 | 0 | 0 | 0 | 0 | 0 | 0 | 0 | 0 | 0 | 0 | 0 | 0 | 0 |
| Sleeper shark | 0 | 0 | 0 | 0 | 0 | 0 | 0 | 0 | 0 | 0 | 0 | 0 | 0 | 0 | 0 | 0 | 0 | 0 | 0 | 0 | 0 | 0 | 0 | 0 | 0 | 0 | 0 | 0 | 0 | 0 |
| W. Pollock Juv | 0 | 0.01 | 0.044 | 0.007 | 0.029 | 0 | 0 | 0.219 | 0.469 | 0.423 | 0.581 | 0.012 | 0.271 | 0 | 0.019 | 0 | 0.01 | 0 | 0.168 | 0 | 0.001 | 0 | 0 | 0 | 0.032 | 0.002 | 0 | 0 | 0 | 0 |
| W. Pollock Ad | 0.2 | 0 | 0.085 | 0 | 0.259 | 0 | 0 | 0 | 0.195 | 0 | 0.213 | 0 | 0.105 | 0 | 0.521 | 0 | 0 | 0 | 0 | 0 | 0 | 0 | 0 | 0 | 0 | 0.405 | 0.449 | 0 | 0.459 | 0.004 |
| P. Cod Juv | 0 | 0 | 0 | 0 | 0 | 0 | 0 | 0 | 0 | 0 | 0 | 0 | 0 | 0 | 0 | 0 | 0 | 0 | 0 | 0 | 0 | 0 | 0 | 0 | 0 | 0 | 0.004 | 0 | 0 | 0 |
| P. Cod Ad | 0 | 0 | 0.001 | 0 | 0.005 | 0 | 0 | 0 | 0 | 0 | 0 | 0 | 0.001 | 0 | 0.047 | 0 | 0 | 0 | 0 | 0 | 0 | 0 | 0 | 0 | 0 | 0.006 | 0 | 0 | 0 | 0 |
| Herring Juv | 0 | 0 | 0 | 0 | 0.002 | 0 | 0 | 0 | 0 | 0 | 0 | 0 | 0 | 0 | 0 | 0 | 0 | 0 | 0 | 0 | 0.002 | 0 | 0 | 0 | 0 | 0 | 0 | 0 | 0 | 0 |

| Group | Sleeper shark | W. Pollock Juv | W. Pollock Ad | P. Cod Juv | P. Cod Ad | Herring Juv | Herring Ad | Arrowtooth Juv | Arrowtooth Ad | Kamchatka fl. Juv | Kamchatka fl. Ad | Gr. Turbot Juv | Gr. Turbot Ad | P. Halibut Juv | P. Halibut Ad | YF. Sole Juv | YF. Sole Ad | FH. Sole Juv | FH. Sole Ad | N. Rock sole Juv | N. Rock sole Ad | AK Plaice | Dover Sole | Rex Sole | Misc. Flatfish | Alaska skate | Other skates | Sablefish Juv | Sablefish Ad | Eelpouts |
|---|---|---|---|---|---|---|---|---|---|---|---|---|---|---|---|---|---|---|---|---|---|---|---|---|---|---|---|---|---|---|
| Herring Ad | 0 | 0 | 0.001 | 0 | 0 | 0 | 0 | 0 | 0.013 | 0 | 0 | 0 | 0.012 | 0 | 0.003 | 0 | 0 | 0 | 0 | 0 | 0 | 0 | 0 | 0 | 0 | 0.002 | 0 | 0 | 0 | 0.001 |
| Arrowtooth Juv | 0 | 0 | 0 | 0 | 0 | 0 | 0 | 0 | 0.003 | 0 | 0 | 0 | 0 | 0 | 0 | 0 | 0 | 0 | 0 | 0 | 0 | 0 | 0 | 0 | 0 | 0 | 0 | 0 | 0 | 0 |
| Arrowtooth Ad | 0.12 | 0 | 0.001 | 0 | 0.001 | 0 | 0 | 0 | 0 | 0 | 0 | 0 | 0 | 0 | 0.001 | 0 | 0 | 0 | 0 | 0 | 0 | 0 | 0 | 0 | 0 | 0.026 | 0 | 0 | 0 | 0 |
| Kamchatka fl. Juv | 0 | 0 | 0 | 0 | 0 | 0 | 0 | 0 | 0 | 0 | 0 | 0 | 0 | 0 | 0 | 0 | 0 | 0 | 0 | 0 | 0 | 0 | 0 | 0 | 0 | 0 | 0 | 0 | 0 | 0 |
| Kamchatka fl. Ad | 0.007 | 0 | 0 | 0 | 0 | 0 | 0 | 0 | 0 | 0 | 0 | 0 | 0 | 0 | 0 | 0 | 0 | 0 | 0 | 0 | 0 | 0 | 0 | 0 | 0 | 0 | 0 | 0 | 0 | 0 |
| Gr. Turbot Juv | 0 | 0 | 0 | 0 | 0 | 0 | 0 | 0 | 0 | 0 | 0 | 0 | 0 | 0 | 0 | 0 | 0 | 0 | 0 | 0 | 0 | 0 | 0 | 0 | 0 | 0 | 0 | 0 | 0 | 0 |
| Gr. Turbot Ad | 0.044 | 0 | 0 | 0 | 0 | 0 | 0 | 0 | 0 | 0 | 0 | 0 | 0 | 0 | 0 | 0 | 0 | 0 | 0 | 0 | 0 | 0 | 0 | 0 | 0 | 0 | 0 | 0 | 0 | 0 |
| P. Halibut Juv | 0 | 0 | 0 | 0 | 0 | 0 | 0 | 0.001 | 0 | 0 | 0 | 0 | 0 | 0 | 0 | 0 | 0 | 0 | 0 | 0 | 0 | 0 | 0 | 0 | 0 | 0.001 | 0 | 0 | 0 | 0 |
| P. Halibut Ad | 0.028 | 0 | 0 | 0 | 0.001 | 0 | 0 | 0 | 0 | 0 | 0 | 0 | 0 | 0 | 0 | 0 | 0 | 0 | 0 | 0 | 0 | 0 | 0 | 0 | 0 | 0 | 0 | 0 | 0 | 0 |
| YF. Sole Juv | 0 | 0 | 0 | 0 | 0 | 0 | 0 | 0 | 0 | 0 | 0 | 0 | 0 | 0 | 0 | 0 | 0 | 0 | 0 | 0 | 0 | 0 | 0 | 0 | 0 | 0 | 0 | 0 | 0 | 0 |
| YF. Sole Ad | 0 | 0 | 0 | 0 | 0.006 | 0 | 0 | 0 | 0 | 0 | 0 | 0 | 0 | 0 | 0.033 | 0 | 0 | 0 | 0 | 0 | 0 | 0 | 0 | 0 | 0 | 0.007 | 0.042 | 0 | 0 | 0 |
| FH. Sole Juv | 0 | 0 | 0 | 0 | 0.001 | 0 | 0 | 0.001 | 0.011 | 0 | 0 | 0 | 0 | 0 | 0.001 | 0 | 0 | 0 | 0.002 | 0 | 0 | 0 | 0 | 0 | 0 | 0 | 0 | 0 | 0 | 0 |
| FH. Sole Ad | 0 | 0 | 0 | 0 | 0.003 | 0 | 0 | 0 | 0 | 0 | 0.001 | 0 | 0 | 0 | 0.011 | 0 | 0 | 0 | 0 | 0 | 0 | 0 | 0 | 0 | 0 | 0 | 0 | 0 | 0 | 0 |
| N. Rock sole Juv | 0 | 0 | 0 | 0 | 0 | 0 | 0 | 0 | 0 | 0 | 0 | 0 | 0 | 0 | 0 | 0 | 0 | 0 | 0 | 0 | 0 | 0 | 0 | 0 | 0 | 0 | 0 | 0 | 0 | 0 |
| N. Rock sole Ad | 0 | 0 | 0 | 0 | 0.013 | 0 | 0 | 0 | 0 | 0 | 0 | 0 | 0 | 0 | 0.032 | 0 | 0 | 0 | 0 | 0 | 0 | 0 | 0 | 0 | 0 | 0.085 | 0.02 | 0 | 0 | 0 |
| AK Plaice | 0 | 0 | 0 | 0 | 0 | 0 | 0 | 0 | 0 | 0 | 0 | 0 | 0 | 0 | 0.005 | 0 | 0 | 0 | 0 | 0 | 0 | 0 | 0 | 0 | 0 | 0 | 0 | 0 | 0 | 0 |
| Dover Sole | 0 | 0 | 0 | 0 | 0 | 0 | 0 | 0 | 0 | 0 | 0 | 0 | 0 | 0 | 0 | 0 | 0 | 0 | 0 | 0 | 0 | 0 | 0 | 0 | 0 | 0 | 0 | 0 | 0 | 0 |
| Rex Sole | 0 | 0 | 0 | 0 | 0 | 0 | 0 | 0 | 0 | 0 | 0 | 0 | 0 | 0 | 0.001 | 0 | 0 | 0 | 0 | 0 | 0 | 0 | 0 | 0 | 0 | 0 | 0 | 0 | 0 | 0 |
| Misc. Flatfish | 0 | 0 | 0 | 0 | 0.001 | 0 | 0 | 0 | 0 | 0 | 0 | 0 | 0 | 0 | 0.027 | 0 | 0 | 0 | 0 | 0 | 0 | 0 | 0 | 0 | 0 | 0.002 | 0 | 0 | 0 | 0 |
| Alaska skate | 0 | 0 | 0 | 0 | 0 | 0 | 0 | 0 | 0 | 0 | 0 | 0 | 0 | 0 | 0 | 0 | 0 | 0 | 0 | 0 | 0 | 0 | 0 | 0 | 0 | 0 | 0 | 0 | 0 | 0 |
| Other skates | 0 | 0 | 0 | 0 | 0 | 0 | 0 | 0 | 0 | 0 | 0 | 0 | 0 | 0 | 0 | 0 | 0 | 0 | 0 | 0 | 0 | 0 | 0 | 0 | 0 | 0 | 0 | 0 | 0 | 0 |
| Sablefish Juv | 0 | 0 | 0 | 0 | 0 | 0 | 0 | 0 | 0 | 0 | 0 | 0 | 0 | 0 | 0 | 0 | 0 | 0 | 0 | 0 | 0 | 0 | 0 | 0 | 0 | 0 | 0 | 0 | 0 | 0 |
| Sablefish Ad | 0 | 0 | 0 | 0 | 0 | 0 | 0 | 0 | 0 | 0 | 0 | 0 | 0 | 0 | 0.001 | 0 | 0 | 0 | 0 | 0 | 0 | 0 | 0 | 0 | 0 | 0 | 0 | 0 | 0 | 0 |
| Eelpouts | 0 | 0 | 0.001 | 0 | 0.04 | 0 | 0 | 0 | 0.043 | 0 | 0.053 | 0 | 0.073 | 0 | 0.009 | 0 | 0 | 0.003 | 0.002 | 0 | 0.001 | 0 | 0 | 0 | 0.042 | 0.087 | 0.032 | 0 | 0 | 0.018 |
| Grenadiers | 0.2 | 0 | 0 | 0 | 0 | 0 | 0 | 0 | 0 | 0 | 0 | 0 | 0 | 0 | 0 | 0 | 0 | 0 | 0 | 0 | 0 | 0 | 0 | 0 | 0 | 0 | 0 | 0 | 0 | 0 |
| Misc. fish deep | 0 | 0 | 0 | 0 | 0 | 0 | 0 | 0 | 0 | 0 | 0 | 0 | 0 | 0 | 0 | 0 | 0 | 0 | 0 | 0 | 0 | 0 | 0 | 0 | 0 | 0 | 0 | 0 | 0 | 0 |
| POP | 0 | 0 | 0 | 0 | 0 | 0 | 0 | 0 | 0 | 0 | 0 | 0 | 0 | 0 | 0 | 0 | 0 | 0 | 0 | 0 | 0 | 0 | 0 | 0 | 0 | 0 | 0 | 0 | 0 | 0 |
| Sharpchin Rock | 0 | 0 | 0 | 0 | 0 | 0 | 0 | 0 | 0 | 0 | 0 | 0 | 0 | 0 | 0 | 0 | 0 | 0 | 0 | 0 | 0 | 0 | 0 | 0 | 0 | 0 | 0 | 0 | 0 | 0 |
| Northern Rock | 0 | 0 | 0 | 0 | 0 | 0 | 0 | 0 | 0 | 0 | 0 | 0 | 0 | 0 | 0 | 0 | 0 | 0 | 0 | 0 | 0 | 0 | 0 | 0 | 0 | 0 | 0 | 0 | 0 | 0 |
| Dusky Rock | 0 | 0 | 0 | 0 | 0 | 0 | 0 | 0 | 0 | 0 | 0 | 0 | 0 | 0 | 0 | 0 | 0 | 0 | 0 | 0 | 0 | 0 | 0 | 0 | 0 | 0 | 0 | 0 | 0 | 0 |
| Shortraker Rock | 0 | 0 | 0 | 0 | 0 | 0 | 0 | 0 | 0 | 0 | 0 | 0 | 0 | 0 | 0 | 0 | 0 | 0 | 0 | 0 | 0 | 0 | 0 | 0 | 0 | 0 | 0 | 0 | 0 | 0 |
| Rougheye Rock | 0 | 0 | 0 | 0 | 0 | 0 | 0 | 0 | 0 | 0 | 0 | 0 | 0 | 0 | 0 | 0 | 0 | 0 | 0 | 0 | 0 | 0 | 0 | 0 | 0 | 0 | 0 | 0 | 0 | 0 |
| Shortspine Thorns | 0 | 0 | 0 | 0 | 0 | 0 | 0 | 0 | 0 | 0 | 0 | 0 | 0 | 0 | 0 | 0 | 0 | 0 | 0 | 0 | 0 | 0 | 0 | 0 | 0 | 0 | 0 | 0 | 0 | 0 |
| Other Sebastes | 0 | 0 | 0 | 0 | 0 | 0 | 0 | 0 | 0 | 0 | 0 | 0 | 0 | 0 | 0 | 0 | 0 | 0 | 0 | 0 | 0 | 0 | 0 | 0 | 0 | 0 | 0 | 0 | 0 | 0 |
| Atka mackerel Juv | 0 | 0 | 0 | 0 | 0 | 0 | 0 | 0 | 0 | 0 | 0 | 0 | 0 | 0 | 0 | 0 | 0 | 0 | 0 | 0 | 0 | 0 | 0 | 0 | 0 | 0 | 0 | 0 | 0 | 0 |
| Atka mackerel Ad | 0 | 0 | 0 | 0 | 0.002 | 0 | 0 | 0 | 0 | 0 | 0 | 0 | 0 | 0 | 0.005 | 0 | 0 | 0 | 0 | 0 | 0 | 0 | 0 | 0 | 0 | 0 | 0 | 0 | 0 | 0 |
| Greenlings | 0 | 0 | 0 | 0 | 0 | 0 | 0 | 0 | 0 | 0 | 0 | 0 | 0 | 0 | 0 | 0 | 0 | 0 | 0 | 0 | 0 | 0 | 0 | 0 | 0 | 0 | 0 | 0 | 0 | 0 |
| Lg. Sculpins | 0 | 0 | 0 | 0 | 0.002 | 0 | 0 | 0 | 0 | 0 | 0 | 0 | 0 | 0 | 0.002 | 0 | 0 | 0 | 0 | 0 | 0 | 0 | 0 | 0 | 0 | 0 | 0 | 0 | 0 | 0 |

| Group | Sleeper shark | W. Pollock Juv | W. Pollock Ad | P. Cod Juv | P. Cod Ad | Herring Juv | Herring Ad | Arrowtooth Juv | Arrowtooth Ad | Kamchatka fl. Juv | Kamchatka fl. Ad | Gr. Turbot Juv | Gr. Turbot Ad | P. Halibut Juv | P. Halibut Ad | YF. Sole Juv | YF. Sole Ad | FH. Sole Juv | FH. Sole Ad | N. Rock sole Juv | N. Rock sole Ad | AK Plaice | Dover Sole | Rex Sole | Misc. Flatfish | Alaska skate | Other skates | Sablefish Juv | Sablefish Ad | Eelpouts |
|---|---|---|---|---|---|---|---|---|---|---|---|---|---|---|---|---|---|---|---|---|---|---|---|---|---|---|---|---|---|---|
| Other sculpins | 0 | 0 | 0.001 | 0.01 | 0.011 | 0 | 0 | 0.026 | 0.009 | 0 | 0.003 | 0 | 0.022 | 0 | 0.022 | 0 | 0.002 | 0.008 | 0.001 | 0.001 | 0 | 0 | 0 | 0 | 0 | 0.024 | 0.002 | 0 | 0 | 0.014 |
| Misc. fish shallow | 0.02 | 0 | 0 | 0 | 0.004 | 0 | 0 | 0.002 | 0 | 0 | 0.008 | 0 | 0.01 | 0 | 0.015 | 0 | 0 | 0 | 0.005 | 0 | 0 | 0 | 0 | 0 | 0 | 0.004 | 0.033 | 0 | 0 | 0.001 |
| Octopi | 0.02 | 0 | 0 | 0 | 0.006 | 0 | 0 | 0 | 0 | 0 | 0 | 0 | 0 | 0 | 0.005 | 0 | 0 | 0 | 0 | 0 | 0 | 0 | 0 | 0 | 0 | 0 | 0 | 0 | 0.039 | 0 |
| Squids | 0.2 | 0 | 0.002 | 0 | 0.001 | 0 | 0 | 0 | 0.009 | 0 | 0.002 | 0 | 0.325 | 0 | 0.003 | 0 | 0 | 0.003 | 0 | 0 | 0 | 0 | 0 | 0 | 0 | 0 | 0 | 0.1 | 0.111 | 0 |
| Salmon returning | 0.05 | 0 | 0 | 0 | 0.002 | 0 | 0 | 0 | 0 | 0 | 0 | 0 | 0 | 0 | 0 | 0 | 0 | 0 | 0 | 0 | 0 | 0 | 0 | 0 | 0 | 0.026 | 0 | 0 | 0 | 0 |
| Salmon outgoing | 0 | 0 | 0 | 0 | 0 | 0 | 0 | 0 | 0 | 0 | 0 | 0 | 0 | 0 | 0 | 0 | 0 | 0 | 0 | 0 | 0 | 0 | 0 | 0 | 0 | 0 | 0 | 0 | 0 | 0 |
| Bathylagidae | 0 | 0 | 0.001 | 0 | 0 | 0 | 0 | 0 | 0 | 0 | 0 | 0 | 0.051 | 0 | 0 | 0 | 0 | 0 | 0 | 0 | 0 | 0 | 0 | 0 | 0 | 0 | 0 | 0 | 0.018 | 0 |
| Myctophidae | 0 | 0.001 | 0.003 | 0 | 0 | 0 | 0 | 0 | 0.003 | 0 | 0.001 | 0 | 0.064 | 0 | 0 | 0 | 0 | 0 | 0 | 0 | 0 | 0 | 0 | 0 | 0 | 0 | 0 | 0 | 0.001 | 0 |
| Capelin | 0 | 0 | 0.001 | 0 | 0.004 | 0 | 0 | 0 | 0.012 | 0 | 0 | 0 | 0 | 0 | 0.039 | 0 | 0 | 0 | 0.001 | 0 | 0 | 0 | 0 | 0 | 0.046 | 0 | 0 | 0 | 0 | 0 |
| Sandlance | 0 | 0.001 | 0.002 | 0.006 | 0.024 | 0 | 0 | 0 | 0 | 0 | 0 | 0 | 0 | 0 | 0.034 | 0.002 | 0.006 | 0 | 0.003 | 0 | 0.141 | 0.007 | 0 | 0 | 0 | 0.029 | 0 | 0 | 0 | 0 |
| Eulachon | 0 | 0 | 0 | 0 | 0 | 0 | 0 | 0 | 0.021 | 0 | 0 | 0 | 0 | 0 | 0 | 0 | 0 | 0 | 0 | 0 | 0 | 0 | 0 | 0 | 0 | 0 | 0 | 0 | 0 | 0 |
| Oth. managed forage | 0 | 0 | 0.001 | 0.003 | 0.008 | 0 | 0 | 0.004 | 0.006 | 0.008 | 0.03 | 0 | 0.01 | 0 | 0.003 | 0 | 0.001 | 0 | 0.008 | 0 | 0.001 | 0 | 0 | 0 | 0.008 | 0.003 | 0 | 0 | 0.308 | 0.012 |
| Oth. pelagic smelt | 0 | 0 | 0 | 0 | 0 | 0 | 0 | 0 | 0 | 0 | 0 | 0 | 0 | 0 | 0 | 0 | 0 | 0 | 0 | 0 | 0 | 0 | 0 | 0 | 0 | 0 | 0 | 0 | 0 | 0 |
| Bairdi | 0 | 0 | 0 | 0 | 0.048 | 0 | 0 | 0 | 0 | 0 | 0.001 | 0 | 0 | 0 | 0.026 | 0 | 0.003 | 0 | 0.007 | 0 | 0 | 0 | 0.015 | 0 | 0.006 | 0.005 | 0.056 | 0 | 0 | 0.001 |
| King Crab | 0 | 0 | 0 | 0 | 0.01 | 0 | 0 | 0 | 0 | 0 | 0 | 0 | 0 | 0 | 0.002 | 0 | 0 | 0 | 0 | 0 | 0 | 0 | 0 | 0 | 0 | 0 | 0 | 0 | 0 | 0 |
| Opilio | 0 | 0 | 0 | 0.001 | 0.074 | 0 | 0 | 0 | 0 | 0 | 0 | 0 | 0 | 0 | 0.034 | 0 | 0.007 | 0 | 0.01 | 0 | 0.001 | 0.006 | 0 | 0 | 0 | 0.031 | 0.022 | 0 | 0 | 0.093 |
| Pandalidae | 0.003 | 0.001 | 0.036 | 0.01 | 0.053 | 0 | 0 | 0.054 | 0.039 | 0.028 | 0.023 | 0 | 0.004 | 0 | 0.01 | 0 | 0 | 0.061 | 0.108 | 0.002 | 0 | 0 | 0.222 | 0 | 0.096 | 0.001 | 0 | 0 | 0.001 | 0 |
| NP shrimp | 0.003 | 0.005 | 0.015 | 0.177 | 0.076 | 0 | 0 | 0.419 | 0.022 | 0.403 | 0.039 | 0.004 | 0.005 | 0.693 | 0.005 | 0.061 | 0.073 | 0.111 | 0.102 | 0.01 | 0.031 | 0 | 0.265 | 0.545 | 0.1 | 0.065 | 0.111 | 0 | 0.005 | 0.014 |
| Sea stars | 0 | 0 | 0 | 0 | 0 | 0 | 0 | 0 | 0 | 0 | 0 | 0 | 0 | 0 | 0 | 0 | 0 | 0 | 0 | 0 | 0 | 0 | 0 | 0 | 0 | 0.001 | 0 | 0 | 0.001 | 0 |
| Brittle stars | 0 | 0 | 0 | 0 | 0 | 0 | 0 | 0 | 0 | 0 | 0 | 0 | 0 | 0 | 0 | 0.01 | 0.032 | 0.142 | 0.248 | 0.001 | 0.013 | 0.002 | 0.01 | 0 | 0.06 | 0 | 0 | 0 | 0 | 0.182 |
| Urchins dollars cucumbers | 0 | 0 | 0 | 0 | 0 | 0 | 0 | 0 | 0 | 0 | 0 | 0 | 0 | 0 | 0 | 0.022 | 0.046 | 0 | 0 | 0.006 | 0.023 | 0.015 | 0 | 0 | 0 | 0 | 0 | 0 | 0 | 0 |
| Snails | 0.003 | 0 | 0 | 0 | 0.006 | 0 | 0 | 0 | 0 | 0 | 0 | 0 | 0 | 0 | 0.002 | 0.007 | 0.01 | 0 | 0.02 | 0 | 0.001 | 0.004 | 0 | 0 | 0 | 0 | 0 | 0 | 0 | 0 |
| Hermit crabs | 0.003 | 0 | 0.001 | 0.007 | 0.07 | 0 | 0 | 0.005 | 0 | 0 | 0 | 0 | 0.003 | 0 | 0.047 | 0 | 0.026 | 0 | 0.016 | 0 | 0.006 | 0 | 0 | 0 | 0.001 | 0.022 | 0.201 | 0 | 0 | 0.003 |
| Misc. crabs | 0 | 0 | 0 | 0 | 0.014 | 0 | 0 | 0 | 0 | 0 | 0 | 0 | 0 | 0 | 0.007 | 0.001 | 0.002 | 0 | 0.001 | 0 | 0.001 | 0 | 0 | 0 | 0 | 0.001 | 0.017 | 0 | 0 | 0.005 |
| Misc. Crustacean | 0 | 0.016 | 0.001 | 0.007 | 0.001 | 0 | 0 | 0 | 0 | 0 | 0 | 0 | 0 | 0 | 0 | 0.435 | 0.009 | 0.012 | 0.001 | 0.004 | 0.012 | 0.002 | 0.002 | 0 | 0.005 | 0 | 0 | 0 | 0 | 0.005 |
| Benthic Amphipods | 0 | 0.057 | 0.02 | 0.392 | 0.028 | 0 | 0 | 0.002 | 0.001 | 0.026 | 0.007 | 0.002 | 0.003 | 0 | 0 | 0.208 | 0.075 | 0.274 | 0.023 | 0.349 | 0.096 | 0.1 | 0.089 | 0.118 | 0.064 | 0.009 | 0.005 | 0 | 0 | 0.328 |
| Anemones | 0 | 0 | 0 | 0 | 0 | 0 | 0 | 0 | 0 | 0 | 0 | 0 | 0 | 0 | 0 | 0 | 0 | 0 | 0 | 0 | 0 | 0 | 0 | 0 | 0 | 0 | 0 | 0 | 0 | 0 |
| Corals | 0 | 0 | 0 | 0 | 0 | 0 | 0 | 0 | 0 | 0 | 0 | 0 | 0 | 0 | 0 | 0 | 0 | 0 | 0 | 0 | 0 | 0 | 0 | 0 | 0 | 0 | 0 | 0 | 0 | 0 |
| Hydroids | 0 | 0 | 0 | 0 | 0 | 0 | 0 | 0 | 0 | 0 | 0 | 0 | 0 | 0 | 0 | 0 | 0 | 0 | 0 | 0 | 0 | 0 | 0 | 0 | 0 | 0 | 0 | 0 | 0 | 0.002 |
| Urochordata | 0 | 0 | 0 | 0 | 0.001 | 0 | 0 | 0 | 0 | 0 | 0 | 0 | 0 | 0 | 0 | 0.005 | 0.005 | 0 | 0.001 | 0.001 | 0.001 | 0 | 0 | 0 | 0 | 0 | 0 | 0.15 | 0 | 0 |
| Sea Pens | 0 | 0 | 0 | 0 | 0 | 0 | 0 | 0 | 0 | 0 | 0 | 0 | 0 | 0 | 0 | 0 | 0 | 0 | 0 | 0 | 0 | 0 | 0 | 0 | 0 | 0 | 0 | 0 | 0 | 0 |
| Sponges | 0 | 0 | 0 | 0 | 0 | 0 | 0 | 0 | 0 | 0 | 0 | 0 | 0 | 0 | 0 | 0 | 0 | 0 | 0 | 0 | 0 | 0 | 0 | 0 | 0 | 0 | 0 | 0 | 0 | 0 |
| Bivalves | 0 | 0.001 | 0 | 0 | 0.002 | 0 | 0 | 0 | 0 | 0 | 0 | 0 | 0 | 0 | 0.004 | 0.053 | 0.115 | 0.002 | 0.024 | 0.061 | 0.073 | 0.155 | 0.011 | 0 | 0.07 | 0 | 0 | 0 | 0 | 0.053 |
| Polychaetes | 0 | 0.006 | 0.002 | 0.041 | 0.036 | 0 | 0 | 0 | 0.001 | 0.029 | 0.002 | 0 | 0 | 0 | 0 | 0.142 | 0.253 | 0.012 | 0.039 | 0.536 | 0.472 | 0.584 | 0.282 | 0.205 | 0.244 | 0.003 | 0.003 | 0 | 0 | 0.239 |
| Misc. worms | 0 | 0 | 0.002 | 0.002 | 0.034 | 0 | 0 | 0 | 0 | 0 | 0.001 | 0 | 0 | 0 | 0.002 | 0.008 | 0.156 | 0.01 | 0.005 | 0.006 | 0.11 | 0.123 | 0.054 | 0.033 | 0.02 | 0 | 0 | 0 | 0 | 0.024 |
| Scyphozoid Jellies | 0 | 0 | 0 | 0 | 0 | 0 | 0 | 0 | 0 | 0 | 0 | 0 | 0 | 0 | 0 | 0 | 0.005 | 0 | 0 | 0 | 0 | 0 | 0 | 0 | 0 | 0 | 0 | 0 | 0.016 | 0 |
| Fish Larvae | 0 | 0 | 0 | 0 | 0 | 0 | 0 | 0 | 0 | 0 | 0 | 0 | 0 | 0 | 0 | 0 | 0 | 0 | 0 | 0 | 0 | 0 | 0 | 0 | 0 | 0 | 0 | 0 | 0 | 0 |
| Chaetognaths | 0 | 0.048 | 0.01 | 0 | 0 | 0 | 0 | 0 | 0 | 0 | 0 | 0 | 0 | 0 | 0 | 0 | 0 | 0.003 | 0.001 | 0 | 0 | 0 | 0 | 0 | 0 | 0 | 0 | 0 | 0 | 0 |

| Group | Sleeper shark | W. Pollock Juv | W. Pollock Ad | P. Cod Juv | P. Cod Ad | Herring Juv | Herring Ad | Arrowtooth Juv | Arrowtooth Ad | Kamchatka fl. Juv | Kamchatka fl. Ad | Gr. Turbot Juv | Gr. Turbot Ad | P. Halibut Juv | P. Halibut Ad | YF. Sole Juv | YF. Sole Ad | FH. Sole Juv | FH. Sole Ad | N. Rock sole Juv | N. Rock sole Ad | AK Plaice | Dover Sole | Rex Sole | Misc. Flatfish | Alaska skate | Other skates | Sablefish Juv | Sablefish Ad | Eelpouts |
|---|---|---|---|---|---|---|---|---|---|---|---|---|---|---|---|---|---|---|---|---|---|---|---|---|---|---|---|---|---|---|
| Euphausiids | 0 | 0.329 | 0.35 | 0.033 | 0.024 | 0.879 | 0.957 | 0.246 | 0.092 | 0.078 | 0.03 | 0.974 | 0.04 | 0 | 0.003 | 0.026 | 0.138 | 0.152 | 0.117 | 0.005 | 0.012 | 0.001 | 0.035 | 0.099 | 0.001 | 0 | 0 | 0.5 | 0 | 0 |
| Mysids | 0 | 0.094 | 0.012 | 0.297 | 0.025 | 0 | 0 | 0.02 | 0.001 | 0.005 | 0.001 | 0 | 0 | 0.306 | 0.001 | 0.011 | 0.005 | 0.196 | 0.046 | 0.017 | 0.001 | 0 | 0.002 | 0 | 0.195 | 0 | 0 | 0 | 0 | 0 |
| Pelagic Amphipods | 0 | 0.005 | 0.016 | 0 | 0 | 0 | 0 | 0 | 0 | 0 | 0 | 0.008 | 0 | 0 | 0 | 0 | 0 | 0 | 0 | 0 | 0 | 0 | 0.012 | 0 | 0.005 | 0 | 0 | 0 | 0 | 0.001 |
| Gelatinous filter feeders | 0 | 0.006 | 0.014 | 0 | 0 | 0 | 0 | 0 | 0 | 0 | 0 | 0 | 0 | 0 | 0 | 0 | 0.002 | 0 | 0 | 0 | 0.001 | 0 | 0 | 0 | 0 | 0 | 0 | 0 | 0 | 0 |
| Pteropods | 0 | 0 | 0.001 | 0 | 0 | 0 | 0 | 0 | 0 | 0 | 0 | 0 | 0 | 0 | 0 | 0 | 0.004 | 0.006 | 0 | 0 | 0.001 | 0 | 0 | 0 | 0 | 0 | 0 | 0.1 | 0 | 0 |
| Copepods | 0 | 0.419 | 0.365 | 0.006 | 0 | 0.121 | 0.043 | 0 | 0 | 0 | 0 | 0 | 0 | 0 | 0 | 0.004 | 0 | 0.004 | 0 | 0 | 0 | 0 | 0 | 0 | 0.003 | 0 | 0 | 0.15 | 0 | 0 |
| Pelagic microbes | 0 | 0 | 0 | 0 | 0 | 0 | 0 | 0 | 0 | 0 | 0 | 0 | 0 | 0 | 0 | 0 | 0 | 0 | 0 | 0 | 0 | 0 | 0 | 0 | 0 | 0 | 0 | 0 | 0 | 0.001 |
| Benthic microbes | 0 | 0 | 0 | 0 | 0 | 0 | 0 | 0 | 0 | 0 | 0 | 0 | 0 | 0 | 0 | 0 | 0 | 0 | 0 | 0 | 0 | 0 | 0 | 0 | 0 | 0 | 0 | 0 | 0 | 0 |
| Macroalgae | 0 | 0 | 0 | 0 | 0 | 0 | 0 | 0 | 0 | 0 | 0 | 0 | 0 | 0 | 0 | 0 | 0 | 0 | 0 | 0 | 0 | 0 | 0 | 0 | 0 | 0 | 0 | 0 | 0 | 0 |
| Lg Phytoplankton | 0 | 0 | 0 | 0 | 0 | 0 | 0 | 0 | 0 | 0 | 0 | 0 | 0 | 0 | 0 | 0 | 0 | 0 | 0 | 0 | 0 | 0 | 0 | 0 | 0 | 0 | 0 | 0 | 0 | 0 |
| Sm Phytoplankton | 0 | 0 | 0 | 0 | 0 | 0 | 0 | 0 | 0 | 0 | 0 | 0 | 0 | 0 | 0 | 0 | 0 | 0 | 0 | 0 | 0 | 0 | 0 | 0 | 0 | 0 | 0 | 0 | 0 | 0 |
| Outside Production | 0 | 0 | 0 | 0 | 0 | 0 | 0 | 0 | 0 | 0 | 0 | 0 | 0 | 0 | 0 | 0 | 0 | 0 | 0 | 0 | 0 | 0 | 0 | 0 | 0 | 0 | 0 | 0 | 0 | 0 |
| Discards | 0 | 0 | 0 | 0 | 0 | 0 | 0 | 0 | 0 | 0 | 0 | 0 | 0 | 0 | 0 | 0 | 0 | 0 | 0 | 0 | 0 | 0 | 0 | 0 | 0 | 0 | 0 | 0 | 0 | 0 |
| Offal | 0.1 | 0 | 0.011 | 0 | 0.073 | 0 | 0 | 0 | 0.048 | 0 | 0.003 | 0 | 0 | 0 | 0.017 | 0.003 | 0.011 | 0 | 0.04 | 0 | 0 | 0 | 0 | 0 | 0 | 0.154 | 0.001 | 0 | 0.039 | 0.001 |
| Pelagic Detritus | 0 | 0 | 0 | 0 | 0 | 0 | 0 | 0 | 0 | 0 | 0 | 0 | 0 | 0 | 0 | 0 | 0 | 0 | 0 | 0 | 0 | 0 | 0 | 0 | 0 | 0 | 0 | 0 | 0 | 0 |
| Benthic Detritus | 0 | 0 | 0 | 0 | 0 | 0 | 0 | 0 | 0 | 0 | 0 | 0 | 0 | 0 | 0 | 0 | 0 | 0 | 0 | 0 | 0 | 0 | 0 | 0 | 0 | 0 | 0 | 0 | 0 | 0 |
| Outside Detritus | 0 | 0 | 0 | 0 | 0 | 0 | 0 | 0 | 0 | 0 | 0 | 0 | 0 | 0 | 0 | 0 | 0 | 0 | 0 | 0 | 0 | 0 | 0 | 0 | 0 | 0 | 0 | 0 | 0 | 0 |

**Table S3:** Diet matrix for the East Bering Sea – continued

| Group | Grenadiers | Misc. fish deep | POP | Sharpchin Rock | Northern Rock | Dusky Rock | Shortraker Rock | Rougheye Rock | Shortspine Thorns | Other Sebastes | Atka mackerel Juv | Atka mackerel Ad | Greenlings | Lg. Sculpins | Other sculpins | Misc. fish shallow | Octopi | Squids | Salmon returning | Salmon outgoing | Bathylagidae | Myctophidae | Capelin | Sandlance | Eulachon | Oth. managed forage | Oth. pelagic smelt | Bairdi | King Crab | Opilio |
|---|---|---|---|---|---|---|---|---|---|---|---|---|---|---|---|---|---|---|---|---|---|---|---|---|---|---|---|---|---|---|
| Transient KIllers | 0 | 0 | 0 | 0 | 0 | 0 | 0 | 0 | 0 | 0 | 0 | 0 | 0 | 0 | 0 | 0 | 0 | 0 | 0 | 0 | 0 | 0 | 0 | 0 | 0 | 0 | 0 | 0 | 0 | 0 |
| Sperm and Beaked Whales | 0 | 0 | 0 | 0 | 0 | 0 | 0 | 0 | 0 | 0 | 0 | 0 | 0 | 0 | 0 | 0 | 0 | 0 | 0 | 0 | 0 | 0 | 0 | 0 | 0 | 0 | 0 | 0 | 0 | 0 |
| Resident Killers | 0 | 0 | 0 | 0 | 0 | 0 | 0 | 0 | 0 | 0 | 0 | 0 | 0 | 0 | 0 | 0 | 0 | 0 | 0 | 0 | 0 | 0 | 0 | 0 | 0 | 0 | 0 | 0 | 0 | 0 |
| Porpoises | 0 | 0 | 0 | 0 | 0 | 0 | 0 | 0 | 0 | 0 | 0 | 0 | 0 | 0 | 0 | 0 | 0 | 0 | 0 | 0 | 0 | 0 | 0 | 0 | 0 | 0 | 0 | 0 | 0 | 0 |
| Belugas | 0 | 0 | 0 | 0 | 0 | 0 | 0 | 0 | 0 | 0 | 0 | 0 | 0 | 0 | 0 | 0 | 0 | 0 | 0 | 0 | 0 | 0 | 0 | 0 | 0 | 0 | 0 | 0 | 0 | 0 |
| Gray Whales | 0 | 0 | 0 | 0 | 0 | 0 | 0 | 0 | 0 | 0 | 0 | 0 | 0 | 0 | 0 | 0 | 0 | 0 | 0 | 0 | 0 | 0 | 0 | 0 | 0 | 0 | 0 | 0 | 0 | 0 |
| Humpbacks | 0 | 0 | 0 | 0 | 0 | 0 | 0 | 0 | 0 | 0 | 0 | 0 | 0 | 0 | 0 | 0 | 0 | 0 | 0 | 0 | 0 | 0 | 0 | 0 | 0 | 0 | 0 | 0 | 0 | 0 |

| Group | Grenadiers | Misc. fish deep | POP | Sharpchin Rock | Northern Rock | Dusky Rock | Shortraker Rock | Rougheye Rock | Shortspine Thorns | Other Sebastes | Atka mackerel Juv | Atka mackerel Ad | Greenlings | Lg. Sculpins | Other sculpins | Misc. fish shallow | Octopi | Squids | Salmon returning | Salmon outgoing | Bathylagidae | Myctophidae | Capelin | Sandlance | Eulachon | Oth. managed forage | Oth. pelagic smelt | Bairdi | King Crab | Opilio |
|---|---|---|---|---|---|---|---|---|---|---|---|---|---|---|---|---|---|---|---|---|---|---|---|---|---|---|---|---|---|---|
| Fin Whales | 0 | 0 | 0 | 0 | 0 | 0 | 0 | 0 | 0 | 0 | 0 | 0 | 0 | 0 | 0 | 0 | 0 | 0 | 0 | 0 | 0 | 0 | 0 | 0 | 0 | 0 | 0 | 0 | 0 | 0 |
| Sei Whales | 0 | 0 | 0 | 0 | 0 | 0 | 0 | 0 | 0 | 0 | 0 | 0 | 0 | 0 | 0 | 0 | 0 | 0 | 0 | 0 | 0 | 0 | 0 | 0 | 0 | 0 | 0 | 0 | 0 | 0 |
| Right Whales | 0 | 0 | 0 | 0 | 0 | 0 | 0 | 0 | 0 | 0 | 0 | 0 | 0 | 0 | 0 | 0 | 0 | 0 | 0 | 0 | 0 | 0 | 0 | 0 | 0 | 0 | 0 | 0 | 0 | 0 |
| Minke Whales | 0 | 0 | 0 | 0 | 0 | 0 | 0 | 0 | 0 | 0 | 0 | 0 | 0 | 0 | 0 | 0 | 0 | 0 | 0 | 0 | 0 | 0 | 0 | 0 | 0 | 0 | 0 | 0 | 0 | 0 |
| Bowhead Whales | 0 | 0 | 0 | 0 | 0 | 0 | 0 | 0 | 0 | 0 | 0 | 0 | 0 | 0 | 0 | 0 | 0 | 0 | 0 | 0 | 0 | 0 | 0 | 0 | 0 | 0 | 0 | 0 | 0 | 0 |
| Sea Otters | 0 | 0 | 0 | 0 | 0 | 0 | 0 | 0 | 0 | 0 | 0 | 0 | 0 | 0 | 0 | 0 | 0 | 0 | 0 | 0 | 0 | 0 | 0 | 0 | 0 | 0 | 0 | 0 | 0 | 0 |
| Walrus Bd Seals | 0 | 0 | 0 | 0 | 0 | 0 | 0 | 0 | 0 | 0 | 0 | 0 | 0 | 0 | 0 | 0 | 0 | 0 | 0 | 0 | 0 | 0 | 0 | 0 | 0 | 0 | 0 | 0 | 0 | 0 |
| N Fur Seal Juv | 0 | 0 | 0 | 0 | 0 | 0 | 0 | 0 | 0 | 0 | 0 | 0 | 0 | 0 | 0 | 0 | 0 | 0 | 0 | 0 | 0 | 0 | 0 | 0 | 0 | 0 | 0 | 0 | 0 | 0 |
| N Fur Seal Ad | 0 | 0 | 0 | 0 | 0 | 0 | 0 | 0 | 0 | 0 | 0 | 0 | 0 | 0 | 0 | 0 | 0 | 0 | 0 | 0 | 0 | 0 | 0 | 0 | 0 | 0 | 0 | 0 | 0 | 0 |
| Steller Sea Lions Juv | 0 | 0 | 0 | 0 | 0 | 0 | 0 | 0 | 0 | 0 | 0 | 0 | 0 | 0 | 0 | 0 | 0 | 0 | 0 | 0 | 0 | 0 | 0 | 0 | 0 | 0 | 0 | 0 | 0 | 0 |
| Steller Sea Lions Ad | 0 | 0 | 0 | 0 | 0 | 0 | 0 | 0 | 0 | 0 | 0 | 0 | 0 | 0 | 0 | 0 | 0 | 0 | 0 | 0 | 0 | 0 | 0 | 0 | 0 | 0 | 0 | 0 | 0 | 0 |
| Resident Seals | 0 | 0 | 0 | 0 | 0 | 0 | 0 | 0 | 0 | 0 | 0 | 0 | 0 | 0 | 0 | 0 | 0 | 0 | 0 | 0 | 0 | 0 | 0 | 0 | 0 | 0 | 0 | 0 | 0 | 0 |
| Wintering Seals | 0 | 0 | 0 | 0 | 0 | 0 | 0 | 0 | 0 | 0 | 0 | 0 | 0 | 0 | 0 | 0 | 0 | 0 | 0 | 0 | 0 | 0 | 0 | 0 | 0 | 0 | 0 | 0 | 0 | 0 |
| Shearwater | 0 | 0 | 0 | 0 | 0 | 0 | 0 | 0 | 0 | 0 | 0 | 0 | 0 | 0 | 0 | 0 | 0 | 0 | 0 | 0 | 0 | 0 | 0 | 0 | 0 | 0 | 0 | 0 | 0 | 0 |
| Murres | 0 | 0 | 0 | 0 | 0 | 0 | 0 | 0 | 0 | 0 | 0 | 0 | 0 | 0 | 0 | 0 | 0 | 0 | 0 | 0 | 0 | 0 | 0 | 0 | 0 | 0 | 0 | 0 | 0 | 0 |
| Kittiwakes | 0 | 0 | 0 | 0 | 0 | 0 | 0 | 0 | 0 | 0 | 0 | 0 | 0 | 0 | 0 | 0 | 0 | 0 | 0 | 0 | 0 | 0 | 0 | 0 | 0 | 0 | 0 | 0 | 0 | 0 |
| Auklets | 0 | 0 | 0 | 0 | 0 | 0 | 0 | 0 | 0 | 0 | 0 | 0 | 0 | 0 | 0 | 0 | 0 | 0 | 0 | 0 | 0 | 0 | 0 | 0 | 0 | 0 | 0 | 0 | 0 | 0 |
| Puffins | 0 | 0 | 0 | 0 | 0 | 0 | 0 | 0 | 0 | 0 | 0 | 0 | 0 | 0 | 0 | 0 | 0 | 0 | 0 | 0 | 0 | 0 | 0 | 0 | 0 | 0 | 0 | 0 | 0 | 0 |
| Fulmars | 0 | 0 | 0 | 0 | 0 | 0 | 0 | 0 | 0 | 0 | 0 | 0 | 0 | 0 | 0 | 0 | 0 | 0 | 0 | 0 | 0 | 0 | 0 | 0 | 0 | 0 | 0 | 0 | 0 | 0 |
| Storm Petrels | 0 | 0 | 0 | 0 | 0 | 0 | 0 | 0 | 0 | 0 | 0 | 0 | 0 | 0 | 0 | 0 | 0 | 0 | 0 | 0 | 0 | 0 | 0 | 0 | 0 | 0 | 0 | 0 | 0 | 0 |
| Cormorants | 0 | 0 | 0 | 0 | 0 | 0 | 0 | 0 | 0 | 0 | 0 | 0 | 0 | 0 | 0 | 0 | 0 | 0 | 0 | 0 | 0 | 0 | 0 | 0 | 0 | 0 | 0 | 0 | 0 | 0 |
| Gulls | 0 | 0 | 0 | 0 | 0 | 0 | 0 | 0 | 0 | 0 | 0 | 0 | 0 | 0 | 0 | 0 | 0 | 0 | 0 | 0 | 0 | 0 | 0 | 0 | 0 | 0 | 0 | 0 | 0 | 0 |
| Albatross Jaeger | 0 | 0 | 0 | 0 | 0 | 0 | 0 | 0 | 0 | 0 | 0 | 0 | 0 | 0 | 0 | 0 | 0 | 0 | 0 | 0 | 0 | 0 | 0 | 0 | 0 | 0 | 0 | 0 | 0 | 0 |
| Sleeper shark | 0 | 0 | 0 | 0 | 0 | 0 | 0 | 0 | 0 | 0 | 0 | 0 | 0 | 0 | 0 | 0 | 0 | 0 | 0 | 0 | 0 | 0 | 0 | 0 | 0 | 0 | 0 | 0 | 0 | 0 |
| W. Pollock Juv | 0 | 0 | 0 | 0 | 0 | 0 | 0 | 0 | 0 | 0 | 0 | 0 | 0.047 | 0.095 | 0 | 0 | 0 | 0 | 0 | 0 | 0 | 0 | 0 | 0 | 0 | 0 | 0 | 0 | 0 | 0 |
| W. Pollock Ad | 0 | 0 | 0 | 0 | 0 | 0 | 0 | 0 | 0 | 0 | 0 | 0 | 0 | 0.035 | 0 | 0 | 0 | 0 | 0 | 0 | 0 | 0 | 0 | 0 | 0 | 0 | 0 | 0 | 0 | 0 |
| P. Cod Juv | 0 | 0 | 0 | 0 | 0 | 0 | 0 | 0 | 0 | 0 | 0 | 0 | 0 | 0 | 0.04 | 0 | 0 | 0 | 0 | 0 | 0 | 0 | 0 | 0 | 0 | 0 | 0 | 0 | 0 | 0 |
| P. Cod Ad | 0 | 0 | 0 | 0 | 0 | 0 | 0 | 0 | 0 | 0 | 0 | 0 | 0 | 0.001 | 0 | 0 | 0 | 0 | 0 | 0 | 0 | 0 | 0 | 0 | 0 | 0 | 0 | 0 | 0 | 0 |
| Herring Juv | 0 | 0 | 0 | 0 | 0 | 0 | 0 | 0 | 0 | 0 | 0 | 0 | 0 | 0.005 | 0 | 0 | 0 | 0 | 0 | 0 | 0 | 0 | 0 | 0 | 0 | 0 | 0 | 0 | 0 | 0 |
| Herring Ad | 0 | 0 | 0 | 0 | 0 | 0 | 0 | 0 | 0 | 0 | 0 | 0 | 0 | 0.002 | 0 | 0 | 0 | 0 | 0 | 0 | 0 | 0 | 0 | 0 | 0 | 0 | 0 | 0 | 0 | 0 |
| Arrowtooth Juv | 0 | 0 | 0 | 0 | 0 | 0 | 0 | 0 | 0 | 0 | 0 | 0 | 0 | 0 | 0 | 0 | 0 | 0 | 0 | 0 | 0 | 0 | 0 | 0 | 0 | 0 | 0 | 0 | 0 | 0 |
| Arrowtooth Ad | 0 | 0 | 0 | 0 | 0 | 0 | 0 | 0 | 0 | 0 | 0 | 0 | 0 | 0 | 0 | 0 | 0 | 0 | 0 | 0 | 0 | 0 | 0 | 0 | 0 | 0 | 0 | 0 | 0 | 0 |
| Kamchatka fl. Juv | 0 | 0 | 0 | 0 | 0 | 0 | 0 | 0 | 0 | 0 | 0 | 0 | 0 | 0 | 0 | 0 | 0 | 0 | 0 | 0 | 0 | 0 | 0 | 0 | 0 | 0 | 0 | 0 | 0 | 0 |
| Kamchatka fl. Ad | 0 | 0 | 0 | 0 | 0 | 0 | 0 | 0 | 0 | 0 | 0 | 0 | 0 | 0 | 0 | 0 | 0 | 0 | 0 | 0 | 0 | 0 | 0 | 0 | 0 | 0 | 0 | 0 | 0 | 0 |
| Gr. Turbot Juv | 0 | 0 | 0 | 0 | 0 | 0 | 0 | 0 | 0 | 0 | 0 | 0 | 0 | 0 | 0 | 0 | 0 | 0 | 0 | 0 | 0 | 0 | 0 | 0 | 0 | 0 | 0 | 0 | 0 | 0 |
| Gr. Turbot Ad | 0 | 0 | 0 | 0 | 0 | 0 | 0 | 0 | 0 | 0 | 0 | 0 | 0 | 0 | 0 | 0 | 0 | 0 | 0 | 0 | 0 | 0 | 0 | 0 | 0 | 0 | 0 | 0 | 0 | 0 |

| Group | Grenadiers | Misc. fish deep | POP | Sharpchin Rock | Northern Rock | Dusky Rock | Shortraker Rock | Rougheye Rock | Shortspine Thorns | Other Sebastes | Atka mackerel Juv | Atka mackerel Ad | Greenlings | Lg. Sculpins | Other sculpins | Misc. fish shallow | Octopi | Squids | Salmon returning | Salmon outgoing | Bathylagidae | Myctophidae | Capelin | Sandlance | Eulachon | Oth. managed forage | Oth. pelagic smelt | Bairdi | King Crab | Opilio |
|---|---|---|---|---|---|---|---|---|---|---|---|---|---|---|---|---|---|---|---|---|---|---|---|---|---|---|---|---|---|---|
| P. Halibut Juv | 0 | 0 | 0 | 0 | 0 | 0 | 0 | 0 | 0 | 0 | 0 | 0 | 0 | 0 | 0 | 0 | 0 | 0 | 0 | 0 | 0 | 0 | 0 | 0 | 0 | 0 | 0 | 0 | 0 | 0 |
| P. Halibut Ad | 0 | 0 | 0 | 0 | 0 | 0 | 0 | 0 | 0 | 0 | 0 | 0 | 0 | 0 | 0 | 0 | 0 | 0 | 0 | 0 | 0 | 0 | 0 | 0 | 0 | 0 | 0 | 0 | 0 | 0 |
| YF. Sole Juv | 0 | 0 | 0 | 0 | 0 | 0 | 0 | 0 | 0 | 0 | 0 | 0 | 0 | 0.022 | 0.01 | 0 | 0 | 0 | 0 | 0 | 0 | 0 | 0 | 0 | 0 | 0 | 0 | 0 | 0 | 0 |
| YF. Sole Ad | 0 | 0 | 0 | 0 | 0 | 0 | 0 | 0 | 0 | 0 | 0 | 0 | 0 | 0.005 | 0 | 0 | 0 | 0 | 0 | 0 | 0 | 0 | 0 | 0 | 0 | 0 | 0 | 0 | 0 | 0 |
| FH. Sole Juv | 0 | 0 | 0 | 0 | 0 | 0 | 0 | 0 | 0 | 0 | 0 | 0 | 0 | 0 | 0 | 0.007 | 0 | 0 | 0 | 0 | 0 | 0 | 0 | 0 | 0 | 0 | 0 | 0 | 0 | 0 |
| FH. Sole Ad | 0 | 0 | 0 | 0 | 0 | 0 | 0 | 0 | 0 | 0 | 0 | 0 | 0 | 0.002 | 0 | 0 | 0 | 0 | 0 | 0 | 0 | 0 | 0 | 0 | 0 | 0 | 0 | 0 | 0 | 0 |
| N. Rock sole Juv | 0 | 0 | 0 | 0 | 0 | 0 | 0 | 0 | 0 | 0 | 0 | 0 | 0 | 0.007 | 0 | 0 | 0 | 0 | 0 | 0 | 0 | 0 | 0 | 0 | 0 | 0 | 0 | 0 | 0 | 0 |
| N. Rock sole Ad | 0 | 0 | 0 | 0 | 0 | 0 | 0 | 0 | 0 | 0 | 0 | 0 | 0 | 0.002 | 0 | 0 | 0 | 0 | 0 | 0 | 0 | 0 | 0 | 0 | 0 | 0 | 0 | 0 | 0 | 0 |
| AK Plaice | 0 | 0 | 0 | 0 | 0 | 0 | 0 | 0 | 0 | 0 | 0 | 0 | 0 | 0.001 | 0 | 0 | 0 | 0 | 0 | 0 | 0 | 0 | 0 | 0 | 0 | 0 | 0 | 0 | 0 | 0 |
| Dover Sole | 0 | 0 | 0 | 0 | 0 | 0 | 0 | 0 | 0 | 0 | 0 | 0 | 0 | 0 | 0 | 0 | 0 | 0 | 0 | 0 | 0 | 0 | 0 | 0 | 0 | 0 | 0 | 0 | 0 | 0 |
| Rex Sole | 0 | 0 | 0 | 0 | 0 | 0 | 0 | 0 | 0 | 0 | 0 | 0 | 0 | 0 | 0 | 0 | 0 | 0 | 0 | 0 | 0 | 0 | 0 | 0 | 0 | 0 | 0 | 0 | 0 | 0 |
| Misc. Flatfish | 0 | 0 | 0 | 0 | 0 | 0 | 0 | 0 | 0 | 0 | 0 | 0 | 0.03 | 0.001 | 0 | 0 | 0 | 0 | 0 | 0 | 0 | 0 | 0 | 0 | 0 | 0 | 0 | 0 | 0 | 0 |
| Alaska skate | 0 | 0 | 0 | 0 | 0 | 0 | 0 | 0 | 0 | 0 | 0 | 0 | 0 | 0 | 0 | 0 | 0 | 0 | 0 | 0 | 0 | 0 | 0 | 0 | 0 | 0 | 0 | 0 | 0 | 0 |
| Other skates | 0 | 0 | 0 | 0 | 0 | 0 | 0 | 0 | 0 | 0 | 0 | 0 | 0 | 0 | 0 | 0 | 0 | 0 | 0 | 0 | 0 | 0 | 0 | 0 | 0 | 0 | 0 | 0 | 0 | 0 |
| Sablefish Juv | 0 | 0 | 0 | 0 | 0 | 0 | 0 | 0 | 0 | 0 | 0 | 0 | 0 | 0 | 0 | 0 | 0 | 0 | 0 | 0 | 0 | 0 | 0 | 0 | 0 | 0 | 0 | 0 | 0 | 0 |
| Sablefish Ad | 0 | 0 | 0 | 0 | 0 | 0 | 0 | 0 | 0 | 0 | 0 | 0 | 0 | 0 | 0 | 0 | 0 | 0 | 0 | 0 | 0 | 0 | 0 | 0 | 0 | 0 | 0 | 0 | 0 | 0 |
| Eelpouts | 0 | 0 | 0 | 0 | 0 | 0 | 0 | 0 | 0 | 0 | 0 | 0 | 0 | 0.033 | 0 | 0 | 0 | 0 | 0 | 0 | 0 | 0 | 0 | 0 | 0 | 0 | 0 | 0 | 0 | 0 |
| Grenadiers | 0 | 0 | 0 | 0 | 0 | 0 | 0 | 0 | 0 | 0 | 0 | 0 | 0 | 0 | 0 | 0 | 0 | 0 | 0 | 0 | 0 | 0 | 0 | 0 | 0 | 0 | 0 | 0 | 0 | 0 |
| Misc. fish deep | 0 | 0 | 0.004 | 0 | 0 | 0 | 0 | 0 | 0 | 0 | 0 | 0 | 0 | 0 | 0 | 0 | 0 | 0 | 0 | 0 | 0 | 0 | 0 | 0 | 0 | 0 | 0 | 0 | 0 | 0 |
| POP | 0 | 0 | 0 | 0 | 0 | 0 | 0 | 0 | 0 | 0 | 0 | 0 | 0 | 0 | 0 | 0 | 0 | 0 | 0 | 0 | 0 | 0 | 0 | 0 | 0 | 0 | 0 | 0 | 0 | 0 |
| Sharpchin Rock | 0 | 0 | 0 | 0 | 0 | 0 | 0 | 0 | 0 | 0 | 0 | 0 | 0 | 0 | 0 | 0 | 0 | 0 | 0 | 0 | 0 | 0 | 0 | 0 | 0 | 0 | 0 | 0 | 0 | 0 |
| Northern Rock | 0 | 0 | 0 | 0 | 0 | 0 | 0 | 0 | 0 | 0 | 0 | 0 | 0 | 0 | 0 | 0 | 0 | 0 | 0 | 0 | 0 | 0 | 0 | 0 | 0 | 0 | 0 | 0 | 0 | 0 |
| Dusky Rock | 0 | 0 | 0 | 0 | 0 | 0 | 0 | 0 | 0 | 0 | 0 | 0 | 0 | 0 | 0 | 0 | 0 | 0 | 0 | 0 | 0 | 0 | 0 | 0 | 0 | 0 | 0 | 0 | 0 | 0 |
| Shortraker Rock | 0 | 0 | 0 | 0 | 0 | 0 | 0 | 0 | 0 | 0 | 0 | 0 | 0 | 0 | 0 | 0 | 0 | 0 | 0 | 0 | 0 | 0 | 0 | 0 | 0 | 0 | 0 | 0 | 0 | 0 |
| Rougheye Rock | 0 | 0 | 0 | 0 | 0 | 0 | 0 | 0 | 0 | 0 | 0 | 0 | 0 | 0 | 0 | 0 | 0 | 0 | 0 | 0 | 0 | 0 | 0 | 0 | 0 | 0 | 0 | 0 | 0 | 0 |
| Shortspine Thorns | 0 | 0 | 0 | 0 | 0 | 0 | 0 | 0 | 0 | 0 | 0 | 0 | 0 | 0 | 0 | 0 | 0 | 0 | 0 | 0 | 0 | 0 | 0 | 0 | 0 | 0 | 0 | 0 | 0 | 0 |
| Other Sebastes | 0 | 0 | 0 | 0 | 0 | 0 | 0 | 0 | 0 | 0 | 0 | 0 | 0 | 0 | 0 | 0 | 0 | 0 | 0 | 0 | 0 | 0 | 0 | 0 | 0 | 0 | 0 | 0 | 0 | 0 |
| Atka mackerel Juv | 0 | 0 | 0 | 0 | 0 | 0 | 0 | 0 | 0 | 0 | 0 | 0 | 0 | 0 | 0 | 0 | 0 | 0 | 0 | 0 | 0 | 0 | 0 | 0 | 0 | 0 | 0 | 0 | 0 | 0 |
| Atka mackerel Ad | 0 | 0 | 0 | 0 | 0 | 0 | 0 | 0 | 0 | 0 | 0 | 0 | 0 | 0 | 0 | 0 | 0 | 0 | 0 | 0 | 0 | 0 | 0 | 0 | 0 | 0 | 0 | 0 | 0 | 0 |
| Greenlings | 0 | 0 | 0 | 0 | 0 | 0 | 0 | 0 | 0 | 0 | 0 | 0 | 0 | 0 | 0 | 0 | 0 | 0 | 0 | 0 | 0 | 0 | 0 | 0 | 0 | 0 | 0 | 0 | 0 | 0 |
| Lg. Sculpins | 0 | 0 | 0 | 0 | 0 | 0 | 0 | 0 | 0 | 0 | 0 | 0 | 0.178 | 0.002 | 0 | 0 | 0 | 0 | 0 | 0 | 0 | 0 | 0 | 0 | 0 | 0 | 0 | 0 | 0 | 0 |
| Other sculpins | 0 | 0 | 0 | 0.001 | 0 | 0 | 0.006 | 0 | 0 | 0.001 | 0 | 0 | 0.15 | 0.021 | 0 | 0 | 0 | 0 | 0 | 0 | 0 | 0 | 0 | 0 | 0 | 0 | 0 | 0 | 0 | 0 |
| Misc. fish shallow | 0 | 0 | 0 | 0 | 0 | 0 | 0 | 0.001 | 0 | 0 | 0 | 0 | 0 | 0.019 | 0.05 | 0 | 0 | 0 | 0 | 0 | 0 | 0 | 0 | 0 | 0 | 0 | 0 | 0 | 0 | 0 |
| Octopi | 0 | 0 | 0 | 0 | 0 | 0 | 0 | 0 | 0 | 0 | 0 | 0 | 0 | 0.014 | 0 | 0 | 0 | 0 | 0 | 0 | 0 | 0 | 0 | 0 | 0 | 0 | 0 | 0 | 0 | 0 |
| Squids | 0.5 | 0.5 | 0 | 0.131 | 0.054 | 0 | 0.009 | 0.62 | 0 | 0.131 | 0 | 0 | 0 | 0 | 0 | 0 | 0 | 0 | 0.2 | 0 | 0 | 0 | 0 | 0 | 0 | 0 | 0 | 0 | 0 | 0 |
| Salmon returning | 0 | 0 | 0 | 0 | 0 | 0 | 0 | 0 | 0 | 0 | 0 | 0 | 0 | 0 | 0 | 0 | 0 | 0 | 0 | 0 | 0 | 0 | 0 | 0 | 0 | 0 | 0 | 0 | 0 | 0 |
| Salmon outgoing | 0 | 0 | 0 | 0 | 0 | 0 | 0 | 0 | 0 | 0 | 0 | 0 | 0 | 0 | 0 | 0 | 0 | 0 | 0 | 0 | 0 | 0 | 0 | 0 | 0 | 0 | 0 | 0 | 0 | 0 |
| Bathylagidae | 0 | 0 | 0.007 | 0 | 0 | 0 | 0 | 0 | 0 | 0 | 0 | 0 | 0 | 0 | 0 | 0 | 0 | 0 | 0 | 0 | 0 | 0 | 0 | 0 | 0 | 0 | 0 | 0 | 0 | 0 |

| Group | Grenadiers | Misc. fish deep | POP | Sharpchin Rock | Northern Rock | Dusky Rock | Shortraker Rock | Rougheye Rock | Shortspine Thorns | Other Sebastes | Atka mackerel Juv | Atka mackerel Ad | Greenlings | Lg. Sculpins | Other sculpins | Misc. fish shallow | Octopi | Squids | Salmon returning | Salmon outgoing | Bathylagidae | Myctophidae | Capelin | Sandlance | Eulachon | Oth. managed forage | Oth. pelagic smelt | Bairdi | King Crab | Opilio |
|---|---|---|---|---|---|---|---|---|---|---|---|---|---|---|---|---|---|---|---|---|---|---|---|---|---|---|---|---|---|---|
| Myctophidae | 0 | 0 | 0.01 | 0.001 | 0 | 0 | 0.003 | 0.002 | 0 | 0.001 | 0 | 0 | 0 | 0 | 0 | 0 | 0 | 0.025 | 0 | 0 | 0 | 0 | 0 | 0 | 0 | 0 | 0 | 0 | 0 | 0 |
| Capelin | 0 | 0 | 0 | 0 | 0 | 0 | 0 | 0 | 0 | 0 | 0 | 0 | 0 | 0.012 | 0 | 0 | 0 | 0.025 | 0 | 0 | 0 | 0 | 0 | 0 | 0 | 0 | 0 | 0 | 0 | 0 |
| Sandlance | 0 | 0 | 0 | 0 | 0 | 0 | 0 | 0 | 0 | 0 | 0 | 0 | 0 | 0.009 | 0 | 0 | 0 | 0.025 | 0 | 0 | 0 | 0 | 0 | 0 | 0 | 0 | 0 | 0 | 0 | 0 |
| Eulachon | 0 | 0 | 0 | 0 | 0 | 0 | 0 | 0 | 0 | 0 | 0 | 0 | 0 | 0 | 0 | 0 | 0 | 0.025 | 0 | 0 | 0 | 0 | 0 | 0 | 0 | 0 | 0 | 0 | 0 | 0 |
| Oth. managed forage | 0 | 0 | 0 | 0 | 0 | 0 | 0 | 0 | 0 | 0 | 0 | 0 | 0 | 0.058 | 0 | 0 | 0 | 0.025 | 0 | 0 | 0 | 0 | 0 | 0 | 0 | 0 | 0 | 0 | 0 | 0 |
| Oth. pelagic smelt | 0 | 0 | 0 | 0 | 0 | 0 | 0 | 0 | 0 | 0 | 0 | 0 | 0 | 0 | 0 | 0 | 0 | 0.025 | 0 | 0 | 0 | 0 | 0 | 0 | 0 | 0 | 0 | 0 | 0 | 0 |
| Bairdi | 0 | 0 | 0 | 0 | 0 | 0 | 0 | 0 | 0 | 0 | 0 | 0.006 | 0 | 0.005 | 0.035 | 0 | 0.05 | 0 | 0 | 0 | 0 | 0 | 0 | 0 | 0 | 0 | 0 | 0 | 0 | 0 |
| King Crab | 0 | 0 | 0 | 0 | 0 | 0 | 0 | 0 | 0 | 0 | 0 | 0 | 0 | 0.018 | 0 | 0 | 0 | 0 | 0 | 0 | 0 | 0 | 0 | 0 | 0 | 0 | 0 | 0 | 0 | 0 |
| Opilio | 0 | 0 | 0 | 0 | 0 | 0 | 0 | 0 | 0 | 0 | 0 | 0 | 0 | 0.039 | 0 | 0 | 0.05 | 0 | 0 | 0 | 0 | 0 | 0 | 0 | 0 | 0 | 0 | 0 | 0 | 0 |
| Pandalidae | 0 | 0 | 0 | 0.167 | 0 | 0 | 0.83 | 0.001 | 0.002 | 0.167 | 0 | 0 | 0 | 0.007 | 0.042 | 0.019 | 0 | 0 | 0 | 0 | 0 | 0 | 0 | 0 | 0 | 0 | 0 | 0 | 0 | 0 |
| NP shrimp | 0.5 | 0.5 | 0 | 0.064 | 0.018 | 0 | 0 | 0.138 | 0.153 | 0.064 | 0 | 0 | 0.43 | 0.19 | 0.501 | 0.207 | 0 | 0 | 0 | 0 | 0 | 0 | 0 | 0 | 0 | 0 | 0 | 0 | 0 | 0 |
| Sea stars | 0 | 0 | 0 | 0 | 0 | 0 | 0 | 0 | 0 | 0 | 0 | 0 | 0 | 0 | 0 | 0 | 0 | 0 | 0 | 0 | 0 | 0 | 0 | 0 | 0 | 0 | 0 | 0 | 0.006 | 0.001 |
| Brittle stars | 0 | 0 | 0 | 0 | 0 | 0 | 0 | 0 | 0 | 0 | 0 | 0 | 0 | 0 | 0 | 0 | 0 | 0 | 0 | 0 | 0 | 0 | 0 | 0 | 0 | 0 | 0 | 0.083 | 0.027 | 0.072 |
| Urchins dollars cucumbers | 0 | 0 | 0 | 0 | 0 | 0 | 0 | 0 | 0 | 0 | 0 | 0 | 0 | 0 | 0 | 0 | 0 | 0 | 0 | 0 | 0 | 0 | 0 | 0 | 0 | 0 | 0 | 0.015 | 0.166 | 0.007 |
| Snails | 0 | 0 | 0 | 0 | 0 | 0 | 0 | 0 | 0 | 0 | 0 | 0 | 0 | 0.001 | 0 | 0 | 0.4 | 0 | 0 | 0 | 0 | 0 | 0 | 0 | 0 | 0 | 0 | 0.038 | 0.198 | 0.048 |
| Hermit crabs | 0 | 0 | 0 | 0.004 | 0.011 | 0 | 0 | 0 | 0 | 0.004 | 0 | 0 | 0 | 0.019 | 0 | 0.001 | 0.1 | 0 | 0 | 0 | 0 | 0 | 0 | 0 | 0 | 0 | 0 | 0.014 | 0.016 | 0.03 |
| Misc. crabs | 0 | 0 | 0 | 0 | 0 | 0 | 0 | 0 | 0 | 0 | 0 | 0 | 0.004 | 0.019 | 0.019 | 0 | 0 | 0 | 0 | 0 | 0 | 0 | 0 | 0 | 0 | 0 | 0 | 0.014 | 0.016 | 0.03 |
| Misc. Crustacean | 0 | 0 | 0 | 0 | 0 | 0 | 0 | 0 | 0 | 0 | 0 | 0 | 0.001 | 0.001 | 0.014 | 0 | 0 | 0 | 0 | 0 | 0 | 0 | 0 | 0 | 0 | 0 | 0 | 0.005 | 0.015 | 0.017 |
| Benthic Amphipods | 0 | 0 | 0.001 | 0.085 | 0 | 0.046 | 0 | 0.014 | 0.362 | 0.085 | 0 | 0 | 0.032 | 0.06 | 0.195 | 0.092 | 0 | 0 | 0 | 0 | 0 | 0 | 0 | 0 | 0 | 0 | 0 | 0.038 | 0.009 | 0.048 |
| Anemones | 0 | 0 | 0 | 0 | 0 | 0 | 0 | 0 | 0 | 0 | 0 | 0 | 0 | 0 | 0 | 0 | 0 | 0 | 0 | 0 | 0 | 0 | 0 | 0 | 0 | 0 | 0 | 0 | 0 | 0.002 |
| Corals | 0 | 0 | 0 | 0 | 0 | 0 | 0 | 0 | 0 | 0 | 0 | 0 | 0 | 0 | 0 | 0 | 0 | 0 | 0 | 0 | 0 | 0 | 0 | 0 | 0 | 0 | 0 | 0 | 0 | 0 |
| Hydroids | 0 | 0 | 0 | 0 | 0 | 0 | 0 | 0 | 0 | 0 | 0 | 0 | 0.001 | 0 | 0.037 | 0 | 0 | 0 | 0 | 0 | 0 | 0 | 0 | 0 | 0 | 0 | 0 | 0.012 | 0.046 | 0.006 |
| Urochordata | 0 | 0 | 0 | 0 | 0 | 0 | 0 | 0 | 0 | 0 | 0 | 0 | 0 | 0 | 0 | 0 | 0 | 0 | 0 | 0 | 0 | 0 | 0 | 0 | 0 | 0 | 0 | 0 | 0.009 | 0 |
| Sea Pens | 0 | 0 | 0 | 0 | 0 | 0 | 0 | 0 | 0 | 0 | 0 | 0 | 0 | 0 | 0 | 0 | 0 | 0 | 0 | 0 | 0 | 0 | 0 | 0 | 0 | 0 | 0 | 0 | 0 | 0 |
| Sponges | 0 | 0 | 0 | 0 | 0 | 0 | 0 | 0 | 0 | 0 | 0 | 0 | 0 | 0 | 0 | 0 | 0 | 0 | 0 | 0 | 0 | 0 | 0 | 0 | 0 | 0 | 0 | 0.002 | 0 | 0.003 |
| Bivalves | 0 | 0 | 0 | 0 | 0 | 0 | 0 | 0 | 0 | 0 | 0 | 0 | 0.012 | 0 | 0 | 0 | 0.4 | 0 | 0 | 0 | 0 | 0 | 0 | 0 | 0 | 0 | 0 | 0.308 | 0.184 | 0.252 |
| Polychaetes | 0 | 0 | 0 | 0.017 | 0 | 0.084 | 0 | 0 | 0 | 0.017 | 0 | 0 | 0.071 | 0.11 | 0.026 | 0 | 0 | 0 | 0 | 0 | 0 | 0 | 0 | 0 | 0 | 0 | 0 | 0.384 | 0.189 | 0.285 |
| Misc. worms | 0 | 0 | 0 | 0 | 0 | 0 | 0 | 0 | 0 | 0 | 0 | 0 | 0 | 0.019 | 0 | 0 | 0 | 0 | 0 | 0 | 0 | 0 | 0 | 0 | 0 | 0 | 0 | 0 | 0.026 | 0.024 |
| Scyphozoid Jellies | 0 | 0 | 0 | 0 | 0 | 0 | 0 | 0 | 0 | 0 | 0 | 0 | 0 | 0 | 0 | 0.064 | 0 | 0 | 0 | 0 | 0 | 0 | 0 | 0 | 0 | 0 | 0 | 0 | 0 | 0 |
| Fish Larvae | 0 | 0 | 0 | 0 | 0 | 0 | 0 | 0 | 0 | 0 | 0 | 0 | 0 | 0 | 0 | 0 | 0 | 0 | 0 | 0 | 0 | 0 | 0 | 0 | 0 | 0 | 0 | 0 | 0 | 0 |
| Chaetognaths | 0 | 0 | 0.002 | 0.016 | 0.058 | 0 | 0 | 0 | 0 | 0.016 | 0 | 0 | 0 | 0 | 0 | 0 | 0 | 0.025 | 0.2 | 0 | 0 | 0 | 0 | 0 | 0 | 0 | 0 | 0 | 0 | 0 |
| Euphausiids | 0 | 0 | 0.213 | 0.188 | 0.04 | 0.68 | 0.02 | 0.194 | 0 | 0.188 | 0.9 | 0.984 | 0 | 0.01 | 0 | 0.012 | 0 | 0.42 | 0.2 | 0.25 | 0.9 | 0.9 | 0.9 | 0.9 | 0.9 | 0.9 | 0.9 | 0 | 0 | 0.001 |
| Mysids | 0 | 0 | 0.712 | 0.129 | 0.005 | 0 | 0.132 | 0.026 | 0.483 | 0.129 | 0 | 0 | 0.043 | 0.076 | 0.003 | 0.035 | 0 | 0.025 | 0 | 0 | 0 | 0 | 0 | 0 | 0 | 0 | 0 | 0 | 0 | 0.004 |
| Pelagic Amphipods | 0 | 0 | 0 | 0 | 0.001 | 0 | 0 | 0 | 0 | 0 | 0 | 0 | 0.001 | 0.046 | 0.008 | 0.014 | 0 | 0.14 | 0 | 0.25 | 0 | 0 | 0 | 0 | 0 | 0 | 0 | 0 | 0 | 0 |

| Group | Grenadiers | Misc. fish deep | POP | Sharpchin Rock | Northern Rock | Dusky Rock | Shortraker Rock | Rougheye Rock | Shortspine Thorns | Other Sebastes | Atka mackerel Juv | Atka mackerel Ad | Greenlings | Lg. Sculpins | Other sculpins | Misc. fish shallow | Octopi | Squids | Salmon returning | Salmon outgoing | Bathylagidae | Myctophidae | Capelin | Sandlance | Eulachon | Oth. managed forage | Oth. pelagic smelt | Bairdi | King Crab | Opilio |
|---|---|---|---|---|---|---|---|---|---|---|---|---|---|---|---|---|---|---|---|---|---|---|---|---|---|---|---|---|---|---|
| Gelatinous filter feeders | 0 | 0 | 0 | 0 | 0 | 0 | 0 | 0 | 0 | 0 | 0 | 0.004 | 0 | 0.025 | 0 | 0.55 | 0 | 0.04 | 0 | 0 | 0 | 0 | 0 | 0 | 0 | 0 | 0 | 0 | 0 | 0 |
| Pteropods | 0 | 0 | 0 | 0.001 | 0.002 | 0 | 0 | 0 | 0 | 0.001 | 0 | 0 | 0 | 0 | 0 | 0 | 0 | 0 | 0.2 | 0 | 0 | 0 | 0 | 0 | 0 | 0 | 0 | 0 | 0 | 0 |
| Copepods | 0 | 0 | 0.051 | 0.197 | 0.809 | 0.19 | 0 | 0.004 | 0 | 0.197 | 0.1 | 0.006 | 0 | 0 | 0.019 | 0 | 0 | 0.2 | 0.2 | 0.5 | 0.1 | 0.1 | 0.1 | 0.1 | 0.1 | 0.1 | 0.1 | 0 | 0 | 0 |
| Pelagic microbes | 0 | 0 | 0 | 0 | 0 | 0 | 0 | 0 | 0 | 0 | 0 | 0 | 0 | 0 | 0 | 0 | 0 | 0 | 0 | 0 | 0 | 0 | 0 | 0 | 0 | 0 | 0 | 0 | 0 | 0 |
| Benthic microbes | 0 | 0 | 0 | 0 | 0 | 0 | 0 | 0 | 0 | 0 | 0 | 0 | 0 | 0 | 0 | 0 | 0 | 0 | 0 | 0 | 0 | 0 | 0 | 0 | 0 | 0 | 0 | 0 | 0 | 0 |
| Macroalgae | 0 | 0 | 0 | 0 | 0 | 0 | 0 | 0 | 0 | 0 | 0 | 0 | 0 | 0 | 0 | 0 | 0 | 0 | 0 | 0 | 0 | 0 | 0 | 0 | 0 | 0 | 0 | 0 | 0.025 | 0.004 |
| Lg Phytoplankton | 0 | 0 | 0 | 0 | 0 | 0 | 0 | 0 | 0 | 0 | 0 | 0 | 0 | 0 | 0 | 0 | 0 | 0 | 0 | 0 | 0 | 0 | 0 | 0 | 0 | 0 | 0 | 0 | 0 | 0 |
| Sm Phytoplankton | 0 | 0 | 0 | 0 | 0 | 0 | 0 | 0 | 0 | 0 | 0 | 0 | 0 | 0 | 0 | 0 | 0 | 0 | 0 | 0 | 0 | 0 | 0 | 0 | 0 | 0 | 0 | 0 | 0 | 0 |
| Outside Production | 0 | 0 | 0 | 0 | 0 | 0 | 0 | 0 | 0 | 0 | 0 | 0 | 0 | 0 | 0 | 0 | 0 | 0 | 0 | 0 | 0 | 0 | 0 | 0 | 0 | 0 | 0 | 0 | 0 | 0 |
| Discards | 0 | 0 | 0 | 0 | 0 | 0 | 0 | 0 | 0 | 0 | 0 | 0 | 0 | 0 | 0 | 0 | 0 | 0 | 0 | 0 | 0 | 0 | 0 | 0 | 0 | 0 | 0 | 0.006 | 0.003 | 0.006 |
| Offal | 0 | 0 | 0 | 0 | 0 | 0 | 0 | 0 | 0 | 0 | 0 | 0 | 0 | 0.008 | 0 | 0 | 0 | 0 | 0 | 0 | 0 | 0 | 0 | 0 | 0 | 0 | 0 | 0.006 | 0.003 | 0.006 |
| Pelagic Detritus | 0 | 0 | 0 | 0 | 0 | 0 | 0 | 0 | 0 | 0 | 0 | 0 | 0 | 0 | 0 | 0 | 0 | 0 | 0 | 0 | 0 | 0 | 0 | 0 | 0 | 0 | 0 | 0 | 0 | 0 |
| Benthic Detritus | 0 | 0 | 0 | 0 | 0 | 0 | 0 | 0 | 0 | 0 | 0 | 0 | 0 | 0 | 0 | 0 | 0 | 0 | 0 | 0 | 0 | 0 | 0 | 0 | 0 | 0 | 0 | 0.078 | 0.06 | 0.154 |
| Outside Detritus | 0 | 0 | 0 | 0 | 0 | 0 | 0 | 0 | 0 | 0 | 0 | 0 | 0 | 0 | 0 | 0 | 0 | 0 | 0 | 0 | 0 | 0 | 0 | 0 | 0 | 0 | 0 | 0 | 0 | 0 |

**Table S3:** Diet matrix for the East Bering Sea – continued

| Group | Pandalidae | NP shrimp | Sea stars | Brittle stars | Urchins dollars cucumbers | Snails | Hermit crabs | Misc. crabs | Misc. Crustacean | Benthic Amphipods | Anemones | Corals | Hydroids | Urochordata | Sea Pens | Sponges | Bivalves | Polychaetes | Misc. worms | Scyphozoid Jellies | Fish Larvae | Chaetognaths | Euphausiids | Mysids | Pelagic Amphipods | Gelatinous filter feeders | Pteropods | Copepods | Pelagic microbes | Benthic microbes |
|---|---|---|---|---|---|---|---|---|---|---|---|---|---|---|---|---|---|---|---|---|---|---|---|---|---|---|---|---|---|---|
| Transient KIllers | 0 | 0 | 0 | 0 | 0 | 0 | 0 | 0 | 0 | 0 | 0 | 0 | 0 | 0 | 0 | 0 | 0 | 0 | 0 | 0 | 0 | 0 | 0 | 0 | 0 | 0 | 0 | 0 | 0 | 0 |
| Sperm and Beaked Whales | 0 | 0 | 0 | 0 | 0 | 0 | 0 | 0 | 0 | 0 | 0 | 0 | 0 | 0 | 0 | 0 | 0 | 0 | 0 | 0 | 0 | 0 | 0 | 0 | 0 | 0 | 0 | 0 | 0 | 0 |
| Resident Killers | 0 | 0 | 0 | 0 | 0 | 0 | 0 | 0 | 0 | 0 | 0 | 0 | 0 | 0 | 0 | 0 | 0 | 0 | 0 | 0 | 0 | 0 | 0 | 0 | 0 | 0 | 0 | 0 | 0 | 0 |
| Porpoises | 0 | 0 | 0 | 0 | 0 | 0 | 0 | 0 | 0 | 0 | 0 | 0 | 0 | 0 | 0 | 0 | 0 | 0 | 0 | 0 | 0 | 0 | 0 | 0 | 0 | 0 | 0 | 0 | 0 | 0 |
| Belugas | 0 | 0 | 0 | 0 | 0 | 0 | 0 | 0 | 0 | 0 | 0 | 0 | 0 | 0 | 0 | 0 | 0 | 0 | 0 | 0 | 0 | 0 | 0 | 0 | 0 | 0 | 0 | 0 | 0 | 0 |
| Gray Whales | 0 | 0 | 0 | 0 | 0 | 0 | 0 | 0 | 0 | 0 | 0 | 0 | 0 | 0 | 0 | 0 | 0 | 0 | 0 | 0 | 0 | 0 | 0 | 0 | 0 | 0 | 0 | 0 | 0 | 0 |

| Group | Pandalidae | NP shrimp | Sea stars | Brittle stars | Urchins dollars cucumbers | Snails | Hermit crabs | Misc. crabs | Misc. Crustacean | Benthic Amphipods | Anemones | Corals | Hydroids | Urochordata | Sea Pens | Sponges | Bivalves | Polychaetes | Misc. worms | Scyphozoid Jellies | Fish Larvae | Chaetognaths | Euphausiids | Mysids | Pelagic Amphipods | Gelatinous filter feeders | Pteropods | Copepods | Pelagic microbes | Benthic microbes |
|---|---|---|---|---|---|---|---|---|---|---|---|---|---|---|---|---|---|---|---|---|---|---|---|---|---|---|---|---|---|---|
| Humpbacks | 0 | 0 | 0 | 0 | 0 | 0 | 0 | 0 | 0 | 0 | 0 | 0 | 0 | 0 | 0 | 0 | 0 | 0 | 0 | 0 | 0 | 0 | 0 | 0 | 0 | 0 | 0 | 0 | 0 | 0 |
| Fin Whales | 0 | 0 | 0 | 0 | 0 | 0 | 0 | 0 | 0 | 0 | 0 | 0 | 0 | 0 | 0 | 0 | 0 | 0 | 0 | 0 | 0 | 0 | 0 | 0 | 0 | 0 | 0 | 0 | 0 | 0 |
| Sei Whales | 0 | 0 | 0 | 0 | 0 | 0 | 0 | 0 | 0 | 0 | 0 | 0 | 0 | 0 | 0 | 0 | 0 | 0 | 0 | 0 | 0 | 0 | 0 | 0 | 0 | 0 | 0 | 0 | 0 | 0 |
| Right Whales | 0 | 0 | 0 | 0 | 0 | 0 | 0 | 0 | 0 | 0 | 0 | 0 | 0 | 0 | 0 | 0 | 0 | 0 | 0 | 0 | 0 | 0 | 0 | 0 | 0 | 0 | 0 | 0 | 0 | 0 |
| Minke Whales | 0 | 0 | 0 | 0 | 0 | 0 | 0 | 0 | 0 | 0 | 0 | 0 | 0 | 0 | 0 | 0 | 0 | 0 | 0 | 0 | 0 | 0 | 0 | 0 | 0 | 0 | 0 | 0 | 0 | 0 |
| Bowhead Whales | 0 | 0 | 0 | 0 | 0 | 0 | 0 | 0 | 0 | 0 | 0 | 0 | 0 | 0 | 0 | 0 | 0 | 0 | 0 | 0 | 0 | 0 | 0 | 0 | 0 | 0 | 0 | 0 | 0 | 0 |
| Sea Otters | 0 | 0 | 0 | 0 | 0 | 0 | 0 | 0 | 0 | 0 | 0 | 0 | 0 | 0 | 0 | 0 | 0 | 0 | 0 | 0 | 0 | 0 | 0 | 0 | 0 | 0 | 0 | 0 | 0 | 0 |
| Walrus Bd Seals | 0 | 0 | 0 | 0 | 0 | 0 | 0 | 0 | 0 | 0 | 0 | 0 | 0 | 0 | 0 | 0 | 0 | 0 | 0 | 0 | 0 | 0 | 0 | 0 | 0 | 0 | 0 | 0 | 0 | 0 |
| N Fur Seal Juv | 0 | 0 | 0 | 0 | 0 | 0 | 0 | 0 | 0 | 0 | 0 | 0 | 0 | 0 | 0 | 0 | 0 | 0 | 0 | 0 | 0 | 0 | 0 | 0 | 0 | 0 | 0 | 0 | 0 | 0 |
| N Fur Seal Ad | 0 | 0 | 0 | 0 | 0 | 0 | 0 | 0 | 0 | 0 | 0 | 0 | 0 | 0 | 0 | 0 | 0 | 0 | 0 | 0 | 0 | 0 | 0 | 0 | 0 | 0 | 0 | 0 | 0 | 0 |
| Steller Sea Lions Juv | 0 | 0 | 0 | 0 | 0 | 0 | 0 | 0 | 0 | 0 | 0 | 0 | 0 | 0 | 0 | 0 | 0 | 0 | 0 | 0 | 0 | 0 | 0 | 0 | 0 | 0 | 0 | 0 | 0 | 0 |
| Steller Sea Lions Ad | 0 | 0 | 0 | 0 | 0 | 0 | 0 | 0 | 0 | 0 | 0 | 0 | 0 | 0 | 0 | 0 | 0 | 0 | 0 | 0 | 0 | 0 | 0 | 0 | 0 | 0 | 0 | 0 | 0 | 0 |
| Resident Seals | 0 | 0 | 0 | 0 | 0 | 0 | 0 | 0 | 0 | 0 | 0 | 0 | 0 | 0 | 0 | 0 | 0 | 0 | 0 | 0 | 0 | 0 | 0 | 0 | 0 | 0 | 0 | 0 | 0 | 0 |
| Wintering Seals | 0 | 0 | 0 | 0 | 0 | 0 | 0 | 0 | 0 | 0 | 0 | 0 | 0 | 0 | 0 | 0 | 0 | 0 | 0 | 0 | 0 | 0 | 0 | 0 | 0 | 0 | 0 | 0 | 0 | 0 |
| Shearwater | 0 | 0 | 0 | 0 | 0 | 0 | 0 | 0 | 0 | 0 | 0 | 0 | 0 | 0 | 0 | 0 | 0 | 0 | 0 | 0 | 0 | 0 | 0 | 0 | 0 | 0 | 0 | 0 | 0 | 0 |
| Murres | 0 | 0 | 0 | 0 | 0 | 0 | 0 | 0 | 0 | 0 | 0 | 0 | 0 | 0 | 0 | 0 | 0 | 0 | 0 | 0 | 0 | 0 | 0 | 0 | 0 | 0 | 0 | 0 | 0 | 0 |
| Kittiwakes | 0 | 0 | 0 | 0 | 0 | 0 | 0 | 0 | 0 | 0 | 0 | 0 | 0 | 0 | 0 | 0 | 0 | 0 | 0 | 0 | 0 | 0 | 0 | 0 | 0 | 0 | 0 | 0 | 0 | 0 |
| Auklets | 0 | 0 | 0 | 0 | 0 | 0 | 0 | 0 | 0 | 0 | 0 | 0 | 0 | 0 | 0 | 0 | 0 | 0 | 0 | 0 | 0 | 0 | 0 | 0 | 0 | 0 | 0 | 0 | 0 | 0 |
| Puffins | 0 | 0 | 0 | 0 | 0 | 0 | 0 | 0 | 0 | 0 | 0 | 0 | 0 | 0 | 0 | 0 | 0 | 0 | 0 | 0 | 0 | 0 | 0 | 0 | 0 | 0 | 0 | 0 | 0 | 0 |
| Fulmars | 0 | 0 | 0 | 0 | 0 | 0 | 0 | 0 | 0 | 0 | 0 | 0 | 0 | 0 | 0 | 0 | 0 | 0 | 0 | 0 | 0 | 0 | 0 | 0 | 0 | 0 | 0 | 0 | 0 | 0 |
| Storm Petrels | 0 | 0 | 0 | 0 | 0 | 0 | 0 | 0 | 0 | 0 | 0 | 0 | 0 | 0 | 0 | 0 | 0 | 0 | 0 | 0 | 0 | 0 | 0 | 0 | 0 | 0 | 0 | 0 | 0 | 0 |
| Cormorants | 0 | 0 | 0 | 0 | 0 | 0 | 0 | 0 | 0 | 0 | 0 | 0 | 0 | 0 | 0 | 0 | 0 | 0 | 0 | 0 | 0 | 0 | 0 | 0 | 0 | 0 | 0 | 0 | 0 | 0 |
| Gulls | 0 | 0 | 0 | 0 | 0 | 0 | 0 | 0 | 0 | 0 | 0 | 0 | 0 | 0 | 0 | 0 | 0 | 0 | 0 | 0 | 0 | 0 | 0 | 0 | 0 | 0 | 0 | 0 | 0 | 0 |
| Albatross Jaeger | 0 | 0 | 0 | 0 | 0 | 0 | 0 | 0 | 0 | 0 | 0 | 0 | 0 | 0 | 0 | 0 | 0 | 0 | 0 | 0 | 0 | 0 | 0 | 0 | 0 | 0 | 0 | 0 | 0 | 0 |
| Sleeper shark | 0 | 0 | 0 | 0 | 0 | 0 | 0 | 0 | 0 | 0 | 0 | 0 | 0 | 0 | 0 | 0 | 0 | 0 | 0 | 0 | 0 | 0 | 0 | 0 | 0 | 0 | 0 | 0 | 0 | 0 |
| W. Pollock Juv | 0 | 0 | 0 | 0 | 0 | 0 | 0 | 0 | 0 | 0 | 0 | 0 | 0 | 0 | 0 | 0 | 0 | 0 | 0 | 0.001 | 0 | 0 | 0 | 0 | 0 | 0 | 0 | 0 | 0 | 0 |
| W. Pollock Ad | 0 | 0 | 0 | 0 | 0 | 0 | 0 | 0 | 0 | 0 | 0 | 0 | 0 | 0 | 0 | 0 | 0 | 0 | 0 | 0 | 0 | 0 | 0 | 0 | 0 | 0 | 0 | 0 | 0 | 0 |
| P. Cod Juv | 0 | 0 | 0 | 0 | 0 | 0 | 0 | 0 | 0 | 0 | 0 | 0 | 0 | 0 | 0 | 0 | 0 | 0 | 0 | 0 | 0 | 0 | 0 | 0 | 0 | 0 | 0 | 0 | 0 | 0 |
| P. Cod Ad | 0 | 0 | 0 | 0 | 0 | 0 | 0 | 0 | 0 | 0 | 0 | 0 | 0 | 0 | 0 | 0 | 0 | 0 | 0 | 0 | 0 | 0 | 0 | 0 | 0 | 0 | 0 | 0 | 0 | 0 |
| Herring Juv | 0 | 0 | 0 | 0 | 0 | 0 | 0 | 0 | 0 | 0 | 0 | 0 | 0 | 0 | 0 | 0 | 0 | 0 | 0 | 0.001 | 0 | 0 | 0 | 0 | 0 | 0 | 0 | 0 | 0 | 0 |
| Herring Ad | 0 | 0 | 0 | 0 | 0 | 0 | 0 | 0 | 0 | 0 | 0 | 0 | 0 | 0 | 0 | 0 | 0 | 0 | 0 | 0 | 0 | 0 | 0 | 0 | 0 | 0 | 0 | 0 | 0 | 0 |
| Arrowtooth Juv | 0 | 0 | 0 | 0 | 0 | 0 | 0 | 0 | 0 | 0 | 0 | 0 | 0 | 0 | 0 | 0 | 0 | 0 | 0 | 0 | 0 | 0 | 0 | 0 | 0 | 0 | 0 | 0 | 0 | 0 |
| Arrowtooth Ad | 0 | 0 | 0 | 0 | 0 | 0 | 0 | 0 | 0 | 0 | 0 | 0 | 0 | 0 | 0 | 0 | 0 | 0 | 0 | 0 | 0 | 0 | 0 | 0 | 0 | 0 | 0 | 0 | 0 | 0 |
| Kamchatka fl. Juv | 0 | 0 | 0 | 0 | 0 | 0 | 0 | 0 | 0 | 0 | 0 | 0 | 0 | 0 | 0 | 0 | 0 | 0 | 0 | 0 | 0 | 0 | 0 | 0 | 0 | 0 | 0 | 0 | 0 | 0 |
| Kamchatka fl. Ad | 0 | 0 | 0 | 0 | 0 | 0 | 0 | 0 | 0 | 0 | 0 | 0 | 0 | 0 | 0 | 0 | 0 | 0 | 0 | 0 | 0 | 0 | 0 | 0 | 0 | 0 | 0 | 0 | 0 | 0 |

| Group | Pandalidae | NP shrimp | Sea stars | Brittle stars | Urchins dollars cucumbers | Snails | Hermit crabs | Misc. crabs | Misc. Crustacean | Benthic Amphipods | Anemones | Corals | Hydroids | Urochordata | Sea Pens | Sponges | Bivalves | Polychaetes | Misc. worms | Scyphozoid Jellies | Fish Larvae | Chaetognaths | Euphausiids | Mysids | Pelagic Amphipods | Gelatinous filter feeders | Pteropods | Copepods | Pelagic microbes | Benthic microbes |
|---|---|---|---|---|---|---|---|---|---|---|---|---|---|---|---|---|---|---|---|---|---|---|---|---|---|---|---|---|---|---|
| Gr. Turbot Juv | 0 | 0 | 0 | 0 | 0 | 0 | 0 | 0 | 0 | 0 | 0 | 0 | 0 | 0 | 0 | 0 | 0 | 0 | 0 | 0 | 0 | 0 | 0 | 0 | 0 | 0 | 0 | 0 | 0 | 0 |
| Gr. Turbot Ad | 0 | 0 | 0 | 0 | 0 | 0 | 0 | 0 | 0 | 0 | 0 | 0 | 0 | 0 | 0 | 0 | 0 | 0 | 0 | 0 | 0 | 0 | 0 | 0 | 0 | 0 | 0 | 0 | 0 | 0 |
| P. Halibut Juv | 0 | 0 | 0 | 0 | 0 | 0 | 0 | 0 | 0 | 0 | 0 | 0 | 0 | 0 | 0 | 0 | 0 | 0 | 0 | 0 | 0 | 0 | 0 | 0 | 0 | 0 | 0 | 0 | 0 | 0 |
| P. Halibut Ad | 0 | 0 | 0 | 0 | 0 | 0 | 0 | 0 | 0 | 0 | 0 | 0 | 0 | 0 | 0 | 0 | 0 | 0 | 0 | 0 | 0 | 0 | 0 | 0 | 0 | 0 | 0 | 0 | 0 | 0 |
| YF. Sole Juv | 0 | 0 | 0 | 0 | 0 | 0 | 0 | 0 | 0 | 0 | 0 | 0 | 0 | 0 | 0 | 0 | 0 | 0 | 0 | 0 | 0 | 0 | 0 | 0 | 0 | 0 | 0 | 0 | 0 | 0 |
| YF. Sole Ad | 0 | 0 | 0 | 0 | 0 | 0 | 0 | 0 | 0 | 0 | 0 | 0 | 0 | 0 | 0 | 0 | 0 | 0 | 0 | 0 | 0 | 0 | 0 | 0 | 0 | 0 | 0 | 0 | 0 | 0 |
| FH. Sole Juv | 0 | 0 | 0 | 0 | 0 | 0 | 0 | 0 | 0 | 0 | 0 | 0 | 0 | 0 | 0 | 0 | 0 | 0 | 0 | 0 | 0 | 0 | 0 | 0 | 0 | 0 | 0 | 0 | 0 | 0 |
| FH. Sole Ad | 0 | 0 | 0 | 0 | 0 | 0 | 0 | 0 | 0 | 0 | 0 | 0 | 0 | 0 | 0 | 0 | 0 | 0 | 0 | 0 | 0 | 0 | 0 | 0 | 0 | 0 | 0 | 0 | 0 | 0 |
| N. Rock sole Juv | 0 | 0 | 0 | 0 | 0 | 0 | 0 | 0 | 0 | 0 | 0 | 0 | 0 | 0 | 0 | 0 | 0 | 0 | 0 | 0 | 0 | 0 | 0 | 0 | 0 | 0 | 0 | 0 | 0 | 0 |
| N. Rock sole Ad | 0 | 0 | 0 | 0 | 0 | 0 | 0 | 0 | 0 | 0 | 0 | 0 | 0 | 0 | 0 | 0 | 0 | 0 | 0 | 0 | 0 | 0 | 0 | 0 | 0 | 0 | 0 | 0 | 0 | 0 |
| AK Plaice | 0 | 0 | 0 | 0 | 0 | 0 | 0 | 0 | 0 | 0 | 0 | 0 | 0 | 0 | 0 | 0 | 0 | 0 | 0 | 0 | 0 | 0 | 0 | 0 | 0 | 0 | 0 | 0 | 0 | 0 |
| Dover Sole | 0 | 0 | 0 | 0 | 0 | 0 | 0 | 0 | 0 | 0 | 0 | 0 | 0 | 0 | 0 | 0 | 0 | 0 | 0 | 0 | 0 | 0 | 0 | 0 | 0 | 0 | 0 | 0 | 0 | 0 |
| Rex Sole | 0 | 0 | 0 | 0 | 0 | 0 | 0 | 0 | 0 | 0 | 0 | 0 | 0 | 0 | 0 | 0 | 0 | 0 | 0 | 0 | 0 | 0 | 0 | 0 | 0 | 0 | 0 | 0 | 0 | 0 |
| Misc. Flatfish | 0 | 0 | 0 | 0 | 0 | 0 | 0 | 0 | 0 | 0 | 0 | 0 | 0 | 0 | 0 | 0 | 0 | 0 | 0 | 0 | 0 | 0 | 0 | 0 | 0 | 0 | 0 | 0 | 0 | 0 |
| Alaska skate | 0 | 0 | 0 | 0 | 0 | 0 | 0 | 0 | 0 | 0 | 0 | 0 | 0 | 0 | 0 | 0 | 0 | 0 | 0 | 0 | 0 | 0 | 0 | 0 | 0 | 0 | 0 | 0 | 0 | 0 |
| Other skates | 0 | 0 | 0 | 0 | 0 | 0 | 0 | 0 | 0 | 0 | 0 | 0 | 0 | 0 | 0 | 0 | 0 | 0 | 0 | 0 | 0 | 0 | 0 | 0 | 0 | 0 | 0 | 0 | 0 | 0 |
| Sablefish Juv | 0 | 0 | 0 | 0 | 0 | 0 | 0 | 0 | 0 | 0 | 0 | 0 | 0 | 0 | 0 | 0 | 0 | 0 | 0 | 0 | 0 | 0 | 0 | 0 | 0 | 0 | 0 | 0 | 0 | 0 |
| Sablefish Ad | 0 | 0 | 0 | 0 | 0 | 0 | 0 | 0 | 0 | 0 | 0 | 0 | 0 | 0 | 0 | 0 | 0 | 0 | 0 | 0 | 0 | 0 | 0 | 0 | 0 | 0 | 0 | 0 | 0 | 0 |
| Eelpouts | 0 | 0 | 0 | 0 | 0 | 0 | 0 | 0 | 0 | 0 | 0 | 0 | 0 | 0 | 0 | 0 | 0 | 0 | 0 | 0 | 0 | 0 | 0 | 0 | 0 | 0 | 0 | 0 | 0 | 0 |
| Grenadiers | 0 | 0 | 0 | 0 | 0 | 0 | 0 | 0 | 0 | 0 | 0 | 0 | 0 | 0 | 0 | 0 | 0 | 0 | 0 | 0 | 0 | 0 | 0 | 0 | 0 | 0 | 0 | 0 | 0 | 0 |
| Misc. fish deep | 0 | 0 | 0 | 0 | 0 | 0 | 0 | 0 | 0 | 0 | 0 | 0 | 0 | 0 | 0 | 0 | 0 | 0 | 0 | 0 | 0 | 0 | 0 | 0 | 0 | 0 | 0 | 0 | 0 | 0 |
| POP | 0 | 0 | 0 | 0 | 0 | 0 | 0 | 0 | 0 | 0 | 0 | 0 | 0 | 0 | 0 | 0 | 0 | 0 | 0 | 0 | 0 | 0 | 0 | 0 | 0 | 0 | 0 | 0 | 0 | 0 |
| Sharpchin Rock | 0 | 0 | 0 | 0 | 0 | 0 | 0 | 0 | 0 | 0 | 0 | 0 | 0 | 0 | 0 | 0 | 0 | 0 | 0 | 0 | 0 | 0 | 0 | 0 | 0 | 0 | 0 | 0 | 0 | 0 |
| Northern Rock | 0 | 0 | 0 | 0 | 0 | 0 | 0 | 0 | 0 | 0 | 0 | 0 | 0 | 0 | 0 | 0 | 0 | 0 | 0 | 0 | 0 | 0 | 0 | 0 | 0 | 0 | 0 | 0 | 0 | 0 |
| Dusky Rock | 0 | 0 | 0 | 0 | 0 | 0 | 0 | 0 | 0 | 0 | 0 | 0 | 0 | 0 | 0 | 0 | 0 | 0 | 0 | 0 | 0 | 0 | 0 | 0 | 0 | 0 | 0 | 0 | 0 | 0 |
| Shortraker Rock | 0 | 0 | 0 | 0 | 0 | 0 | 0 | 0 | 0 | 0 | 0 | 0 | 0 | 0 | 0 | 0 | 0 | 0 | 0 | 0 | 0 | 0 | 0 | 0 | 0 | 0 | 0 | 0 | 0 | 0 |
| Rougheye Rock | 0 | 0 | 0 | 0 | 0 | 0 | 0 | 0 | 0 | 0 | 0 | 0 | 0 | 0 | 0 | 0 | 0 | 0 | 0 | 0 | 0 | 0 | 0 | 0 | 0 | 0 | 0 | 0 | 0 | 0 |
| Shortspine Thorns | 0 | 0 | 0 | 0 | 0 | 0 | 0 | 0 | 0 | 0 | 0 | 0 | 0 | 0 | 0 | 0 | 0 | 0 | 0 | 0 | 0 | 0 | 0 | 0 | 0 | 0 | 0 | 0 | 0 | 0 |
| Other Sebastes | 0 | 0 | 0 | 0 | 0 | 0 | 0 | 0 | 0 | 0 | 0 | 0 | 0 | 0 | 0 | 0 | 0 | 0 | 0 | 0 | 0 | 0 | 0 | 0 | 0 | 0 | 0 | 0 | 0 | 0 |
| Atka mackerel Juv | 0 | 0 | 0 | 0 | 0 | 0 | 0 | 0 | 0 | 0 | 0 | 0 | 0 | 0 | 0 | 0 | 0 | 0 | 0 | 0 | 0 | 0 | 0 | 0 | 0 | 0 | 0 | 0 | 0 | 0 |
| Atka mackerel Ad | 0 | 0 | 0 | 0 | 0 | 0 | 0 | 0 | 0 | 0 | 0 | 0 | 0 | 0 | 0 | 0 | 0 | 0 | 0 | 0 | 0 | 0 | 0 | 0 | 0 | 0 | 0 | 0 | 0 | 0 |
| Greenlings | 0 | 0 | 0 | 0 | 0 | 0 | 0 | 0 | 0 | 0 | 0 | 0 | 0 | 0 | 0 | 0 | 0 | 0 | 0 | 0 | 0 | 0 | 0 | 0 | 0 | 0 | 0 | 0 | 0 | 0 |
| Lg. Sculpins | 0 | 0 | 0 | 0 | 0 | 0 | 0 | 0 | 0 | 0 | 0 | 0 | 0 | 0 | 0 | 0 | 0 | 0 | 0 | 0 | 0 | 0 | 0 | 0 | 0 | 0 | 0 | 0 | 0 | 0 |
| Other sculpins | 0 | 0 | 0 | 0 | 0 | 0 | 0 | 0 | 0 | 0 | 0 | 0 | 0 | 0 | 0 | 0 | 0 | 0 | 0 | 0 | 0 | 0 | 0 | 0 | 0 | 0 | 0 | 0 | 0 | 0 |
| Misc. fish shallow | 0 | 0 | 0 | 0 | 0 | 0 | 0 | 0 | 0 | 0 | 0 | 0 | 0 | 0 | 0 | 0 | 0 | 0 | 0 | 0 | 0 | 0 | 0 | 0 | 0 | 0 | 0 | 0 | 0 | 0 |

| Group | Pandalidae | NP shrimp | Sea stars | Brittle stars | Urchins dollars cucumbers | Snails | Hermit crabs | Misc. crabs | Misc. Crustacean | Benthic Amphipods | Anemones | Corals | Hydroids | Urochordata | Sea Pens | Sponges | Bivalves | Polychaetes | Misc. worms | Scyphozoid Jellies | Fish Larvae | Chaetognaths | Euphausiids | Mysids | Pelagic Amphipods | Gelatinous filter feeders | Pteropods | Copepods | Pelagic microbes | Benthic microbes |
|---|---|---|---|---|---|---|---|---|---|---|---|---|---|---|---|---|---|---|---|---|---|---|---|---|---|---|---|---|---|---|
| Octopi | 0 | 0 | 0 | 0 | 0 | 0 | 0 | 0 | 0 | 0 | 0 | 0 | 0 | 0 | 0 | 0 | 0 | 0 | 0 | 0 | 0 | 0 | 0 | 0 | 0 | 0 | 0 | 0 | 0 | 0 |
| Squids | 0 | 0 | 0 | 0 | 0 | 0 | 0 | 0 | 0 | 0 | 0 | 0 | 0 | 0 | 0 | 0 | 0 | 0 | 0 | 0.001 | 0 | 0 | 0 | 0 | 0 | 0 | 0 | 0 | 0 | 0 |
| Salmon returning | 0 | 0 | 0 | 0 | 0 | 0 | 0 | 0 | 0 | 0 | 0 | 0 | 0 | 0 | 0 | 0 | 0 | 0 | 0 | 0 | 0 | 0 | 0 | 0 | 0 | 0 | 0 | 0 | 0 | 0 |
| Salmon outgoing | 0 | 0 | 0 | 0 | 0 | 0 | 0 | 0 | 0 | 0 | 0 | 0 | 0 | 0 | 0 | 0 | 0 | 0 | 0 | 0 | 0 | 0 | 0 | 0 | 0 | 0 | 0 | 0 | 0 | 0 |
| Bathylagidae | 0 | 0 | 0 | 0 | 0 | 0 | 0 | 0 | 0 | 0 | 0 | 0 | 0 | 0 | 0 | 0 | 0 | 0 | 0 | 0.001 | 0 | 0 | 0 | 0 | 0 | 0 | 0 | 0 | 0 | 0 |
| Myctophidae | 0 | 0 | 0 | 0 | 0 | 0 | 0 | 0 | 0 | 0 | 0 | 0 | 0 | 0 | 0 | 0 | 0 | 0 | 0 | 0.001 | 0 | 0 | 0 | 0 | 0 | 0 | 0 | 0 | 0 | 0 |
| Capelin | 0 | 0 | 0 | 0 | 0 | 0 | 0 | 0 | 0 | 0 | 0 | 0 | 0 | 0 | 0 | 0 | 0 | 0 | 0 | 0.001 | 0 | 0 | 0 | 0 | 0 | 0 | 0 | 0 | 0 | 0 |
| Sandlance | 0 | 0 | 0 | 0 | 0 | 0 | 0 | 0 | 0 | 0 | 0 | 0 | 0 | 0 | 0 | 0 | 0 | 0 | 0 | 0.001 | 0 | 0 | 0 | 0 | 0 | 0 | 0 | 0 | 0 | 0 |
| Eulachon | 0 | 0 | 0 | 0 | 0 | 0 | 0 | 0 | 0 | 0 | 0 | 0 | 0 | 0 | 0 | 0 | 0 | 0 | 0 | 0.001 | 0 | 0 | 0 | 0 | 0 | 0 | 0 | 0 | 0 | 0 |
| Oth. managed forage | 0 | 0 | 0 | 0 | 0 | 0 | 0 | 0 | 0 | 0 | 0 | 0 | 0 | 0 | 0 | 0 | 0 | 0 | 0 | 0.001 | 0 | 0 | 0 | 0 | 0 | 0 | 0 | 0 | 0 | 0 |
| Oth. pelagic smelt | 0 | 0 | 0 | 0 | 0 | 0 | 0 | 0 | 0 | 0 | 0 | 0 | 0 | 0 | 0 | 0 | 0 | 0 | 0 | 0.001 | 0 | 0 | 0 | 0 | 0 | 0 | 0 | 0 | 0 | 0 |
| Bairdi | 0 | 0 | 0.005 | 0 | 0 | 0 | 0 | 0 | 0 | 0 | 0 | 0 | 0 | 0 | 0 | 0 | 0 | 0 | 0 | 0 | 0 | 0 | 0 | 0 | 0 | 0 | 0 | 0 | 0 | 0 |
| King Crab | 0 | 0 | 0 | 0 | 0 | 0 | 0 | 0 | 0 | 0 | 0 | 0 | 0 | 0 | 0 | 0 | 0 | 0 | 0 | 0 | 0 | 0 | 0 | 0 | 0 | 0 | 0 | 0 | 0 | 0 |
| Opilio | 0 | 0 | 0.005 | 0 | 0 | 0 | 0 | 0 | 0 | 0 | 0 | 0 | 0 | 0 | 0 | 0 | 0 | 0 | 0 | 0 | 0 | 0 | 0 | 0 | 0 | 0 | 0 | 0 | 0 | 0 |
| Pandalidae | 0 | 0 | 0.005 | 0 | 0 | 0 | 0 | 0 | 0 | 0 | 0 | 0 | 0 | 0 | 0 | 0 | 0 | 0 | 0 | 0 | 0 | 0 | 0 | 0 | 0 | 0 | 0 | 0 | 0 | 0 |
| NP shrimp | 0 | 0 | 0.005 | 0 | 0 | 0 | 0 | 0 | 0 | 0 | 0 | 0 | 0 | 0 | 0 | 0 | 0 | 0 | 0 | 0 | 0 | 0 | 0 | 0 | 0 | 0 | 0 | 0 | 0 | 0 |
| Sea stars | 0 | 0 | 0 | 0 | 0 | 0 | 0 | 0 | 0 | 0 | 0 | 0 | 0 | 0 | 0 | 0 | 0 | 0 | 0 | 0 | 0 | 0 | 0 | 0 | 0 | 0 | 0 | 0 | 0 | 0 |
| Brittle stars | 0 | 0 | 0.03 | 0 | 0 | 0 | 0 | 0 | 0 | 0 | 0 | 0 | 0 | 0 | 0 | 0 | 0 | 0 | 0 | 0 | 0 | 0 | 0 | 0 | 0 | 0 | 0 | 0 | 0 | 0 |
| Urchins dollars cucumbers | 0 | 0 | 0 | 0 | 0 | 0 | 0 | 0 | 0 | 0 | 0 | 0 | 0 | 0 | 0 | 0 | 0 | 0 | 0 | 0 | 0 | 0 | 0 | 0 | 0 | 0 | 0 | 0 | 0 | 0 |
| Snails | 0 | 0 | 0.005 | 0 | 0 | 0 | 0 | 0 | 0 | 0 | 0 | 0 | 0 | 0 | 0 | 0 | 0 | 0 | 0 | 0 | 0 | 0 | 0 | 0 | 0 | 0 | 0 | 0 | 0 | 0 |
| Hermit crabs | 0 | 0 | 0.005 | 0 | 0 | 0 | 0 | 0 | 0 | 0 | 0 | 0 | 0 | 0 | 0 | 0 | 0 | 0 | 0 | 0 | 0 | 0 | 0 | 0 | 0 | 0 | 0 | 0 | 0 | 0 |
| Misc. crabs | 0 | 0 | 0 | 0 | 0 | 0 | 0 | 0 | 0 | 0 | 0 | 0 | 0 | 0 | 0 | 0 | 0 | 0 | 0 | 0 | 0 | 0 | 0 | 0 | 0 | 0 | 0 | 0 | 0 | 0 |
| Misc. Crustacean | 0 | 0 | 0 | 0 | 0 | 0 | 0 | 0 | 0 | 0 | 0.125 | 0.125 | 0.125 | 0.125 | 0.125 | 0.125 | 0.125 | 0 | 0 | 0 | 0 | 0 | 0 | 0 | 0 | 0 | 0 | 0 | 0 | 0 |
| Benthic Amphipods | 0.3 | 0.3 | 0 | 0.1 | 0 | 0 | 0 | 0 | 0 | 0 | 0.125 | 0.125 | 0.125 | 0.125 | 0.125 | 0.125 | 0.125 | 0 | 0 | 0 | 0 | 0 | 0 | 0 | 0 | 0 | 0 | 0 | 0 | 0 |
| Anemones | 0 | 0 | 0 | 0 | 0 | 0 | 0 | 0 | 0 | 0 | 0 | 0 | 0 | 0 | 0 | 0 | 0 | 0 | 0 | 0 | 0 | 0 | 0 | 0 | 0 | 0 | 0 | 0 | 0 | 0 |
| Corals | 0 | 0 | 0 | 0 | 0 | 0 | 0 | 0 | 0 | 0 | 0 | 0 | 0 | 0 | 0 | 0 | 0 | 0 | 0 | 0 | 0 | 0 | 0 | 0 | 0 | 0 | 0 | 0 | 0 | 0 |
| Hydroids | 0 | 0 | 0 | 0 | 0 | 0 | 0 | 0 | 0 | 0 | 0 | 0 | 0 | 0 | 0 | 0 | 0 | 0 | 0 | 0 | 0 | 0 | 0 | 0 | 0 | 0 | 0 | 0 | 0 | 0 |
| Urochordata | 0 | 0 | 0 | 0 | 0 | 0 | 0 | 0 | 0 | 0 | 0 | 0 | 0 | 0 | 0 | 0 | 0 | 0 | 0 | 0 | 0 | 0 | 0 | 0 | 0 | 0 | 0 | 0 | 0 | 0 |
| Sea Pens | 0 | 0 | 0 | 0 | 0 | 0 | 0 | 0 | 0 | 0 | 0 | 0 | 0 | 0 | 0 | 0 | 0 | 0 | 0 | 0 | 0 | 0 | 0 | 0 | 0 | 0 | 0 | 0 | 0 | 0 |
| Sponges | 0 | 0 | 0 | 0 | 0 | 0 | 0 | 0 | 0 | 0 | 0 | 0 | 0 | 0 | 0 | 0 | 0 | 0 | 0 | 0 | 0 | 0 | 0 | 0 | 0 | 0 | 0 | 0 | 0 | 0 |
| Bivalves | 0.1 | 0.1 | 0.91 | 0 | 0 | 0.2 | 0.25 | 0.25 | 0 | 0 | 0 | 0 | 0 | 0 | 0 | 0 | 0 | 0 | 0 | 0 | 0 | 0 | 0 | 0 | 0 | 0 | 0 | 0 | 0 | 0 |
| Polychaetes | 0 | 0 | 0.02 | 0 | 0 | 0.2 | 0.25 | 0.25 | 0 | 0 | 0 | 0 | 0 | 0 | 0 | 0 | 0 | 0 | 0 | 0 | 0 | 0 | 0 | 0 | 0 | 0 | 0 | 0 | 0 | 0 |
| Misc. worms | 0 | 0 | 0.01 | 0 | 0 | 0.2 | 0.25 | 0.25 | 0 | 0 | 0 | 0 | 0 | 0 | 0 | 0 | 0 | 0 | 0 | 0 | 0 | 0 | 0 | 0 | 0 | 0 | 0 | 0 | 0 | 0 |
| Scyphozoid Jellies | 0 | 0 | 0 | 0 | 0 | 0 | 0 | 0 | 0 | 0 | 0 | 0 | 0 | 0 | 0 | 0 | 0 | 0 | 0 | 0 | 0 | 0 | 0 | 0 | 0 | 0 | 0 | 0 | 0 | 0 |
| Fish Larvae | 0 | 0 | 0 | 0 | 0 | 0 | 0 | 0 | 0 | 0 | 0 | 0 | 0 | 0 | 0 | 0 | 0 | 0 | 0 | 0.05 | 0 | 0 | 0 | 0 | 0 | 0 | 0 | 0 | 0 | 0 |

| Group | Pandalidae | NP shrimp | Sea stars | Brittle stars | Urchins dollars cucumbers | Snails | Hermit crabs | Misc. crabs | Misc. Crustacean | Benthic Amphipods | Anemones | Corals | Hydroids | Urochordata | Sea Pens | Sponges | Bivalves | Polychaetes | Misc. worms | Scyphozoid Jellies | Fish Larvae | Chaetognaths | Euphausiids | Mysids | Pelagic Amphipods | Gelatinous filter feeders | Pteropods | Copepods | Pelagic microbes | Benthic microbes |
|---|---|---|---|---|---|---|---|---|---|---|---|---|---|---|---|---|---|---|---|---|---|---|---|---|---|---|---|---|---|---|
| Chaetognaths | 0 | 0 | 0 | 0 | 0 | 0 | 0 | 0 | 0 | 0 | 0 | 0 | 0 | 0 | 0 | 0 | 0 | 0 | 0 | 0 | 0 | 0 | 0 | 0 | 0 | 0 | 0 | 0 | 0 | 0 |
| Euphausiids | 0.2 | 0.2 | 0 | 0 | 0 | 0 | 0 | 0 | 0 | 0 | 0 | 0 | 0 | 0 | 0 | 0 | 0 | 0 | 0 | 0.64 | 0 | 0.05 | 0 | 0 | 0 | 0 | 0 | 0 | 0 | 0 |
| Mysids | 0 | 0 | 0 | 0 | 0 | 0 | 0 | 0 | 0 | 0 | 0 | 0 | 0 | 0 | 0 | 0 | 0 | 0 | 0 | 0 | 0 | 0.05 | 0 | 0 | 0 | 0 | 0 | 0 | 0 | 0 |
| Pelagic Amphipods | 0 | 0 | 0 | 0 | 0 | 0 | 0 | 0 | 0 | 0 | 0 | 0 | 0 | 0 | 0 | 0 | 0 | 0 | 0 | 0 | 0 | 0.05 | 0 | 0 | 0 | 0 | 0 | 0 | 0 | 0 |
| Gelatinous filter feeders | 0 | 0 | 0 | 0 | 0 | 0 | 0 | 0 | 0 | 0 | 0 | 0 | 0 | 0 | 0 | 0 | 0 | 0 | 0 | 0.05 | 0 | 0.05 | 0 | 0 | 0 | 0 | 0 | 0 | 0 | 0 |
| Pteropods | 0 | 0 | 0 | 0 | 0 | 0 | 0 | 0 | 0 | 0 | 0 | 0 | 0 | 0 | 0 | 0 | 0 | 0 | 0 | 0 | 0 | 0.05 | 0 | 0 | 0 | 0 | 0 | 0 | 0 | 0 |
| Copepods | 0 | 0 | 0 | 0 | 0 | 0 | 0 | 0 | 0 | 0 | 0 | 0 | 0 | 0 | 0 | 0 | 0 | 0 | 0 | 0.15 | 0.25 | 0.25 | 0.25 | 0.25 | 0.25 | 0.25 | 0.25 | 0 | 0 | 0 |
| Pelagic microbes | 0 | 0 | 0 | 0 | 0 | 0 | 0 | 0 | 0 | 0 | 0 | 0 | 0 | 0 | 0 | 0 | 0 | 0 | 0 | 0.05 | 0.15 | 0.15 | 0.15 | 0.15 | 0.15 | 0.15 | 0.15 | 0.5 | 0 | 0 |
| Benthic microbes | 0 | 0 | 0 | 0 | 0 | 0 | 0 | 0 | 0.5 | 0.5 | 0.15 | 0.15 | 0.15 | 0.15 | 0.15 | 0.15 | 0.15 | 0.5 | 0.5 | 0 | 0 | 0 | 0 | 0 | 0 | 0 | 0 | 0 | 0 | 0 |
| Macroalgae | 0 | 0 | 0 | 0 | 0.25 | 0.2 | 0 | 0 | 0 | 0 | 0 | 0 | 0 | 0 | 0 | 0 | 0 | 0 | 0 | 0 | 0 | 0 | 0 | 0 | 0 | 0 | 0 | 0 | 0 | 0 |
| Lg Phytoplankton | 0 | 0 | 0 | 0 | 0 | 0 | 0 | 0 | 0 | 0 | 0 | 0 | 0 | 0 | 0 | 0 | 0 | 0 | 0 | 0.05 | 0.5 | 0.25 | 0.5 | 0.5 | 0.5 | 0.5 | 0.5 | 0.25 | 0 | 0 |
| Sm Phytoplankton | 0 | 0 | 0 | 0 | 0 | 0 | 0 | 0 | 0 | 0 | 0 | 0 | 0 | 0 | 0 | 0 | 0 | 0 | 0 | 0 | 0.1 | 0.1 | 0.1 | 0.1 | 0.1 | 0.1 | 0.1 | 0.25 | 0.7 | 0 |
| Outside Production | 0 | 0 | 0 | 0 | 0 | 0 | 0 | 0 | 0 | 0 | 0 | 0 | 0 | 0 | 0 | 0 | 0 | 0 | 0 | 0 | 0 | 0 | 0 | 0 | 0 | 0 | 0 | 0 | 0 | 0 |
| Discards | 0 | 0 | 0 | 0 | 0 | 0 | 0 | 0 | 0 | 0 | 0 | 0 | 0 | 0 | 0 | 0 | 0 | 0 | 0 | 0 | 0 | 0 | 0 | 0 | 0 | 0 | 0 | 0 | 0 | 0 |
| Offal | 0 | 0 | 0 | 0 | 0 | 0 | 0 | 0 | 0 | 0 | 0 | 0 | 0 | 0 | 0 | 0 | 0 | 0 | 0 | 0 | 0 | 0 | 0 | 0 | 0 | 0 | 0 | 0 | 0 | 0 |
| Pelagic Detritus | 0 | 0 | 0 | 0 | 0 | 0 | 0 | 0 | 0 | 0 | 0 | 0 | 0 | 0 | 0 | 0 | 0 | 0 | 0 | 0 | 0 | 0 | 0 | 0 | 0 | 0 | 0 | 0 | 0.3 | 0 |
| Benthic Detritus | 0.4 | 0.4 | 0 | 0.9 | 0.75 | 0.2 | 0.25 | 0.25 | 0.5 | 0.5 | 0.6 | 0.6 | 0.6 | 0.6 | 0.6 | 0.6 | 0.6 | 0.5 | 0.5 | 0 | 0 | 0 | 0 | 0 | 0 | 0 | 0 | 0 | 0 | 1 |
| Outside Detritus | 0 | 0 | 0 | 0 | 0 | 0 | 0 | 0 | 0 | 0 | 0 | 0 | 0 | 0 | 0 | 0 | 0 | 0 | 0 | 0 | 0 | 0 | 0 | 0 | 0 | 0 | 0 | 0 | 0 | 0 |

**Table S4:** Landed catch for the East Bering Sea Ecopath model (converted from the Rpath version of the model; units $tkm^{-2}year^{-1}$)

| Group | Pollock Trawl | Cod Trawl | Cod Pots | Cod Longline | Atka Trawl | RSflats_TWL | YFSflats_TWL | ATFflats_TWL | FHSflats_TWL | Oth_Flatfish_TWL | Turbot Trawl | Turbot Longline | Sablefish Longline | Rockfish Trawl | Halibut Longline | Crab Pots | Salmon Fishery | Herring Fishery | Indigenous | Subsistence |
|---|---|---|---|---|---|---|---|---|---|---|---|---|---|---|---|---|---|---|---|---|
| Transient KIllers | 0 | 0 | 0 | 0 | 0 | 0 | 0 | 0 | 0 | 0 | 0 | 0 | 0 | 0 | 0 | 0 | 0 | 0 | 0 | 0 |
| Sperm and Beaked Whales | 0 | 0 | 0 | 0 | 0 | 0 | 0 | 0 | 0 | 0 | 0 | 0 | 0 | 0 | 0 | 0 | 0 | 0 | 0 | 0 |
| Resident Killers | 0 | 0 | 0 | 0 | 0 | 0 | 0 | 0 | 0 | 0 | 0 | 0 | 0 | 0 | 0 | 0 | 0 | 0 | 0 | 0 |
| Porpoises | 0 | 0 | 0 | 0 | 0 | 0 | 0 | 0 | 0 | 0 | 0 | 0 | 0 | 0 | 0 | 0 | 0 | 0 | 0 | 0 |
| Belugas | 0 | 0 | 0 | 0 | 0 | 0 | 0 | 0 | 0 | 0 | 0 | 0 | 0 | 0 | 0 | 0 | 0 | 0 | 2.52E-05 | 0 |
| Gray Whales | 0 | 0 | 0 | 0 | 0 | 0 | 0 | 0 | 0 | 0 | 0 | 0 | 0 | 0 | 0 | 0 | 0 | 0 | 0 | 0 |
| Humpbacks | 0 | 0 | 0 | 0 | 0 | 0 | 0 | 0 | 0 | 0 | 0 | 0 | 0 | 0 | 0 | 0 | 0 | 0 | 0 | 0 |
| Fin Whales | 0 | 0 | 0 | 0 | 0 | 0 | 0 | 0 | 0 | 0 | 0 | 0 | 0 | 0 | 0 | 0 | 0 | 0 | 0 | 0 |
| Sei Whales | 0 | 0 | 0 | 0 | 0 | 0 | 0 | 0 | 0 | 0 | 0 | 0 | 0 | 0 | 0 | 0 | 0 | 0 | 0 | 0 |
| Right Whales | 0 | 0 | 0 | 0 | 0 | 0 | 0 | 0 | 0 | 0 | 0 | 0 | 0 | 0 | 0 | 0 | 0 | 0 | 0 | 0 |
| Minke Whales | 0 | 0 | 0 | 0 | 0 | 0 | 0 | 0 | 0 | 0 | 0 | 0 | 0 | 0 | 0 | 0 | 0 | 0 | 1.83E-07 | 0 |
| Bowhead Whales | 0 | 0 | 0 | 0 | 0 | 0 | 0 | 0 | 0 | 0 | 0 | 0 | 0 | 0 | 0 | 0 | 0 | 0 | 0 | 0 |
| Sea Otters | 0 | 0 | 0 | 0 | 0 | 0 | 0 | 0 | 0 | 0 | 0 | 0 | 0 | 0 | 0 | 0 | 0 | 0 | 2.97E-08 | 0 |
| Walrus Bd Seals | 0 | 0 | 0 | 0 | 0 | 0 | 0 | 0 | 0 | 0 | 0 | 0 | 0 | 0 | 0 | 0 | 0 | 0 | 3.09E-05 | 0 |
| N Fur Seal Juv | 0 | 0 | 0 | 0 | 0 | 0 | 0 | 0 | 0 | 0 | 0 | 0 | 0 | 0 | 0 | 0 | 0 | 0 | 0 | 0 |
| N Fur Seal Ad | 0 | 0 | 0 | 0 | 0 | 0 | 0 | 0 | 0 | 0 | 0 | 0 | 0 | 0 | 0 | 0 | 0 | 0 | 3.68E-05 | 0 |
| Steller Sea Lions Juv | 0 | 0 | 0 | 0 | 0 | 0 | 0 | 0 | 0 | 0 | 0 | 0 | 0 | 0 | 0 | 0 | 0 | 0 | 0 | 0 |
| Steller Sea Lions Ad | 0 | 0 | 0 | 0 | 0 | 0 | 0 | 0 | 0 | 0 | 0 | 0 | 0 | 0 | 0 | 0 | 0 | 0 | 4.90E-05 | 0 |
| Resident Seals | 0 | 0 | 0 | 0 | 0 | 0 | 0 | 0 | 0 | 0 | 0 | 0 | 0 | 0 | 0 | 0 | 0 | 0 | 1.48E-05 | 0 |
| Wintering Seals | 0 | 0 | 0 | 0 | 0 | 0 | 0 | 0 | 0 | 0 | 0 | 0 | 0 | 0 | 0 | 0 | 0 | 0 | 3.16E-05 | 0 |
| Shearwater | 3.00E-10 | 0 | 0 | 7.00E-10 | 0 | 0 | 0 | 0 | 0 | 0 | 0 | 0 | 0 | 0 | 0 | 0 | 0 | 0 | 0 | 0 |
| Murres | 5.40E-09 | 0 | 0 | 1.43E-08 | 0 | 0 | 0 | 0 | 0 | 0 | 0 | 0 | 0 | 0 | 0 | 0 | 0 | 0 | 0 | 0 |
| Kittiwakes | 4.00E-10 | 0 | 0 | 1.20E-09 | 0 | 0 | 0 | 0 | 0 | 0 | 0 | 0 | 0 | 0 | 0 | 0 | 0 | 0 | 0 | 0 |
| Auklets | 1.20E-09 | 0 | 0 | 3.10E-09 | 0 | 0 | 0 | 0 | 0 | 0 | 0 | 0 | 0 | 0 | 0 | 0 | 0 | 0 | 0 | 0 |
| Puffins | 3.00E-10 | 0 | 0 | 8.00E-10 | 0 | 0 | 0 | 0 | 0 | 0 | 0 | 0 | 0 | 0 | 0 | 0 | 0 | 0 | 0 | 0 |
| Fulmars | 3.00E-10 | 0 | 0 | 9.00E-10 | 0 | 0 | 0 | 0 | 0 | 0 | 0 | 0 | 0 | 0 | 0 | 0 | 0 | 0 | 0 | 0 |
| Storm Petrels | 0 | 0 | 0 | 0 | 0 | 0 | 0 | 0 | 0 | 0 | 0 | 0 | 0 | 0 | 0 | 0 | 0 | 0 | 0 | 0 |
| Cormorants | 1.00E-10 | 0 | 0 | 3.00E-10 | 0 | 0 | 0 | 0 | 0 | 0 | 0 | 0 | 0 | 0 | 0 | 0 | 0 | 0 | 0 | 0 |
| Gulls | 1.00E-10 | 0 | 0 | 2.00E-10 | 0 | 0 | 0 | 0 | 0 | 0 | 0 | 0 | 0 | 0 | 0 | 0 | 0 | 0 | 0 | 0 |
| Albatross Jaeger | 1.00E-10 | 0 | 0 | 2.00E-10 | 0 | 0 | 0 | 0 | 0 | 0 | 0 | 0 | 0 | 0 | 0 | 0 | 0 | 0 | 0 | 0 |
| Sleeper shark | 1.43E-05 | 0 | 0 | 1.11E-07 | 0 | 0 | 0 | 0 | 3.00E-10 | 0 | 0 | 1.06E-06 | 0 | 0 | 0 | 0 | 0 | 0 | 0 | 0 |
| W. Pollock Juv | 0 | 0 | 0 | 0 | 0 | 0 | 0 | 0 | 0 | 0 | 0 | 0 | 0 | 0 | 0 | 0 | 0 | 0 | 0 | 0 |
| W. Pollock Ad | 3.91E-01 | 5.33E-03 | 1.50E-08 | 4.42E-04 | 3.15E-05 | 2.28E-03 | 3.91E-03 | 2.98E-05 | 6.67E-04 | 0 | 1.82E-07 | 2.69E-06 | 0 | 6.38E-06 | 0 | 0 | 0 | 0 | 0 | 0 |

| Group | Pollock Trawl | Cod Trawl | Cod Pots | Cod Longline | Atka Trawl | RSflats_TWL | YFSflats_TWL | ATFflats_TWL | FHSflats_TWL | Oth_Flatfish_TWL | Turbot Trawl | Turbot Longline | Sablefish Longline | Rockfish Trawl | Halibut Longline | Crab Pots | Salmon Fishery | Herring Fishery | Indigenous | Subsistence |
|---|---|---|---|---|---|---|---|---|---|---|---|---|---|---|---|---|---|---|---|---|
| P. Cod Juv | 0 | 0 | 0 | 0 | 0 | 0 | 0 | 0 | 0 | 0 | 0 | 0 | 0 | 0 | 0 | 0 | 0 | 0 | 0 | 0 |
| P. Cod Ad | 2.96E-02 | 5.83E-02 | 9.14E-03 | 1.01E-01 | 1.14E-04 | 7.20E-03 | 1.56E-02 | 2.26E-05 | 2.04E-03 | 0 | 4.54E-05 | 1.08E-04 | 2.94E-05 | 5.96E-04 | 0 | 0 | 0 | 0 | 0 | 0 |
| Herring Juv | 0 | 0 | 0 | 0 | 0 | 0 | 0 | 0 | 0 | 0 | 0 | 0 | 0 | 0 | 0 | 0 | 0 | 0 | 0 | 0 |
| Herring Ad | 0 | 0 | 0 | 0 | 0 | 0 | 0 | 0 | 0 | 0 | 0 | 0 | 0 | 0 | 0 | 0 | 0 | 2.15E-02 | 0 | 0 |
| Arrowtooth Juv | 0 | 0 | 0 | 0 | 0 | 0 | 0 | 0 | 0 | 0 | 0 | 0 | 0 | 0 | 0 | 0 | 0 | 0 | 0 | 0 |
| Arrowtooth Ad | 3.26E-03 | 6.28E-04 | 5.57E-07 | 9.58E-04 | 2.28E-05 | 7.79E-05 | 2.94E-04 | 1.53E-04 | 1.50E-05 | 0 | 1.68E-04 | 1.45E-04 | 8.97E-06 | 1.50E-04 | 0 | 0 | 0 | 0 | 0 | 0 |
| Kamchatka fl. Juv | 0 | 0 | 0 | 0 | 0 | 0 | 0 | 0 | 0 | 0 | 0 | 0 | 0 | 0 | 0 | 0 | 0 | 0 | 0 | 0 |
| Kamchatka fl. Ad | 0 | 0 | 0 | 0 | 0 | 0 | 0 | 0 | 0 | 0 | 0 | 0 | 0 | 0 | 0 | 0 | 0 | 0 | 0 | 0 |
| Gr. Turbot Juv | 0 | 0 | 0 | 0 | 0 | 0 | 0 | 0 | 0 | 0 | 0 | 0 | 0 | 0 | 0 | 0 | 0 | 0 | 0 | 0 |
| Gr. Turbot Ad | 5.05E-05 | 1.47E-05 | 2.09E-06 | 1.46E-04 | 1.06E-05 | 5.93E-06 | 7.83E-05 | 2.67E-06 | 1.51E-05 | 0 | 1.82E-03 | 1.90E-03 | 3.03E-05 | 5.58E-05 | 0 | 0 | 0 | 0 | 0 | 0 |
| P. Halibut Juv | 0 | 0 | 0 | 0 | 0 | 0 | 0 | 0 | 0 | 0 | 0 | 0 | 0 | 0 | 0 | 0 | 0 | 0 | 0 | 0 |
| P. Halibut Ad | 0 | 0 | 0 | 0 | 0 | 0 | 0 | 0 | 0 | 0 | 0 | 0 | 0 | 0 | 8.05E-03 | 0 | 0 | 0 | 0 | 0 |
| YF. Sole Juv | 0 | 0 | 0 | 0 | 0 | 0 | 0 | 0 | 0 | 0 | 0 | 0 | 0 | 0 | 0 | 0 | 0 | 0 | 0 | 0 |
| YF. Sole Ad | 4.29E-04 | 1.07E-03 | 1.27E-08 | 6.06E-07 | 5.20E-08 | 2.54E-03 | 1.30E-01 | 1.18E-06 | 3.83E-03 | 0 | 1.46E-08 | 1.60E-08 | 0 | 1.46E-05 | 0 | 0 | 0 | 0 | 0 | 0 |
| FH. Sole Juv | 0 | 0 | 0 | 0 | 0 | 0 | 0 | 0 | 0 | 0 | 0 | 0 | 0 | 0 | 0 | 0 | 0 | 0 | 0 | 0 |
| FH. Sole Ad | 1.17E-03 | 3.28E-04 | 6.37E-08 | 1.42E-05 | 9.81E-06 | 6.94E-04 | 2.18E-03 | 1.33E-05 | 1.94E-03 | 0 | 2.78E-06 | 3.82E-07 | 2.89E-08 | 7.74E-05 | 0 | 0 | 0 | 0 | 0 | 0 |
| N. Rock sole Juv | 0 | 0 | 0 | 0 | 0 | 0 | 0 | 0 | 0 | 0 | 0 | 0 | 0 | 0 | 0 | 0 | 0 | 0 | 0 | 0 |
| N. Rock sole Ad | 2.62E-03 | 1.64E-03 | 5.22E-08 | 4.49E-06 | 5.18E-06 | 2.79E-02 | 5.97E-03 | 7.31E-06 | 1.61E-03 | 0 | 3.06E-08 | 0 | 0 | 7.50E-06 | 0 | 0 | 0 | 0 | 0 | 0 |
| AK Plaice | 1.24E-03 | 3.47E-04 | 6.73E-08 | 1.50E-05 | 1.04E-05 | 7.33E-04 | 2.30E-03 | 1.41E-05 | 2.05E-03 | 1.54E-06 | 2.94E-06 | 4.03E-07 | 3.05E-08 | 8.18E-05 | 0 | 0 | 0 | 0 | 0 | 0 |
| Dover Sole | 1.38E-08 | 0 | 0 | 0 | 0 | 0 | 0 | 2.41E-07 | 9.14E-08 | 0 | 0 | 0 | 0 | 0 | 0 | 0 | 0 | 0 | 0 | 0 |
| Rex Sole | 8.55E-05 | 1.71E-04 | 0 | 0 | 4.54E-05 | 2.87E-05 | 2.74E-05 | 4.46E-06 | 7.01E-05 | 0 | 5.10E-05 | 0 | 0 | 2.64E-05 | 0 | 0 | 0 | 0 | 0 | 0 |
| Misc. Flatfish | 2.95E-04 | 8.28E-05 | 1.60E-08 | 3.59E-06 | 2.47E-06 | 1.75E-04 | 5.49E-04 | 3.36E-06 | 4.90E-04 | 0 | 7.01E-07 | 9.61E-08 | 7.30E-09 | 1.95E-05 | 0 | 0 | 0 | 0 | 0 | 0 |
| Alaska skate | 2.05E-04 | 5.99E-05 | 2.12E-07 | 3.11E-03 | 0 | 5.00E-05 | 1.87E-04 | 1.94E-06 | 3.57E-05 | 0 | 1.17E-05 | 1.73E-06 | 0 | 3.14E-05 | 0 | 0 | 0 | 0 | 0 | 0 |
| Other skates | 3.32E-05 | 9.69E-06 | 3.43E-08 | 5.03E-04 | 0 | 8.08E-06 | 3.03E-05 | 3.14E-07 | 5.77E-06 | 0 | 1.89E-06 | 2.79E-07 | 0 | 5.07E-06 | 0 | 0 | 0 | 0 | 0 | 0 |
| Sablefish Juv | 0 | 0 | 0 | 0 | 0 | 0 | 0 | 0 | 0 | 0 | 0 | 0 | 0 | 0 | 0 | 0 | 0 | 0 | 0 | 0 |
| Sablefish Ad | 7.32E-06 | 1.46E-05 | 1.45E-05 | 1.93E-04 | 4.79E-05 | 1.93E-05 | 2.70E-06 | 3.08E-06 | 7.51E-06 | 0 | 3.37E-04 | 5.25E-04 | 1.16E-03 | 5.91E-06 | 3.35E-04 | 0 | 0 | 0 | 0 | 0 |
| Eelpouts | 0 | 0 | 0 | 0 | 0 | 0 | 0 | 0 | 0 | 0 | 0 | 0 | 0 | 0 | 0 | 0 | 0 | 0 | 0 | 0 |
| Grenadiers | 3.19E-05 | 0 | 1.91E-08 | 0 | 0 | 0 | 1.79E-07 | 1.56E-08 | 0 | 0 | 1.81E-07 | 4.07E-06 | 0 | 0 | 0 | 0 | 0 | 0 | 0 | 0 |
| Misc. fish deep | 0 | 0 | 0 | 0 | 0 | 0 | 0 | 0 | 0 | 0 | 0 | 0 | 0 | 0 | 0 | 0 | 0 | 0 | 0 | 0 |
| POP | 8.66E-05 | 4.06E-04 | 4.13E-08 | 4.15E-06 | 1.13E-04 | 1.33E-05 | 1.24E-06 | 1.01E-06 | 2.51E-05 | 0 | 5.45E-08 | 2.23E-06 | 4.50E-07 | 7.66E-03 | 0 | 0 | 0 | 0 | 0 | 0 |
| Sharpchin Rock | 1.74E-06 | 6.44E-06 | 1.00E-09 | 1.33E-06 | 1.01E-07 | 1.55E-07 | 2.60E-09 | 3.71E-07 | 4.80E-09 | 0 | 3.50E-09 | 2.77E-07 | 5.88E-07 | 6.18E-05 | 0 | 0 | 0 | 0 | 0 | 0 |
| Northern Rock | 3.30E-05 | 1.22E-04 | 1.85E-08 | 2.53E-05 | 1.91E-06 | 2.93E-06 | 4.88E-08 | 7.05E-06 | 9.13E-08 | 0 | 6.74E-08 | 5.26E-06 | 1.12E-05 | 1.17E-03 | 0 | 0 | 0 | 0 | 0 | 0 |
| Dusky Rock | 2.34E-06 | 2.05E-06 | 1.38E-08 | 2.10E-06 | 0 | 0 | 9.29E-08 | 1.36E-08 | 0 | 0 | 0 | 8.56E-08 | 4.24E-08 | 0 | 0 | 0 | 0 | 0 | 0 | 0 |
| Shortraker Rock | 8.54E-06 | 9.47E-07 | 7.00E-10 | 9.82E-06 | 6.66E-06 | 0 | 1.03E-08 | 1.10E-06 | 1.73E-06 | 0 | 2.34E-05 | 7.72E-06 | 3.84E-06 | 1.80E-04 | 0 | 0 | 0 | 0 | 0 | 0 |
| Rougheye Rock | 5.57E-07 | 1.25E-07 | 0 | 6.09E-06 | 4.57E-06 | 0 | 6.90E-09 | 6.08E-07 | 7.13E-07 | 0 | 1.47E-05 | 3.82E-06 | 2.72E-06 | 1.19E-04 | 0 | 0 | 0 | 0 | 0 | 0 |

| Group | Pollock Trawl | Cod Trawl | Cod Pots | Cod Longline | Atka Trawl | RSflats_TWL | YFSflats_TWL | ATFflats_TWL | FHSflats_TWL | Oth_Flatfish_TWL | Turbot Trawl | Turbot Longline | Sablefish Longline | Rockfish Trawl | Halibut Longline | Crab Pots | Salmon Fishery | Herring Fishery | Indigenous | Subsistence |
|---|---|---|---|---|---|---|---|---|---|---|---|---|---|---|---|---|---|---|---|---|
| Shortspine Thorns | 4.27E-06 | 6.74E-06 | 3.40E-09 | 1.22E-06 | 3.31E-06 | 0 | 2.59E-07 | 1.30E-06 | 3.59E-06 | 0 | 2.43E-05 | 1.23E-05 | 7.04E-05 | 0 | 0 | 0 | 0 | 0 | 0 | 0 |
| Other Sebastes | 1.12E-05 | 7.75E-06 | 9.78E-07 | 5.49E-05 | 9.99E-06 | 7.56E-06 | 2.29E-06 | 6.72E-06 | 3.29E-06 | 0 | 6.63E-05 | 3.21E-05 | 5.06E-05 | 7.84E-05 | 0 | 0 | 0 | 0 | 0 | 0 |
| Atka mackerel Juv | 0 | 0 | 0 | 0 | 0 | 0 | 0 | 0 | 0 | 0 | 0 | 0 | 0 | 0 | 0 | 0 | 0 | 0 | 0 | 0 |
| Atka mackerel Ad | 5.31E-05 | 2.02E-05 | 9.67E-08 | 1.65E-07 | 1.63E-03 | 1.44E-05 | 1.56E-07 | 1.19E-05 | 7.38E-06 | 0 | 1.22E-04 | 0 | 0 | 1.68E-04 | 0 | 0 | 0 | 0 | 0 | 0 |
| Greenlings | 0 | 0 | 0 | 0 | 0 | 0 | 0 | 0 | 0 | 0 | 0 | 0 | 0 | 0 | 0 | 0 | 0 | 0 | 0 | 0 |
| Lg. Sculpins | 8.05E-05 | 1.28E-05 | 2.12E-06 | 9.62E-06 | 0 | 1.68E-05 | 4.46E-04 | 0 | 5.05E-06 | 0 | 1.62E-07 | 6.00E-10 | 0 | 0 | 0 | 0 | 0 | 0 | 0 | 0 |
| Other sculpins | 0 | 0 | 0 | 0 | 0 | 0 | 0 | 0 | 0 | 0 | 0 | 0 | 0 | 0 | 0 | 0 | 0 | 0 | 0 | 0 |
| Misc. fish shallow | 9.52E-06 | 8.93E-07 | 1.04E-08 | 8.31E-08 | 0 | 2.88E-08 | 7.22E-07 | 0 | 0 | 0 | 0 | 0 | 0 | 0 | 0 | 0 | 0 | 0 | 0 | 0 |
| Octopi | 1.45E-06 | 6.18E-06 | 4.27E-05 | 2.97E-06 | 0 | 2.59E-05 | 6.61E-07 | 0 | 0 | 0 | 0 | 6.70E-09 | 0 | 0 | 0 | 0 | 0 | 0 | 0 | 0 |
| Squids | 3.58E-04 | 9.62E-05 | 0 | 0 | 6.40E-09 | 0 | 8.24E-08 | 6.00E-10 | 6.52E-07 | 0 | 3.37E-07 | 0 | 0 | 9.75E-08 | 0 | 0 | 0 | 0 | 0 | 0 |
| Salmon returning | 0 | 0 | 0 | 0 | 0 | 0 | 0 | 0 | 0 | 0 | 0 | 0 | 0 | 0 | 0 | 0 | 1.69E-01 | 0 | 0 | 2.01E-06 |
| Salmon outgoing | 0 | 0 | 0 | 0 | 0 | 0 | 0 | 0 | 0 | 0 | 0 | 0 | 0 | 0 | 0 | 0 | 0 | 0 | 0 | 0 |
| Bathylagidae | 0 | 0 | 0 | 0 | 0 | 0 | 0 | 0 | 0 | 0 | 0 | 0 | 0 | 0 | 0 | 0 | 0 | 0 | 0 | 0 |
| Myctophidae | 2.32E-07 | 0 | 0 | 0 | 0 | 0 | 0 | 0 | 0 | 0 | 0 | 0 | 0 | 0 | 0 | 0 | 0 | 0 | 0 | 0 |
| Capelin | 1.24E-08 | 0 | 0 | 0 | 0 | 0 | 0 | 0 | 0 | 0 | 0 | 0 | 0 | 0 | 0 | 0 | 0 | 0 | 0 | 0 |
| Sandlance | 4.00E-10 | 0 | 0 | 0 | 0 | 0 | 0 | 0 | 0 | 0 | 0 | 0 | 0 | 0 | 0 | 0 | 0 | 0 | 0 | 0 |
| Eulachon | 1.61E-06 | 0 | 0 | 0 | 0 | 0 | 0 | 0 | 0 | 0 | 0 | 0 | 0 | 0 | 0 | 0 | 0 | 0 | 0 | 0 |
| Oth. managed forage | 4.02E-07 | 0 | 0 | 0 | 0 | 0 | 0 | 0 | 0 | 0 | 0 | 0 | 0 | 0 | 0 | 0 | 0 | 0 | 0 | 0 |
| Oth. pelagic smelt | 5.76E-05 | 0 | 0 | 0 | 0 | 1.60E-08 | 9.90E-08 | 0 | 1.70E-09 | 0 | 0 | 0 | 0 | 0 | 0 | 0 | 0 | 0 | 0 | 0 |
| Bairdi | 0 | 0 | 0 | 0 | 0 | 0 | 0 | 0 | 0 | 0 | 0 | 0 | 0 | 0 | 0 | 4.74E-02 | 0 | 0 | 0 | 0 |
| King Crab | 0 | 0 | 0 | 0 | 0 | 0 | 0 | 0 | 0 | 0 | 0 | 0 | 0 | 0 | 0 | 1.88E-02 | 0 | 0 | 0 | 0 |
| Opilio | 0 | 0 | 0 | 0 | 0 | 0 | 0 | 0 | 0 | 0 | 0 | 0 | 0 | 0 | 0 | 3.01E-01 | 0 | 0 | 0 | 0 |
| Pandalidae | 1.62E-07 | 0 | 1.00E-10 | 0 | 0 | 0 | 4.27E-08 | 0 | 1.96E-08 | 0 | 0 | 0 | 0 | 0 | 0 | 0 | 0 | 0 | 0 | 0 |
| NP shrimp | 0 | 0 | 0 | 0 | 0 | 0 | 0 | 0 | 0 | 0 | 0 | 0 | 0 | 0 | 0 | 0 | 0 | 0 | 0 | 0 |
| Sea stars | 4.18E-06 | 3.08E-08 | 2.00E-08 | 2.54E-07 | 0 | 8.16E-06 | 2.56E-05 | 0 | 2.04E-07 | 0 | 2.08E-07 | 2.20E-09 | 0 | 0 | 0 | 0 | 0 | 0 | 0 | 0 |
| Brittle stars | 0 | 0 | 0 | 0 | 0 | 0 | 0 | 0 | 0 | 0 | 0 | 0 | 0 | 0 | 0 | 0 | 0 | 0 | 0 | 0 |
| Urchins dollars cucumbers | 3.94E-08 | 2.25E-08 | 2.00E-10 | 0 | 0 | 4.32E-08 | 2.43E-07 | 0 | 0 | 0 | 0 | 0 | 0 | 0 | 0 | 0 | 0 | 0 | 0 | 0 |
| Snails | 6.70E-09 | 0 | 1.87E-08 | 0 | 0 | 0 | 1.63E-07 | 0 | 0 | 0 | 0 | 0 | 0 | 0 | 0 | 0 | 0 | 0 | 0 | 0 |
| Hermit crabs | 1.46E-07 | 5.47E-08 | 8.46E-08 | 4.80E-09 | 0 | 1.33E-06 | 1.66E-06 | 0 | 1.94E-08 | 0 | 0 | 0 | 0 | 0 | 0 | 0 | 0 | 0 | 0 | 0 |
| Misc. crabs | 0 | 0 | 0 | 0 | 0 | 0 | 0 | 0 | 0 | 0 | 0 | 0 | 0 | 0 | 0 | 3.46E-04 | 0 | 0 | 0 | 0 |
| Misc. Crustacean | 0 | 0 | 0 | 0 | 0 | 0 | 0 | 0 | 0 | 0 | 0 | 0 | 0 | 0 | 0 | 0 | 0 | 0 | 0 | 0 |
| Benthic Amphipods | 0 | 0 | 0 | 0 | 0 | 0 | 0 | 0 | 0 | 0 | 0 | 0 | 0 | 0 | 0 | 0 | 0 | 0 | 0 | 0 |
| Anemones | 1.27E-07 | 0 | 0 | 1.51E-07 | 0 | 1.25E-07 | 2.62E-08 | 0 | 2.00E-09 | 0 | 0 | 0 | 0 | 0 | 0 | 0 | 0 | 0 | 0 | 0 |
| Corals | 0 | 1.93E-08 | 0 | 0 | 0 | 0 | 1.26E-07 | 0 | 0 | 0 | 0 | 0 | 0 | 0 | 0 | 0 | 0 | 0 | 0 | 0 |
| Hydroids | 0 | 0 | 0 | 0 | 0 | 0 | 0 | 0 | 0 | 0 | 0 | 0 | 0 | 0 | 0 | 0 | 0 | 0 | 0 | 0 |

| Group | Pollock Trawl | Cod Trawl | Cod Pots | Cod Longline | Atka Trawl | RSflats_TWL | YFSflats_TWL | ATFflats_TWL | FHSflats_TWL | Oth_Flatfish_TWL | Turbot Trawl | Turbot Longline | Sablefish Longline | Rockfish Trawl | Halibut Longline | Crab Pots | Salmon Fishery | Herring Fishery | Indigenous | Subsistence |
|---|---|---|---|---|---|---|---|---|---|---|---|---|---|---|---|---|---|---|---|---|
| Urochordata | 0 | 0 | 0 | 0 | 0 | 1.96E-06 | 4.02E-06 | 0 | 1.08E-07 | 0 | 0 | 0 | 0 | 0 | 0 | 0 | 0 | 0 | 0 | 0 |
| Sea Pens | 1.80E-07 | 0 | 0 | 0 | 0 | 0 | 0 | 0 | 0 | 0 | 0 | 0 | 0 | 0 | 0 | 0 | 0 | 0 | 0 | 0 |
| Sponges | 3.00E-09 | 1.47E-06 | 1.03E-08 | 0 | 0 | 0 | 4.34E-07 | 0 | 5.00E-10 | 0 | 0 | 0 | 0 | 0 | 0 | 0 | 0 | 0 | 0 | 0 |
| Bivalves | 0 | 0 | 2.33E-08 | 0 | 0 | 0 | 0 | 0 | 0 | 0 | 0 | 0 | 0 | 0 | 0 | 0 | 0 | 0 | 0 | 0 |
| Polychaetes | 0 | 0 | 0 | 0 | 0 | 0 | 0 | 0 | 0 | 0 | 0 | 0 | 0 | 0 | 0 | 0 | 0 | 0 | 0 | 0 |
| Misc. worms | 0 | 0 | 0 | 0 | 0 | 0 | 0 | 0 | 0 | 0 | 0 | 0 | 0 | 0 | 0 | 0 | 0 | 0 | 0 | 0 |
| Scyphozoid Jellies | 3.33E-03 | 7.84E-07 | 2.44E-08 | 0 | 0 | 3.72E-06 | 2.91E-05 | 0 | 1.03E-08 | 0 | 0 | 0 | 0 | 0 | 0 | 0 | 0 | 0 | 0 | 0 |
| Fish Larvae | 0 | 0 | 0 | 0 | 0 | 0 | 0 | 0 | 0 | 0 | 0 | 0 | 0 | 0 | 0 | 0 | 0 | 0 | 0 | 0 |
| Chaetognaths | 0 | 0 | 0 | 0 | 0 | 0 | 0 | 0 | 0 | 0 | 0 | 0 | 0 | 0 | 0 | 0 | 0 | 0 | 0 | 0 |
| Euphausiids | 0 | 0 | 0 | 0 | 0 | 0 | 0 | 0 | 0 | 0 | 0 | 0 | 0 | 0 | 0 | 0 | 0 | 0 | 0 | 0 |
| Mysids | 0 | 0 | 0 | 0 | 0 | 0 | 0 | 0 | 0 | 0 | 0 | 0 | 0 | 0 | 0 | 0 | 0 | 0 | 0 | 0 |
| Pelagic Amphipods | 0 | 0 | 0 | 0 | 0 | 0 | 0 | 0 | 0 | 0 | 0 | 0 | 0 | 0 | 0 | 0 | 0 | 0 | 0 | 0 |
| Gelatinous filter feeders | 0 | 0 | 0 | 0 | 0 | 0 | 0 | 0 | 0 | 0 | 0 | 0 | 0 | 0 | 0 | 0 | 0 | 0 | 0 | 0 |
| Pteropods | 0 | 0 | 0 | 0 | 0 | 0 | 0 | 0 | 0 | 0 | 0 | 0 | 0 | 0 | 0 | 0 | 0 | 0 | 0 | 0 |
| Copepods | 0 | 0 | 0 | 0 | 0 | 0 | 0 | 0 | 0 | 0 | 0 | 0 | 0 | 0 | 0 | 0 | 0 | 0 | 0 | 0 |
| Pelagic microbes | 0 | 0 | 0 | 0 | 0 | 0 | 0 | 0 | 0 | 0 | 0 | 0 | 0 | 0 | 0 | 0 | 0 | 0 | 0 | 0 |
| Benthic microbes | 0 | 0 | 0 | 0 | 0 | 0 | 0 | 0 | 0 | 0 | 0 | 0 | 0 | 0 | 0 | 0 | 0 | 0 | 0 | 0 |
| Macroalgae | 0 | 0 | 0 | 0 | 0 | 0 | 0 | 0 | 0 | 0 | 0 | 0 | 0 | 0 | 0 | 0 | 0 | 0 | 0 | 0 |
| Lg Phytoplankton | 0 | 0 | 0 | 0 | 0 | 0 | 0 | 0 | 0 | 0 | 0 | 0 | 0 | 0 | 0 | 0 | 0 | 0 | 0 | 0 |
| Sm Phytoplankton | 0 | 0 | 0 | 0 | 0 | 0 | 0 | 0 | 0 | 0 | 0 | 0 | 0 | 0 | 0 | 0 | 0 | 0 | 0 | 0 |
| Outside Production | 0 | 0 | 0 | 0 | 0 | 0 | 0 | 0 | 0 | 0 | 0 | 0 | 0 | 0 | 0 | 0 | 0 | 0 | 0 | 0 |
| Discards | 0 | 0 | 0 | 0 | 0 | 0 | 0 | 0 | 0 | 0 | 0 | 0 | 0 | 0 | 0 | 0 | 0 | 0 | 0 | 0 |
| Offal | 0 | 0 | 0 | 0 | 0 | 0 | 0 | 0 | 0 | 0 | 0 | 0 | 0 | 0 | 0 | 0 | 0 | 0 | 0 | 0 |
| Pelagic Detritus | 0 | 0 | 0 | 0 | 0 | 0 | 0 | 0 | 0 | 0 | 0 | 0 | 0 | 0 | 0 | 0 | 0 | 0 | 0 | 0 |
| Benthic Detritus | 0 | 0 | 0 | 0 | 0 | 0 | 0 | 0 | 0 | 0 | 0 | 0 | 0 | 0 | 0 | 0 | 0 | 0 | 0 | 0 |
| Outside Detritus | 0 | 0 | 0 | 0 | 0 | 0 | 0 | 0 | 0 | 0 | 0 | 0 | 0 | 0 | 0 | 0 | 0 | 0 | 0 | 0 |

**Table S5:** Discards for the East Bering Sea Ecopath model (converted from the Rpath version of the model; units tkm$^{-2}$year$^{-1}$)

| Group | Pollock Trawl | Cod Trawl | Cod Pots | Cod Longline | Atka Trawl | RSflats_TWL | YFSflats_TWL | ATFflats_TWL | FHSflats_TWL | Oth_Flatfish_TWL | Turbot Trawl | Turbot Longline | Sablefish Longline | Rockfish Trawl | Halibut Longline | Crab Pots | Salmon Fishery | Herring Fishery | Indigenous | Subsistence |
|---|---|---|---|---|---|---|---|---|---|---|---|---|---|---|---|---|---|---|---|---|
| Transient KIllers | 0 | 0 | 0 | 0 | 0 | 0 | 0 | 0 | 0 | 0 | 0 | 0 | 0 | 0 | 0 | 0 | 0 | 0 | 0 | 0 |
| Sperm and Beaked Whales | 0 | 0 | 0 | 0 | 0 | 0 | 0 | 0 | 0 | 0 | 0 | 0 | 0 | 0 | 0 | 0 | 0 | 0 | 0 | 0 |
| Resident Killers | 2.29E-06 | 5.74E-07 | 0 | 1.24E-06 | 5.16E-08 | 6.27E-07 | 7.84E-07 | 3.92E-08 | 1.18E-07 | 0 | 4.95E-08 | 7.90E-09 | 6.00E-10 | 5.19E-08 | 0 | 0 | 0 | 0 | 0 | 0 |
| Porpoises | 3.49E-07 | 8.73E-08 | 0 | 1.82E-07 | 7.80E-09 | 9.54E-08 | 1.19E-07 | 6.00E-09 | 1.79E-08 | 0 | 7.50E-09 | 1.00E-09 | 1.00E-10 | 7.90E-09 | 0 | 0 | 0 | 0 | 0 | 0 |
| Belugas | 0 | 0 | 0 | 0 | 0 | 0 | 0 | 0 | 0 | 0 | 0 | 0 | 0 | 0 | 0 | 0 | 0 | 0 | 2.52E-08 | 0 |
| Gray Whales | 0 | 0 | 0 | 0 | 0 | 0 | 0 | 0 | 0 | 0 | 0 | 0 | 0 | 0 | 0 | 0 | 0 | 0 | 0 | 0 |
| Humpbacks | 8.35E-06 | 2.09E-06 | 0 | 0 | 1.88E-07 | 2.28E-06 | 2.85E-06 | 1.43E-07 | 4.28E-07 | 0 | 1.80E-07 | 0 | 0 | 1.89E-07 | 0 | 0 | 0 | 0 | 0 | 0 |
| Fin Whales | 0 | 0 | 0 | 0 | 0 | 0 | 0 | 0 | 0 | 0 | 0 | 0 | 0 | 0 | 0 | 0 | 0 | 0 | 0 | 0 |
| Sei Whales | 0 | 0 | 0 | 0 | 0 | 0 | 0 | 0 | 0 | 0 | 0 | 0 | 0 | 0 | 0 | 0 | 0 | 0 | 0 | 0 |
| Right Whales | 0 | 0 | 0 | 0 | 0 | 0 | 0 | 0 | 0 | 0 | 0 | 0 | 0 | 0 | 0 | 0 | 0 | 0 | 0 | 0 |
| Minke Whales | 1.20E-06 | 3.00E-07 | 0 | 0 | 2.70E-08 | 3.28E-07 | 4.11E-07 | 2.05E-08 | 6.16E-08 | 0 | 2.59E-08 | 0 | 0 | 2.72E-08 | 0 | 0 | 0 | 0 | 2.00E-10 | 0 |
| Bowhead Whales | 0 | 0 | 0 | 0 | 0 | 0 | 0 | 0 | 0 | 0 | 0 | 0 | 0 | 0 | 0 | 0 | 0 | 0 | 0 | 0 |
| Sea Otters | 0 | 0 | 0 | 0 | 0 | 0 | 0 | 0 | 0 | 0 | 0 | 0 | 0 | 0 | 0 | 0 | 0 | 0 | 0 | 0 |
| Walrus Bd Seals | 2.96E-06 | 7.41E-07 | 0 | 0 | 6.66E-08 | 8.10E-07 | 1.01E-06 | 5.06E-08 | 1.52E-07 | 0 | 6.39E-08 | 0 | 0 | 6.71E-08 | 0 | 0 | 0 | 0 | 3.09E-08 | 0 |
| N Fur Seal Juv | 0 | 0 | 0 | 0 | 0 | 0 | 0 | 0 | 0 | 0 | 0 | 0 | 0 | 0 | 0 | 0 | 0 | 0 | 0 | 0 |
| N Fur Seal Ad | 6.15E-08 | 1.54E-08 | 0 | 0 | 1.40E-09 | 1.68E-08 | 2.10E-08 | 1.10E-09 | 3.20E-09 | 0 | 1.30E-09 | 0 | 0 | 1.40E-09 | 0 | 0 | 0 | 0 | 3.68E-08 | 0 |
| Steller Sea Lions Juv | 0 | 0 | 0 | 0 | 0 | 0 | 0 | 0 | 0 | 0 | 0 | 0 | 0 | 0 | 0 | 0 | 0 | 0 | 0 | 0 |
| Steller Sea Lions Ad | 7.50E-07 | 1.53E-07 | 0 | 6.40E-08 | 1.25E-08 | 1.80E-07 | 2.18E-07 | 1.10E-08 | 3.30E-08 | 0 | 1.68E-08 | 3.50E-10 | 2.50E-11 | 1.25E-08 | 0 | 0 | 0 | 0 | 1.54E-08 | 0 |
| Resident Seals | 9.33E-08 | 2.33E-08 | 3.29E-08 | 4.36E-08 | 2.10E-09 | 2.55E-08 | 3.19E-08 | 1.60E-09 | 4.80E-09 | 0 | 2.00E-09 | 3.00E-10 | 0 | 2.10E-09 | 0 | 0 | 0 | 0 | 1.48E-08 | 0 |
| Wintering Seals | 1.30E-07 | 3.25E-08 | 0 | 2.34E-08 | 2.90E-09 | 3.55E-08 | 4.44E-08 | 2.20E-09 | 6.70E-09 | 0 | 2.80E-09 | 1.00E-10 | 0 | 2.90E-09 | 0 | 0 | 0 | 0 | 3.16E-08 | 0 |
| Shearwater | 4.71E-08 | 6.00E-09 | 3.00E-09 | 1.45E-06 | 1.50E-09 | 0 | 2.90E-09 | 0 | 0 | 0 | 1.60E-09 | 2.72E-08 | 1.37E-08 | 0 | 0 | 0 | 0 | 0 | 0 | 0 |
| Murres | 9.60E-07 | 1.23E-07 | 6.10E-08 | 2.95E-05 | 3.02E-08 | 0 | 5.81E-08 | 0 | 0 | 0 | 3.17E-08 | 5.53E-07 | 2.79E-07 | 0 | 0 | 0 | 0 | 0 | 0 | 0 |
| Kittiwakes | 7.81E-08 | 1.00E-08 | 5.00E-09 | 2.40E-06 | 2.50E-09 | 0 | 4.70E-09 | 0 | 0 | 0 | 2.60E-09 | 4.50E-08 | 2.27E-08 | 0 | 0 | 0 | 0 | 0 | 0 | 0 |
| Auklets | 2.07E-07 | 2.64E-08 | 1.31E-08 | 6.35E-06 | 6.50E-09 | 0 | 1.25E-08 | 0 | 0 | 0 | 6.80E-09 | 1.19E-07 | 6.01E-08 | 0 | 0 | 0 | 0 | 0 | 0 | 0 |
| Puffins | 5.60E-08 | 7.20E-09 | 3.60E-09 | 1.72E-06 | 1.80E-09 | 0 | 3.40E-09 | 0 | 0 | 0 | 1.90E-09 | 3.23E-08 | 1.63E-08 | 0 | 0 | 0 | 0 | 0 | 0 | 0 |
| Fulmars | 6.13E-08 | 7.80E-09 | 3.90E-09 | 1.88E-06 | 1.90E-09 | 0 | 3.70E-09 | 0 | 0 | 0 | 2.00E-09 | 3.53E-08 | 1.78E-08 | 0 | 0 | 0 | 0 | 0 | 0 | 0 |
| Storm Petrels | 2.00E-10 | 0 | 0 | 6.30E-09 | 0 | 0 | 0 | 0 | 0 | 0 | 0 | 1.00E-10 | 1.00E-10 | 0 | 0 | 0 | 0 | 0 | 0 | 0 |
| Cormorants | 1.77E-08 | 2.30E-09 | 1.10E-09 | 5.42E-07 | 6.00E-10 | 0 | 1.10E-09 | 0 | 0 | 0 | 6.00E-10 | 1.02E-08 | 5.10E-09 | 0 | 0 | 0 | 0 | 0 | 0 | 0 |
| Gulls | 1.24E-08 | 1.60E-09 | 8.00E-10 | 3.80E-07 | 4.00E-10 | 0 | 7.00E-10 | 0 | 0 | 0 | 4.00E-10 | 7.10E-09 | 3.60E-09 | 0 | 0 | 0 | 0 | 0 | 0 | 0 |
| Albatross Jaeger | 1.18E-08 | 1.50E-09 | 7.00E-10 | 3.62E-07 | 4.00E-10 | 0 | 7.00E-10 | 0 | 0 | 0 | 4.00E-10 | 6.80E-09 | 3.40E-09 | 0 | 0 | 0 | 0 | 0 | 0 | 0 |
| Sleeper shark | 2.39E-04 | 3.03E-05 | 0 | 1.81E-04 | 0 | 0 | 1.69E-07 | 5.06E-07 | 4.90E-06 | 0 | 8.68E-06 | 4.81E-05 | 2.53E-05 | 5.71E-05 | 1.01E-04 | 0 | 0 | 0 | 0 | 0 |
| W. Pollock Juv | 0 | 0 | 0 | 0 | 0 | 0 | 0 | 0 | 0 | 0 | 0 | 0 | 0 | 0 | 0 | 0 | 0 | 0 | 0 | 0 |
| W. Pollock Ad | 1.96E+00 | 4.90E-02 | 1.01E-05 | 5.91E-03 | 2.29E-04 | 3.20E-02 | 4.48E-02 | 1.06E-04 | 7.60E-03 | 0 | 1.55E-05 | 9.45E-07 | 4.63E-07 | 7.77E-04 | 0 | 0 | 0 | 0 | 0 | 0 |

| Group | Pollock Trawl | Cod Trawl | Cod Pots | Cod Longline | Atka Trawl | RSflats_TWL | YFSflats_TWL | ATFflats_TWL | FHSflats_TWL | Oth_Flatfish_TWL | Turbot Trawl | Turbot Longline | Sablefish Longline | Rockfish Trawl | Halibut Longline | Crab Pots | Salmon Fishery | Herring Fishery | Indigenous | Subsistence |
|---|---|---|---|---|---|---|---|---|---|---|---|---|---|---|---|---|---|---|---|---|
| P. Cod Juv | 0 | 0 | 0 | 0 | 0 | 0 | 0 | 0 | 0 | 0 | 0 | 0 | 0 | 0 | 0 | 0 | 0 | 0 | 0 | 0 |
| P. Cod Ad | 1.24E-02 | 6.21E-02 | 8.97E-03 | 1.01E-01 | 2.55E-05 | 4.60E-03 | 7.63E-03 | 8.05E-07 | 6.98E-04 | 0 | 5.26E-07 | 5.52E-06 | 3.25E-06 | 9.18E-05 | 9.13E-04 | 0 | 0 | 0 | 0 | 0 |
| Herring Juv | 0 | 0 | 0 | 0 | 0 | 0 | 0 | 0 | 0 | 0 | 0 | 0 | 0 | 0 | 0 | 0 | 0 | 0 | 0 | 0 |
| Herring Ad | 6.36E-03 | 3.11E-06 | 0 | 0 | 0 | 5.02E-05 | 1.13E-03 | 7.81E-08 | 6.37E-05 | 0 | 0 | 0 | 0 | 3.89E-07 | 0 | 0 | 0 | 2.15E-05 | 0 | 0 |
| Arrowtooth Juv | 0 | 0 | 0 | 0 | 0 | 0 | 0 | 0 | 0 | 0 | 0 | 0 | 0 | 0 | 0 | 0 | 0 | 0 | 0 | 0 |
| Arrowtooth Ad | 6.59E-03 | 5.96E-03 | 3.78E-06 | 3.27E-03 | 2.45E-04 | 3.26E-03 | 2.99E-03 | 1.02E-04 | 4.29E-03 | 0 | 1.00E-03 | 3.62E-04 | 2.33E-04 | 1.63E-03 | 2.29E-04 | 0 | 0 | 0 | 0 | 0 |
| Kamchatka fl. Juv | 0 | 0 | 0 | 0 | 0 | 0 | 0 | 0 | 0 | 0 | 0 | 0 | 0 | 0 | 0 | 0 | 0 | 0 | 0 | 0 |
| Kamchatka fl. Ad | 0 | 0 | 0 | 0 | 0 | 0 | 0 | 0 | 0 | 0 | 0 | 0 | 0 | 0 | 0 | 0 | 0 | 0 | 0 | 0 |
| Gr. Turbot Juv | 0 | 0 | 0 | 0 | 0 | 0 | 0 | 0 | 0 | 0 | 0 | 0 | 0 | 0 | 0 | 0 | 0 | 0 | 0 | 0 |
| Gr. Turbot Ad | 2.88E-04 | 8.39E-05 | 6.56E-06 | 4.74E-04 | 4.13E-05 | 3.40E-05 | 3.05E-05 | 2.67E-06 | 2.65E-04 | 0 | 7.67E-04 | 8.31E-04 | 1.95E-03 | 4.00E-05 | 0 | 0 | 0 | 0 | 0 | 0 |
| P. Halibut Juv | 0 | 0 | 0 | 0 | 0 | 0 | 0 | 0 | 0 | 0 | 0 | 0 | 0 | 0 | 0 | 0 | 0 | 0 | 0 | 0 |
| P. Halibut Ad | 2.37E-03 | 3.93E-03 | 6.53E-06 | 8.08E-04 | 1.96E-05 | 2.01E-03 | 1.17E-03 | 8.95E-05 | 9.30E-05 | 0 | 4.60E-04 | 1.30E-06 | 5.39E-05 | 2.08E-04 | 2.00E-04 | 0 | 0 | 0 | 0 | 0 |
| YF. Sole Juv | 0 | 0 | 0 | 0 | 0 | 0 | 0 | 0 | 0 | 0 | 0 | 0 | 0 | 0 | 0 | 0 | 0 | 0 | 0 | 0 |
| YF. Sole Ad | 1.36E-03 | 1.34E-03 | 2.57E-05 | 1.34E-04 | 9.35E-05 | 5.80E-03 | 8.68E-02 | 4.92E-06 | 3.16E-03 | 0 | 3.58E-08 | 1.80E-08 | 5.94E-08 | 1.35E-05 | 0 | 0 | 0 | 0 | 0 | 0 |
| FH. Sole Juv | 0 | 0 | 0 | 0 | 0 | 0 | 0 | 0 | 0 | 0 | 0 | 0 | 0 | 0 | 0 | 0 | 0 | 0 | 0 | 0 |
| FH. Sole Ad | 4.59E-03 | 2.38E-03 | 4.97E-07 | 1.98E-04 | 4.41E-05 | 3.05E-03 | 9.66E-03 | 4.34E-06 | 2.10E-03 | 0 | 9.67E-06 | 2.90E-05 | 6.14E-07 | 7.27E-05 | 0 | 0 | 0 | 0 | 0 | 0 |
| N. Rock sole Juv | 0 | 0 | 0 | 0 | 0 | 0 | 0 | 0 | 0 | 0 | 0 | 0 | 0 | 0 | 0 | 0 | 0 | 0 | 0 | 0 |
| N. Rock sole Ad | 8.75E-03 | 7.97E-03 | 5.35E-07 | 2.99E-05 | 6.29E-05 | 4.13E-02 | 1.31E-02 | 5.19E-06 | 1.51E-03 | 0 | 7.66E-07 | 1.91E-06 | 3.50E-08 | 1.85E-05 | 0 | 0 | 0 | 0 | 0 | 0 |
| AK Plaice | 4.85E-03 | 2.52E-03 | 5.25E-07 | 2.09E-04 | 4.66E-05 | 3.22E-03 | 1.02E-02 | 4.59E-06 | 2.22E-03 | 2.49E-06 | 1.02E-05 | 3.06E-05 | 6.49E-07 | 7.68E-05 | 0 | 0 | 0 | 0 | 0 | 0 |
| Dover Sole | 1.10E-08 | 6.89E-08 | 0 | 7.50E-09 | 9.00E-10 | 6.40E-09 | 6.02E-07 | 7.61E-08 | 6.93E-08 | 0 | 1.48E-08 | 1.11E-08 | 0 | 5.36E-08 | 0 | 0 | 0 | 0 | 0 | 0 |
| Rex Sole | 7.41E-05 | 2.23E-05 | 2.20E-09 | 0 | 9.02E-06 | 1.25E-05 | 9.39E-06 | 1.39E-06 | 2.49E-05 | 0 | 1.88E-05 | 0 | 0 | 0 | 0 | 0 | 0 | 0 | 0 | 0 |
| Misc. Flatfish | 1.16E-03 | 6.00E-04 | 1.25E-07 | 4.98E-05 | 1.11E-05 | 7.69E-04 | 2.44E-03 | 1.09E-06 | 5.30E-04 | 0 | 2.44E-06 | 7.30E-06 | 1.55E-07 | 1.83E-05 | 0 | 0 | 0 | 0 | 0 | 0 |
| Alaska skate | 8.49E-04 | 1.65E-03 | 2.77E-08 | 1.45E-02 | 9.53E-06 | 1.09E-03 | 1.84E-03 | 7.94E-06 | 3.26E-04 | 0 | 4.96E-05 | 9.42E-05 | 1.19E-04 | 7.31E-05 | 6.94E-06 | 0 | 0 | 0 | 0 | 0 |
| Other skates | 1.37E-04 | 2.67E-04 | 4.50E-09 | 2.34E-03 | 1.54E-06 | 1.76E-04 | 2.98E-04 | 1.28E-06 | 5.26E-05 | 0 | 8.01E-06 | 1.52E-05 | 1.92E-05 | 1.18E-05 | 1.18E-04 | 0 | 0 | 0 | 0 | 0 |
| Sablefish Juv | 0 | 0 | 0 | 0 | 0 | 0 | 0 | 0 | 0 | 0 | 0 | 0 | 0 | 0 | 0 | 0 | 0 | 0 | 0 | 0 |
| Sablefish Ad | 3.94E-06 | 5.69E-07 | 3.41E-08 | 2.07E-05 | 9.26E-06 | 3.45E-06 | 1.63E-08 | 1.21E-06 | 1.41E-05 | 0 | 2.45E-05 | 5.18E-06 | 6.38E-06 | 4.96E-06 | 0 | 0 | 0 | 0 | 0 | 0 |
| Eelpouts | 0 | 0 | 0 | 0 | 0 | 0 | 0 | 0 | 0 | 0 | 0 | 0 | 0 | 0 | 0 | 0 | 0 | 0 | 0 | 0 |
| Grenadiers | 9.05E-05 | 6.13E-06 | 6.98E-06 | 7.38E-04 | 1.84E-06 | 0 | 1.47E-06 | 7.28E-07 | 7.80E-06 | 0 | 6.97E-05 | 1.62E-03 | 1.07E-03 | 6.46E-04 | 0 | 0 | 0 | 0 | 0 | 0 |
| Misc. fish deep | 0 | 0 | 0 | 0 | 0 | 0 | 0 | 0 | 0 | 0 | 0 | 0 | 0 | 0 | 0 | 0 | 0 | 0 | 0 | 0 |
| POP | 4.44E-04 | 5.24E-04 | 7.70E-09 | 8.60E-06 | 4.04E-05 | 7.43E-05 | 4.81E-06 | 4.96E-07 | 2.12E-04 | 0 | 2.92E-07 | 2.66E-07 | 7.96E-08 | 6.49E-04 | 3.90E-06 | 0 | 0 | 0 | 0 | 0 |
| Sharpchin Rock | 2.26E-06 | 7.14E-06 | 8.10E-09 | 7.42E-07 | 4.21E-06 | 6.77E-08 | 9.00E-10 | 2.27E-07 | 2.49E-08 | 0 | 2.66E-08 | 2.30E-09 | 3.56E-07 | 7.13E-06 | 3.90E-06 | 0 | 0 | 0 | 0 | 0 |
| Northern Rock | 4.29E-05 | 1.36E-04 | 1.53E-07 | 1.41E-05 | 7.99E-05 | 1.29E-06 | 1.74E-08 | 2.56E-06 | 4.74E-07 | 0 | 5.06E-07 | 4.36E-08 | 6.76E-06 | 1.35E-04 | 3.90E-06 | 0 | 0 | 0 | 0 | 0 |
| Dusky Rock | 3.43E-06 | 3.09E-06 | 2.50E-09 | 5.33E-07 | 1.89E-06 | 1.41E-07 | 0 | 0 | 4.14E-07 | 0 | 0 | 0 | 0 | 0 | 3.90E-06 | 0 | 0 | 0 | 0 | 0 |
| Shortraker Rock | 1.00E-10 | 0 | 0 | 6.00E-10 | 2.00E-10 | 4.00E-10 | 0 | 0 | 1.00E-10 | 0 | 0 | 2.00E-10 | 3.00E-10 | 1.70E-09 | 3.90E-06 | 0 | 0 | 0 | 0 | 0 |
| Rougheye Rock | 1.00E-10 | 0 | 0 | 4.00E-10 | 1.00E-10 | 3.00E-10 | 0 | 0 | 1.00E-10 | 0 | 0 | 1.00E-10 | 2.00E-10 | 1.10E-09 | 3.90E-06 | 0 | 0 | 0 | 0 | 0 |

| Group | Pollock Trawl | Cod Trawl | Cod Pots | Cod Longline | Atka Trawl | RSflats_TWL | YFSflats_TWL | ATFflats_TWL | FHSflats_TWL | Oth_Flatfish_TWL | Turbot Trawl | Turbot Longline | Sablefish Longline | Rockfish Trawl | Halibut Longline | Crab Pots | Salmon Fishery | Herring Fishery | Indigenous | Subsistence |
|---|---|---|---|---|---|---|---|---|---|---|---|---|---|---|---|---|---|---|---|---|
| Shortspine Thorns | 3.55E-06 | 7.09E-07 | 0 | 1.86E-07 | 0 | 0 | 0 | 2.32E-08 | 1.75E-07 | 0 | 2.11E-07 | 3.34E-07 | 3.24E-07 | 0 | 3.90E-06 | 0 | 0 | 0 | 0 | 0 |
| Other Sebastes | 4.02E-05 | 9.01E-05 | 1.53E-06 | 5.90E-05 | 6.91E-05 | 6.29E-06 | 1.77E-06 | 1.18E-06 | 2.53E-06 | 0 | 8.19E-06 | 3.77E-06 | 4.19E-05 | 5.26E-05 | 3.90E-06 | 0 | 0 | 0 | 0 | 0 |
| Atka mackerel Juv | 4.00E-10 | 0 | 0 | 0 | 0 | 0 | 0 | 0 | 0 | 0 | 0 | 0 | 0 | 0 | 0 | 0 | 0 | 0 | 0 | 0 |
| Atka mackerel Ad | 3.18E-04 | 2.92E-03 | 3.76E-04 | 1.51E-03 | 3.18E-05 | 7.77E-04 | 3.10E-03 | 3.71E-06 | 1.93E-04 | 0 | 2.58E-05 | 1.20E-06 | 5.08E-06 | 9.76E-05 | 2.40E-05 | 0 | 0 | 0 | 0 | 0 |
| Greenlings | 0 | 0 | 0 | 0 | 0 | 0 | 0 | 0 | 0 | 0 | 0 | 0 | 0 | 0 | 0 | 0 | 0 | 0 | 0 | 0 |
| Lg. Sculpins | 9.84E-06 | 2.81E-06 | 5.96E-08 | 1.35E-07 | 5.98E-06 | 6.38E-06 | 5.64E-07 | 0 | 2.65E-07 | 0 | 9.75E-06 | 1.59E-08 | 0 | 0 | 0 | 0 | 0 | 0 | 0 | 0 |
| Other sculpins | 2.28E-06 | 7.30E-05 | 6.73E-05 | 3.12E-05 | 1.13E-07 | 5.11E-05 | 2.27E-06 | 6.60E-08 | 6.81E-07 | 0 | 5.05E-07 | 5.94E-08 | 2.90E-08 | 0 | 0 | 0 | 0 | 0 | 0 | 0 |
| Misc. fish shallow | 7.85E-04 | 5.85E-05 | 1.71E-07 | 1.47E-08 | 1.94E-06 | 1.44E-07 | 3.43E-06 | 4.78E-07 | 3.05E-06 | 0 | 8.03E-06 | 6.00E-10 | 4.40E-09 | 1.45E-05 | 0 | 0 | 0 | 0 | 0 | 0 |
| Octopi | 4.20E-04 | 5.39E-05 | 0 | 7.91E-07 | 2.39E-08 | 9.34E-06 | 8.87E-06 | 4.76E-07 | 5.16E-07 | 0 | 1.05E-07 | 0 | 7.90E-09 | 6.17E-06 | 0 | 0 | 1.69E-04 | 0 | 0 | 2.01E-06 |
| Squids | 0 | 0 | 0 | 0 | 0 | 0 | 0 | 0 | 0 | 0 | 0 | 0 | 0 | 0 | 0 | 0 | 0 | 0 | 0 | 0 |
| Salmon returning | 0 | 0 | 0 | 0 | 0 | 0 | 0 | 0 | 0 | 0 | 0 | 0 | 0 | 0 | 0 | 0 | 0 | 0 | 0 | 0 |
| Salmon outgoing | 1.81E-07 | 0 | 1.10E-09 | 0 | 0 | 0 | 9.40E-09 | 0 | 3.00E-10 | 0 | 9.60E-09 | 0 | 0 | 0 | 0 | 0 | 0 | 0 | 0 | 0 |
| Bathylagidae | 4.00E-10 | 0 | 0 | 0 | 0 | 0 | 0 | 0 | 0 | 0 | 0 | 0 | 0 | 0 | 0 | 0 | 0 | 0 | 0 | 0 |
| Myctophidae | 1.40E-09 | 0 | 0 | 0 | 0 | 9.90E-09 | 3.31E-08 | 0 | 0 | 0 | 0 | 0 | 0 | 0 | 0 | 0 | 0 | 0 | 0 | 0 |
| Capelin | 1.93E-06 | 0 | 0 | 0 | 4.60E-09 | 0 | 4.80E-09 | 0 | 0 | 0 | 0 | 0 | 0 | 0 | 0 | 0 | 0 | 0 | 0 | 0 |
| Sandlance | 1.64E-07 | 2.25E-06 | 0 | 1.02E-08 | 0 | 9.17E-07 | 4.37E-07 | 0 | 1.06E-06 | 0 | 0 | 0 | 0 | 0 | 0 | 0 | 0 | 0 | 0 | 0 |
| Eulachon | 4.15E-05 | 1.33E-06 | 0 | 0 | 0 | 2.21E-06 | 5.79E-06 | 2.02E-08 | 3.77E-07 | 0 | 1.57E-08 | 0 | 0 | 0 | 0 | 0 | 0 | 0 | 0 | 0 |
| Oth. managed forage | 5.13E-04 | 3.27E-04 | 1.26E-04 | 4.56E-06 | 2.50E-08 | 4.12E-04 | 3.65E-04 | 1.09E-06 | 1.31E-04 | 0 | 5.66E-06 | 3.00E-10 | 1.70E-09 | 2.39E-06 | 0 | 4.74E-05 | 0 | 0 | 0 | 0 |
| Oth. pelagic smelt | 7.98E-05 | 1.41E-05 | 2.01E-04 | 7.68E-07 | 1.45E-07 | 3.75E-04 | 9.46E-05 | 1.51E-06 | 1.06E-05 | 0 | 2.55E-06 | 9.50E-09 | 4.28E-07 | 3.75E-07 | 0 | 1.88E-05 | 0 | 0 | 0 | 0 |
| Bairdi | 2.62E-03 | 1.55E-04 | 2.34E-05 | 3.54E-05 | 4.46E-07 | 1.00E-03 | 2.02E-03 | 2.64E-06 | 5.28E-04 | 0 | 1.42E-04 | 1.00E-09 | 5.93E-08 | 2.80E-06 | 0 | 3.01E-04 | 0 | 0 | 0 | 0 |
| King Crab | 7.21E-08 | 1.14E-07 | 8.00E-10 | 2.00E-10 | 9.50E-09 | 1.50E-07 | 1.11E-06 | 2.19E-08 | 3.34E-07 | 0 | 1.03E-07 | 0 | 0 | 7.44E-08 | 0 | 0 | 0 | 0 | 0 | 0 |
| Opilio | 0 | 0 | 0 | 0 | 0 | 0 | 0 | 0 | 0 | 0 | 0 | 0 | 0 | 0 | 0 | 0 | 0 | 0 | 0 | 0 |
| Pandalidae | 2.84E-05 | 2.13E-04 | 2.74E-05 | 3.12E-04 | 7.29E-07 | 1.26E-03 | 4.48E-03 | 3.38E-06 | 1.36E-04 | 0 | 1.11E-05 | 3.15E-07 | 8.80E-07 | 1.62E-05 | 1.66E-06 | 0 | 0 | 0 | 0 | 0 |
| NP shrimp | 0 | 0 | 0 | 0 | 0 | 0 | 0 | 0 | 0 | 0 | 0 | 0 | 0 | 0 | 0 | 0 | 0 | 0 | 0 | 0 |
| Sea stars | 1.58E-06 | 2.47E-05 | 7.08E-07 | 1.76E-06 | 1.26E-06 | 2.69E-05 | 7.22E-06 | 1.82E-08 | 6.58E-07 | 0 | 5.04E-07 | 2.78E-08 | 2.43E-07 | 3.11E-08 | 0 | 0 | 0 | 0 | 0 | 0 |
| Brittle stars | 3.82E-07 | 8.40E-07 | 0 | 0 | 0 | 5.71E-07 | 1.62E-05 | 0 | 0 | 0 | 0 | 0 | 0 | 0 | 0 | 0 | 0 | 0 | 0 | 0 |
| Urchins dollars cucumbers | 4.56E-06 | 1.62E-05 | 2.12E-06 | 9.16E-07 | 5.50E-07 | 5.56E-05 | 2.05E-04 | 9.65E-08 | 6.52E-06 | 0 | 1.89E-07 | 1.41E-08 | 4.00E-10 | 2.96E-07 | 0 | 0 | 0 | 0 | 0 | 0 |
| Snails | 0 | 0 | 0 | 0 | 0 | 0 | 0 | 0 | 0 | 0 | 0 | 0 | 0 | 0 | 0 | 3.46E-07 | 0 | 0 | 0 | 0 |
| Hermit crabs | 0 | 0 | 0 | 0 | 0 | 0 | 0 | 0 | 0 | 0 | 0 | 0 | 0 | 0 | 0 | 0 | 0 | 0 | 0 | 0 |
| Misc. crabs | 0 | 0 | 0 | 0 | 0 | 0 | 0 | 0 | 0 | 0 | 0 | 0 | 0 | 0 | 0 | 0 | 0 | 0 | 0 | 0 |
| Misc. Crustacean | 4.53E-06 | 2.71E-05 | 3.54E-08 | 2.22E-04 | 2.99E-07 | 7.00E-05 | 4.96E-05 | 1.77E-07 | 6.69E-06 | 0 | 2.47E-06 | 9.21E-08 | 2.05E-06 | 2.99E-08 | 0 | 0 | 0 | 0 | 0 | 0 |
| Benthic Amphipods | 1.24E-07 | 2.72E-06 | 6.54E-08 | 2.20E-06 | 1.05E-08 | 1.28E-06 | 1.12E-05 | 0 | 2.09E-07 | 0 | 8.84E-08 | 2.84E-08 | 2.45E-07 | 4.30E-07 | 0 | 0 | 0 | 0 | 0 | 0 |
| Anemones | 0 | 0 | 0 | 0 | 0 | 0 | 0 | 0 | 0 | 0 | 0 | 0 | 0 | 0 | 0 | 0 | 0 | 0 | 0 | 0 |
| Corals | 1.67E-06 | 5.14E-05 | 9.40E-09 | 2.23E-06 | 4.01E-08 | 8.58E-05 | 1.33E-04 | 3.70E-09 | 3.10E-05 | 0 | 5.07E-07 | 0 | 2.00E-10 | 8.37E-07 | 0 | 0 | 0 | 0 | 0 | 0 |
| Hydroids | 5.07E-07 | 4.51E-07 | 0 | 5.25E-06 | 7.47E-08 | 2.41E-07 | 0 | 0 | 1.10E-08 | 0 | 7.20E-09 | 9.00E-10 | 0 | 2.44E-07 | 0 | 0 | 0 | 0 | 0 | 0 |

| Group | Pollock Trawl | Cod Trawl | Cod Pots | Cod Longline | Atka Trawl | RSflats_TWL | YFSflats_TWL | ATFflats_TWL | FHSflats_TWL | Oth_Flatfish_TWL | Turbot Trawl | Turbot Longline | Sablefish Longline | Rockfish Trawl | Halibut Longline | Crab Pots | Salmon Fishery | Herring Fishery | Indigenous | Subsistence |
|---|---|---|---|---|---|---|---|---|---|---|---|---|---|---|---|---|---|---|---|---|
| Urochordata | 1.09E-05 | 9.27E-05 | 2.77E-07 | 2.33E-06 | 1.56E-06 | 4.84E-04 | 4.77E-05 | 5.89E-08 | 1.89E-06 | 0 | 2.65E-08 | 1.03E-08 | 5.40E-09 | 5.47E-06 | 0 | 0 | 0 | 0 | 0 | 0 |
| Sea Pens | 0 | 0 | 3.57E-08 | 7.55E-08 | 0 | 0 | 0 | 0 | 0 | 0 | 0 | 0 | 0 | 0 | 0 | 0 | 0 | 0 | 0 | 0 |
| Sponges | 0 | 0 | 0 | 0 | 0 | 0 | 0 | 0 | 0 | 0 | 0 | 0 | 0 | 0 | 0 | 0 | 0 | 0 | 0 | 0 |
| Bivalves | 0 | 0 | 0 | 0 | 0 | 0 | 0 | 0 | 0 | 0 | 0 | 0 | 0 | 0 | 0 | 0 | 0 | 0 | 0 | 0 |
| Polychaetes | 1.24E-02 | 1.17E-03 | 6.83E-06 | 1.83E-05 | 2.57E-05 | 7.65E-04 | 8.32E-04 | 1.06E-06 | 2.52E-05 | 0 | 8.43E-07 | 3.00E-09 | 0 | 7.22E-07 | 0 | 0 | 0 | 0 | 0 | 0 |
| Misc. worms | 0 | 0 | 0 | 0 | 0 | 0 | 0 | 0 | 0 | 0 | 0 | 0 | 0 | 0 | 0 | 0 | 0 | 0 | 0 | 0 |
| Scyphozoid Jellies | 0 | 0 | 0 | 0 | 0 | 0 | 0 | 0 | 0 | 0 | 0 | 0 | 0 | 0 | 0 | 0 | 0 | 0 | 0 | 0 |
| Fish Larvae | 0 | 0 | 0 | 0 | 0 | 0 | 0 | 0 | 0 | 0 | 0 | 0 | 0 | 0 | 0 | 0 | 0 | 0 | 0 | 0 |
| Chaetognaths | 0 | 0 | 0 | 0 | 0 | 0 | 0 | 0 | 0 | 0 | 0 | 0 | 0 | 0 | 0 | 0 | 0 | 0 | 0 | 0 |
| Euphausiids | 0 | 0 | 0 | 0 | 0 | 0 | 0 | 0 | 0 | 0 | 0 | 0 | 0 | 0 | 0 | 0 | 0 | 0 | 0 | 0 |
| Mysids | 0 | 0 | 0 | 0 | 0 | 0 | 0 | 0 | 0 | 0 | 0 | 0 | 0 | 0 | 0 | 0 | 0 | 0 | 0 | 0 |
| Pelagic Amphipods | 0 | 0 | 0 | 0 | 0 | 0 | 0 | 0 | 0 | 0 | 0 | 0 | 0 | 0 | 0 | 0 | 0 | 0 | 0 | 0 |
| Gelatinous filter feeders | 0 | 0 | 0 | 0 | 0 | 0 | 0 | 0 | 0 | 0 | 0 | 0 | 0 | 0 | 0 | 0 | 0 | 0 | 0 | 0 |
| Pteropods | 0 | 0 | 0 | 0 | 0 | 0 | 0 | 0 | 0 | 0 | 0 | 0 | 0 | 0 | 0 | 0 | 0 | 0 | 0 | 0 |
| Copepods | 0 | 0 | 0 | 0 | 0 | 0 | 0 | 0 | 0 | 0 | 0 | 0 | 0 | 0 | 0 | 0 | 0 | 0 | 0 | 0 |
| Pelagic microbes | 0 | 0 | 0 | 0 | 0 | 0 | 0 | 0 | 0 | 0 | 0 | 0 | 0 | 0 | 0 | 0 | 0 | 0 | 0 | 0 |
| Benthic microbes | 0 | 0 | 0 | 0 | 0 | 0 | 0 | 0 | 0 | 0 | 0 | 0 | 0 | 0 | 0 | 0 | 0 | 0 | 0 | 0 |
| Macroalgae | 0 | 0 | 0 | 0 | 0 | 0 | 0 | 0 | 0 | 0 | 0 | 0 | 0 | 0 | 0 | 0 | 0 | 0 | 0 | 0 |
| Lg Phytoplankton | 0 | 0 | 0 | 0 | 0 | 0 | 0 | 0 | 0 | 0 | 0 | 0 | 0 | 0 | 0 | 0 | 0 | 0 | 0 | 0 |
| Sm Phytoplankton | 0 | 0 | 0 | 0 | 0 | 0 | 0 | 0 | 0 | 0 | 0 | 0 | 0 | 0 | 0 | 0 | 0 | 0 | 0 | 0 |
| Outside Production | 0 | 0 | 0 | 0 | 0 | 0 | 0 | 0 | 0 | 0 | 0 | 0 | 0 | 0 | 0 | 0 | 0 | 0 | 0 | 0 |
| Discards | 0 | 0 | 0 | 0 | 0 | 0 | 0 | 0 | 0 | 0 | 0 | 0 | 0 | 0 | 0 | 0 | 0 | 0 | 0 | 0 |
| Offal | 0 | 0 | 0 | 0 | 0 | 0 | 0 | 0 | 0 | 0 | 0 | 0 | 0 | 0 | 0 | 0 | 0 | 0 | 0 | 0 |
| Pelagic Detritus | 0 | 0 | 0 | 0 | 0 | 0 | 0 | 0 | 0 | 0 | 0 | 0 | 0 | 0 | 0 | 0 | 0 | 0 | 0 | 0 |
| Benthic Detritus | 0 | 0 | 0 | 0 | 0 | 0 | 0 | 0 | 0 | 0 | 0 | 0 | 0 | 0 | 0 | 0 | 0 | 0 | 0 | 0 |
| Outside Detritus | 4E-10 | 0 | 0 | 0 | 0 | 0 | 0 | 0 | 0 | 0 | 0 | 0 | 0 | 0 | 0 | 0 | 0 | 0 | 0 | 0 |

**Table S6:** Ecopath base parameters for the SE Australian Ecopath model

| Group | Trophic Level | Biomass (tkm$^{-2}$) | Production / Biomass (year$^{-1}$) | Consumption / Biomass (year$^{-1}$) | Ecotrophic Efficiency | Biomass Accumulation (tkm$^{-2}$) | Biomass Accumulation year$^{-1}$) |
|---|---|---|---|---|---|---|---|
| Toothed whale | 4.563 | 0.013 | 0.02 | 13 | 0.634 | 0 | 0 |
| Baleen whale | 3.754 | 0.006 | 0.02 | 11.2 | 0 | 0 | 0 |
| Seal | 4.622 | 0.051 | 0.18 | 38.898 | 0.724 | 0.004 | 0.105 |
| Seabirds | 4.481 | 0.003 | 0.8 | 80 | 0.363 | 0 | 0 |
| Penguins | 4.389 | 0.001 | 0.8 | 80 | 0.692 | 0 | 0 |
| Tuna/billfish | 4.579 | 1.131 | 0.48 | 6.8 | 0.4 | 0 | 0 |
| Pelagic sharks | 4.916 | 0.004 | 0.2 | 1.2 | 0.95 | 0 | 0 |
| Demersal sharks | 4.466 | 1.215 | 0.18 | 1.8 | 0.806 | 0 | 0 |
| Rays | 3.604 | 1.2 | 0.35 | 3.5 | 0.661 | 0 | 0 |
| Warehous | 3.962 | 0.9 | 0.28 | 2.4 | 0.787 | 0 | 0 |
| Redbait | 3.506 | 2.2 | 0.49 | 4.1 | 0.974 | 0 | 0 |
| Redfish | 3.948 | 1.07 | 0.31 | 3.4 | 0.646 | 0 | 0 |
| Ling | 4.457 | 0.44 | 0.22 | 2.4 | 0.767 | 0 | 0 |
| Dories | 4.579 | 0.39 | 0.3 | 2.8 | 0.826 | 0 | 0 |
| Jack mackerel | 3.524 | 6 | 0.47 | 3.3 | 0.647 | 0 | 0 |
| Jackass morwong | 3.399 | 0.628 | 0.22 | 2.9 | 0.975 | 0 | 0 |
| Flathead | 4.305 | 0.434 | 0.52 | 3.5 | 0.803 | 0 | 0 |
| Gemfish | 4.791 | 0.22 | 0.44 | 2.1 | 0.893 | 0 | 0 |
| ShOceanPerch | 4.219 | 0.274 | 0.26 | 2.6 | 0.995 | 0 | 0 |
| Chinaman leatherjacket | 4.21 | 0.011 | 0.36 | 2.3 | 0.565 | 0 | 0 |
| Cucumberfish | 3.451 | 2.49 | 0.52 | 4.7 | 0.987 | 0 | 0 |
| Whiting | 3.358 | 1.691 | 0.9 | 5.4 | 0.799 | 0 | 0 |
| Cardinal | 4.089 | 4.088 | 0.77 | 6.4 | 0.95 | 0 | 0 |
| ShSmInvertFeeder | 3.412 | 5.21 | 0.65 | 4.67 | 1 | 0 | 0 |
| ShSmPredator | 3.989 | 0.8 | 0.55 | 4.46 | 0.999 | 0 | 0 |
| ShMedInvertFeeder | 3.941 | 1.35 | 0.36 | 3.4 | 0.957 | 0 | 0 |
| ShMedPredator | 4.363 | 0.51 | 0.4 | 2.93 | 0.996 | 0 | 0 |
| ShLInvertFeeder | 3.584 | 0.12 | 0.21 | 2 | 0.437 | 0 | 0 |
| ShLPredator | 4.385 | 3 | 0.22 | 1.84 | 0.51 | 0 | 0 |
| Blue-eye trevalla | 4.177 | 0.35 | 0.2 | 1.4 | 0.557 | 0 | 0 |
| Blue grenadier | 4.39 | 0.82 | 0.27 | 2.9 | 0.973 | 0 | 0 |
| SlopeOceanPerch | 4.23 | 0.18 | 0.26 | 3.1 | 0.935 | 0 | 0 |
| Deepsea Cod | 3.775 | 0.47 | 0.25 | 2.2 | 0.294 | 0 | 0 |
| Oreos | 3.639 | 0.084 | 0.35 | 2.7 | 0.655 | 0 | 0 |
| SlopeSmInvertFeeder | 3.474 | 0.2 | 0.47 | 4.13 | 0.985 | 0 | 0 |
| SlopeSmPredator | 3.888 | 0.453 | 0.4 | 3.24 | 0.743 | 0 | 0 |
| SlopeMInverFeeder | 3.594 | 3.3 | 0.29 | 3.5 | 0.616 | 0 | 0 |
| SlopeMPredator | 4.712 | 0.3 | 0.305 | 2.5 | 0.961 | 0 | 0 |
| SlopeLInvertFeeder | 3.932 | 1.229 | 0.44 | 2.9 | 0.95 | 0 | 0 |
| SlopeLPredator | 4.636 | 0.12 | 0.2 | 2.34 | 0.512 | 0 | 0 |
| PelSmInvertFeeder | 3.061 | 6.193 | 0.76 | 8.85 | 0.98 | 0 | 0 |
| PelMInvertFeeder | 3.345 | 0.131 | 0.46 | 3.4 | 0.88 | 0 | 0 |
| PelMPredator | 4.162 | 0.32 | 0.32 | 2.85 | 0.974 | 0 | 0 |
| PelLInvertFeeder | 3.396 | 0.039 | 0.12 | 2.6 | 0.935 | 0 | 0 |
| PelLPredator | 4.19 | 0.009 | 0.26 | 3.1 | 0.95 | 0 | 0 |
| Mesopelagics | 3.248 | 200 | 0.83 | 8 | 0.871 | 0 | 0 |
| Squid | 4.046 | 1.64 | 2.75 | 10 | 0.996 | 0 | 0 |
| PelagicPrawns | 2.2 | 3.024 | 1.6 | 10 | 0.95 | 0 | 0 |
| Macrobenthos | 2.52 | 26.319 | 1.6 | 6 | 0.95 | 0 | 0 |
| Megabenthos | 3.216 | 7.203 | 2.5 | 5.85 | 0.95 | 0 | 0 |

| Group | Trophic Level | Biomass (tkm$^{-2}$) | Production / Biomass (year$^{-1}$) | Consumption / Biomass (year$^{-1}$) | Ecotrophic Efficiency | Biomass Accumulation (tkm$^{-2}$) | Biomass Accumulation year$^{-1}$) |
|---|---|---|---|---|---|---|---|
| Polychaeta | 2.126 | 5.749 | 2 | 22 | 0.95 | 0 | 0 |
| Gelatinous nekton | 2.9 | 2.575 | 3 | 10 | 0.95 | 0 | 0 |
| Euphausids | 2.2 | 122.163 | 5 | 32 | 0.95 | 0 | 0 |
| L zooplankton | 2.2 | 10.938 | 5 | 32 | 0.95 | 0 | 0 |
| Sm zooplankton | 2 | 32.601 | 20 | 70 | 0.95 | 0 | 0 |
| Primary producers | 1 | 15 | 298 | | 0.634 | 0 | 0 |
| Benthic producer | 1 | 0.006 | 4.43 | | 0 | 0 | 0 |
| Detritus | 1 | 200 | | | 0.724 | 0.004 | 0.105 |
| Discards | 1 | 0.17 | | | 0.363 | 0 | 0 |

**Table S7:** Diet matrix for the SE Australian Ecopath model

| Group | Toothed whale | Baleen whale | Seal | Seabirds | Penguins | Tuna/billfish | Pelagic sharks | Demersal sharks | Rays | Warehous | Redbait | Redfish | Ling | Dories | Jack mackerel | Jackass morwong | Flathead | Gemfish | ShOceanPerch | Chinaman leatherjacket | Cucumberfish | Whiting | Cardinal | ShSmInvertFeeder | ShSmPredator | ShMedInvertFeeder | ShMedPredator | ShLInvertFeeder | ShLPredator | Blue-eye trevalla |
|---|---|---|---|---|---|---|---|---|---|---|---|---|---|---|---|---|---|---|---|---|---|---|---|---|---|---|---|---|---|---|
| Toothed whale | 0.001 | 0 | 0 | 0 | 0 | 0 | 0 | 0 | 0 | 0 | 0 | 0 | 0 | 0 | 0 | 0 | 0 | 0 | 0 | 0 | 0 | 0 | 0 | 0 | 0 | 0 | 0 | 0 | 0 | 0 |
| Baleen whale | 0 | 0 | 0 | 0 | 0 | 0 | 0 | 0 | 0 | 0 | 0 | 0 | 0 | 0 | 0 | 0 | 0 | 0 | 0 | 0 | 0 | 0 | 0 | 0 | 0 | 0 | 0 | 0 | 0 | 0 |
| Seal | 0.002 | 0 | 0 | 0 | 0 | 0 | 0.025 | 0 | 0 | 0 | 0 | 0 | 0 | 0 | 0 | 0 | 0 | 0 | 0 | 0 | 0 | 0 | 0 | 0 | 0 | 0 | 0 | 0 | 0 | 0 |
| Seabirds | 0.005 | 0 | 0 | 0 | 0 | 0 | 0.009 | 0 | 0 | 0 | 0 | 0 | 0 | 0 | 0 | 0 | 0 | 0 | 0 | 0 | 0 | 0 | 0 | 0 | 0 | 0 | 0 | 0 | 0 | 0 |
| Penguins | 0.003 | 0 | 0 | 0 | 0 | 0 | 0.009 | 0 | 0 | 0 | 0 | 0 | 0 | 0 | 0 | 0 | 0 | 0 | 0 | 0 | 0 | 0 | 0 | 0 | 0 | 0 | 0 | 0 | 0 | 0 |
| Tuna/billfish | 0.002 | 0 | 0 | 0 | 0 | 0 | 0.025 | 0 | 0 | 0 | 0 | 0 | 0 | 0 | 0 | 0 | 0 | 0 | 0 | 0 | 0 | 0 | 0 | 0 | 0 | 0 | 0 | 0 | 0 | 0 |
| Pelagic sharks | 0.002 | 0 | 0 | 0 | 0 | 0 | 0.006 | 0 | 0 | 0 | 0 | 0 | 0 | 0 | 0 | 0 | 0 | 0 | 0 | 0 | 0 | 0 | 0 | 0 | 0 | 0 | 0 | 0 | 0 | 0 |
| Demersal sharks | 0 | 0 | 0 | 0 | 0 | 0 | 0.134 | 0 | 0 | 0 | 0 | 0 | 0 | 0 | 0 | 0 | 0 | 0 | 0 | 0 | 0 | 0 | 0 | 0 | 0 | 0 | 0.01 | 0 | 0.03 | 0 |
| Rays | 0 | 0 | 0 | 0 | 0 | 0 | 0.134 | 0.006 | 0 | 0 | 0 | 0 | 0 | 0 | 0 | 0 | 0 | 0 | 0 | 0 | 0 | 0 | 0 | 0 | 0 | 0 | 0.01 | 0 | 0.05 | 0 |
| Warehous | 0 | 0 | 0.045 | 0 | 0.1 | 0 | 0.048 | 0 | 0 | 0 | 0 | 0 | 0 | 0 | 0 | 0 | 0 | 0 | 0 | 0 | 0 | 0 | 0 | 0 | 0 | 0 | 0 | 0 | 0.01 | 0 |
| Redbait | 0 | 0.01 | 0.251 | 0.01 | 0 | 0.159 | 0 | 0.008 | 0 | 0 | 0 | 0 | 0 | 0.096 | 0 | 0 | 0 | 0 | 0 | 0 | 0 | 0 | 0 | 0 | 0 | 0 | 0.05 | 0 | 0.017 | 0 |
| Redfish | 0 | 0 | 0.024 | 0 | 0 | 0 | 0.009 | 0.023 | 0 | 0 | 0 | 0 | 0 | 0.039 | 0 | 0 | 0 | 0 | 0 | 0 | 0 | 0 | 0 | 0 | 0 | 0 | 0 | 0 | 0.002 | 0 |
| Ling | 0 | 0 | 0.012 | 0 | 0 | 0 | 0.001 | 0.001 | 0 | 0 | 0 | 0 | 0 | 0 | 0 | 0 | 0.014 | 0 | 0 | 0 | 0 | 0 | 0 | 0 | 0 | 0 | 0 | 0 | 0 | 0 |
| Dories | 0 | 0 | 0.01 | 0 | 0 | 0 | 0.008 | 0.001 | 0 | 0 | 0 | 0 | 0 | 0 | 0 | 0 | 0.01 | 0.05 | 0 | 0 | 0 | 0 | 0 | 0 | 0 | 0 | 0.03 | 0 | 0 | 0 |
| Jack mackerel | 0.078 | 0 | 0.3 | 0 | 0 | 0.228 | 0.045 | 0.21 | 0 | 0 | 0 | 0 | 0 | 0.245 | 0 | 0 | 0.005 | 0.249 | 0 | 0 | 0 | 0 | 0 | 0 | 0 | 0 | 0.06 | 0 | 0.032 | 0 |
| Jackass morwong | 0 | 0 | 0.016 | 0 | 0 | 0 | 0 | 0.011 | 0 | 0 | 0 | 0 | 0 | 0 | 0 | 0 | 0 | 0 | 0 | 0 | 0 | 0 | 0 | 0 | 0 | 0 | 0.02 | 0 | 0.009 | 0 |
| Flathead | 0 | 0 | 0.035 | 0 | 0 | 0 | 0 | 0.007 | 0 | 0 | 0 | 0 | 0 | 0 | 0 | 0.001 | 0 | 0 | 0 | 0 | 0 | 0 | 0 | 0 | 0 | 0 | 0.01 | 0 | 0 | 0 |
| Gemfish | 0 | 0 | 0.015 | 0 | 0 | 0 | 0.008 | 0 | 0 | 0 | 0 | 0 | 0 | 0 | 0 | 0 | 0 | 0 | 0 | 0 | 0 | 0 | 0 | 0 | 0 | 0 | 0 | 0 | 0 | 0 |
| ShOceanPerch | 0 | 0 | 0.005 | 0 | 0 | 0 | 0 | 0.001 | 0 | 0 | 0 | 0 | 0 | 0 | 0 | 0 | 0.002 | 0 | 0 | 0 | 0 | 0 | 0 | 0 | 0 | 0 | 0 | 0 | 0.002 | 0 |
| Chinaman leatherjacket | 0 | 0 | 0 | 0 | 0 | 0 | 0 | 0 | 0 | 0 | 0 | 0 | 0 | 0 | 0 | 0 | 0 | 0 | 0 | 0 | 0 | 0 | 0 | 0 | 0 | 0 | 0 | 0 | 0 | 0 |
| Cucumberfish | 0 | 0 | 0 | 0 | 0 | 0 | 0 | 0.007 | 0 | 0 | 0 | 0.01 | 0.08 | 0.152 | 0 | 0.002 | 0.128 | 0.05 | 0.08 | 0 | 0 | 0 | 0 | 0 | 0.1 | 0.04 | 0.04 | 0 | 0.04 | 0 |
| Whiting | 0 | 0 | 0.01 | 0 | 0 | 0 | 0 | 0.003 | 0.02 | 0 | 0 | 0 | 0 | 0.006 | 0 | 0 | 0.1 | 0 | 0 | 0 | 0 | 0 | 0 | 0 | 0.1 | 0 | 0.002 | 0 | 0.083 | 0 |
| Cardinal | 0 | 0 | 0.008 | 0 | 0 | 0 | 0 | 0.037 | 0 | 0.018 | 0 | 0.253 | 0.192 | 0.16 | 0.006 | 0 | 0.05 | 0.071 | 0.074 | 0 | 0 | 0 | 0 | 0 | 0.101 | 0 | 0.025 | 0 | 0.181 | 0 |
| ShSmInvertFeeder | 0 | 0 | 0.092 | 0 | 0 | 0 | 0 | 0.03 | 0.022 | 0 | 0 | 0.01 | 0.477 | 0.224 | 0.001 | 0.007 | 0.085 | 0 | 0.083 | 0.1 | 0.01 | 0.005 | 0 | 0.002 | 0.151 | 0.064 | 0.251 | 0 | 0.014 | 0 |
| ShSmPredator | 0 | 0 | 0.051 | 0 | 0 | 0 | 0 | 0 | 0 | 0 | 0 | 0.01 | 0 | 0 | 0 | 0 | 0 | 0 | 0 | 0.1 | 0.005 | 0 | 0 | 0.001 | 0 | 0 | 0.012 | 0 | 0 | 0 |
| ShMedInvertFeeder | 0 | 0 | 0 | 0.005 | 0 | 0 | 0 | 0.013 | 0 | 0 | 0 | 0.001 | 0 | 0.019 | 0 | 0.004 | 0.025 | 0 | 0.082 | 0 | 0.001 | 0.005 | 0 | 0.001 | 0.005 | 0.001 | 0.025 | 0 | 0.01 | 0 |
| ShMedPredator | 0 | 0 | 0.04 | 0.005 | 0 | 0 | 0 | 0.005 | 0 | 0 | 0 | 0 | 0 | 0.017 | 0 | 0.002 | 0.005 | 0 | 0 | 0 | 0 | 0 | 0 | 0 | 0 | 0 | 0 | 0 | 0 | 0 |
| ShLInvertFeeder | 0 | 0 | 0 | 0 | 0.1 | 0 | 0 | 0 | 0 | 0 | 0 | 0 | 0 | 0 | 0 | 0 | 0 | 0 | 0 | 0 | 0 | 0 | 0 | 0 | 0 | 0 | 0 | 0 | 0 | 0 |
| ShLPredator | 0 | 0 | 0 | 0 | 0 | 0 | 0 | 0.008 | 0 | 0 | 0 | 0 | 0 | 0 | 0 | 0 | 0.01 | 0.169 | 0.007 | 0 | 0 | 0 | 0 | 0 | 0 | 0 | 0.02 | 0 | 0 | 0 |
| Blue-eye trevalla | 0.01 | 0 | 0 | 0 | 0 | 0 | 0 | 0.001 | 0 | 0 | 0 | 0 | 0 | 0 | 0 | 0 | 0 | 0 | 0 | 0 | 0 | 0 | 0 | 0 | 0 | 0 | 0 | 0 | 0 | 0 |
| Blue grenadier | 0 | 0 | 0 | 0 | 0 | 0 | 0 | 0 | 0 | 0 | 0 | 0 | 0 | 0 | 0 | 0 | 0 | 0.06 | 0 | 0 | 0 | 0 | 0 | 0 | 0 | 0 | 0 | 0 | 0 | 0.01 |
| SlopeOceanPerch | 0 | 0 | 0 | 0 | 0 | 0 | 0 | 0 | 0 | 0 | 0 | 0.001 | 0 | 0 | 0 | 0 | 0 | 0 | 0 | 0 | 0 | 0 | 0 | 0 | 0 | 0 | 0 | 0 | 0.002 | 0 |
| Deepsea Cod | 0 | 0 | 0 | 0 | 0 | 0 | 0 | 0 | 0 | 0 | 0 | 0 | 0 | 0 | 0 | 0 | 0 | 0 | 0 | 0 | 0 | 0 | 0 | 0 | 0 | 0 | 0 | 0 | 0 | 0 |
| Oreos | 0 | 0 | 0 | 0 | 0 | 0 | 0 | 0 | 0 | 0 | 0 | 0 | 0 | 0 | 0 | 0 | 0 | 0 | 0 | 0 | 0 | 0 | 0 | 0 | 0 | 0 | 0 | 0 | 0 | 0 |
| SlopeSmInvertFeeder | 0 | 0.02 | 0 | 0 | 0 | 0 | 0 | 0 | 0.001 | 0 | 0 | 0 | 0 | 0 | 0 | 0 | 0 | 0.016 | 0 | 0.1 | 0 | 0 | 0 | 0 | 0 | 0 | 0 | 0 | 0 | 0.01 |

| Group | Toothed whale | Baleen whale | Seal | Seabirds | Penguins | Tuna/billfish | Pelagic sharks | Demersal sharks | Rays | Warehous | Redbait | Redfish | Ling | Dories | Jack mackerel | Jackass morwong | Flathead | Gemfish | ShOceanPerch | Chinaman leatherjacket | Cucumberfish | Whiting | Cardinal | ShSmInvertFeeder | ShSmPredator | ShMedInvertFeeder | ShMedPredator | ShLInvertFeeder | ShLPredator | Blue-eye trevalla |
|---|---|---|---|---|---|---|---|---|---|---|---|---|---|---|---|---|---|---|---|---|---|---|---|---|---|---|---|---|---|---|
| SlopeSmPredator | 0 | 0.01 | 0 | 0 | 0 | 0 | 0 | 0.006 | 0.004 | 0 | 0 | 0.001 | 0.01 | 0 | 0 | 0 | 0 | 0 | 0.014 | 0.1 | 0.001 | 0 | 0 | 0 | 0 | 0 | 0 | 0 | 0 | 0.01 |
| SlopeMInverFeeder | 0 | 0 | 0 | 0 | 0 | 0 | 0 | 0.022 | 0.01 | 0 | 0 | 0.004 | 0 | 0 | 0 | 0.002 | 0 | 0.111 | 0 | 0 | 0 | 0 | 0 | 0 | 0 | 0 | 0 | 0 | 0 | 0.01 |
| SlopeMPredator | 0 | 0 | 0 | 0 | 0.1 | 0 | 0 | 0.006 | 0 | 0 | 0 | 0 | 0 | 0 | 0 | 0 | 0 | 0 | 0 | 0 | 0 | 0 | 0 | 0 | 0 | 0 | 0 | 0 | 0 | 0 |
| SlopeLInvertFeeder | 0 | 0 | 0 | 0 | 0 | 0 | 0 | 0.005 | 0 | 0 | 0 | 0.014 | 0.056 | 0 | 0 | 0 | 0.105 | 0 | 0.068 | 0 | 0 | 0 | 0 | 0 | 0.005 | 0 | 0 | 0 | 0 | 0 |
| SlopeLPredator | 0 | 0 | 0 | 0 | 0 | 0 | 0 | 0 | 0 | 0 | 0 | 0 | 0 | 0 | 0 | 0 | 0 | 0 | 0 | 0 | 0 | 0 | 0 | 0 | 0 | 0 | 0 | 0 | 0 | 0 |
| PelSmInvertFeeder | 0.098 | 0.04 | 0 | 0.005 | 0.45 | 0.009 | 0 | 0.001 | 0 | 0 | 0 | 0 | 0 | 0 | 0 | 0 | 0.24 | 0 | 0 | 0 | 0 | 0 | 0 | 0 | 0 | 0 | 0 | 0 | 0.28 | 0 |
| PelMInvertFeeder | 0.049 | 0 | 0 | 0 | 0 | 0 | 0.01 | 0.005 | 0 | 0 | 0 | 0 | 0 | 0 | 0 | 0 | 0 | 0 | 0 | 0 | 0 | 0 | 0 | 0 | 0 | 0 | 0 | 0 | 0 | 0 |
| PelMPredator | 0.01 | 0 | 0 | 0 | 0 | 0.02 | 0.03 | 0.01 | 0 | 0 | 0 | 0 | 0 | 0 | 0 | 0 | 0 | 0 | 0 | 0 | 0 | 0 | 0 | 0 | 0 | 0 | 0 | 0 | 0 | 0 |
| PelLInvertFeeder | 0.025 | 0 | 0 | 0 | 0 | 0 | 0.045 | 0 | 0 | 0 | 0 | 0 | 0 | 0 | 0 | 0 | 0 | 0 | 0 | 0 | 0 | 0 | 0 | 0 | 0 | 0 | 0 | 0 | 0 | 0 |
| PelLPredator | 0 | 0 | 0 | 0 | 0 | 0 | 0.039 | 0 | 0 | 0 | 0 | 0 | 0 | 0 | 0 | 0 | 0 | 0 | 0 | 0 | 0 | 0 | 0 | 0 | 0 | 0 | 0 | 0 | 0 | 0 |
| Mesopelagics | 0.098 | 0.02 | 0 | 0.135 | 0 | 0.005 | 0 | 0 | 0 | 0.002 | 0.065 | 0.068 | 0 | 0 | 0.313 | 0.001 | 0.038 | 0.18 | 0 | 0 | 0 | 0 | 0.837 | 0 | 0 | 0 | 0 | 0 | 0.12 | 0.05 |
| Squid | 0.149 | 0 | 0.055 | 0.27 | 0.1 | 0.07 | 0.1 | 0.222 | 0 | 0.046 | 0 | 0.049 | 0 | 0.007 | 0 | 0.065 | 0 | 0.044 | 0.021 | 0 | 0.018 | 0.007 | 0 | 0.002 | 0 | 0.003 | 0.1 | 0 | 0.025 | 0.2 |
| PelagicPrawns | 0 | 0 | 0 | 0 | 0.05 | 0 | 0 | 0.029 | 0.084 | 0 | 0 | 0.025 | 0.011 | 0.012 | 0 | 0 | 0 | 0 | 0.014 | 0 | 0.006 | 0 | 0.01 | 0.027 | 0 | 0 | 0.05 | 0 | 0.075 | 0.01 |
| Macrobenthos | 0 | 0 | 0 | 0.025 | 0 | 0 | 0.045 | 0.005 | 0.114 | 0.021 | 0 | 0 | 0 | 0.001 | 0.009 | 0.004 | 0.006 | 0 | 0.029 | 0.1 | 0.139 | 0.48 | 0 | 0.215 | 0.1 | 0.1 | 0.05 | 0.36 | 0 | 0 |
| Megabenthos | 0 | 0 | 0 | 0 | 0.05 | 0 | 0.018 | 0.257 | 0.342 | 0.005 | 0 | 0.053 | 0.103 | 0.008 | 0.011 | 0.109 | 0.05 | 0 | 0.096 | 0.1 | 0.117 | 0.012 | 0.013 | 0.197 | 0.1 | 0.575 | 0.151 | 0.28 | 0.008 | 0 |
| Polychaeta | 0 | 0 | 0 | 0 | 0 | 0 | 0 | 0.003 | 0.209 | 0 | 0 | 0.001 | 0 | 0 | 0 | 0.124 | 0.002 | 0 | 0.007 | 0 | 0.05 | 0.464 | 0 | 0.144 | 0 | 0.05 | 0 | 0.21 | 0 | 0 |
| Gelatinous nekton | 0 | 0 | 0 | 0.005 | 0 | 0 | 0 | 0.001 | 0 | 0.901 | 0.412 | 0.001 | 0 | 0 | 0.011 | 0 | 0.024 | 0 | 0.347 | 0.4 | 0.073 | 0 | 0 | 0.003 | 0.056 | 0 | 0 | 0 | 0.001 | 0.7 |
| Euphausids | 0 | 0.1 | 0 | 0 | 0.05 | 0.01 | 0 | 0 | 0 | 0.005 | 0.15 | 0.09 | 0.03 | 0.005 | 0.094 | 0.425 | 0.001 | 0 | 0 | 0 | 0.285 | 0.016 | 0.052 | 0.016 | 0.005 | 0 | 0 | 0 | 0.001 | 0 |
| L zooplankton | 0 | 0 | 0 | 0 | 0 | 0 | 0 | 0.002 | 0.058 | 0 | 0.116 | 0.284 | 0.036 | 0.006 | 0.365 | 0.003 | 0.049 | 0 | 0.04 | 0 | 0.212 | 0.005 | 0.084 | 0.102 | 0.194 | 0.153 | 0.08 | 0.15 | 0.004 | 0 |
| Sm zooplankton | 0 | 0 | 0 | 0 | 0 | 0 | 0 | 0.026 | 0.126 | 0.002 | 0.256 | 0.126 | 0.005 | 0.003 | 0.19 | 0.252 | 0.05 | 0 | 0.038 | 0 | 0.084 | 0.006 | 0.004 | 0.29 | 0.082 | 0.011 | 0 | 0 | 0.004 | 0 |
| Primary producers | 0 | 0 | 0 | 0 | 0 | 0 | 0 | 0 | 0 | 0 | 0 | 0 | 0 | 0 | 0 | 0 | 0 | 0 | 0 | 0 | 0 | 0 | 0 | 0 | 0 | 0 | 0 | 0 | 0 | 0 |
| Benthic producer | 0 | 0 | 0 | 0 | 0 | 0 | 0 | 0 | 0 | 0 | 0 | 0 | 0 | 0 | 0 | 0 | 0 | 0 | 0 | 0 | 0 | 0 | 0 | 0 | 0 | 0.003 | 0 | 0 | 0 | 0 |
| Detritus | 0 | 0 | 0 | 0 | 0 | 0 | 0 | 0 | 0 | 0 | 0 | 0 | 0 | 0 | 0 | 0 | 0 | 0 | 0 | 0 | 0 | 0 | 0 | 0 | 0 | 0 | 0 | 0 | 0 | 0 |
| Discards | 0 | 0 | 0.03 | 0.04 | 0 | 0 | 0 | 0.028 | 0.01 | 0 | 0 | 0 | 0 | 0 | 0 | 0 | 0 | 0 | 0 | 0 | 0 | 0 | 0 | 0 | 0 | 0 | 0.003 | 0 | 0 | 0 |
| Imports | 0.47 | 0.8 | 0 | 0.5 | 0 | 0.5 | 0.25 | 0 | 0 | 0 | 0 | 0 | 0 | 0 | 0 | 0 | 0 | 0 | 0 | 0 | 0 | 0 | 0 | 0 | 0 | 0 | 0 | 0 | 0 | 0 |

**Table S7:** Diet matrix for the SE Australian Ecopath model - continued

| Group | Blue grenadier | SlopeOceanPerch | Deepsea Cod | Oreos | SlopeSmInvertFeeder | SlopeSmPredator | SlopeMInverFeeder | SlopeMPredator | SlopeLInvertFeeder | SlopeLPredator | PelSmInvertFeeder | PelMInvertFeeder | PelMPredator | PelLInvertFeeder | PelLPredator | Mesopelagics | Squid | PelagicPrawns | Macrobenthos | Megabenthos | Polychaeta | Gelatinous nekton | Euphausids | L zooplankton | Sm zooplankton |
|---|---|---|---|---|---|---|---|---|---|---|---|---|---|---|---|---|---|---|---|---|---|---|---|---|---|
| Toothed whale | 0 | 0 | 0 | 0 | 0 | 0 | 0 | 0 | 0 | 0 | 0 | 0 | 0 | 0 | 0 | 0 | 0 | 0 | 0 | 0 | 0 | 0 | 0 | 0 | 0 |
| Baleen whale | 0 | 0 | 0 | 0 | 0 | 0 | 0 | 0 | 0 | 0 | 0 | 0 | 0 | 0 | 0 | 0 | 0 | 0 | 0 | 0 | 0 | 0 | 0 | 0 | 0 |
| Seal | 0 | 0 | 0 | 0 | 0 | 0 | 0 | 0 | 0 | 0 | 0 | 0 | 0 | 0 | 0 | 0 | 0 | 0 | 0 | 0 | 0 | 0 | 0 | 0 | 0 |
| Seabirds | 0 | 0 | 0 | 0 | 0 | 0 | 0 | 0 | 0 | 0 | 0 | 0 | 0 | 0 | 0 | 0 | 0 | 0 | 0 | 0 | 0 | 0 | 0 | 0 | 0 |
| Penguins | 0 | 0 | 0 | 0 | 0 | 0 | 0 | 0 | 0 | 0 | 0 | 0 | 0 | 0 | 0 | 0 | 0 | 0 | 0 | 0 | 0 | 0 | 0 | 0 | 0 |
| Tuna/billfish | 0 | 0 | 0 | 0 | 0 | 0 | 0 | 0 | 0 | 0 | 0 | 0 | 0 | 0 | 0 | 0 | 0 | 0 | 0 | 0 | 0 | 0 | 0 | 0 | 0 |
| Pelagic sharks | 0 | 0 | 0 | 0 | 0 | 0 | 0 | 0 | 0 | 0 | 0 | 0 | 0 | 0 | 0 | 0 | 0 | 0 | 0 | 0 | 0 | 0 | 0 | 0 | 0 |
| Demersal sharks | 0 | 0 | 0 | 0 | 0 | 0 | 0 | 0 | 0 | 0 | 0 | 0 | 0 | 0 | 0 | 0 | 0 | 0 | 0 | 0 | 0 | 0 | 0 | 0 | 0 |
| Rays | 0 | 0 | 0 | 0 | 0 | 0 | 0 | 0 | 0 | 0.01 | 0 | 0 | 0 | 0 | 0 | 0 | 0 | 0 | 0 | 0 | 0 | 0 | 0 | 0 | 0 |
| Warehous | 0 | 0 | 0 | 0 | 0 | 0 | 0 | 0 | 0 | 0.01 | 0 | 0 | 0 | 0 | 0 | 0 | 0 | 0 | 0 | 0 | 0 | 0 | 0 | 0 | 0 |
| Redbait | 0 | 0 | 0 | 0 | 0 | 0 | 0 | 0 | 0 | 0 | 0 | 0 | 0.05 | 0 | 0 | 0 | 0.01 | 0 | 0 | 0 | 0 | 0 | 0 | 0 | 0 |
| Redfish | 0 | 0 | 0 | 0 | 0 | 0 | 0 | 0 | 0 | 0 | 0 | 0 | 0 | 0 | 0 | 0 | 0.002 | 0 | 0 | 0 | 0 | 0 | 0 | 0 | 0 |
| Ling | 0 | 0 | 0 | 0 | 0 | 0 | 0 | 0 | 0 | 0 | 0 | 0 | 0 | 0 | 0 | 0 | 0.001 | 0 | 0 | 0 | 0 | 0 | 0 | 0 | 0 |
| Dories | 0 | 0 | 0 | 0 | 0 | 0 | 0 | 0 | 0 | 0 | 0 | 0 | 0 | 0 | 0 | 0 | 0 | 0 | 0 | 0 | 0 | 0 | 0 | 0 | 0 |
| Jack mackerel | 0 | 0.06 | 0 | 0 | 0 | 0 | 0 | 0 | 0 | 0 | 0 | 0 | 0 | 0 | 0 | 0 | 0.001 | 0 | 0 | 0 | 0 | 0 | 0 | 0 | 0 |
| Jackass morwong | 0 | 0 | 0 | 0 | 0 | 0 | 0 | 0 | 0 | 0 | 0 | 0 | 0 | 0 | 0 | 0 | 0 | 0 | 0 | 0 | 0 | 0 | 0 | 0 | 0 |
| Flathead | 0 | 0 | 0 | 0 | 0 | 0 | 0 | 0 | 0 | 0 | 0 | 0 | 0 | 0 | 0 | 0 | 0 | 0 | 0 | 0 | 0 | 0 | 0 | 0 | 0 |
| Gemfish | 0.016 | 0.059 | 0 | 0 | 0 | 0 | 0 | 0 | 0 | 0 | 0 | 0 | 0 | 0 | 0 | 0 | 0 | 0 | 0 | 0 | 0 | 0 | 0 | 0 | 0 |
| ShOceanPerch | 0 | 0 | 0 | 0 | 0 | 0 | 0 | 0 | 0 | 0 | 0 | 0 | 0 | 0 | 0 | 0 | 0.001 | 0 | 0 | 0 | 0 | 0 | 0 | 0 | 0 |
| Chinaman leatherjacket | 0 | 0 | 0 | 0 | 0 | 0 | 0 | 0 | 0 | 0 | 0 | 0 | 0 | 0 | 0 | 0 | 0 | 0 | 0 | 0 | 0 | 0 | 0 | 0 | 0 |
| Cucumberfish | 0 | 0.009 | 0 | 0 | 0 | 0 | 0 | 0.094 | 0 | 0 | 0 | 0 | 0 | 0 | 0 | 0 | 0 | 0 | 0 | 0 | 0 | 0 | 0 | 0 | 0 |
| Whiting | 0 | 0 | 0 | 0 | 0 | 0 | 0 | 0 | 0 | 0 | 0 | 0 | 0 | 0 | 0 | 0 | 0.001 | 0 | 0 | 0 | 0 | 0 | 0 | 0 | 0 |
| Cardinal | 0.015 | 0.001 | 0 | 0 | 0 | 0 | 0 | 0.249 | 0 | 0 | 0 | 0 | 0 | 0 | 0 | 0 | 0 | 0 | 0 | 0 | 0 | 0 | 0 | 0 | 0 |
| ShSmInvertFeeder | 0.009 | 0.023 | 0 | 0 | 0 | 0 | 0 | 0.036 | 0 | 0 | 0 | 0 | 0.205 | 0 | 0 | 0 | 0.005 | 0 | 0 | 0 | 0 | 0 | 0 | 0 | 0 |
| ShSmPredator | 0 | 0 | 0 | 0 | 0 | 0 | 0 | 0.171 | 0 | 0 | 0 | 0 | 0.015 | 0 | 0 | 0 | 0.005 | 0 | 0 | 0 | 0 | 0 | 0 | 0 | 0 |
| ShMedInvertFeeder | 0 | 0 | 0 | 0 | 0 | 0 | 0 | 0.099 | 0 | 0 | 0 | 0 | 0 | 0 | 0 | 0 | 0.003 | 0 | 0 | 0 | 0 | 0 | 0 | 0 | 0 |
| ShMedPredator | 0 | 0 | 0 | 0 | 0 | 0 | 0 | 0.05 | 0 | 0 | 0 | 0 | 0 | 0 | 0 | 0 | 0.003 | 0 | 0 | 0 | 0 | 0 | 0 | 0 | 0 |
| ShLInvertFeeder | 0 | 0 | 0 | 0 | 0 | 0 | 0 | 0 | 0 | 0 | 0 | 0 | 0 | 0 | 0 | 0 | 0 | 0 | 0 | 0 | 0 | 0 | 0 | 0 | 0 |
| ShLPredator | 0 | 0 | 0 | 0 | 0 | 0 | 0 | 0 | 0 | 0 | 0 | 0 | 0 | 0 | 0 | 0 | 0.008 | 0 | 0 | 0 | 0 | 0 | 0 | 0 | 0 |
| Blue-eye trevalla | 0 | 0 | 0 | 0 | 0 | 0 | 0 | 0 | 0 | 0 | 0 | 0 | 0 | 0 | 0 | 0 | 0 | 0 | 0 | 0 | 0 | 0 | 0 | 0 | 0 |
| Blue grenadier | 0.041 | 0 | 0 | 0 | 0 | 0 | 0 | 0 | 0 | 0 | 0 | 0 | 0 | 0 | 0 | 0 | 0 | 0 | 0 | 0 | 0 | 0 | 0 | 0 | 0 |
| SlopeOceanPerch | 0 | 0.005 | 0 | 0 | 0 | 0 | 0 | 0.015 | 0 | 0 | 0 | 0 | 0 | 0 | 0 | 0 | 0 | 0 | 0 | 0 | 0 | 0 | 0 | 0 | 0 |
| Deepsea Cod | 0 | 0 | 0 | 0 | 0 | 0 | 0 | 0.01 | 0 | 0 | 0 | 0 | 0 | 0 | 0 | 0 | 0 | 0 | 0 | 0 | 0 | 0 | 0 | 0 | 0 |
| Oreos | 0 | 0 | 0 | 0 | 0 | 0 | 0 | 0.005 | 0 | 0 | 0 | 0 | 0 | 0 | 0 | 0 | 0 | 0 | 0 | 0 | 0 | 0 | 0 | 0 | 0 |
| SlopeSmInvertFeeder | 0.018 | 0.004 | 0.05 | 0 | 0 | 0 | 0 | 0 | 0 | 0.16 | 0 | 0 | 0 | 0 | 0 | 0 | 0 | 0 | 0 | 0 | 0 | 0 | 0 | 0 | 0 |

| Group | Blue grenadier | SlopeOceanPerch | Deepsea Cod | Oreos | SlopeSmInvertFeeder | SlopeSmPredator | SlopeMInverFeeder | SlopeMPredator | SlopeLInvertFeeder | SlopeLPredator | PelSmInvertFeeder | PelMInvertFeeder | PelMPredator | PelLInvertFeeder | PelLPredator | Mesopelagics | Squid | PelagicPrawns | Macrobenthos | Megabenthos | Polychaeta | Gelatinous nekton | Euphausids | L zooplankton | Sm zooplankton |
|---|---|---|---|---|---|---|---|---|---|---|---|---|---|---|---|---|---|---|---|---|---|---|---|---|---|
| SlopeSmPredator | 0 | 0 | 0.05 | 0 | 0 | 0 | 0 | 0.011 | 0 | 0.02 | 0 | 0 | 0 | 0 | 0 | 0 | 0 | 0 | 0 | 0 | 0 | 0 | 0 | 0 | 0 |
| SlopeMInverFeeder | 0.118 | 0.206 | 0 | 0 | 0 | 0 | 0.006 | 0.02 | 0 | 0.2 | 0 | 0 | 0 | 0 | 0 | 0 | 0 | 0 | 0 | 0 | 0 | 0 | 0 | 0 | 0 |
| SlopeMPredator | 0 | 0 | 0 | 0 | 0 | 0 | 0 | 0 | 0 | 0.025 | 0 | 0 | 0 | 0 | 0 | 0 | 0.002 | 0 | 0 | 0 | 0 | 0 | 0 | 0 | 0 |
| SlopeLInvertFeeder | 0 | 0 | 0 | 0 | 0 | 0 | 0 | 0 | 0 | 0 | 0 | 0 | 0 | 0 | 0 | 0 | 0 | 0 | 0 | 0 | 0 | 0 | 0 | 0 | 0 |
| SlopeLPredator | 0.008 | 0 | 0 | 0 | 0 | 0 | 0 | 0 | 0 | 0 | 0 | 0 | 0 | 0 | 0 | 0 | 0 | 0 | 0 | 0 | 0 | 0 | 0 | 0 | 0 |
| PelSmInvertFeeder | 0 | 0 | 0 | 0 | 0 | 0 | 0 | 0 | 0 | 0 | 0 | 0.05 | 0.2 | 0 | 0.48 | 0 | 0.16 | 0 | 0 | 0 | 0 | 0 | 0 | 0 | 0 |
| PelMInvertFeeder | 0 | 0 | 0 | 0 | 0 | 0 | 0 | 0 | 0 | 0 | 0 | 0 | 0 | 0 | 0.02 | 0 | 0.002 | 0 | 0 | 0 | 0 | 0 | 0 | 0 | 0 |
| PelMPredator | 0 | 0 | 0 | 0 | 0 | 0 | 0 | 0 | 0 | 0 | 0 | 0 | 0 | 0 | 0 | 0 | 0.002 | 0 | 0 | 0 | 0 | 0 | 0 | 0 | 0 |
| PelLInvertFeeder | 0 | 0 | 0 | 0 | 0 | 0 | 0 | 0 | 0 | 0 | 0 | 0 | 0 | 0 | 0 | 0 | 0 | 0 | 0 | 0 | 0 | 0 | 0 | 0 | 0 |
| PelLPredator | 0 | 0 | 0 | 0 | 0 | 0 | 0 | 0 | 0 | 0 | 0 | 0 | 0 | 0 | 0 | 0 | 0 | 0 | 0 | 0 | 0 | 0 | 0 | 0 | 0 |
| Mesopelagics | 0.717 | 0.068 | 0 | 0.381 | 0 | 0.534 | 0.251 | 0.189 | 0.061 | 0.025 | 0.024 | 0.06 | 0.42 | 0.152 | 0 | 0.07 | 0.215 | 0 | 0 | 0 | 0 | 0 | 0 | 0 | 0 |
| Squid | 0.03 | 0.001 | 0 | 0 | 0 | 0 | 0.012 | 0 | 0.003 | 0.35 | 0.003 | 0 | 0.01 | 0.02 | 0.3 | 0.002 | 0.11 | 0 | 0 | 0 | 0 | 0 | 0 | 0 | 0 |
| PelagicPrawns | 0.009 | 0 | 0 | 0 | 0.037 | 0.006 | 0.034 | 0.012 | 0 | 0.05 | 0 | 0 | 0 | 0 | 0 | 0 | 0.15 | 0 | 0 | 0 | 0 | 0 | 0 | 0 | 0 |
| Macrobenthos | 0.002 | 0.078 | 0.7 | 0.09 | 0.189 | 0.052 | 0.002 | 0 | 0.404 | 0 | 0.003 | 0.1 | 0 | 0.01 | 0 | 0 | 0 | 0 | 0 | 0.5 | 0.05 | 0 | 0 | 0 | 0 |
| Megabenthos | 0.007 | 0.176 | 0.2 | 0.03 | 0.252 | 0.136 | 0.1 | 0 | 0.525 | 0.15 | 0 | 0 | 0 | 0 | 0 | 0 | 0.2 | 0 | 0 | 0 | 0 | 0 | 0 | 0 | 0 |
| Polychaeta | 0 | 0.008 | 0 | 0 | 0.367 | 0.137 | 0.014 | 0.039 | 0.004 | 0 | 0 | 0 | 0 | 0 | 0 | 0 | 0 | 0 | 0 | 0.05 | 0 | 0 | 0 | 0 | 0 |
| Gelatinous nekton | 0.002 | 0.258 | 0 | 0 | 0.003 | 0 | 0 | 0.001 | 0 | 0 | 0 | 0.05 | 0 | 0.009 | 0 | 0 | 0 | 0 | 0 | 0 | 0 | 0 | 0 | 0 | 0 |
| Euphausids | 0 | 0.004 | 0 | 0 | 0.051 | 0.008 | 0 | 0 | 0 | 0 | 0 | 0.3 | 0 | 0 | 0 | 0.785 | 0.07 | 0 | 0 | 0 | 0 | 0 | 0 | 0 | 0 |
| L zooplankton | 0.009 | 0.041 | 0 | 0.399 | 0.008 | 0.045 | 0.581 | 0 | 0.003 | 0 | 0.239 | 0.3 | 0.05 | 0.762 | 0.2 | 0 | 0.05 | 0 | 0.1 | 0 | 0 | 0 | 0 | 0 | 0 |
| Sm zooplankton | 0.001 | 0.001 | 0 | 0.1 | 0.092 | 0.083 | 0.001 | 0 | 0 | 0 | 0.707 | 0.14 | 0.05 | 0.047 | 0 | 0.143 | 0 | 0.2 | 0.4 | 0.4 | 0.05 | 0.9 | 0.2 | 0.2 | 0 |
| Primary producers | 0 | 0 | 0 | 0 | 0 | 0 | 0 | 0 | 0 | 0 | 0.024 | 0 | 0 | 0 | 0 | 0 | 0 | 0.8 | 0 | 0 | 0.5 | 0.1 | 0.8 | 0.8 | 1 |
| Benthic producer | 0 | 0 | 0 | 0 | 0 | 0 | 0 | 0 | 0 | 0 | 0 | 0 | 0 | 0 | 0 | 0 | 0 | 0 | 0 | 0 | 0 | 0 | 0 | 0 | 0 |
| Detritus | 0 | 0 | 0 | 0 | 0 | 0 | 0 | 0 | 0 | 0 | 0 | 0 | 0 | 0 | 0 | 0 | 0 | 0 | 0.5 | 0.05 | 0.4 | 0 | 0 | 0 | 0 |
| Discards | 0 | 0 | 0 | 0 | 0 | 0 | 0 | 0 | 0 | 0 | 0 | 0 | 0 | 0 | 0 | 0 | 0 | 0 | 0 | 0 | 0 | 0 | 0 | 0 | 0 |
| Imports | 0 | 0 | 0 | 0 | 0 | 0 | 0 | 0 | 0 | 0 | 0 | 0 | 0 | 0 | 0 | 0 | 0 | 0 | 0 | 0 | 0 | 0 | 0 | 0 | 0 |

**Table S8**: Landed catch for the SE Australian Ecopath model (units tkm$^{-2}$year$^{-1}$)

| Group | Trawl | Non-trawl | Line | NSW trawl | Vic trawl | Scallop | Squid | Trap | Danish Seine | Tuna Longline | Recreational |
|---|---|---|---|---|---|---|---|---|---|---|---|
| Toothed whale | 0 | 0 | 0 | 0 | 0 | 0 | 0 | 0 | 0 | 0 | 0 |
| Baleen whale | 0 | 0 | 0 | 0 | 0 | 0 | 0 | 0 | 0 | 0 | 0 |
| Seal | 0 | 0 | 0 | 0 | 0 | 0 | 0 | 0 | 0 | 0 | 0 |
| Seabirds | 0 | 0 | 0 | 0 | 0 | 0 | 0 | 0 | 0 | 0 | 0 |
| Penguins | 0 | 0 | 0 | 0 | 0 | 0 | 0 | 0 | 0 | 0 | 0 |
| Tuna/billfish | 0 | 4.97E-04 | 1.45E-02 | 1.00E-07 | 0 | 0 | 0 | 0 | 0 | 6.18E-03 | 8.49E-03 |
| Pelagic sharks | 0 | 0 | 8.58E-05 | 7.00E-07 | 0 | 0 | 0 | 0 | 0 | 2.37E-04 | 1.31E-04 |
| Demersal sharks | 8.04E-04 | 5.61E-03 | 1.25E-04 | 8.14E-04 | 1.23E-04 | 0 | 0 | 0 | 5.43E-04 | 1.20E-05 | 1.29E-03 |
| Rays | 1.21E-03 | 5.00E-07 | 3.60E-06 | 1.72E-04 | 3.00E-05 | 0 | 0 | 0 | 2.40E-05 | 0 | 1.56E-04 |
| Warehous | 5.70E-02 | 7.69E-03 | 0 | 2.21E-05 | 1.28E-03 | 0 | 0 | 0 | 1.08E-04 | 0 | 3.60E-04 |
| Redbait | 1.00E-05 | 0 | 0 | 0 | 0 | 0 | 0 | 0 | 0 | 0 | 0 |
| Redfish | 2.08E-02 | 3.00E-07 | 0 | 8.91E-05 | 1.59E-04 | 0 | 0 | 0 | 3.00E-07 | 0 | 3.19E-05 |
| Ling | 1.47E-02 | 6.75E-04 | 5.38E-04 | 1.50E-05 | 9.52E-04 | 0 | 0 | 0 | 5.70E-05 | 0 | 0 |
| Dories | 7.49E-03 | 1.56E-05 | 0 | 3.26E-04 | 3.53E-04 | 0 | 0 | 0 | 3.14E-04 | 0 | 0 |
| Jack mackerel | 8.28E-04 | 7.76E-03 | 0 | 3.64E-05 | 2.70E-05 | 0 | 0 | 0 | 3.40E-05 | 0 | 6.67E-05 |
| Jackass morwong | 1.58E-02 | 2.60E-06 | 3.00E-07 | 2.75E-04 | 0 | 0 | 0 | 0 | 2.78E-04 | 0 | 0 |
| Flathead | 3.89E-02 | 8.90E-06 | 2.00E-07 | 1.89E-03 | 8.76E-04 | 0 | 0 | 0 | 1.96E-02 | 0 | 6.04E-03 |
| Gemfish | 2.02E-03 | 6.70E-06 | 1.24E-05 | 0 | 4.10E-05 | 0 | 0 | 0 | 0 | 0 | 0 |
| ShOceanPerch | 3.75E-03 | 0 | 9.86E-05 | 7.38E-05 | 1.53E-04 | 0 | 0 | 0 | 3.00E-06 | 0 | 0 |
| Chinaman leatherjacket | 4.88E-04 | 0 | 0 | 0 | 0 | 0 | 0 | 0 | 2.00E-05 | 0 | 0 |
| Cucumberfish | 0 | 0 | 0 | 0 | 0 | 0 | 0 | 0 | 0 | 0 | 0 |
| Whiting | 8.20E-06 | 3.00E-07 | 0 | 4.75E-05 | 6.00E-06 | 0 | 0 | 0 | 1.98E-02 | 0 | 0 |
| Cardinal | 0 | 0 | 0 | 0 | 0 | 0 | 0 | 0 | 0 | 0 | 0 |
| ShSmInvertFeeder | 5.32E-05 | 3.89E-05 | 6.12E-05 | 1.78E-05 | 2.00E-07 | 0 | 0 | 3.30E-06 | 1.68E-04 | 0 | 2.53E-04 |
| ShSmPredator | 3.75E-04 | 0 | 5.00E-07 | 5.70E-06 | 0 | 0 | 0 | 0 | 3.00E-06 | 0 | 2.08E-05 |
| ShMedInvertFeeder | 1.17E-03 | 7.26E-04 | 8.82E-05 | 5.42E-05 | 8.55E-04 | 0 | 0 | 0 | 2.60E-04 | 0 | 1.50E-03 |
| ShMedPredator | 1.36E-03 | 7.69E-04 | 2.54E-03 | 4.39E-04 | 8.80E-05 | 0 | 0 | 4.05E-04 | 6.91E-04 | 0 | 1.88E-04 |
| ShLInvertFeeder | 0 | 5.60E-06 | 3.60E-06 | 3.25E-05 | 0 | 0 | 0 | 0 | 3.30E-05 | 0 | 1.82E-04 |
| ShLPredator | 1.26E-02 | 5.54E-04 | 2.09E-04 | 8.76E-03 | 2.46E-04 | 0 | 0 | 5.00E-07 | 4.80E-05 | 2.23E-04 | 2.82E-03 |
| Blue-eye trevalla | 3.20E-04 | 5.72E-04 | 6.79E-04 | 0 | 5.00E-06 | 0 | 0 | 0 | 3.00E-07 | 0 | 0 |
| Blue grenadier | 4.85E-03 | 2.85E-05 | 1.00E-07 | 0 | 2.09E-04 | 0 | 0 | 0 | 0 | 0 | 0 |
| SlopeOceanPerch | 4.21E-03 | 0 | 0 | 0 | 0 | 0 | 0 | 0 | 0 | 0 | 0 |
| Deepsea Cod | 3.20E-04 | 0 | 0 | 0 | 9.00E-06 | 0 | 0 | 0 | 0 | 0 | 0 |
| Oreos | 6.64E-04 | 0 | 0 | 0 | 0 | 0 | 0 | 0 | 0 | 0 | 0 |
| SlopeSmInvertFeeder | 7.23E-05 | 0 | 0 | 0 | 0 | 0 | 0 | 0 | 0 | 0 | 0 |

| Group | Trawl | Non-trawl | Line | NSW trawl | Vic trawl | Scallop | Squid | Trap | Danish Seine | Tuna Longline | Recreational |
|---|---|---|---|---|---|---|---|---|---|---|---|
| SlopeSmPredator | 2.07E-04 | 0 | 0 | 0 | 0 | 0 | 0 | 0 | 0 | 0 | 0 |
| SlopeMInverFeeder | 1.70E-06 | 2.00E-07 | 0 | 0 | 0 | 0 | 0 | 0 | 0 | 0 | 0 |
| SlopeMPredator | 3.06E-04 | 2.73E-05 | 0 | 0 | 0 | 0 | 0 | 0 | 0 | 0 | 5.42E-07 |
| SlopeLInvertFeeder | 0 | 0 | 0 | 0 | 0 | 0 | 0 | 0 | 0 | 0 | 0 |
| SlopeLPredator | 1.93E-03 | 2.61E-05 | 1.39E-04 | 1.70E-06 | 0 | 0 | 0 | 0 | 0 | 0 | 0 |
| PelSmInvertFeeder | 0 | 5.76E-02 | 1.73E-05 | 0 | 0 | 0 | 0 | 0 | 0 | 0 | 3.39E-05 |
| PelMInvertFeeder | 7.23E-05 | 3.50E-05 | 7.80E-06 | 2.30E-06 | 0 | 0 | 0 | 0 | 1.00E-07 | 0 | 2.70E-04 |
| PelMPredator | 4.80E-06 | 1.63E-02 | 2.60E-04 | 1.55E-04 | 0 | 0 | 0 | 0 | 1.60E-05 | 3.30E-05 | 4.74E-03 |
| PelLInvertFeeder | 0 | 0 | 0 | 0 | 0 | 0 | 0 | 0 | 0 | 0 | 0 |
| PelLPredator | 0 | 4.90E-06 | 1.44E-05 | 6.00E-07 | 0 | 0 | 0 | 0 | 1.00E-07 | 2.20E-05 | 2.02E-03 |
| Mesopelagics | 0 | 0 | 0 | 0 | 0 | 0 | 0 | 0 | 0 | 0 | 0 |
| Squid | 2.81E-03 | 1.14E-05 | 8.00E-07 | 2.51E-04 | 3.12E-04 | 0 | 4.00E-06 | 8.60E-06 | 0 | 0 | 2.29E-04 |
| PelagicPrawns | 0 | 2.20E-06 | 0 | 1.60E-04 | 3.60E-05 | 0 | 0 | 0 | 0 | 0 | 0 |
| Macrobenthos | 0 | 0 | 0 | 3.84E-04 | 3.07E-04 | 0 | 0 | 0 | 0 | 0 | 0 |
| Megabenthos | 1.13E-02 | 1.50E-06 | 0 | 8.22E-04 | 1.08E-03 | 4.96E-03 | 0 | 1.31E-04 | 0 | 0 | 0 |
| Polychaeta | 0 | 0 | 0 | 0 | 0 | 0 | 0 | 0 | 0 | 0 | 0 |
| Gelatinous nekton | 0 | 0 | 0 | 0 | 0 | 0 | 0 | 0 | 0 | 0 | 0 |
| Euphausids | 0 | 0 | 0 | 0 | 0 | 0 | 0 | 0 | 0 | 0 | 0 |
| L zooplankton | 0 | 0 | 0 | 0 | 0 | 0 | 0 | 0 | 0 | 0 | 0 |
| Sm zooplankton | 0 | 0 | 0 | 0 | 0 | 0 | 0 | 0 | 0 | 0 | 0 |
| Primary producers | 0 | 0 | 0 | 0 | 0 | 0 | 0 | 0 | 0 | 0 | 0 |
| Benthic producer | 0 | 0 | 0 | 0 | 0 | 0 | 0 | 0 | 0 | 0 | 0 |
| Detritus | 0 | 0 | 0 | 0 | 0 | 0 | 0 | 0 | 0 | 0 | 0 |
| Discards | 0 | 0 | 0 | 0 | 0 | 0 | 0 | 0 | 0 | 0 | 0 |
| Imports | 0 | 0 | 0 | 0 | 0 | 0 | 0 | 0 | 0 | 0 | 0 |

**Table S9**: Discards for the SE Australian Ecopath model (units tkm$^{-2}$year$^{-1}$)

| Group | Trawl | Non-trawl | Line | NSW trawl | Vic trawl | Scallop | Squid | Trap | Danish Seine | Tuna Longline | Recreational |
|---|---|---|---|---|---|---|---|---|---|---|---|
| Toothed whale | 0 | 0 | 0 | 0 | 0 | 0 | 0 | 0 | 0 | 0 | 0 |
| Baleen whale | 0 | 0 | 0 | 0 | 0 | 0 | 0 | 0 | 0 | 0 | 0 |
| Seal | 4.45E-04 | 0 | 0 | 3.27E-05 | 1.57E-05 | 0 | 0 | 0 | 0 | 0 | 0 |
| Seabirds | 0 | 0 | 0 | 0 | 0 | 0 | 0 | 0 | 0 | 0 | 0 |
| Penguins | 0 | 0 | 0 | 0 | 0 | 0 | 0 | 0 | 0 | 0 | 0 |
| Tuna/billfish | 0 | 5.00E-06 | 1.45E-04 | 0 | 0 | 0 | 0 | 0 | 0 | 1.23E-05 | 0 |
| Pelagic sharks | 0 | 0 | 9.00E-07 | 0 | 0 | 0 | 0 | 0 | 0 | 4.00E-09 | 0 |
| Demersal sharks | 4.97E-03 | 5.61E-05 | 1.30E-06 | 8.15E-04 | 1.23E-04 | 0 | 0 | 0 | 4.95E-03 | 1.00E-07 | 0 |
| Rays | 3.05E-03 | 5.00E-09 | 4.00E-08 | 1.73E-04 | 3.00E-05 | 0 | 0 | 0 | 8.46E-03 | 0 | 0 |
| Warehous | 1.69E-03 | 7.69E-05 | 0 | 2.33E-05 | 1.36E-03 | 0 | 0 | 0 | 5.97E-05 | 0 | 0 |
| Redbait | 0 | 0 | 0 | 0 | 0 | 0 | 0 | 0 | 0 | 0 | 0 |
| Redfish | 2.00E-02 | 3.00E-09 | 0 | 9.10E-05 | 1.62E-04 | 0 | 0 | 0 | 1.20E-06 | 0 | 0 |
| Ling | 1.67E-04 | 6.80E-06 | 5.40E-06 | 1.52E-05 | 9.66E-04 | 0 | 0 | 0 | 2.11E-05 | 0 | 0 |
| Dories | 3.39E-03 | 2.00E-07 | 0 | 3.28E-04 | 3.56E-04 | 0 | 0 | 0 | 7.76E-05 | 0 | 0 |
| Jack mackerel | 1.31E-02 | 7.76E-05 | 0 | 3.64E-05 | 2.66E-05 | 0 | 0 | 0 | 2.94E-05 | 0 | 0 |
| Jackass morwong | 4.24E-04 | 3.00E-08 | 3.00E-09 | 2.79E-04 | 1.50E-05 | 0 | 0 | 0 | 2.80E-06 | 0 | 0 |
| Flathead | 7.21E-04 | 1.00E-07 | 2.00E-09 | 1.97E-03 | 9.11E-04 | 0 | 0 | 0 | 5.00E-04 | 0 | 0 |
| Gemfish | 1.59E-03 | 1.00E-07 | 1.00E-07 | 1.16E-04 | 4.10E-05 | 0 | 0 | 0 | 1.01E-05 | 0 | 0 |
| ShOceanPerch | 3.79E-03 | 0 | 1.00E-06 | 7.41E-05 | 1.54E-04 | 0 | 0 | 0 | 2.90E-06 | 0 | 0 |
| Chinaman leatherjacket | 1.12E-05 | 0 | 0 | 8.00E-07 | 4.00E-07 | 0 | 0 | 0 | 0 | 0 | 0 |
| Cucumberfish | 1.08E-03 | 0 | 0 | 7.93E-05 | 3.82E-05 | 0 | 0 | 0 | 8.10E-04 | 0 | 0 |
| Whiting | 5.00E-07 | 3.00E-09 | 0 | 4.75E-05 | 5.90E-06 | 0 | 0 | 0 | 9.91E-04 | 0 | 0 |
| Cardinal | 3.60E-05 | 0 | 0 | 3.21E-04 | 1.14E-05 | 0 | 0 | 0 | 0 | 0 | 0 |
| ShSmInvertFeeder | 1.47E-02 | 4.00E-07 | 6.00E-07 | 1.78E-05 | 2.00E-07 | 0 | 0 | 3.00E-08 | 1.19E-02 | 0 | 0 |
| ShSmPredator | 3.77E-04 | 0 | 1.00E-08 | 5.70E-06 | 1.33E-05 | 0 | 0 | 0 | 1.86E-05 | 0 | 0 |
| ShMedInvertFeeder | 9.10E-03 | 7.30E-06 | 9.00E-07 | 5.43E-05 | 8.56E-04 | 0 | 0 | 0 | 9.71E-03 | 0 | 0 |
| ShMedPredator | 1.68E-04 | 7.70E-06 | 2.54E-05 | 4.40E-04 | 8.76E-05 | 0 | 0 | 4.00E-06 | 6.02E-05 | 0 | 0 |
| ShLInvertFeeder | 0 | 1.00E-07 | 3.60E-08 | 0 | 0 | 0 | 0 | 0 | 0 | 0 | 0 |
| ShLPredator | 1.42E-02 | 5.50E-06 | 2.10E-06 | 8.87E-03 | 2.49E-04 | 0 | 0 | 0 | 3.20E-06 | 0 | 0 |
| Blue-eye trevalla | 0 | 5.70E-06 | 6.80E-06 | 0 | 0 | 0 | 0 | 0 | 0 | 0 | 0 |
| Blue grenadier | 4.80E-06 | 3.00E-07 | 1.00E-09 | 0 | 2.10E-04 | 0 | 0 | 0 | 0 | 0 | 0 |
| SlopeOceanPerch | 2.36E-04 | 0 | 0 | 0 | 0 | 0 | 0 | 0 | 0 | 0 | 0 |
| Deepsea Cod | 3.20E-05 | 0 | 0 | 0 | 0 | 0 | 0 | 0 | 0 | 0 | 0 |
| Oreos | 2.30E-06 | 0 | 0 | 0 | 0 | 0 | 0 | 0 | 0 | 0 | 0 |
| SlopeSmInvertFeeder | 8.11E-05 | 0 | 0 | 0 | 0 | 0 | 0 | 0 | 2.50E-06 | 0 | 0 |

| Group | Trawl | Non-trawl | Line | NSW trawl | Vic trawl | Scallop | Squid | Trap | Danish Seine | Tuna Longline | Recreational |
|---|---|---|---|---|---|---|---|---|---|---|---|
| SlopeSmPredator | 1.31E-03 | 0 | 0 | 0 | 0 | 0 | 0 | 0 | 0 | 0 | 0 |
| SlopeMInverFeeder | 8.12E-03 | 2.00E-09 | 0 | 0 | 0 | 0 | 0 | 0 | 0 | 0 | 0 |
| SlopeMPredator | 2.27E-04 | 3.00E-07 | 0 | 0 | 0 | 0 | 0 | 0 | 0 | 0 | 0 |
| SlopeLInvertFeeder | 1.00E-07 | 0 | 0 | 0 | 0 | 0 | 0 | 0 | 0 | 0 | 0 |
| SlopeLPredator | 1.84E-03 | 3.00E-07 | 1.40E-06 | 1.70E-06 | 0 | 0 | 0 | 0 | 8.00E-07 | 0 | 0 |
| PelSmInvertFeeder | 0 | 5.76E-04 | 2.00E-07 | 0 | 0 | 0 | 0 | 0 | 0 | 0 | 0 |
| PelMInvertFeeder | 9.00E-07 | 4.00E-07 | 1.00E-07 | 2.30E-06 | 0 | 0 | 0 | 0 | 1.70E-06 | 0 | 0 |
| PelMPredator | 8.70E-06 | 1.63E-04 | 2.60E-06 | 1.55E-04 | 0 | 0 | 0 | 0 | 0 | 0 | 0 |
| PelLInvertFeeder | 0 | 0 | 0 | 0 | 0 | 0 | 0 | 0 | 0 | 0 | 0 |
| PelLPredator | 0 | 5.00E-08 | 1.00E-07 | 0 | 0 | 0 | 0 | 0 | 0 | 0 | 0 |
| Mesopelagics | 8.10E-07 | 0 | 0 | 0 | 0 | 0 | 0 | 0 | 0 | 0 | 0 |
| Squid | 0 | 1.00E-07 | 1.00E-08 | 0 | 0 | 0 | 0 | 1.00E-07 | 0 | 0 | 0 |
| PelagicPrawns | 0 | 2.00E-08 | 0 | 0 | 0 | 0 | 0 | 0 | 0 | 0 | 0 |
| Macrobenthos | 1.15E-03 | 0 | 0 | 3.84E-04 | 3.07E-04 | 0 | 0 | 0 | 0 | 0 | 0 |
| Megabenthos | 1.53E-04 | 1.00E-08 | 0 | 8.31E-04 | 1.09E-03 | 2.48E-04 | 0 | 1.30E-06 | 0 | 0 | 0 |
| Polychaeta | 0 | 0 | 0 | 0 | 0 | 0 | 0 | 0 | 0 | 0 | 0 |
| Gelatinous nekton | 1.00E-06 | 0 | 0 | 1.00E-06 | 9.00E-08 | 0 | 0 | 0 | 0 | 0 | 0 |
| Euphausids | 0 | 0 | 0 | 0 | 0 | 0 | 0 | 0 | 0 | 0 | 0 |
| L zooplankton | 0 | 0 | 0 | 0 | 0 | 0 | 0 | 0 | 0 | 0 | 0 |
| Sm zooplankton | 0 | 0 | 0 | 0 | 0 | 0 | 0 | 0 | 0 | 0 | 0 |
| Primary producers | 0 | 0 | 0 | 0 | 0 | 0 | 0 | 0 | 0 | 0 | 0 |
| Benthic producer | 0 | 0 | 0 | 0 | 0 | 0 | 0 | 0 | 0 | 0 | 0 |
| Detritus | 0 | 0 | 0 | 0 | 0 | 0 | 0 | 0 | 0 | 0 | 0 |
| Discards | 0 | 0 | 0 | 0 | 0 | 0 | 0 | 0 | 0 | 0 | 0 |
| Imports | 0 | 0 | 0 | 0 | 0 | 0 | 0 | 0 | 0 | 0 | 0 |

**Table S10:** Ecopath base parameters for the Kerala Ecopath model

| Group | Trophic Level | Biomass (tkm$^{-2}$) | Production / Biomass (year$^{-1}$) | Consumption / Biomass (year$^{-1}$) | Ecotrophic Efficiency |
|---|---|---|---|---|---|
| Marine Mammals | 3.976 | 0.019 | 0.1 | 12.75 | 0 |
| Sharks coastal | 4.699 | 0.042 | 1.181 | 3.7 | 0.747 |
| Sharks offshore | 4.991 | 0.037 | 1.098 | 3.43 | 0.806 |
| Sharks demersal | 4.772 | 0.018 | 0.93 | 4.463 | 0.476 |
| Guitarfishes & Rays | 4.35 | 0.077 | 0.89 | 2.964 | 0.364 |
| Large pelagics-offshore | 4.482 | 0.342 | 1.053 | 2.698 | 0.663 |
| Large Pelagics-inshore | 4.751 | 0.205 | 1.206 | 5.091 | 0.989 |
| Large Bentho Pelagics shelf | 4.477 | 0.044 | 0.694 | 4.757 | 0.934 |
| Large benthopelagic deep | 3.899 | 0.62 | 0.563 | 2.254 | 0.98 |
| Medium Bentho Pelagics shelf | 4.183 | 1.519 | 1.425 | 8.904 | 0.996 |
| Medium benthopelagic deep | 4.468 | 0.707 | 1.012 | 6.467 | 0.95 |
| Small Bentho Pelagics shelf | 4.002 | 1.55 | 2.105 | 10.95 | 0.919 |
| Small benthopelagic deep | 3.767 | 3.5 | 1.713 | 13 | 0.998 |
| Mesopelagics | 3.56 | 1.2 | 1.68 | 11.602 | 0.762 |
| Tunas coastal | 4.234 | 0.44 | 1.552 | 13.6 | 0.673 |
| Tunas offshore | 4.323 | 0.193 | 1.172 | 17.603 | 0.743 |
| Ribbonfishes | 4.549 | 0.852 | 1.292 | 3.75 | 0.938 |
| Large Benthic Carnivores shelf | 4.129 | 1.667 | 0.778 | 5.324 | 0.993 |
| Large Benthic Carnivores deep | 4.43 | 0.145 | 0.623 | 4.32 | 0.95 |
| Medium Benthic Carnivores shelf | 3.653 | 1.687 | 1.386 | 7.697 | 0.792 |
| Medium Benthic Carnivores deep | 3.953 | 4.795 | 0.87 | 4.538 | 0.95 |
| Small Benthic Carnivores shelf | 3.019 | 2.95 | 2.17 | 11.272 | 0.968 |
| Small Benthic Carnivores deep | 3.775 | 8.812 | 0.925 | 4.484 | 0.95 |
| Benthic omnivores shelf | 2.598 | 0.288 | 0.94 | 26.98 | 0.98 |
| Threadfin breams | 3.811 | 1.364 | 2.046 | 10.938 | 1 |
| Indian Mackerel | 2.491 | 1.889 | 2.259 | 21.7 | 1 |
| Oil Sardine | 2.204 | 7.856 | 2.605 | 27.6 | 1 |
| Other Clupeids inshore | 3.299 | 1.5 | 2.884 | 20.7 | 0.999 |
| Anchovies | 3.457 | 8 | 2.39 | 17.133 | 0.994 |
| Crabs & Lobsters | 3.343 | 0.091 | 4.785 | 8.5 | 0.997 |
| Deep Sea Shrimps | 3.151 | 0.75 | 3.941 | 10.9 | 0.97 |
| Coastal Shrimps | 3.152 | 5.3 | 6.87 | 19.2 | 0.999 |
| Squids | 3.717 | 1.592 | 4.25 | 16.64 | 0.946 |
| Cuttlefishes | 4.221 | 0.59 | 3.552 | 16.64 | 0.998 |
| Octopus | 3.913 | 0.15 | 3.81 | 12.5 | 0.999 |
| Commercial molluscs | 2.364 | 0.44 | 4.301 | 12.5 | 0.996 |
| Benthic detritivore | 2.295 | 53.478 | 2.9 | 9 | 0.95 |
| Macrozobenthos (Benthic carnivore) | 3.278 | 15 | 3.1 | 12.482 | 0.991 |
| Benthic grazers | 2 | 30.558 | 0.73 | 2.5 | 0.95 |
| Shelf filter feeders | 2 | 5 | 1.6 | 12 | 0.719 |
| Deep filter feeders | 2 | 4 | 1.2 | 12 | 0.274 |
| Gelatinous zooplankton | 2.645 | 3.212 | 4.85 | 16.167 | 0.75 |
| Large Zooplankton | 2.58 | 6.538 | 35.3 | 160 | 0.75 |
| Micro Zooplankton | 2 | 14 | 67.5 | 250 | 0.818 |
| Phytoplankton | 1 | 58.5 | 70 | | 0.953 |
| Macroalgae | 1 | 1554.967 | 2.78 | | 0.95 |
| Detritus | 1 | 15 | | | 0.9 |

**Table S11:** Diet matrix for the Kerala Ecopath model

| Group | Marine Mammals | Sharks coastal | Sharks offshore | Sharks demersal | Guitarfishes & Rays | Large pelagics-offshore | Large Pelagics-inshore | Large Bentho Pelagics shelf | Large benthopelagic deep | Medium Bentho Pelagics shelf | Medium benthopelagic deep | Small Bentho Pelagics shelf | Small benthopelagic deep | Mesopelagics | Tunas coastal | Tunas offshore | Ribbonfishes | Large Benthic Carnivores shelf | Large Benthic Carnivores deep | Medium Benthic Carnivores shelf | Medium Benthic Carnivores deep | Small Benthic Carnivores shelf | Small Benthic Carnivores deep | Benthic omnivores shelf | Threadfin breams | Indian Mackerel | Oil Sardine | Other Clupeids inshore | Anchovies | Crabs & Lobsters |
|---|---|---|---|---|---|---|---|---|---|---|---|---|---|---|---|---|---|---|---|---|---|---|---|---|---|---|---|---|---|---|
| Marine Mammals | 0 | 0 | 0 | 0 | 0 | 0 | 0 | 0 | 0 | 0 | 0 | 0 | 0 | 0 | 0 | 0 | 0 | 0 | 0 | 0 | 0 | 0 | 0 | 0 | 0 | 0 | 0 | 0 | 0 | 0 |
| Sharks coastal | 0 | 0 | 0 | 0 | 0 | 0 | 0 | 0 | 0 | 0 | 0 | 0 | 0 | 0 | 0 | 0 | 0 | 0 | 0 | 0 | 0 | 0 | 0 | 0 | 0 | 0 | 0 | 0 | 0 | 0 |
| Sharks offshore | 0 | 0 | 0 | 0 | 0 | 0 | 0 | 0 | 0 | 0 | 0 | 0 | 0 | 0 | 0 | 0 | 0 | 0 | 0 | 0 | 0 | 0 | 0 | 0 | 0 | 0 | 0 | 0 | 0 | 0 |
| Sharks demersal | 0 | 0 | 0 | 0 | 0 | 0 | 0 | 0 | 0 | 0 | 0 | 0 | 0 | 0 | 0 | 0 | 0 | 0 | 0 | 0 | 0 | 0 | 0 | 0 | 0 | 0 | 0 | 0 | 0 | 0 |
| Guitarfishes & Rays | 0 | 0 | 0 | 0 | 0 | 0 | 0 | 0 | 0 | 0 | 0 | 0 | 0 | 0 | 0 | 0 | 0 | 0 | 0 | 0 | 0 | 0 | 0 | 0 | 0 | 0 | 0 | 0 | 0 | 0 |
| Large pelagics-offshore | 0 | 0 | 0 | 0 | 0 | 0 | 0.053 | 0 | 0 | 0.005 | 0 | 0 | 0 | 0 | 0 | 0 | 0 | 0 | 0 | 0 | 0 | 0 | 0 | 0 | 0 | 0 | 0 | 0 | 0 | 0 |
| Large Pelagics-inshore | 0 | 0.054 | 0 | 0 | 0 | 0 | 0 | 0 | 0 | 0 | 0 | 0 | 0 | 0 | 0 | 0 | 0 | 0 | 0 | 0 | 0 | 0 | 0 | 0 | 0 | 0 | 0 | 0 | 0.054 | 0 |
| Large Bentho Pelagics shelf | 0 | 0 | 0 | 0 | 0 | 0 | 0 | 0 | 0 | 0 | 0 | 0 | 0 | 0 | 0 | 0 | 0 | 0 | 0 | 0 | 0 | 0 | 0 | 0 | 0 | 0 | 0 | 0 | 0 | 0 |
| Large benthopelagic deep | 0 | 0 | 0 | 0 | 0 | 0.003 | 0 | 0 | 0 | 0 | 0 | 0 | 0 | 0 | 0 | 0.1 | 0 | 0 | 0 | 0 | 0 | 0 | 0 | 0 | 0 | 0 | 0 | 0 | 0 | 0 |
| Medium Bentho Pelagics shelf | 0 | 0.059 | 0.16 | 0 | 0 | 0.042 | 0.184 | 0 | 0 | 0.02 | 0 | 0 | 0 | 0 | 0 | 0 | 0.19 | 0.002 | 0 | 0 | 0 | 0 | 0 | 0 | 0 | 0 | 0 | 0 | 0.059 | 0.16 |
| Medium benthopelagic deep | 0 | 0 | 0 | 0 | 0 | 0 | 0 | 0 | 0 | 0 | 0 | 0 | 0 | 0 | 0 | 0.2 | 0 | 0 | 0 | 0 | 0 | 0 | 0 | 0 | 0 | 0 | 0 | 0 | 0 | 0 |
| Small Bentho Pelagics shelf | 0 | 0.359 | 0 | 0.005 | 0 | 0 | 0.11 | 0.078 | 0 | 0.111 | 0.009 | 0 | 0 | 0 | 0.018 | 0.013 | 0.053 | 0 | 0 | 0.003 | 0 | 0 | 0 | 0 | 0 | 0 | 0 | 0 | 0.359 | 0 |
| Small benthopelagic deep | 0 | 0 | 0.159 | 0.116 | 0.01 | 0.234 | 0 | 0 | 0 | 0 | 0.255 | 0 | 0.05 | 0 | 0.2 | 0 | 0 | 0 | 0 | 0 | 0.05 | 0 | 0 | 0 | 0 | 0 | 0 | 0 | 0 | 0.159 |
| Mesopelagics | 0 | 0 | 0 | 0 | 0 | 0 | 0 | 0 | 0 | 0.002 | 0.21 | 0 | 0 | 0.01 | 0.05 | 0 | 0 | 0 | 0.1 | 0 | 0.01 | 0 | 0 | 0 | 0 | 0 | 0 | 0 | 0 | 0 |
| Tunas coastal | 0 | 0 | 0 | 0 | 0 | 0 | 0.001 | 0 | 0 | 0.003 | 0 | 0 | 0 | 0 | 0 | 0 | 0 | 0 | 0 | 0.003 | 0 | 0.003 | 0 | 0 | 0 | 0 | 0 | 0 | 0 | 0 |
| Tunas offshore | 0 | 0 | 0.345 | 0 | 0 | 0 | 0 | 0 | 0 | 0 | 0 | 0 | 0 | 0 | 0 | 0 | 0 | 0 | 0 | 0 | 0 | 0 | 0 | 0 | 0 | 0 | 0 | 0 | 0 | 0.345 |
| Ribbonfishes | 0 | 0 | 0 | 0 | 0.001 | 0 | 0.059 | 0 | 0 | 0.007 | 0 | 0 | 0 | 0 | 0 | 0 | 0.055 | 0.02 | 0 | 0.007 | 0 | 0.001 | 0 | 0 | 0 | 0 | 0 | 0 | 0 | 0 |
| Large Benthic Carnivores shelf | 0.113 | 0 | 0 | 0 | 0.007 | 0.005 | 0.093 | 0.126 | 0 | 0.008 | 0 | 0 | 0 | 0 | 0.01 | 0 | 0.038 | 0.019 | 0 | 0.008 | 0 | 0 | 0 | 0 | 0.002 | 0 | 0 | 0.113 | 0 | 0 |
| Large Benthic Carnivores deep | 0 | 0 | 0 | 0 | 0 | 0 | 0 | 0 | 0 | 0 | 0 | 0 | 0 | 0 | 0 | 0 | 0 | 0 | 0.011 | 0 | 0 | 0 | 0 | 0 | 0 | 0 | 0 | 0 | 0 | 0 |
| Medium Benthic Carnivores shelf | 0 | 0.025 | 0 | 0 | 0 | 0.003 | 0.043 | 0 | 0 | 0.006 | 0 | 0 | 0 | 0 | 0 | 0 | 0 | 0.025 | 0.013 | 0.029 | 0 | 0 | 0 | 0 | 0.001 | 0 | 0 | 0 | 0.025 | 0 |
| Medium Benthic Carniv. deep | 0 | 0 | 0.095 | 0.55 | 0 | 0 | 0.026 | 0 | 0 | 0 | 0 | 0 | 0 | 0 | 0 | 0 | 0 | 0 | 0 | 0.003 | 0.01 | 0 | 0 | 0 | 0 | 0 | 0 | 0 | 0 | 0.095 |
| Small Benthic Carnivores shelf | 0 | 0 | 0.02 | 0 | 0.093 | 0.026 | 0.009 | 0 | 0 | 0.059 | 0 | 0 | 0 | 0 | 0.001 | 0 | 0 | 0.11 | 0.243 | 0.066 | 0 | 0.001 | 0 | 0 | 0.08 | 0 | 0 | 0 | 0 | 0.02 |
| Small Benthic Carnivores deep | 0 | 0 | 0 | 0 | 0 | 0 | 0 | 0 | 0 | 0 | 0.055 | 0 | 0.04 | 0 | 0 | 0 | 0 | 0 | 0 | 0.02 | 0.11 | 0 | 0 | 0 | 0 | 0 | 0 | 0 | 0 | 0 |
| Benthic omnivores shelf | 0 | 0.05 | 0 | 0.05 | 0.05 | 0 | 0 | 0 | 0 | 0 | 0 | 0 | 0 | 0 | 0 | 0 | 0 | 0.055 | 0 | 0 | 0 | 0 | 0 | 0 | 0 | 0 | 0 | 0 | 0.05 | 0 |

| Group | Marine Mammals | Sharks coastal | Sharks offshore | Sharks demersal | Guitarfishes & Rays | Large pelagics-offshore | Large Pelagics-inshore | Large Bentho Pelagics shelf | Large benthopelagic deep | Medium Bentho Pelagics shelf | Medium benthopelagic deep | Small Bentho Pelagics shelf | Small benthopelagic deep | Mesopelagics | Tunas coastal | Tunas offshore | Ribbonfishes | Large Benthic Carnivores shelf | Large Benthic Carnivores deep | Medium Benthic Carnivores shelf | Medium Benthic Carnivores deep | Small Benthic Carnivores shelf | Small Benthic Carnivores deep | Benthic omnivores shelf | Threadfin breams | Indian Mackerel | Oil Sardine | Other Clupeids inshore | Anchovies | Crabs & Lobsters |
|---|---|---|---|---|---|---|---|---|---|---|---|---|---|---|---|---|---|---|---|---|---|---|---|---|---|---|---|---|---|---|
| Threadfin breams | 0.116 | 0 | 0 | 0 | 0.325 | 0.048 | 0.097 | 0 | 0 | 0.04 | 0.015 | 0 | 0 | 0 | 0 | 0 | 0 | 0.032 | 0.096 | 0.03 | 0 | 0 | 0 | 0 | 0.006 | 0 | 0 | 0.116 | 0 | 0 |
| Indian Mackerel | 0.099 | 0 | 0 | 0 | 0 | 0.035 | 0.026 | 0.008 | 0 | 0.054 | 0 | 0 | 0 | 0 | 0 | 0.2 | 0 | 0 | 0 | 0.009 | 0 | 0 | 0 | 0 | 0 | 0 | 0 | 0.099 | 0 | 0 |
| Oil Sardine | 0.35 | 0 | 0 | 0 | 0 | 0.047 | 0.007 | 0.031 | 0 | 0.06 | 0 | 0 | 0 | 0 | 0.158 | 0.1 | 0.038 | 0.014 | 0 | 0.029 | 0 | 0 | 0 | 0 | 0 | 0 | 0 | 0.35 | 0 | 0 |
| Other Clupeids inshore | 0.015 | 0 | 0.042 | 0 | 0 | 0.007 | 0.027 | 0 | 0 | 0.029 | 0 | 0 | 0 | 0 | 0.006 | 0 | 0.012 | 0 | 0 | 0.001 | 0 | 0 | 0 | 0 | 0 | 0 | 0 | 0.015 | 0 | 0.042 |
| Anchovies | 0.012 | 0.096 | 0 | 0.098 | 0.013 | 0.434 | 0.04 | 0.633 | 0 | 0.206 | 0 | 0.061 | 0 | 0 | 0.203 | 0 | 0.38 | 0.166 | 0 | 0.038 | 0 | 0.07 | 0 | 0 | 0.02 | 0 | 0 | 0.012 | 0.096 | 0 |
| Crabs & Lobsters | 0.035 | 0.019 | 0 | 0 | 0.018 | 0.004 | 0.007 | 0 | 0 | 0.005 | 0 | 0 | 0 | 0 | 0.002 | 0 | 0 | 0.005 | 0 | 0.003 | 0 | 0.001 | 0 | 0 | 0 | 0 | 0 | 0.035 | 0.019 | 0 |
| Deep Sea Shrimps | 0.12 | 0 | 0 | 0 | 0.216 | 0.005 | 0.001 | 0.009 | 0 | 0 | 0.02 | 0 | 0 | 0 | 0.025 | 0 | 0 | 0.007 | 0.049 | 0.042 | 0.02 | 0 | 0.001 | 0 | 0 | 0 | 0 | 0.12 | 0 | 0 |
| Coastal Shrimps | 0.099 | 0.012 | 0 | 0 | 0.251 | 0.002 | 0.001 | 0.092 | 0 | 0.129 | 0 | 0.1 | 0 | 0 | 0.326 | 0 | 0.14 | 0.073 | 0.184 | 0.1 | 0.01 | 0.055 | 0.001 | 0 | 0.1 | 0 | 0 | 0.099 | 0.012 | 0 |
| Squids | 0.04 | 0 | 0.13 | 0.181 | 0 | 0.073 | 0.021 | 0.004 | 0.1 | 0.033 | 0.05 | 0.016 | 0.01 | 0.01 | 0 | 0.1 | 0.046 | 0.13 | 0.02 | 0.05 | 0.08 | 0 | 0 | 0 | 0.066 | 0 | 0 | 0.04 | 0 | 0.13 |
| Cuttlefishes | 0 | 0.017 | 0 | 0 | 0 | 0 | 0.002 | 0 | 0.1 | 0 | 0 | 0 | 0 | 0 | 0 | 0.06 | 0 | 0.045 | 0.02 | 0.05 | 0.011 | 0 | 0 | 0 | 0.01 | 0 | 0 | 0 | 0.017 | 0 |
| Octopus | 0 | 0.187 | 0.042 | 0 | 0 | 0.009 | 0 | 0 | 0 | 0.008 | 0 | 0 | 0 | 0 | 0 | 0 | 0 | 0 | 0.19 | 0.002 | 0.006 | 0 | 0 | 0 | 0.001 | 0 | 0 | 0 | 0.187 | 0.042 |
| Commercial molluscs | 0 | 0.107 | 0 | 0 | 0 | 0 | 0 | 0 | 0 | 0 | 0.001 | 0 | 0 | 0 | 0.001 | 0 | 0 | 0.001 | 0 | 0.014 | 0.001 | 0.003 | 0 | 0 | 0.02 | 0 | 0 | 0 | 0.107 | 0 |
| Benthic detritivore | 0 | 0 | 0 | 0 | 0 | 0 | 0.067 | 0 | 0.01 | 0.072 | 0.02 | 0.15 | 0.05 | 0 | 0 | 0.1 | 0.03 | 0 | 0 | 0.042 | 0.187 | 0.09 | 0.3 | 0.025 | 0.081 | 0 | 0 | 0 | 0 | 0 |
| Macrozobenthos (Benthic carnivore) | 0 | 0.015 | 0 | 0 | 0.016 | 0.02 | 0.126 | 0.007 | 0.065 | 0.07 | 0.31 | 0.523 | 0.1 | 0 | 0 | 0.125 | 0 | 0.096 | 0.075 | 0.029 | 0.216 | 0.11 | 0.4 | 0 | 0.33 | 0 | 0 | 0 | 0.015 | 0 |
| Benthic grazers | 0 | 0 | 0 | 0 | 0 | 0 | 0 | 0 | 0.025 | 0.067 | 0 | 0.05 | 0 | 0 | 0 | 0 | 0 | 0.14 | 0 | 0.11 | 0.053 | 0.048 | 0 | 0 | 0.19 | 0 | 0 | 0 | 0 | 0 |
| Shelf filter feeders | 0 | 0 | 0 | 0 | 0 | 0 | 0 | 0 | 0 | 0 | 0 | 0 | 0 | 0 | 0 | 0 | 0 | 0.06 | 0 | 0.046 | 0.073 | 0.052 | 0 | 0 | 0.02 | 0 | 0 | 0 | 0 | 0 |
| Deep filter feeders | 0 | 0 | 0 | 0 | 0 | 0 | 0 | 0 | 0 | 0 | 0 | 0 | 0 | 0 | 0 | 0 | 0 | 0 | 0 | 0 | 0 | 0 | 0 | 0 | 0 | 0 | 0 | 0 | 0 | 0 |
| Gelatinous zooplankton | 0 | 0 | 0 | 0 | 0 | 0 | 0 | 0 | 0.2 | 0 | 0.055 | 0 | 0.2 | 0.105 | 0 | 0 | 0 | 0 | 0 | 0 | 0 | 0 | 0 | 0 | 0.031 | 0 | 0 | 0 | 0 | 0 |
| Large Zooplankton | 0 | 0.001 | 0 | 0 | 0 | 0 | 0 | 0.001 | 0.5 | 0 | 0 | 0.1 | 0.55 | 0.81 | 0 | 0.002 | 0 | 0 | 0 | 0.016 | 0.163 | 0.04 | 0.299 | 0.2 | 0 | 0.187 | 0 | 0 | 0.001 | 0 |
| Micro Zooplankton | 0 | 0 | 0.004 | 0 | 0 | 0 | 0 | 0 | 0 | 0 | 0 | 0 | 0 | 0.055 | 0 | 0 | 0 | 0 | 0 | 0.145 | 0 | 0.179 | 0 | 0.25 | 0 | 0.196 | 0.204 | 0 | 0 | 0.004 |
| Phytoplankton | 0 | 0 | 0 | 0 | 0 | 0 | 0 | 0 | 0 | 0 | 0 | 0 | 0 | 0.01 | 0 | 0 | 0 | 0 | 0 | 0.002 | 0 | 0.028 | 0 | 0 | 0 | 0.511 | 0.796 | 0 | 0 | 0 |
| Macroalgae | 0 | 0 | 0 | 0 | 0 | 0 | 0 | 0 | 0 | 0 | 0 | 0 | 0 | 0 | 0 | 0 | 0 | 0 | 0 | 0 | 0 | 0 | 0 | 0.5 | 0 | 0 | 0 | 0 | 0 | 0 |
| Detritus | 0 | 0 | 0.004 | 0 | 0 | 0.005 | 0 | 0.012 | 0 | 0.002 | 0 | 0 | 0 | 0 | 0 | 0 | 0.018 | 0 | 0 | 0.103 | 0 | 0.32 | 0 | 0.025 | 0.042 | 0.107 | 0 | 0 | 0 | 0.004 |

**Table S11:** Diet matrix for the Kerala Ecopath model - continued

| Group | Deep Sea Shrimps | Coastal Shrimps | Squids | Cuttlefishes | Octopus | Commercial molluscs | Benthic detritivore | Macrozobenthos (Benthic carnivore) | Benthic grazers | Shelf filter feeders | Deep filter feeders | Gelatinous zooplankton | Large Zooplankton | Micro Zooplankton |
|---|---|---|---|---|---|---|---|---|---|---|---|---|---|---|
| Marine Mammals | 0 | 0 | 0 | 0 | 0 | 0 | 0 | 0 | 0 | 0 | 0 | 0 | 0 | 0 |
| Sharks coastal | 0 | 0 | 0 | 0 | 0 | 0 | 0 | 0 | 0 | 0 | 0 | 0 | 0 | 0 |
| Sharks offshore | 0 | 0 | 0 | 0 | 0 | 0 | 0 | 0 | 0 | 0 | 0 | 0 | 0 | 0 |
| Sharks demersal | 0 | 0 | 0 | 0 | 0 | 0 | 0 | 0 | 0 | 0 | 0 | 0 | 0 | 0 |
| Guitarfishes & Rays | 0 | 0 | 0 | 0 | 0 | 0 | 0 | 0 | 0 | 0 | 0 | 0 | 0 | 0 |
| Large pelagics-offshore | 0 | 0 | 0 | 0 | 0 | 0 | 0 | 0 | 0 | 0 | 0 | 0 | 0 | 0 |
| Large Pelagics-inshore | 0 | 0 | 0 | 0 | 0 | 0 | 0 | 0 | 0 | 0 | 0 | 0 | 0 | 0 |
| Large Bentho Pelagics shelf | 0 | 0 | 0 | 0 | 0 | 0 | 0 | 0 | 0 | 0 | 0 | 0 | 0 | 0 |
| Large benthopelagic deep | 0 | 0 | 0 | 0 | 0 | 0 | 0 | 0 | 0 | 0 | 0 | 0 | 0 | 0 |
| Medium Bentho Pelagics shelf | 0 | 0 | 0 | 0 | 0 | 0 | 0 | 0 | 0 | 0 | 0 | 0 | 0 | 0 |
| Medium benthopelagic deep | 0 | 0 | 0 | 0 | 0 | 0 | 0 | 0 | 0 | 0 | 0 | 0 | 0 | 0 |
| Small Bentho Pelagics shelf | 0 | 0 | 0 | 0.03 | 0.1 | 0 | 0 | 0 | 0 | 0 | 0 | 0 | 0 | 0 |
| Small benthopelagic deep | 0 | 0 | 0 | 0 | 0 | 0 | 0 | 0 | 0 | 0 | 0 | 0 | 0 | 0 |
| Mesopelagics | 0 | 0 | 0.01 | 0 | 0 | 0 | 0 | 0 | 0 | 0 | 0 | 0 | 0 | 0 |
| Tunas coastal | 0 | 0 | 0 | 0 | 0 | 0 | 0 | 0 | 0 | 0 | 0 | 0 | 0 | 0 |
| Tunas offshore | 0 | 0 | 0 | 0 | 0 | 0 | 0 | 0 | 0 | 0 | 0 | 0 | 0 | 0 |
| Ribbonfishes | 0 | 0 | 0 | 0 | 0 | 0 | 0 | 0 | 0 | 0 | 0 | 0 | 0 | 0 |
| Large Benthic Carnivores shelf | 0 | 0 | 0.002 | 0 | 0 | 0 | 0 | 0 | 0 | 0 | 0 | 0 | 0 | 0 |
| Large Benthic Carnivores deep | 0 | 0 | 0.001 | 0 | 0 | 0 | 0 | 0 | 0 | 0 | 0 | 0 | 0 | 0 |
| Medium Benthic Carnivores shelf | 0 | 0 | 0.001 | 0.03 | 0 | 0 | 0 | 0 | 0 | 0 | 0 | 0 | 0 | 0 |
| Medium Benthic Carniv. deep | 0 | 0 | 0.022 | 0.05 | 0 | 0 | 0 | 0 | 0 | 0 | 0 | 0 | 0 | 0 |
| Small Benthic Carnivores shelf | 0 | 0 | 0.001 | 0.06 | 0.09 | 0 | 0 | 0 | 0 | 0 | 0 | 0 | 0 | 0 |
| Small Benthic Carnivores deep | 0 | 0 | 0.001 | 0.15 | 0.01 | 0 | 0 | 0 | 0 | 0 | 0 | 0 | 0 | 0 |
| Benthic omnivores shelf | 0 | 0 | 0 | 0 | 0 | 0 | 0 | 0 | 0 | 0 | 0 | 0 | 0 | 0 |
| Threadfin breams | 0 | 0 | 0.001 | 0 | 0 | 0 | 0 | 0 | 0 | 0 | 0 | 0 | 0 | 0 |
| Indian Mackerel | 0 | 0 | 0.026 | 0 | 0 | 0 | 0 | 0 | 0 | 0 | 0 | 0 | 0 | 0 |
| Oil Sardine | 0 | 0 | 0.224 | 0.05 | 0 | 0 | 0 | 0 | 0 | 0 | 0 | 0 | 0 | 0 |
| Other Clupeids inshore | 0 | 0 | 0.055 | 0.013 | 0 | 0 | 0 | 0 | 0 | 0 | 0 | 0 | 0 | 0 |
| Anchovies | 0 | 0 | 0.03 | 0.1 | 0 | 0 | 0 | 0 | 0 | 0 | 0 | 0 | 0 | 0 |
| Crabs & Lobsters | 0 | 0 | 0 | 0.002 | 0.045 | 0 | 0 | 0 | 0 | 0 | 0 | 0 | 0 | 0 |
| Deep Sea Shrimps | 0 | 0 | 0.005 | 0 | 0.1 | 0 | 0 | 0 | 0 | 0 | 0 | 0 | 0 | 0 |

| Group | Deep Sea Shrimps | Coastal Shrimps | Squids | Cuttlefishes | Octopus | Commercial molluscs | Benthic detritivore | Macrozobenthos (Benthic carnivore) | Benthic grazers | Shelf filter feeders | Deep filter feeders | Gelatinous zooplankton | Large Zooplankton | Micro Zooplankton |
|---|---|---|---|---|---|---|---|---|---|---|---|---|---|---|
| Coastal Shrimps | 0 | 0 | 0.04 | 0.07 | 0.1 | 0 | 0 | 0 | 0 | 0 | 0 | 0 | 0 | 0 |
| Squids | 0 | 0.01 | 0.048 | 0.045 | 0 | 0 | 0 | 0 | 0 | 0 | 0 | 0 | 0 | 0 |
| Cuttlefishes | 0 | 0 | 0 | 0.1 | 0 | 0 | 0 | 0 | 0 | 0 | 0 | 0 | 0 | 0 |
| Octopus | 0 | 0 | 0 | 0.002 | 0 | 0 | 0 | 0 | 0 | 0 | 0 | 0 | 0 | 0 |
| Commercial molluscs | 0 | 0 | 0 | 0 | 0 | 0.13 | 0 | 0.001 | 0 | 0 | 0 | 0 | 0 | 0 |
| Benthic detritivore | 0.11 | 0.667 | 0.057 | 0.121 | 0.051 | 0 | 0.03 | 0.5 | 0 | 0 | 0 | 0 | 0 | 0 |
| Macrozobenthos (Benthic carnivore) | 0 | 0.094 | 0.122 | 0.026 | 0.051 | 0 | 0 | 0.035 | 0 | 0 | 0 | 0 | 0 | 0 |
| Benthic grazers | 0.052 | 0 | 0 | 0.001 | 0.041 | 0 | 0 | 0 | 0 | 0 | 0 | 0 | 0 | 0 |
| Shelf filter feeders | 0 | 0 | 0 | 0.025 | 0.011 | 0 | 0 | 0.001 | 0 | 0 | 0 | 0 | 0 | 0 |
| Deep filter feeders | 0.05 | 0 | 0 | 0.025 | 0 | 0 | 0 | 0.001 | 0 | 0 | 0 | 0 | 0 | 0 |
| Gelatinous zooplankton | 0 | 0 | 0.072 | 0 | 0 | 0 | 0 | 0 | 0 | 0 | 0 | 0 | 0 | 0 |
| Large Zooplankton | 0.417 | 0.019 | 0.207 | 0.1 | 0.401 | 0.08 | 0.08 | 0.245 | 0 | 0 | 0 | 0.25 | 0 | 0 |
| Micro Zooplankton | 0.248 | 0.016 | 0.076 | 0 | 0 | 0.06 | 0.13 | 0.16 | 0 | 0 | 0 | 0.25 | 0.58 | 0 |
| Phytoplankton | 0 | 0.004 | 0 | 0 | 0 | 0.12 | 0 | 0.007 | 0.12 | 0.8 | 0.8 | 0.5 | 0.22 | 1 |
| Macroalgae | 0 | 0 | 0 | 0 | 0 | 0 | 0.01 | 0 | 0.62 | 0 | 0 | 0 | 0 | 0 |
| Detritus | 0.123 | 0.19 | 0 | 0 | 0 | 0.61 | 0.75 | 0.05 | 0.26 | 0.2 | 0.2 | 0 | 0.2 | 0 |

**Table S12:** Landed catch for the Kerala Ecopath model (units tkm$^{-2}$year$^{-1}$). The Fleet names come from the Indian catch database codes. No discards are included in this model.

| Group | MDTN | MGN | MHL | MOTHS | MPS | MRS | MTN | NM | OBBS | OBGN | OBHL | OBOTHS | OBRS | OBTN | OBSS |
|---|---|---|---|---|---|---|---|---|---|---|---|---|---|---|---|
| Marine Mammals | 0 | 0 | 0 | 0 | 0 | 0 | 0 | 0 | 0 | 0 | 0 | 0 | 0 | 0 | 0 |
| Sharks coastal | 1.14E-03 | 1.85E-03 | 1.03E-03 | 1.32E-02 | 0 | 4.83E-08 | 1.08E-04 | 5.19E-05 | 4.56E-07 | 9.76E-04 | 6.14E-05 | 1.81E-04 | 7.40E-06 | 9.55E-08 | 0 |
| Sharks offshore | 1.01E-03 | 2.03E-03 | 9.48E-04 | 1.02E-02 | 0 | 4.97E-07 | 4.16E-05 | 3.87E-05 | 0 | 9.94E-04 | 3.52E-04 | 6.54E-04 | 4.57E-06 | 0 | 0 |
| Sharks demersal | 1.29E-03 | 3.46E-04 | 4.10E-04 | 4.89E-03 | 0 | 3.81E-06 | 1.15E-04 | 2.11E-06 | 5.35E-07 | 6.77E-04 | 1.29E-05 | 9.49E-05 | 0 | 1.55E-07 | 0 |
| Guitarfishes & Rays | 4.08E-03 | 1.65E-03 | 4.60E-04 | 9.35E-03 | 0 | 2.54E-04 | 1.04E-03 | 1.80E-04 | 2.12E-05 | 6.10E-03 | 1.34E-03 | 4.02E-04 | 5.76E-06 | 1.64E-05 | 0 |
| Large pelagics-offshore | 1.76E-03 | 5.70E-03 | 2.57E-03 | 2.81E-02 | 2.39E-06 | 1.03E-04 | 6.62E-05 | 6.62E-06 | 6.63E-05 | 1.10E-02 | 7.39E-03 | 1.29E-03 | 4.19E-04 | 4.66E-07 | 0 |
| Large Pelagics-inshore | 2.74E-02 | 2.02E-03 | 2.12E-03 | 1.75E-02 | 1.81E-04 | 4.27E-03 | 1.24E-03 | 1.08E-04 | 2.51E-03 | 4.25E-02 | 1.47E-02 | 1.43E-03 | 1.88E-03 | 5.14E-05 | 1.55E-06 |
| Large Bentho Pelagics shelf | 5.13E-03 | 3.83E-04 | 4.39E-04 | 2.79E-03 | 1.72E-04 | 1.52E-03 | 1.10E-04 | 5.69E-05 | 2.96E-04 | 1.38E-03 | 1.81E-03 | 4.46E-05 | 1.02E-04 | 1.71E-06 | 0 |
| Large benthopelagic deep | 2.58E-05 | 0 | 0 | 4.99E-07 | 0 | 0 | 0 | 0 | 0 | 3.74E-07 | 0 | 1.21E-06 | 0 | 0 | 0 |

| **Group** | MDTN | MGN | MHL | MOTHS | MPS | MRS | MTN | NM | OBBS | OBGN | OBHL | OBOTHS | OBRS | OBTN | OBSS |
|---|---|---|---|---|---|---|---|---|---|---|---|---|---|---|---|
| Medium Bentho Pelagics shelf | 2.62E-01 | 9.72E-04 | 3.87E-04 | 2.71E-02 | 5.42E-03 | 7.11E-02 | 8.30E-03 | 4.26E-03 | 3.24E-02 | 2.44E-02 | 1.31E-02 | 3.40E-04 | 4.31E-02 | 2.62E-04 | 2.48E-03 |
| Medium benthopelagic deep | 8.55E-05 | 8.51E-06 | 1.67E-06 | 1.56E-05 | 0 | 3.61E-06 | 1.12E-05 | 1.05E-07 | 1.99E-06 | 1.28E-05 | 5.19E-05 | 0 | 8.69E-05 | 0 | 0 |
| Small Bentho Pelagics shelf | 1.59E-02 | 2.14E-04 | 5.27E-06 | 1.25E-03 | 1.32E-04 | 6.00E-03 | 1.35E-03 | 4.36E-04 | 6.72E-03 | 2.75E-03 | 1.32E-03 | 6.49E-05 | 3.14E-02 | 2.85E-04 | 1.94E-06 |
| Small benthopelagic deep | 9.12E-05 | 0 | 6.11E-07 | 4.22E-07 | 0 | 0 | 0 | 4.55E-07 | 0 | 0 | 0 | 0 | 0 | 0 | 0 |
| Mesopelagics | 6.73E-06 | 0 | 0 | 0 | 0 | 0 | 0 | 0 | 0 | 0 | 0 | 0 | 0 | 0 | 0 |
| Tunas coastal | 2.03E-03 | 2.63E-03 | 1.90E-03 | 1.32E-02 | 9.88E-04 | 1.15E-02 | 2.58E-05 | 5.92E-05 | 1.48E-03 | 5.90E-02 | 4.92E-02 | 1.10E-03 | 8.50E-04 | 6.29E-05 | 3.13E-04 |
| Tunas offshore | 1.01E-03 | 7.38E-03 | 2.45E-03 | 3.51E-02 | 1.62E-05 | 1.61E-04 | 8.74E-06 | 0 | 0 | 9.62E-03 | 4.90E-03 | 1.45E-03 | 5.39E-05 | 0 | 0 |
| Ribbonfishes | 1.46E-01 | 3.81E-04 | 4.21E-03 | 1.99E-02 | 1.70E-06 | 6.60E-04 | 5.80E-03 | 4.77E-05 | 5.25E-03 | 1.01E-02 | 2.14E-03 | 1.94E-04 | 4.78E-04 | 2.07E-04 | 2.68E-04 |
| Large Benthic Carnivores shelf | 1.51E-01 | 1.93E-03 | 1.76E-03 | 1.78E-02 | 8.64E-05 | 2.83E-03 | 5.62E-03 | 1.07E-03 | 2.57E-03 | 1.81E-02 | 5.30E-03 | 9.24E-05 | 4.91E-03 | 7.30E-05 | 2.81E-04 |
| Large Benthic Carnivores deep | 3.57E-03 | 2.39E-04 | 9.60E-04 | 4.86E-03 | 1.84E-08 | 1.43E-07 | 2.03E-04 | 5.72E-06 | 1.70E-05 | 1.84E-03 | 1.34E-03 | 3.98E-05 | 2.57E-07 | 5.75E-07 | 0 |
| Medium Benthic Carnivores shelf | 1.25E-01 | 1.03E-03 | 2.62E-04 | 6.56E-03 | 1.62E-03 | 9.05E-03 | 1.51E-02 | 5.46E-03 | 1.19E-02 | 1.81E-02 | 3.99E-03 | 8.92E-05 | 2.18E-02 | 2.39E-03 | 0 |
| Medium Benthic Carniv. deep | 1.29E-03 | 4.66E-07 | 1.65E-06 | 9.12E-05 | 0 | 0 | 1.00E-05 | 0 | 0 | 6.17E-06 | 4.06E-06 | 6.57E-06 | 0 | 0 | 0 |
| Small Benthic Carnivores shelf | 9.10E-02 | 4.13E-06 | 5.13E-06 | 3.58E-03 | 1.60E-05 | 6.82E-03 | 7.74E-02 | 1.17E-02 | 4.71E-03 | 1.70E-02 | 1.57E-04 | 2.29E-06 | 1.64E-02 | 8.59E-03 | 5.06E-04 |
| Small Benthic Carnivores deep | 8.17E-04 | 1.17E-05 | 1.83E-06 | 3.21E-05 | 0 | 3.73E-07 | 1.99E-07 | 0 | 0 | 2.23E-05 | 0 | 0 | 0 | 0 | 0 |
| Benthic omnivores shelf | 1.74E-04 | 9.82E-06 | 1.15E-05 | 1.53E-05 | 0 | 2.28E-04 | 9.00E-05 | 3.05E-04 | 3.33E-05 | 2.21E-04 | 2.42E-05 | 0 | 3.53E-04 | 5.94E-07 | 0 |
| Threadfin breams | 4.28E-01 | 1.02E-05 | 4.09E-06 | 2.99E-02 | 0 | 9.08E-05 | 8.11E-03 | 1.10E-04 | 1.14E-03 | 3.01E-03 | 1.65E-03 | 5.67E-07 | 4.33E-05 | 9.95E-06 | 0 |
| Indian Mackerel | 9.37E-02 | 6.34E-05 | 5.98E-08 | 1.25E-02 | 2.26E-02 | 2.26E-01 | 3.93E-03 | 6.17E-03 | 6.86E-03 | 1.54E-01 | 8.02E-03 | 2.10E-04 | 9.14E-02 | 5.71E-05 | 8.39E-04 |
| Oil Sardine | 2.30E-02 | 7.30E-06 | 0 | 3.67E-03 | 6.49E-03 | 1.18E+00 | 1.55E-02 | 3.09E-02 | 6.11E-03 | 5.59E-02 | 0 | 1.75E-06 | 1.00E+00 | 7.20E-05 | 0 |
| Other Clupeids inshore | 3.33E-02 | 2.38E-04 | 0 | 1.21E-03 | 2.38E-04 | 8.35E-02 | 1.23E-02 | 2.47E-02 | 2.88E-02 | 3.39E-02 | 2.28E-05 | 5.73E-06 | 9.11E-02 | 1.47E-03 | 1.52E-03 |
| Anchovies | 5.02E-02 | 5.24E-06 | 0 | 1.16E-03 | 8.98E-08 | 7.56E-02 | 8.16E-03 | 4.79E-03 | 2.09E-02 | 9.16E-04 | 0 | 1.80E-06 | 2.79E-01 | 1.11E-03 | 2.65E-03 |
| Crabs & Lobsters | 2.22E-02 | 0 | 0 | 3.35E-04 | 2.64E-07 | 3.47E-06 | 1.19E-02 | 2.43E-03 | 1.97E-04 | 1.56E-03 | 0 | 7.05E-06 | 8.47E-05 | 2.57E-03 | 0 |
| Deep Sea Shrimps | 9.39E-02 | 0 | 0 | 1.43E-03 | 0 | 0 | 1.74E-04 | 0 | 0 | 0 | 0 | 0 | 0 | 2.39E-05 | 0 |
| Coastal Shrimps | 1.78E-01 | 0 | 0 | 1.13E-03 | 3.39E-05 | 3.01E-02 | 9.19E-02 | 3.51E-03 | 5.04E-03 | 1.56E-03 | 0 | 6.58E-06 | 4.94E-02 | 3.28E-02 | 0 |
| Squids | 1.47E-01 | 6.18E-05 | 1.60E-03 | 1.11E-02 | 1.38E-05 | 8.65E-04 | 8.20E-03 | 6.86E-04 | 1.73E-02 | 1.53E-03 | 7.57E-03 | 6.11E-05 | 1.09E-03 | 1.54E-03 | 1.42E-04 |
| Cuttlefishes | 1.44E-01 | 6.12E-05 | 5.43E-03 | 1.59E-02 | 5.14E-07 | 2.65E-05 | 2.46E-03 | 1.27E-04 | 1.37E-05 | 2.54E-05 | 3.69E-02 | 3.42E-05 | 1.62E-03 | 8.59E-04 | 0 |
| Octopus | 4.21E-02 | 1.14E-05 | 1.04E-06 | 1.77E-03 | 0 | 0 | 1.04E-03 | 3.15E-06 | 2.59E-07 | 1.78E-07 | 2.57E-05 | 0 | 1.44E-07 | 0 | 0 |
| Commercial molluscs | 2.53E-03 | 0 | 0 | 2.53E-08 | 0 | 0 | 5.71E-03 | 1.21E-04 | 0 | 1.85E-04 | 0 | 6.05E-06 | 0 | 0 | 0 |
| Benthic detritivore | 0 | 0 | 0 | 0 | 0 | 0 | 0 | 0 | 0 | 0 | 0 | 0 | 0 | 0 | 0 |
| Macrozobenthos (Benthic carnivore) | 3.21E-03 | 0 | 0 | 2.30E-08 | 1.86E-07 | 6.01E-07 | 1.82E-02 | 3.22E-07 | 0 | 4.03E-07 | 0 | 0 | 4.73E-05 | 3.97E-03 | 0 |
| Benthic grazers | 1.49E-06 | 0 | 0 | 0 | 0 | 0 | 4.35E-07 | 0 | 0 | 0 | 0 | 0 | 0 | 0 | 0 |
| Shelf filter feeders | 0 | 0 | 0 | 0 | 0 | 0 | 0 | 0 | 0 | 0 | 0 | 0 | 0 | 0 | 0 |
| Deep filter feeders | 0 | 0 | 0 | 0 | 0 | 0 | 0 | 0 | 0 | 0 | 0 | 0 | 0 | 0 | 0 |
| Gelatinous zooplankton | 0 | 0 | 0 | 0 | 0 | 0 | 0 | 0 | 0 | 0 | 0 | 0 | 0 | 0 | 0 |
| Large Zooplankton | 0 | 0 | 0 | 0 | 0 | 3.31E-05 | 4.78E-05 | 6.42E-06 | 0 | 0 | 0 | 0 | 1.93E-05 | 6.31E-06 | 0 |
| Micro Zooplankton | 0 | 0 | 0 | 0 | 0 | 0 | 0 | 0 | 0 | 0 | 0 | 0 | 0 | 0 | 0 |
| Phytoplankton | 0 | 0 | 0 | 0 | 0 | 0 | 0 | 0 | 0 | 0 | 0 | 0 | 0 | 0 | 0 |
| Macroalgae | 0 | 0 | 0 | 0 | 0 | 0 | 0 | 0 | 0 | 0 | 0 | 0 | 0 | 0 | 0 |
| Detritus | 0 | 0 | 0 | 0 | 0 | 0 | 0 | 0 | 0 | 0 | 0 | 0 | 0 | 0 | 0 |

**Table S13:** Ecopath base parameters for the north central Chilean Ecopath model

| Group | Trophic Level | Biomass ($tkm^{-2}$) | Production / Biomass ($year^{-1}$) | Consumption / Biomass ($year^{-1}$) | Ecotrophic Efficiency |
|---|---|---|---|---|---|
| Phytoplankton | 1 | 302 | 120 | | 0.594 |
| Zooplankton | 2.25 | 45.229 | 242.5 | 956 | 0.99 |
| Euphausiids | 2.688 | 112.295 | 13 | 31.707 | 0.99 |
| Small fish | 2.448 | 43.557 | 2.88 | 14.2 | 0.95 |
| Anchovy | 2.442 | 12.6 | 4.799 | 12.04 | 0.95 |
| Sardine | 3.369 | 1.046 | 2.14 | 14.2 | 0.81 |
| Horse mackerel | 3.385 | 20.441 | 0.53 | 10.8 | 0.296 |
| Jack mackerel | 3.66 | 99.459 | 0.17 | 14.2 | 0.189 |
| Hake | 3.231 | 10.092 | 1.55 | 5.159 | 0.749 |
| Sea lions | 3.897 | 0.144 | 0.25 | 20 | 0 |
| Marine birds | 3.703 | 0.065 | 0.5 | 20 | 0 |
| Jumbo squid | 3.434 | 7.45 | 3 | 8.571 | 0.535 |
| Demersal crustaceans | 2 | 4.443 | 4 | 14.104 | 0.958 |
| Sword fish | 3.815 | 0.019 | 0.5 | 5 | 0.95 |
| Sharks | 4.046 | 0.008 | 0.362 | 2.413 | 0.95 |
| Detritus | 1 | 1 | | | 0.614 |

**Table S14:** Diet matrix for the north central Chilean Ecopath model

| Group | Zooplankton | Euphausiids | Small fish | Anchovy | Sardine | Horse mackerel | Jack mackerel | Hake | Sea lions | Marine birds | Jumbo squid | Demersal crustaceans | Sword fish | Sharks |
|---|---|---|---|---|---|---|---|---|---|---|---|---|---|---|
| Phytoplankton | 0.45 | 0.45 | 0.6 | 0.6 | 0.05 | 0 | 0 | 0 | 0 | 0 | 0 | 0 | 0 | 0 |
| Zooplankton | 0.2 | 0.55 | 0.1 | 0.1 | 0.535 | 0.63 | 0.02 | 0 | 0 | 0 | 0 | 0 | 0 | 0 |
| Euphausiids | 0 | 0 | 0.14 | 0.14 | 0.415 | 0.26 | 0.9 | 0.053 | 0 | 0 | 0 | 0 | 0 | 0 |
| Small fish | 0 | 0 | 0 | 0 | 0 | 0.028 | 0.08 | 0 | 0 | 0 | 0 | 0 | 0 | 0.22 |
| Anchovy | 0 | 0 | 0.015 | 0.015 | 0 | 0.082 | 0 | 0.145 | 0.155 | 0.682 | 0.198 | 0 | 0 | 0 |
| Sardine | 0 | 0 | 0 | 0 | 0 | 0 | 0 | 0.015 | 0.015 | 0.076 | 0.014 | 0 | 0 | 0 |
| Horse mackerel | 0 | 0 | 0 | 0 | 0 | 0 | 0 | 0.009 | 0.06 | 0 | 0.009 | 0 | 0 | 0.29 |
| Jack mackerel | 0 | 0 | 0 | 0 | 0 | 0 | 0 | 0.009 | 0.06 | 0 | 0.009 | 0 | 0 | 0 |
| Hake | 0 | 0 | 0 | 0 | 0 | 0 | 0 | 0.085 | 0.18 | 0.242 | 0.1 | 0 | 0.45 | 0 |
| Sea lions | 0 | 0 | 0 | 0 | 0 | 0 | 0 | 0 | 0 | 0 | 0 | 0 | 0 | 0 |
| Marine birds | 0 | 0 | 0 | 0 | 0 | 0 | 0 | 0 | 0 | 0 | 0 | 0 | 0 | 0 |
| Jumbo squid | 0 | 0 | 0 | 0 | 0 | 0 | 0 | 0 | 0 | 0 | 0.15 | 0 | 0 | 0.305 |
| Demersal crustaceans | 0 | 0 | 0 | 0 | 0 | 0 | 0 | 0.23 | 0.08 | 0 | 0.068 | 0 | 0.23 | 0.1 |
| Sword fish | 0 | 0 | 0 | 0 | 0 | 0 | 0 | 0 | 0 | 0 | 0 | 0 | 0 | 0.025 |
| Sharks | 0 | 0 | 0 | 0 | 0 | 0 | 0 | 0 | 0 | 0 | 0 | 0 | 0 | 0 |
| Detritus | 0.35 | 0 | 0 | 0 | 0 | 0 | 0 | 0 | 0 | 0 | 0 | 1 | 0 | 0 |
| Import | 0 | 0 | 0.145 | 0.145 | 0 | 0 | 0 | 0.456 | 0.45 | 0 | 0.453 | 0 | 0.32 | 0.06 |

**Table S15:** Landed catch for the north central Chilean Ecopath model (units tkm$^{-2}$year$^{-1}$). No discards are included in this model.

| Group | Purse seine | Trawler | Longline | Small longline | Squider |
|---|---|---|---|---|---|
| Phytoplankton | 0 | 0 | 0 | 0 | 0 |
| Zooplankton | 0 | 0 | 0 | 0 | 0 |
| Euphausiids | 0 | 0 | 0 | 0 | 0 |
| Small fish | 0 | 0 | 0 | 0 | 0 |
| Anchovy | 6.26E+00 | 0 | 0 | 0 | 0 |
| Sardine | 1.05E-02 | 0 | 0 | 0 | 0 |
| Horse mackerel | 2.04E+00 | 0 | 0 | 0 | 0 |
| Jack mackerel | 2.04E+00 | 0 | 0 | 0 | 0 |
| Hake | 0 | 0 | 0 | 2.13E-02 | 0 |
| Sea lions | 0 | 0 | 0 | 0 | 0 |
| Marine birds | 0 | 0 | 0 | 0 | 0 |
| Jumbo squid | 0 | 0 | 0 | 0 | 2.37E+00 |
| Demersal crustaceans | 0 | 4.55E-01 | 0 | 0 | 0 |
| Sword fish | 0 | 0 | 8.75E-03 | 0 | 0 |
| Sharks | 0 | 0 | 2.66E-03 | 0 | 0 |
| Detritus | 0 | 0 | 0 | 0 | 0 |
| Import | 0 | 0 | 0 | 0 | 0 |

**Figure S2:** Catch time series (in tonnes) for historical baseline run of the East Bering Sea EwE model

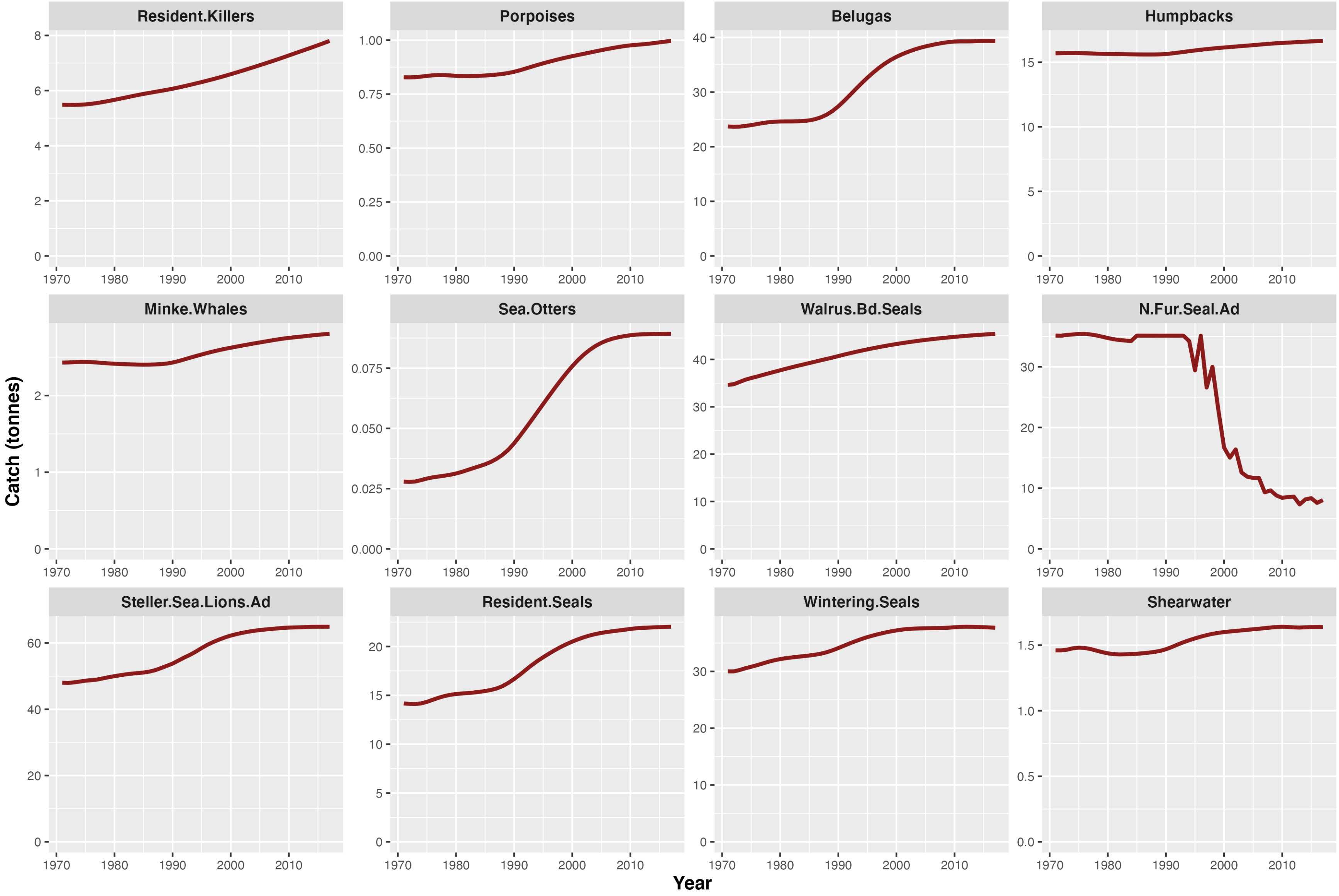

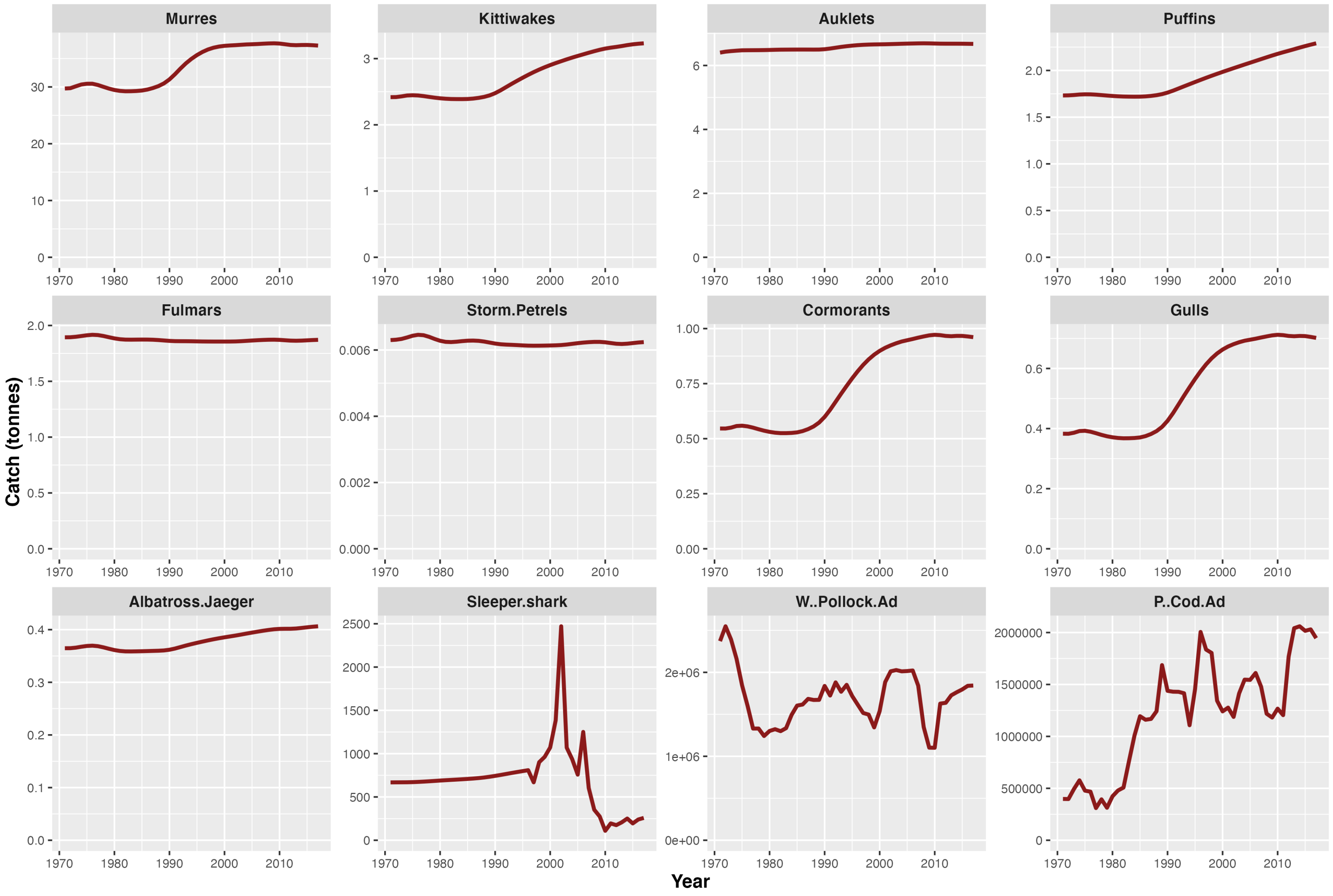

Murres
Kittiwakes
Auklets
Puffins
Fulmars
Storm.Petrels
Cormorants
Gulls
Albatross.Jaeger
Sleeper.shark
W..Pollock.Ad
P..Cod.Ad
Catch (tonnes)
Year

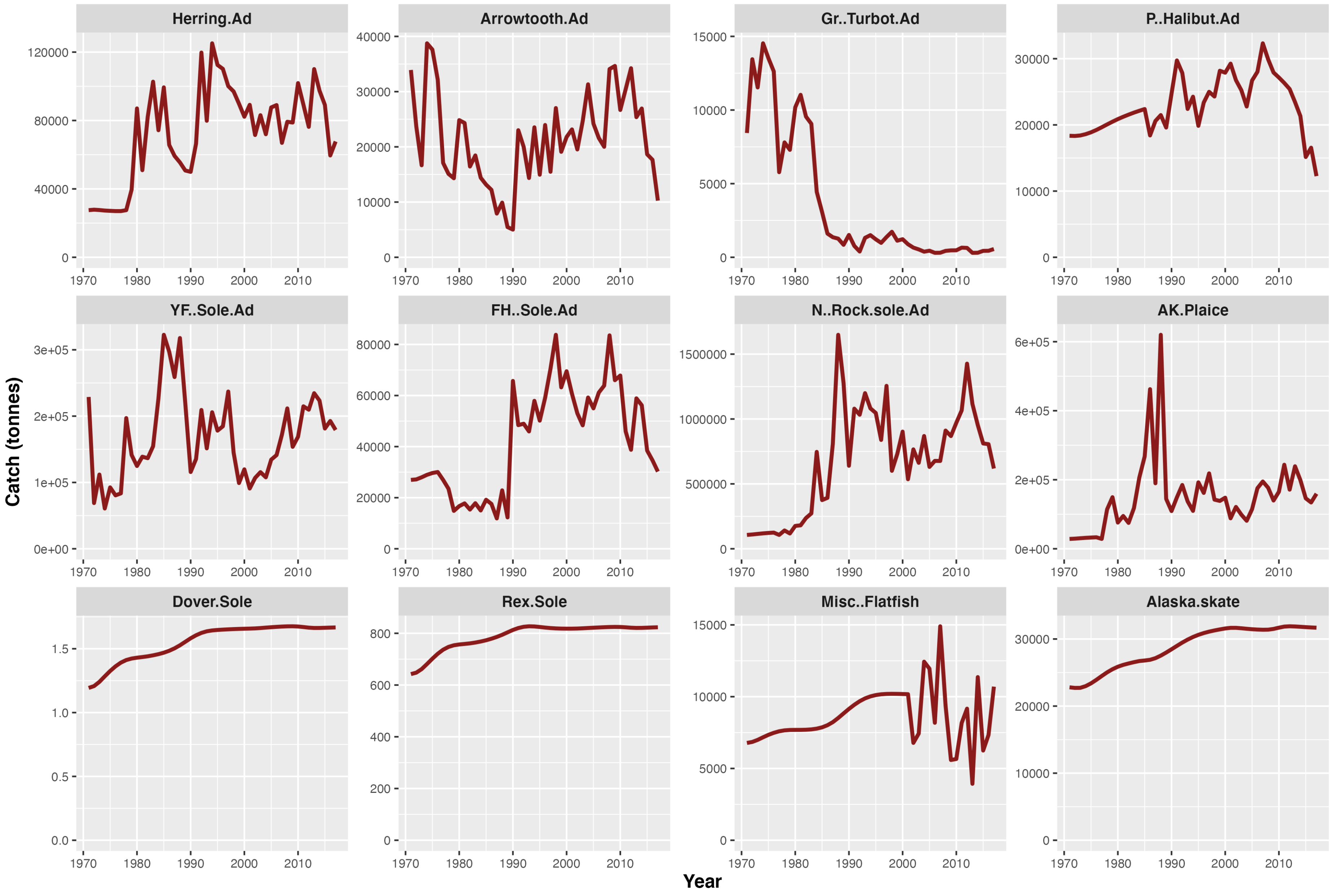
Herring.Ad
Arrowtooth.Ad
Gr..Turbot.Ad
P..Halibut.Ad
YF..Sole.Ad
FH..Sole.Ad
N..Rock.sole.Ad
AK.Plaice
Dover.Sole
Rex.Sole
Misc..Flatfish
Alaska.skate
Catch (tonnes)
Year

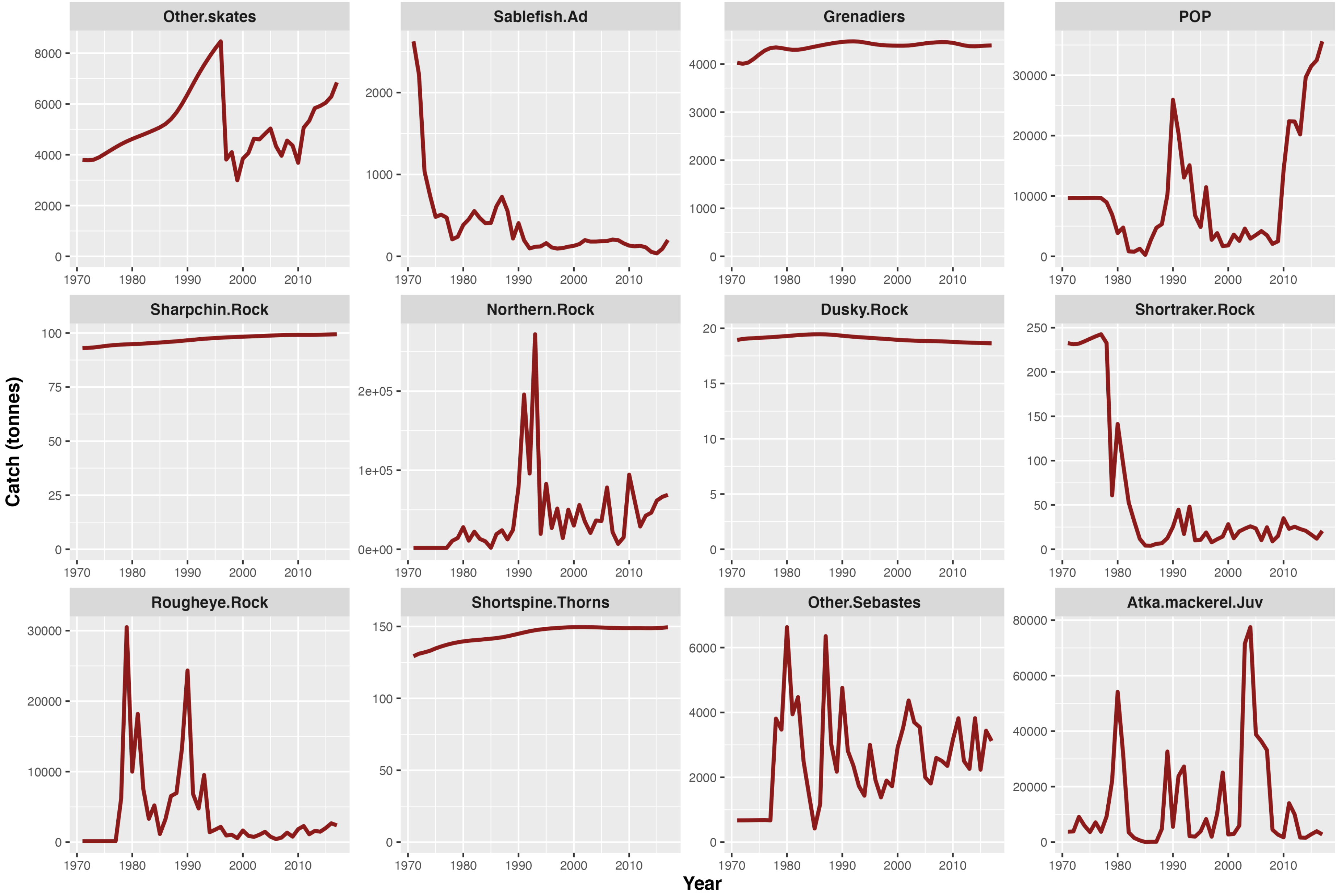
Other.skates
Sablefish.Ad
Grenadiers
POP
Sharpchin.Rock
Northern.Rock
Dusky.Rock
Shortraker.Rock
Rougheye.Rock
Shortspine.Thorns
Other.Sebastes
Atka.mackerel.Juv
Catch (tonnes)
Year

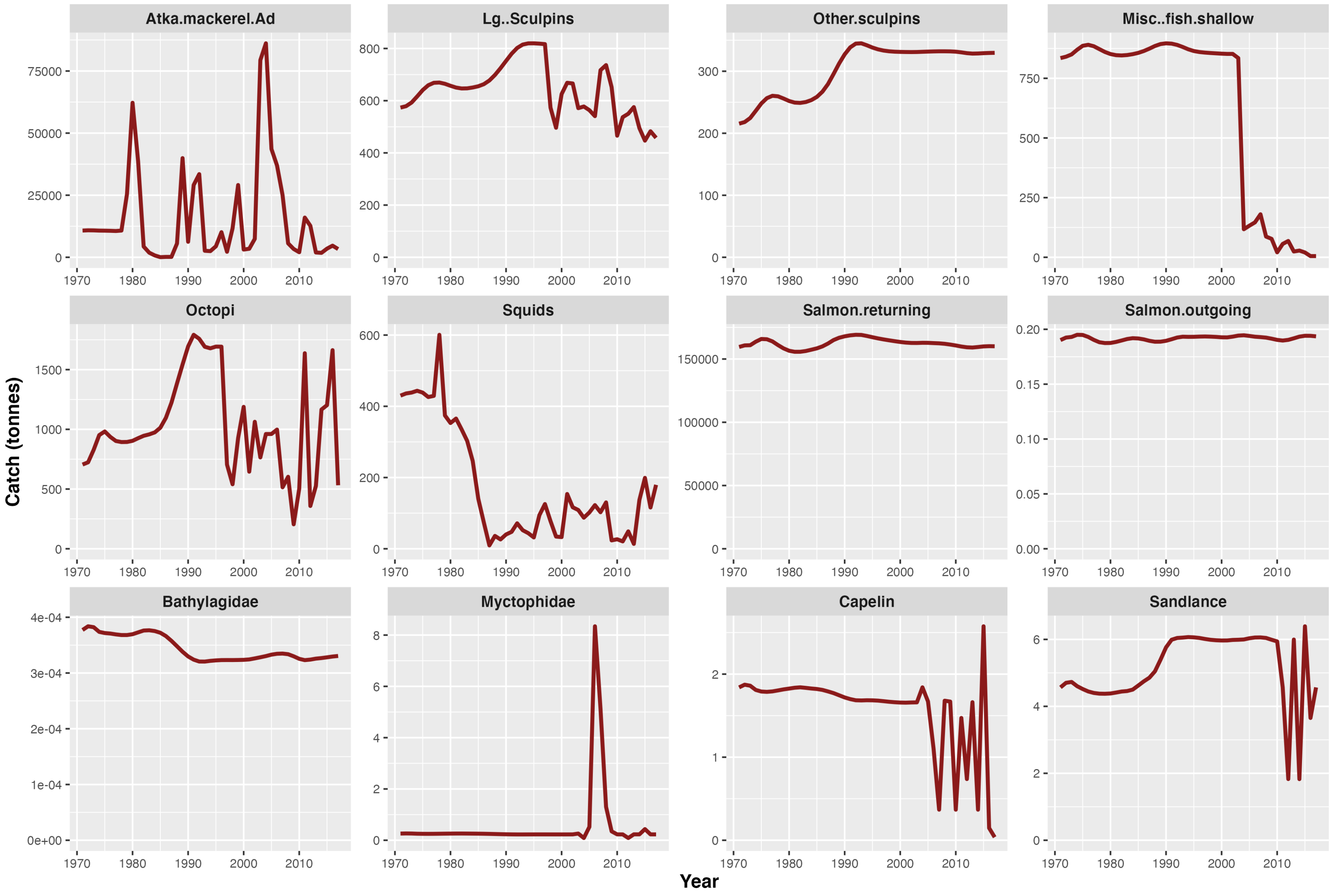
Atka.mackerel.Ad
Lg..Sculpins
Other.sculpins
Misc..fish.shallow
Octopi
Squids
Salmon.returning
Salmon.outgoing
Bathylagidae
Myctophidae
Capelin
Sandlance
Catch (tonnes)
Year

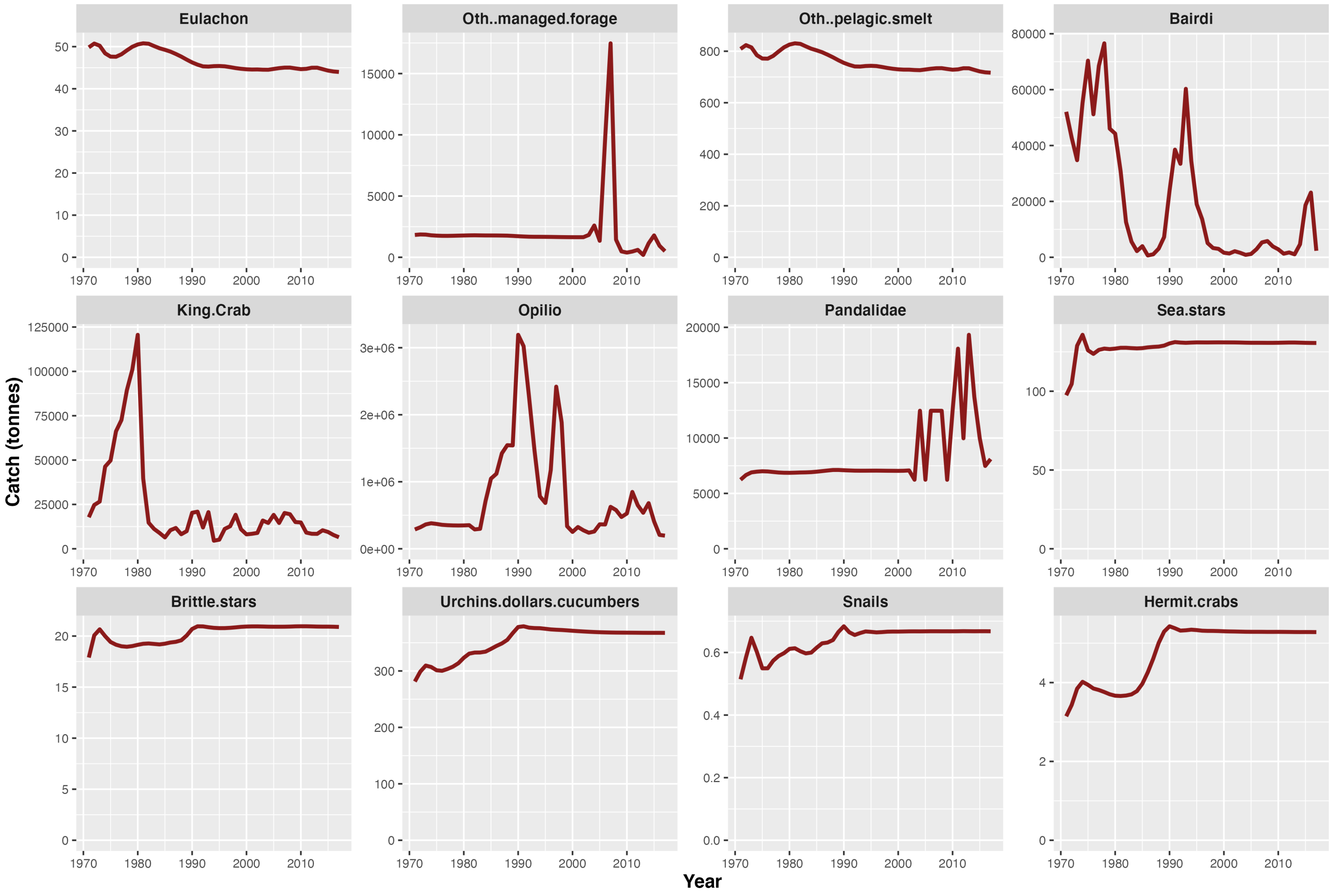

Eulachon
Oth..managed.forage
Oth..pelagic.smelt
Bairdi
King.Crab
Opilio
Pandalidae
Sea.stars
Brittle.stars
Urchins.dollars.cucumbers
Snails
Hermit.crabs
Catch (tonnes)
Year

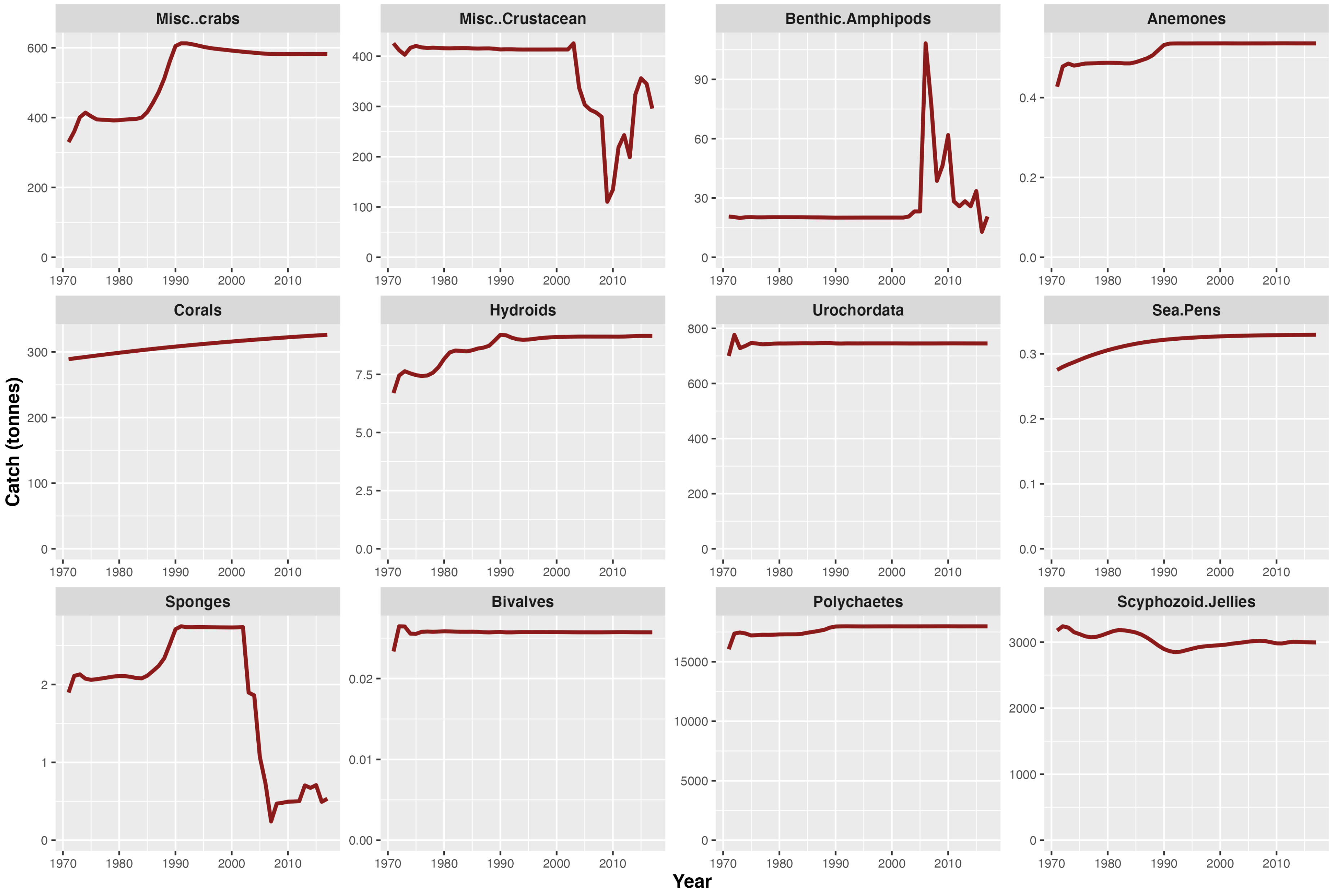
Misc..crabs
Misc..Crustacean
Benthic.Amphipods
Anemones
Corals
Hydroids
Urochordata
Sea.Pens
Sponges
Bivalves
Polychaetes
Scyphozoid.Jellies
Catch (tonnes)
Year
1970
1980
1990
2000
2010

**Figure S3:** Catch time series for the historical baseline run of the SE Australian (eastern Bass Strait) Ecopath model.

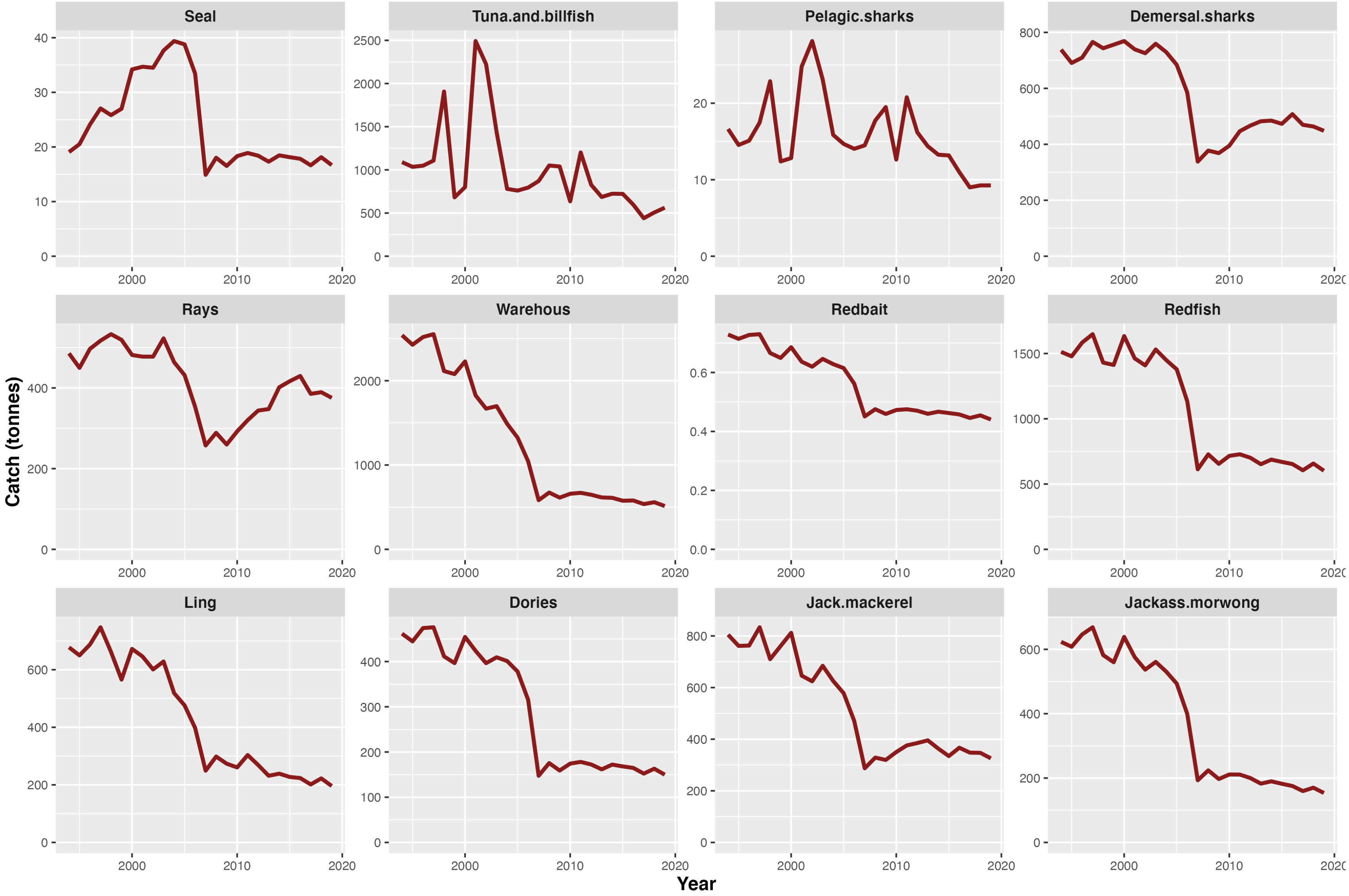

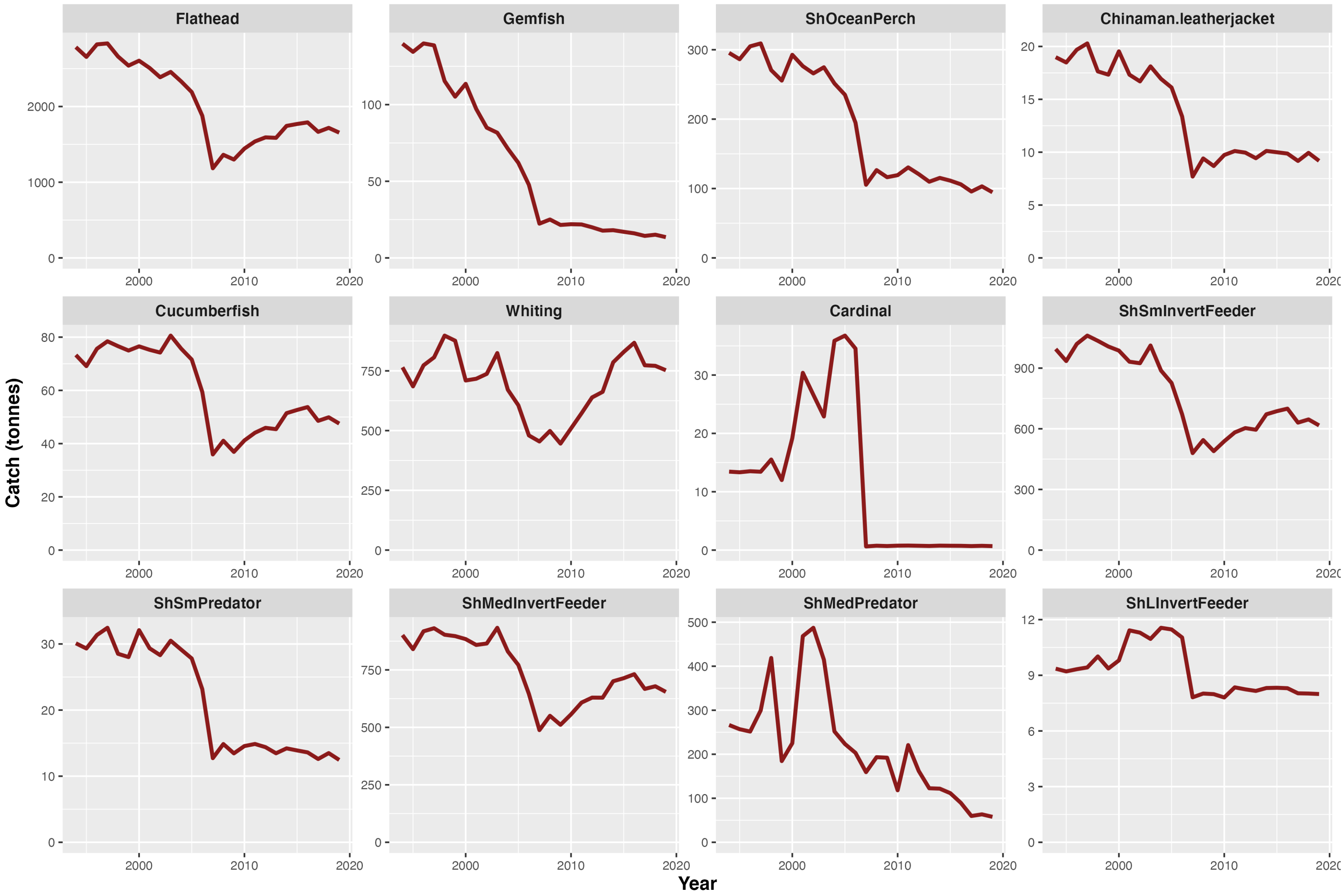

Flathead
Gemfish
ShOceanPerch
Chinaman.leatherjacket
Cucumberfish
Whiting
Cardinal
ShSmInvertFeeder
ShSmPredator
ShMedInvertFeeder
ShMedPredator
ShLInvertFeeder
Catch (tonnes)
Year

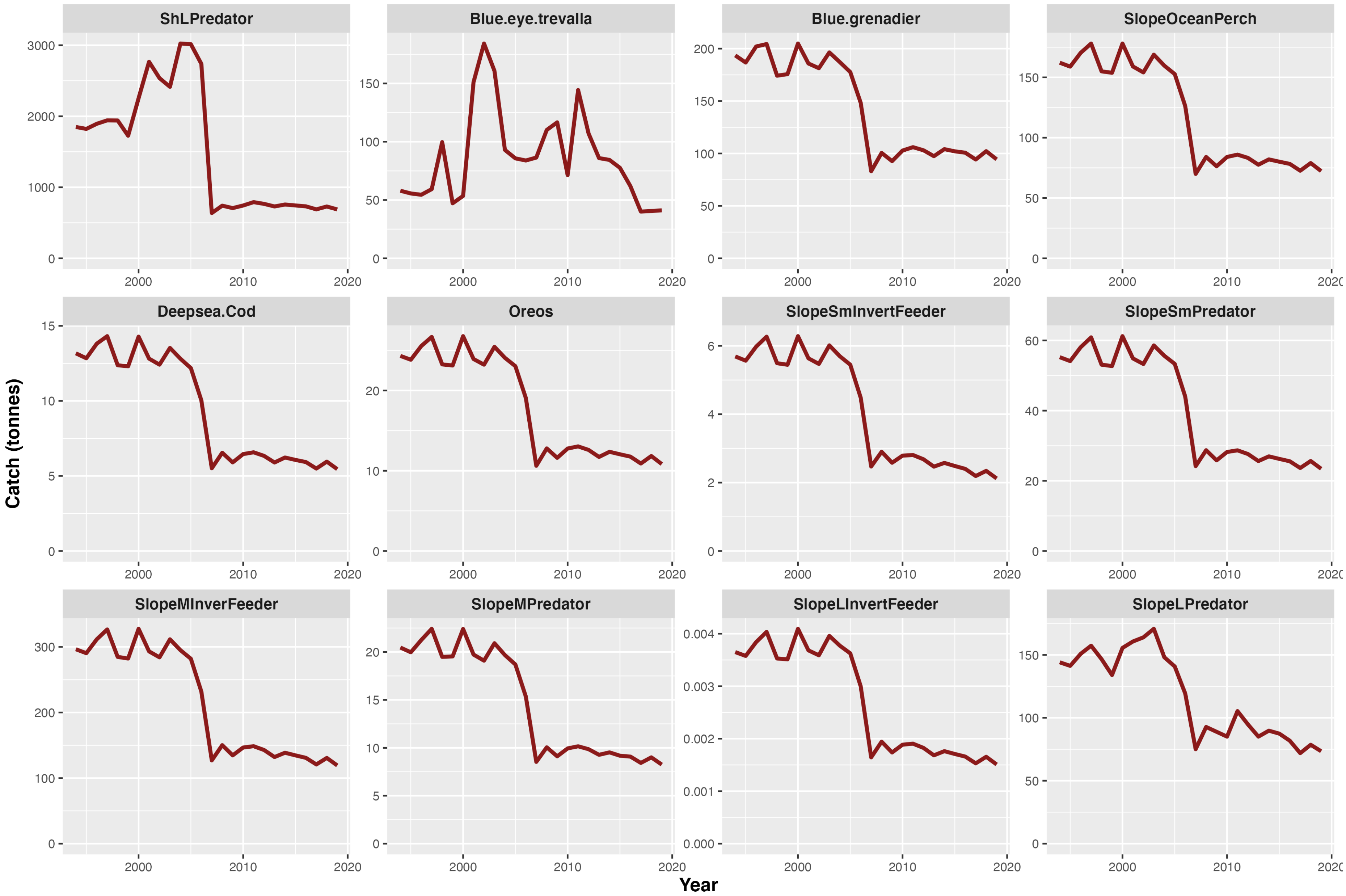

ShLPredator
Blue.eye.trevalla
Blue.grenadier
SlopeOceanPerch
Deepsea.Cod
Oreos
SlopeSmInvertFeeder
SlopeSmPredator
SlopeMInverFeeder
SlopeMPredator
SlopeLInvertFeeder
SlopeLPredator
Catch (tonnes)
Year

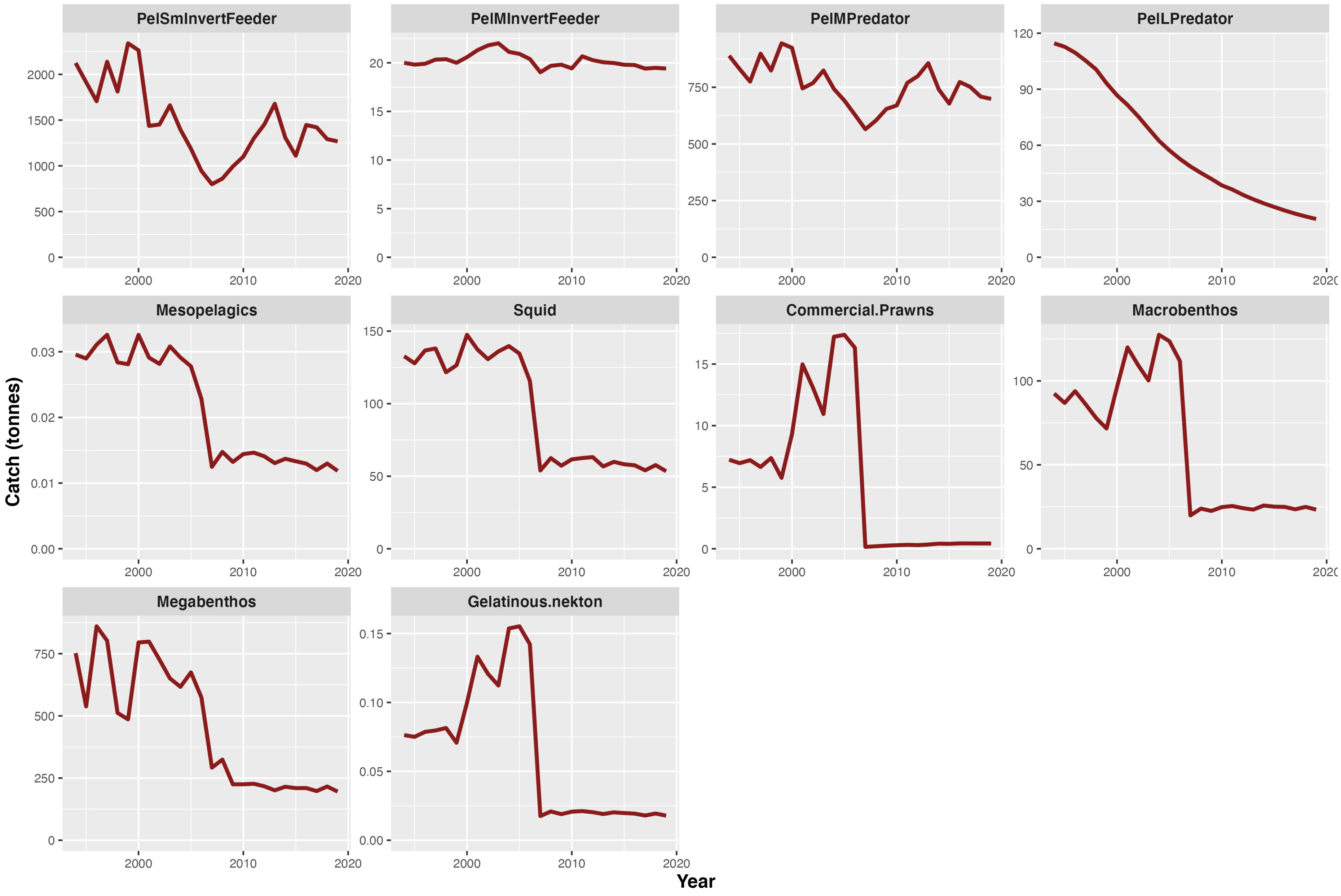

PelSmInvertFeeder
PelMInvertFeeder
PelMPredator
PelLPredator
Mesopelagics
Squid
Commercial.Prawns
Macrobenthos
Megabenthos
Gelatinous.nekton
Catch (tonnes)
Year

**Figure S4:** Catch time series for the historical baseline run of the Kerala EwE model.

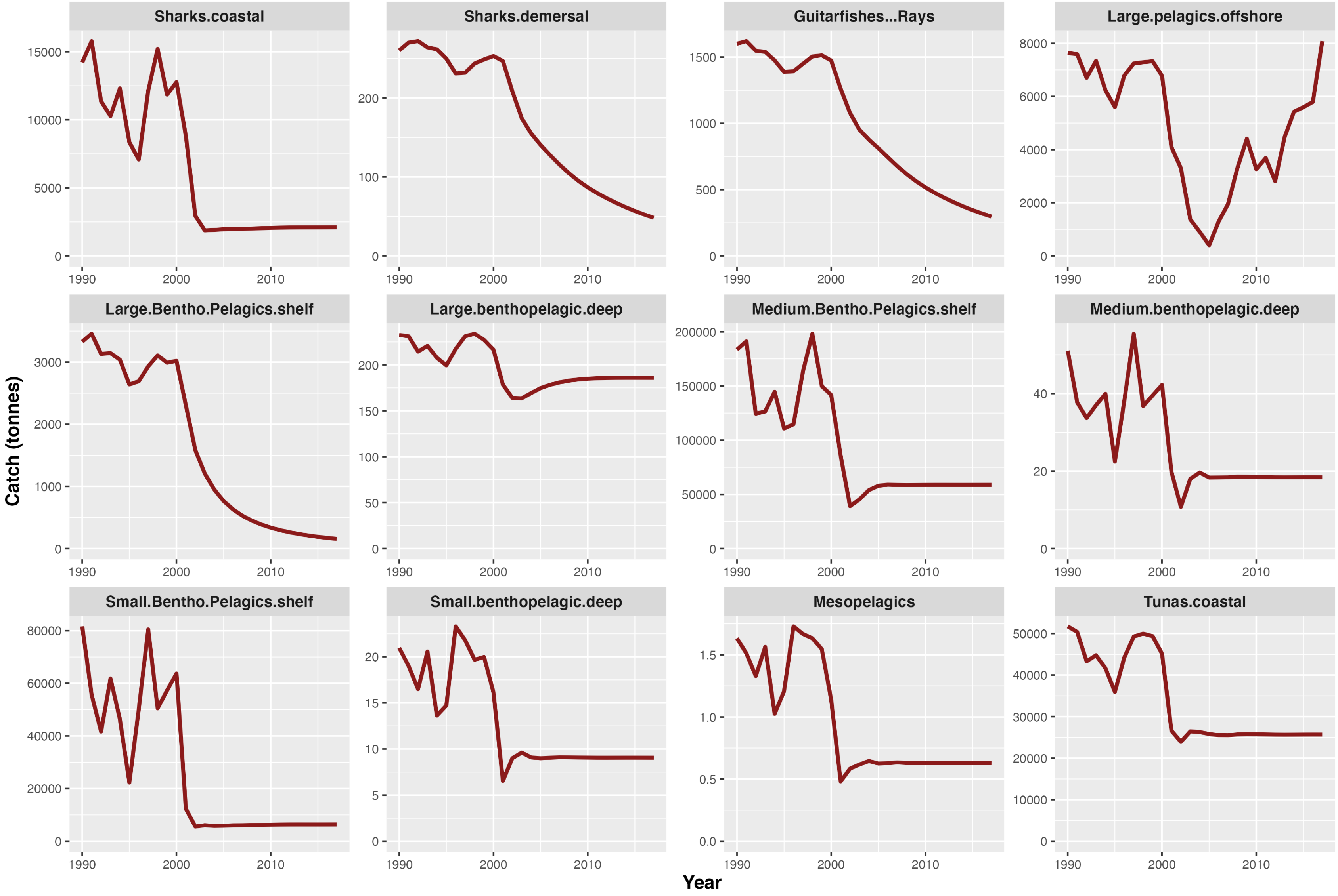

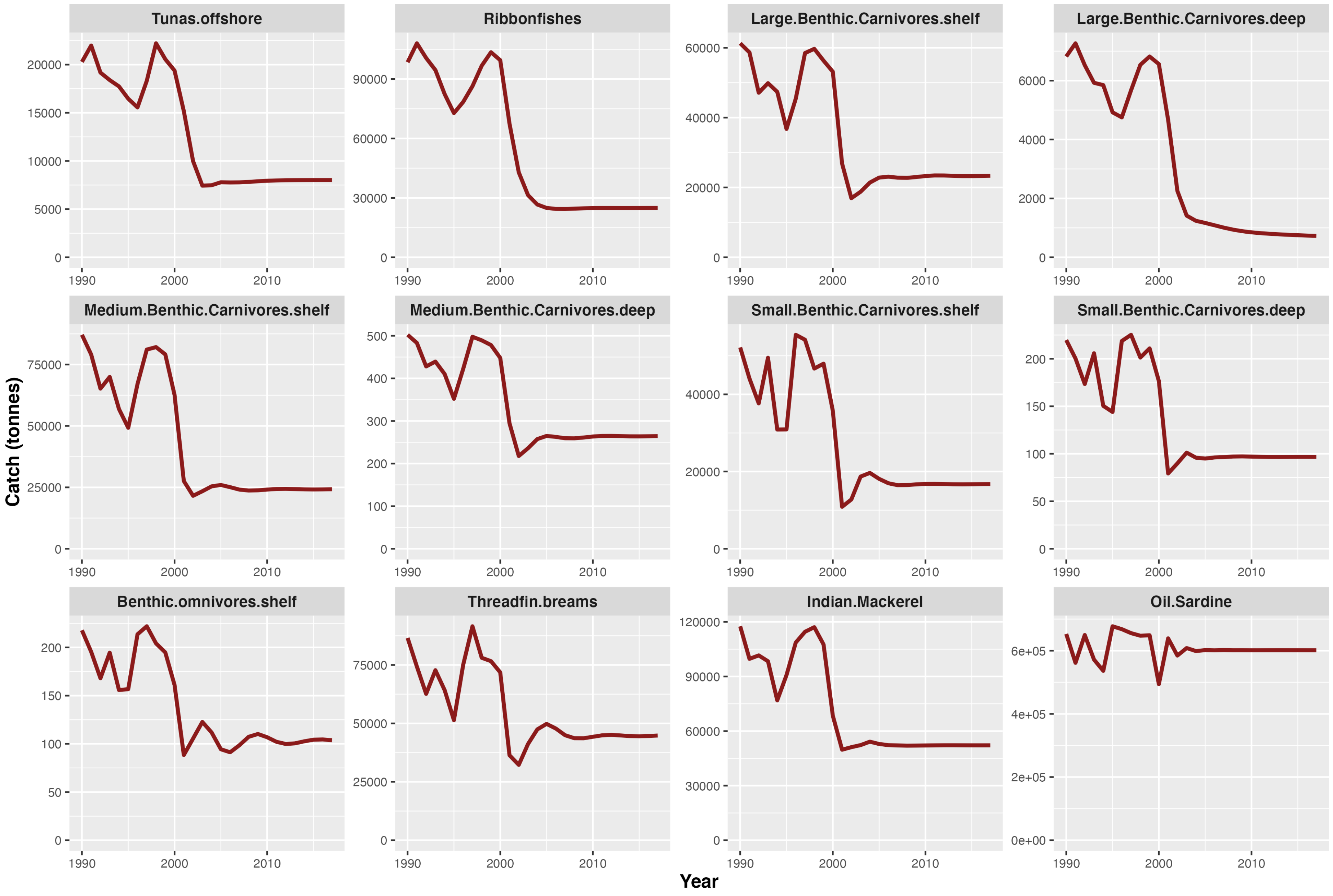

Tunas.offshore
Ribbonfishes
Large.Benthic.Carnivores.shelf
Large.Benthic.Carnivores.deep
Medium.Benthic.Carnivores.shelf
Medium.Benthic.Carnivores.deep
Small.Benthic.Carnivores.shelf
Small.Benthic.Carnivores.deep
Benthic.omnivores.shelf
Threadfin.breams
Indian.Mackerel
Oil.Sardine
Catch (tonnes)
Year

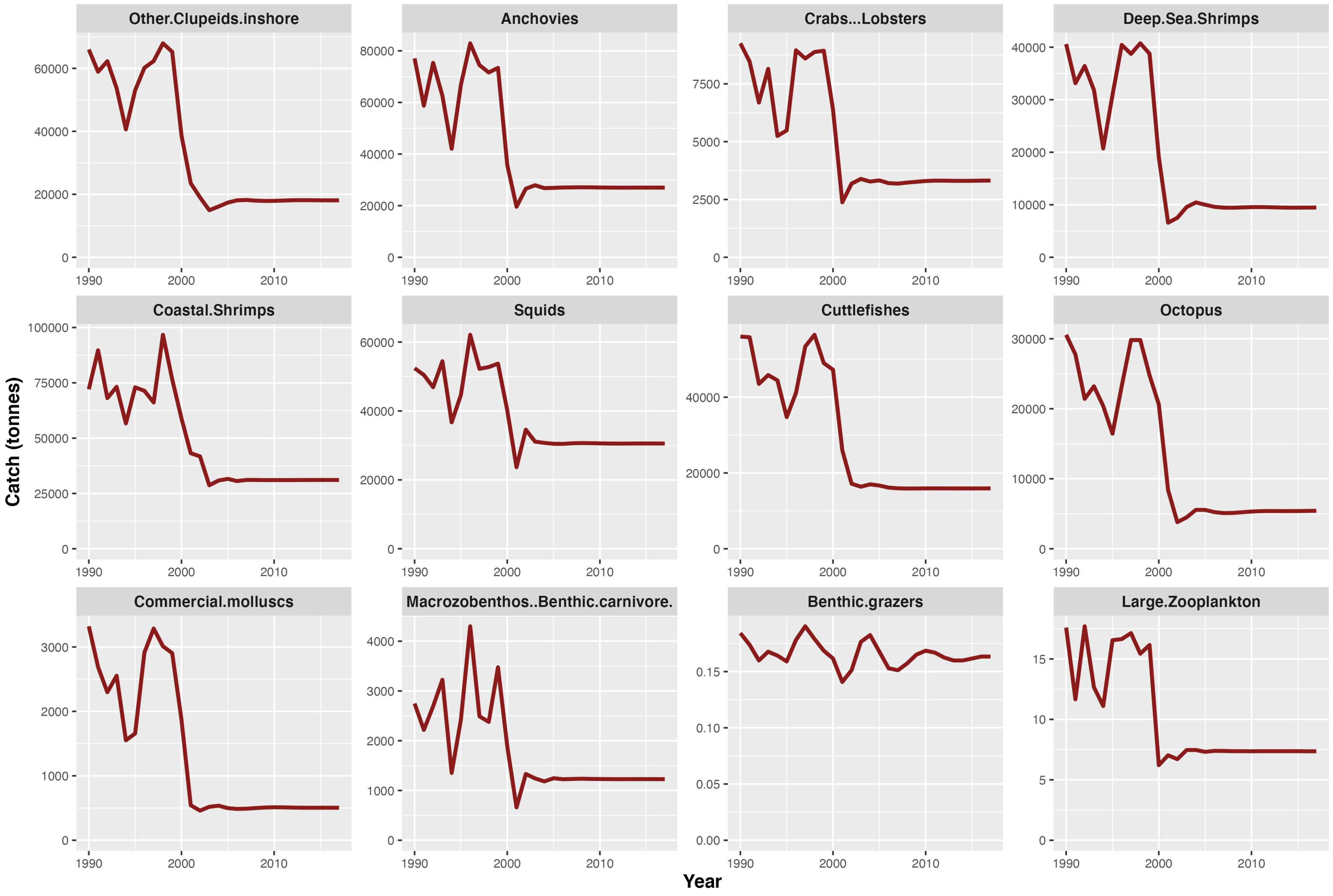
Other.Clupeids.inshore
Anchovies
Crabs...Lobsters
Deep.Sea.Shrimps
Coastal.Shrimps
Squids
Cuttlefishes
Octopus
Commercial.molluscs
Macrozobenthos..Benthic.carnivore.
Benthic.grazers
Large.Zooplankton
Catch (tonnes)
Year

**Figure S5:** Catch time series for the historical baseline run of the north central Chilean EwE model.

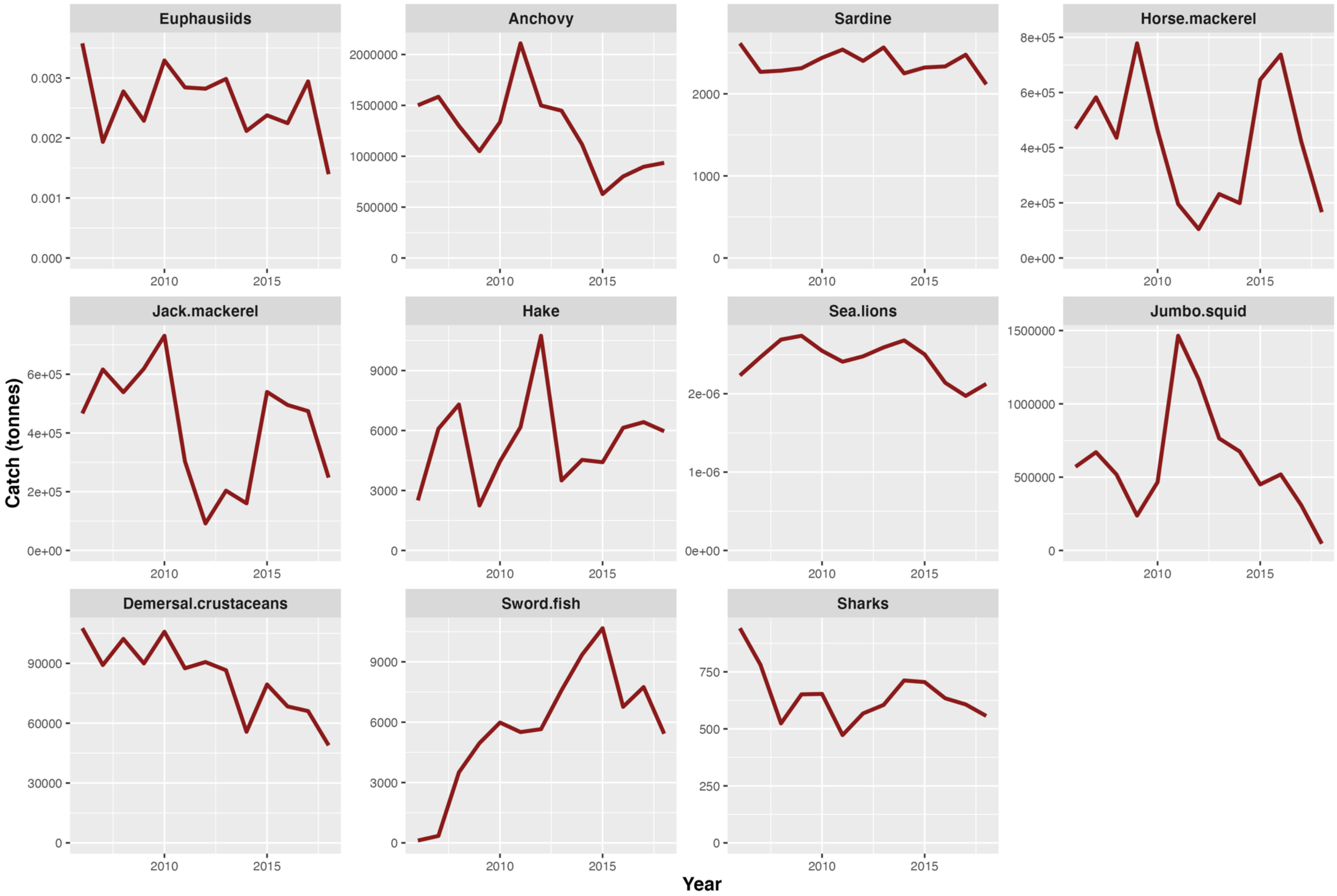
